# Five-leaf generalizations of the *D*-statistic reveal the directionality of admixture

**DOI:** 10.1101/2024.02.24.581856

**Authors:** Kalle Leppälä, Flavio Augusto da Silva Coelho, Michaela Richter, Victor A. Albert, Charlotte Lindqvist

## Abstract

Over the past 15 years, the *D*-statistic, a four-taxon test for organismal admixture (hybridization, or introgression) which incorporates single nucleotide polymorphism data with allelic patterns ABBA and BABA, has seen considerable use. This statistic seeks to discern significant deviation from either a given species tree assumption, or from the balanced incomplete lineage sorting that could otherwise defy this species tree. However, while the *D*-statistic can successfully discriminate admixture from incomplete lineage sorting, it is not a simple matter to determine the directionality of admixture using only four-leaf tree models. As such, methods have been developed that use 5 leaves to evaluate admixture. Among these, the *D*_FOIL_ method, which tests allelic patterns on the “symmetric” tree *S* = (((1, 2), (3, 4)), 5), succeeds in finding admixture direction for many five-taxon examples. However, *D*_FOIL_ does not make full use of all symmetry, nor can *D*_FOIL_ function properly when ancient samples are included because of the reliance on singleton patterns (such as BAAAA and ABAAA). Here, we take inspiration from *D*_FOIL_ to develop a new and completely general family of five-leaf admixture tests, dubbed Δ-statistics, that can either incorporate or exclude the singleton allelic patterns depending on individual taxon and age sampling choices. We describe two new shapes that are also fully testable, namely the “asymmetric” tree *A* = ((((1, 2), 3), 4), 5) and the “quasisymmetric” tree *Q* = (((1, 2), 3), (4, 5)), which can considerably supplement the “symmetric” *S* = (((1, 2), (3, 4)), 5) model used by *D*_FOIL_. We demonstrate the consistency of Δ-statistics under various simulated scenarios, and provide empirical examples using data from black, brown and polar bears, the latter also including two ancient polar bear samples from previous studies. Recently *D*_FOIL_ and one of these ancient samples was used to argue for a dominant polar bear → brown bear introgression direction. However, we find, using both this ancient polar bear and our own, that by far the strongest signal using both *D*_FOIL_ and Δ-statistics on tree *S* is actually bidirectional gene flow of indistinguishable direction. Further experiments on trees *A* and *Q* instead highlight what were likely two phases of admixture: one with stronger brown bear → polar bear introgression in ancient times, and a more recent phase with predominant polar bear → brown bear directionality.

Code and documentation available at https://github.com/KalleLeppala/Delta-statistics.

## 1 Introduction

While phylogenetic tree reconstructions model lineage splits, they do not directly accommodate admixture among populations or species. Demonstrations of introgression among hominin clades (Green et al., 2010; Meyer et al., 2012) were among the first population genomic studies to point out that the evolutionary histories of many organismal groups involve not only lineage bifurcations, but also (sometimes adaptively important) reticulations. Importantly, cross-lineage admixture accommodates the inheritance of traits that might otherwise appear convergently evolved to stem from non-cladistic gene flow instead. Explorations of inter-lineage admixtures at the nucleotide sequence level permit revelation of both the structural and functional impacts of such events. For example, our own research on brown and polar bears has underscored how introgression may have had adaptive consequences following the earlier split of these species into discrete lineages (Lan et al., 2022; Miller et al., 2012). As underscored in that work, alleles introgressed from lower latitude, ecologically generalist brown bear lineages may have influenced exquisitely arctic specific adaptations of polar bears, or vice versa. In this example, it can be appreciated that uncovering the *direction* of admixture events can be of vital importance for understanding both past and future functional impacts of introgression.

Methods developed to reveal introgression from single nucleotide polymorphism data range from likelihood based (for an overview of the multispecies coalescent see (Jiao, Flouri, & Yang, 2021)) to parsimony based. The latter style, comparing allelic state patterns on phylogenetic graphs, may appear mathematically crude compared to a more involved modelling approach, but simple methods such as the *D*-statistic itself that are easily understood, realized and interpreted often enjoy lasting popularity. The classic *D*-statistic (Green et al., 2010; Durand, Patterson, Reich, & Slatkin, 2011; Patterson et al., 2012; related insights already in Huson, Klöpper, Lockhart, & Steel, 2005; Kulathinal, Stevison, & Noor, 2009) leverages symmetry in the phylogenetic tree (((1, 2), 3), 4) to detect evidence of introgression between populations. Allelic patterns BABA (meaning chromosomes from populations 2 and 4 carry allele A while chromosomes from populations 1 and 3 carry allele B at a locus) and ABBA are discordant with the underlying phylogeny, but are still sometimes observed due to incomplete lineage sorting (ILS), gene flow, and convergent nucleotide substitutions. Because ILS does not prefer one pattern over the other, its contribution to

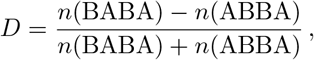

where *n*(**·**) refers to the number of observations, is expected to be zero. Note that the sign is reversed in this re-definition from Patterson et al., 2012 compared to the original Green et al., 2010 to make *D* comparable to the *f*_4_-statistic (Reich, Thangaraj, Patterson, Price, & Singh, 2009); both versions continue to appear in the literature. At a scale where recurrent substitutions can be seen as rare insignificant events, the deviation of *D* from zero therefore points to introgression: between populations 1 and 3 or 2 and 4 (or a ghost population stemming from 123 or 1234) when *D* is positive, and between 2 and 3 or 1 and 4 (or a ghost population stemming from 123 or 1234) when *D* is negative. A key feature of the *D*-statistic is that it requires no assumptions on lengths of the branches of the phylogenetic tree, or mutation and coalescence rates among them, and is therefore suitable for the analysis of ancient samples, with their potentially confounding DNA damage profiles and certain “death date” in the past whereafter mutations can no longer occur as compared to modern samples. This is the reason why the *D*-statistic does not make use of the in principle informative symmetry between the singleton patterns BAAA and ABAA, such as the recently introduced *D*^+^-statistic does (Lopez Fang, Peede, Ortega-Del Vecchyo, McTavish, & Huerta-Sánchez, 2024).

Unfortunately, the *D*-statistic is unable to detect the direction of gene flow. While it is sometimes possible to infer gene flow direction in a sample of only four populations, using *f*_2_-statistic fitting for instance (Leppälä, Nielsen, & Mailund, 2017), the task becomes much less difficult when a fifth population is included in the analysis. This has been a major motivation behind attempts to generalize the classic *D*-statistic to five-leaf trees, such as the Partitioned *D* (Eaton & Ree, 2013) and notably *D*_FOIL_ (Pease & Hahn, 2015) methodologies.

The *D*_FOIL_ method assumes the phylogenetic tree *S* = (((1, 2), (3, 4)), 5), which the authors call *symmetric*, (pictured in Figure 1 Panel I). Counts of allelic patterns are compiled into a *signature*: a 4-tuple (*D*_FO_ *D*_IL_ *D*_FI_ *D*_OL_) of four statistics, each expected to be zero without admixture. When they deviate from zero, the four signs of the signature can be compared with predicted behaviour of various gene flow events; for example, gene flow 1 → 3 should produce the signature (+ + + 0). The authors state that the tree *A* = ((((1, 2), 3), 4), 5), which they label *asymmetric* (pictured in Figure 1 Panel II), does not exhibit enough symmetry for construction of a system of statistics analogous to *D*_FOIL_. However, in our previous work (Lan et al., 2022, Figure 3) we have already leveraged symmetry within the “asymmetric” tree *A*; namely the equal probability between gene trees ((1, 3), (2, 4)), 5) and ((2, 3), (1, 4)), 5) under the null hypothesis of no gene flow. We will demonstrate with proof how allelic pattern counts can also be used to infer gene flow direction in trees of form *A*.

**Figure 1:**
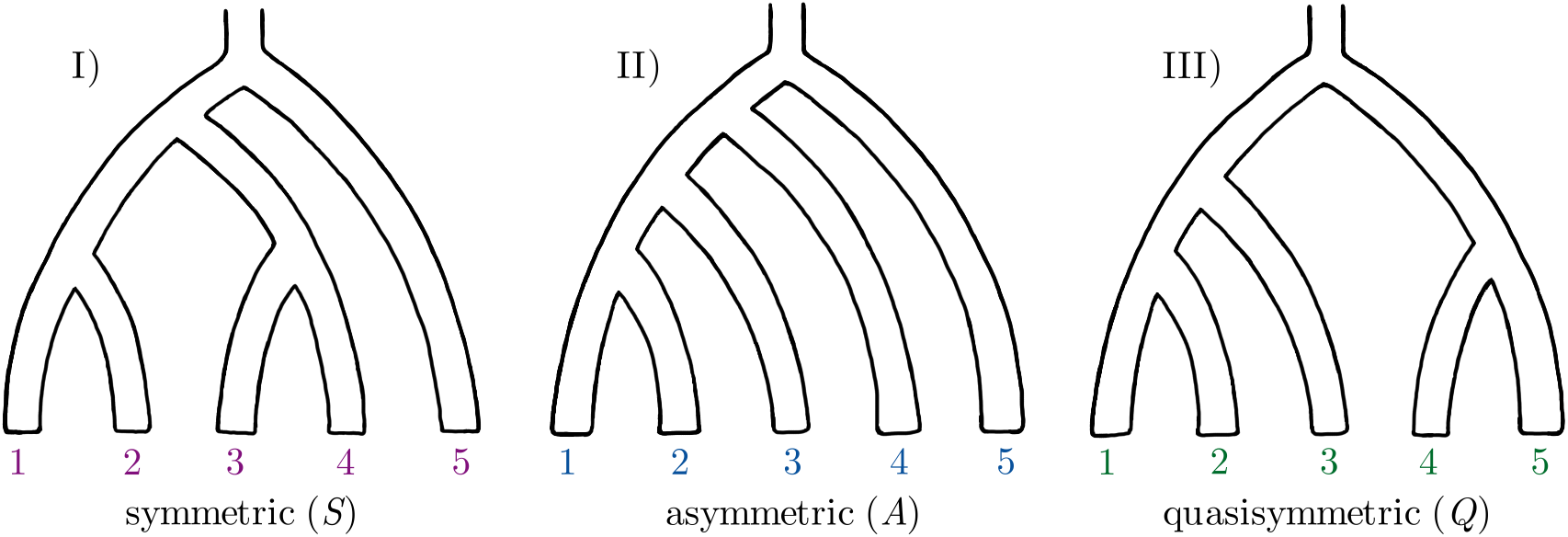
The three possible phylogenetic trees with five leaves.

The current work presents systems of allelic pattern counts that we call Δ-statistics. We consider *S, A*, and a third tree *Q* = (((1, 2), 3), (4, 5)), which we dub *quasisymmetric* (pictured in Figure 1, Panel III)), and make full use of the symmetry within each. Together, *S, A* and *Q* comprise a complete treatise of trees subject to five-leaf relabeling. Some of these Δ-statistics are based on singleton patterns and are therefore not generally applicable to ancient data (for reasons discussed below), but we see the subset of Δ-statistic that doesn’t rely on singleton patterns as the generalization of the classic *D*-statistic for five leaf trees.

Our Δ-statistic method is inspired by the *D*_FOIL_ method, over which it has some advantages:

- Inclusion of trees *A* and *Q*.
- Inclusion of several scenarios within tree *S* that were not considered by Pease and Hahn (2015).
- When *not* using singleton patterns (applicable to ancient data; as with classic *D*), our method provides increased power, while retaining the ability to detect the same set of admixture scenarios within tree *S* as *D*_FOIL_ does, and additionally gene flow into population 5.
- When resorting to singleton patterns (generally not applicable to ancient data, as with *D*_FOIL_), it is able to distinguish more admixture scenarios within tree *S* than what *D*_FOIL_ permits.

We test the Δ-statistics with simulations, and apply the method to further analyse the brown bear – polar bear introgression problem.

## 2 Methods

### Definition of Δ-statistics

The *unscaled* Δ-statistics have the form *n*(*L*) − *n*(*R*), where *L* and *R* (“left” and “right”) are two disjoint sets of allelic patterns expected to be observed with equal probability under the null hypothesis of no gene flow, ignoring the possibility of reverse and convergent substitutions. The *scaled* Δ-statistic, decorated with a star, is then (*n*(*L*) − *n*(*R*))/(*n*(*L*) + *n*(*R*)). In this general framework, *D*- and *D*_FOIL_-statistics are special cases of scaled Δ-statistics.

Because tree *Q* does not have an outgroup, and because we wish to permit admixture to involve the root or population 5 in trees *S* and *A*, we make the *outgroup mutation assumption*. That is, we adopt a notation where allele A is the one observed more often (at least three times out of five) than the other allele B, while remaining agnostic about the ancestral state.

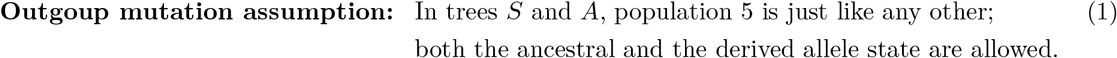

In contrast, the *D*- and the *D*_FOIL_-statistics assume that the ancestral states are known and that the outgroup always carries the ancestral allele, be it A or B. That is an unnecessary complication (Pease and Hahn also write “not strictly necessary”) and in fact the allele frequency-based formulae of the *D*- and *D*_FOIL_-statistics, as presented in Harris and DeGiorgio (2017), implicitly make the outgroup mutation assumption (1) as well. In our case, a data preprocessing step of filtering out the patterns where 5 carries the derived allele would not only be unnecessary but actively harmful, although it would only affect some of the Δ-statistics, and only when there is gene flow involving the (would-be) outgroup 5. See the end of Appendix 2 for more detail. Consider tree *S* without gene flow events, and the coalescent process connecting lineages 1–5 at some locus. If we observe the allelic pattern AAABB, caused by a single mutation, then one of two events must have occurred:

- Lineages 4 and 5 first coalesced with each other (first as in closest to the present; in the context of coalescence processes we consider time running backwards). A subsequent mutation occurred before coalescence with any other lineages, only affecting 4 and 5. Allele A is now ancestral. An example of this scenario is depicted in Figure 2 Panel I), where the black box is replaced with the upper of the two bracketed options.
- Lineages 1, 2 and 3 all coalesced with each other in some order before coalescence with lineages 4 or 5. A subsequent mutation occurred, only affecting 1 and 2 and 3. Allele B is now ancestral. An example of this scenario is depicted in Figure 2 Panel II), where the black box is with the upper of the two bracketed options.

**Figure 2:**
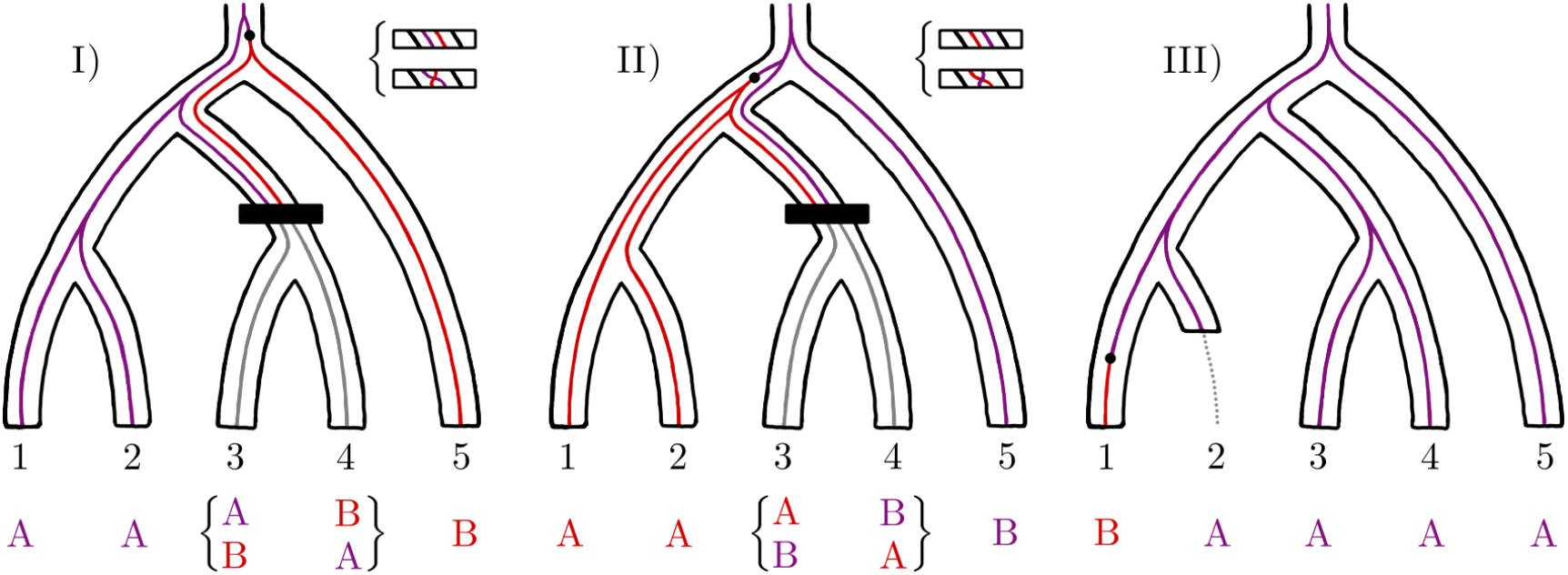
The pattern AAABB results either from a mutation affecting leaves 4 and 5 as in Panel I), or a mutation affecting leaves 1, 2 and 3 as in Panel II). The letter B always denotes the rarer allele, not necessarily the derived allele (red). Symmetry between lineages 3 and 4 is represented as the two black boxes, each hiding two equally likely events that visualize how the probability of pattern AABAB must be the same as the probability of AAABB. We call {AABAB, AAABB} an equal probability set. In Panel III) {BAAAA, ABAAA} is not an equal probability set because the synchronization assumption (2) is violated. Here, 2 is an ancient sample that was long dead when a mutation occurred in lineage 1.

**Figure 3:**
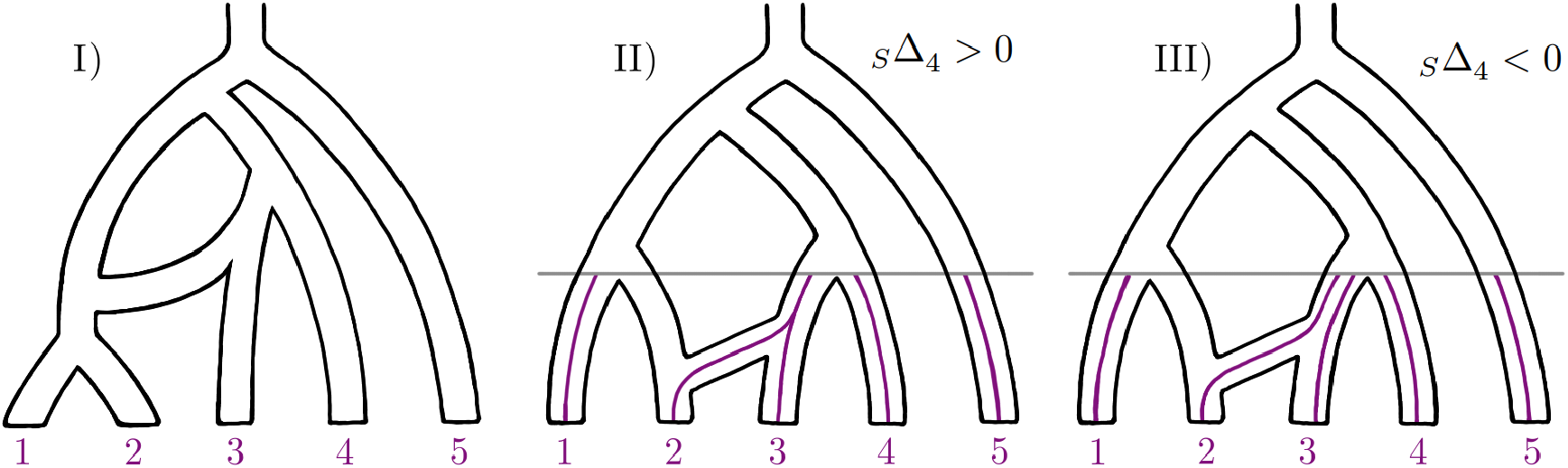
Panel I) depicts the admixture graph meant by 3 → 12. Panels II) and III) demonstrate inconsistency of _*S*_Δ_4_ under the gene flow scenario 3 → 2. In Panel II), 2 and 3 coalesce before meeting 4, and so _*S*_Δ_4_ *>* 0. In Panel III), 2 and 3 do not coalesce before meeting 4, and so _*S*_Δ_4_ *<* 0.

In both scenarios, as soon as lineages 3 and 4 are in the same population, their role is symmetrical concerning coalescent history — what happens to 3 could just as well have happened to 4. This is visualized in Figure 2 Panels I) and II), with the black boxes representing two equally likely options, upper and lower. In both scenarios above, the lower option would have resulted in pattern AABAB, which therefore has exactly the same probability as AAABB. The two allelic patterns form an *equal probability set* {AABAB, AAABB} for tree *S*; we do not know the probability of observing either of these patterns, but we know they are the same by symmetry. Note that if we filter the data such that lineage 5 always carries the ancestral allele, giving up the outgroup mutation assumption (1), we simply censor the latter of the two scenarios listed above, and the set is still an equal probability set.

Symmetry between singleton patterns depends on mutations occurring with the same probability in different populations. The patterns BAAAA and ABAAA each have the same probability if the mutation occurred after populations 1 and 2 merged. However, if the mutation occurred in one of the terminal branches, there are many potential reasons for asymmetry, such as different generation times, mutation rates, and in the case of ancient samples, DNA degradation or simply fewer generations. In Figure 2 Panel III) we have depicted such a situation. In the following, equal probability sets of singleton patterns, such as the set {BAAAA, ABAAA}, are burdened with the *synchronization assumption*.

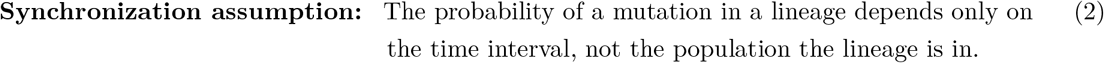

Notice that this assumption concerns the entire species tree, not only the terminal branches. We also point out that, theoretically, even ancient samples can satisfy the synchronization assumption when mutations on the terminal branches are so rare that they do not influence the analysis (see simulation in Appendix 5). In fact, methods using the perspective of drift instead of the perspective of coalescence, such as *f*_2_-, *f*_3_- and *f*_4_-statistics (Reich et al., 2009; Patterson et al., 2012), typically rely on a simplification that variants are already polymorphic at the root, which is simply a special case of the synchronization assumption. If an investigator cannot commit to the synchronization assumption using their data, they can simply resort to the subset of Δ-statistics constructed from equal probability sets that do not depend on it.

Given an equal probability set, we pick one element for *L* and one for *R* to construct a *binomial* Δ-statistic *n*(*L*) − *n*(*R*). The binomial Δ-statistics are random variables that have mean zero under the null hypothesis of no admixture. Which element goes to *L* and which goes to *R* is arbitrary; our choices and the full collection of equal probability sets for trees *S, A* and *Q* can be found in Table 1. Even though 4 “choose” 2 is 4!/(2! × 2!) = 6, only (any) three differences of patterns from the four-element equal probability sets in Table 1 are linearly independent. In those cases we still constructed four binomial Δ-statistics, the remaining linear relation leaving the notational ambiguity _*S*_Δ_1_ + _*S*_Δ_4_ = _*S*_Δ_2_ + _*S*_Δ_3_ and _*Q*_Δ_1_ + _*Q*_Δ_4_ = _*Q*_Δ_2_ + _*Q*_Δ_3_. The missing two binomial Δ-statistics in both instances can be expressed with the four binomial Δ-statistics already defined. For example, in tree *S*:

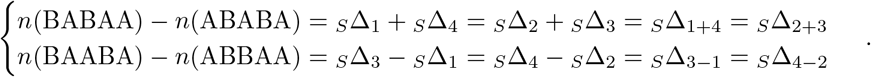

**Table 1:** Equal probability sets and binomial Δ-statistics constructed from them. Rows highlighted grey require the synchronization assumption (2).

| tree | equal probability set | binomial Δ-statistics |
| --- | --- | --- |
| $S$ | $\{BABAA, BAABA, ABBAA, ABABA\}$ | $s\Delta_1 = n(BABAA) - n(BAABA)$<br>$s\Delta_2 = n(ABBAA) - n(ABABA)$<br>$s\Delta_3 = n(BABAA) - n(ABBAA)$<br>$s\Delta_4 = n(BAABA) - n(ABABA)$ |
| | $\{BAAAB, ABAAB\}$ | $s\Delta_5 = n(BAAAB) - n(ABAAB)$ |
| | $\{AABAB, AAABB\}$ | $s\Delta_6 = n(AABAB) - n(AAABB)$ |
| | $\{BAAAA, ABAAA\}$ | $s\Delta_7 = n(BAAAA) - n(ABAAA)$ |
| | $\{AABAA, AAABA\}$ | $s\Delta_8 = n(AABAA) - n(AAABA)$ |
| $A$ | $\{BABAA, ABBAA\}$ | $A\Delta_1 = n(BABAA) - n(ABBAA)$ |
| | $\{BAABA, ABABA\}$ | $A\Delta_2 = n(BAABA) - n(ABABA)$ |
| | $\{BAAAB, ABAAB\}$ | $A\Delta_3 = n(BAAAB) - n(ABAAB)$ |
| | $\{BAAAA, ABAAA\}$ | $A\Delta_4 = n(BAAAA) - n(ABAAA)$ |
| $Q$ | $\{BAABA, BAAAB, ABABA, ABAAB\}$ | $Q\Delta_1 = n(BAABA) - n(BAAAB)$<br>$Q\Delta_2 = n(ABABA) - n(ABAAB)$<br>$Q\Delta_3 = n(BAABA) - n(ABABA)$<br>$Q\Delta_4 = n(BAAAB) - n(ABAAB)$ |
| | $\{BABAA, ABBAA\}$ | $Q\Delta_5 = n(BABAA) - n(ABBAA)$ |
| | $\{AABBA, AABAB\}$ | $Q\Delta_6 = n(AABBA) - n(AABAB)$ |
| | $\{BAAAA, ABAAA\}$ | $Q\Delta_7 = n(BAAAA) - n(ABAAA)$ |
| | $\{AAABA, AAAAB\}$ | $Q\Delta_8 = n(AAABA) - n(AAAAB)$ |

This technique and notation can be expanded to construct a large number of Δ-statistics as sums and differences of binomial Δ-statistics, all of which are expected to be zero by the linearity of the mean. For the purpose of scaling and statistical inference, we must understand the sets *L* and *R* as disjoint. That is, for _*S*_Δ_1+4_ we have *L* = {BABAA} ≠ {BABAA, BAABA} and *R* = {ABABA} ≠ {BAABA, ABABA}, because BAABA cannot exist in both sets.

This large family of Δ-statistics, combined from the binomial Δ-statistics, includes as special cases the *D*_FOIL_-statistics (Pease & Hahn, 2015) and the Partitioned *D* (Eaton & Ree, 2013), which can be expressed as:

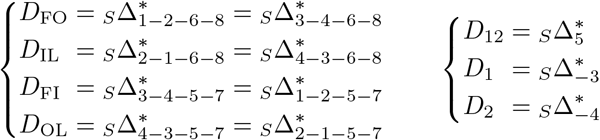

The star stands for the scaling (*n*(*L*) − *n*(*R*))/(*n*(*L*) + *n*(*R*)). There also exist some informative Δ-statistics such as *n*(BAABA) + *n*(ABABA) + 2*n*(AABAB) − *n*(BAAAB) − *n*(ABAAB) − 2*n*(AABBA) in tree *A* that cannot be constructed from the binomial statistics.

We are now prepared to present our preferred twenty Δ-statistics. For the symmetric tree, *S*, we define

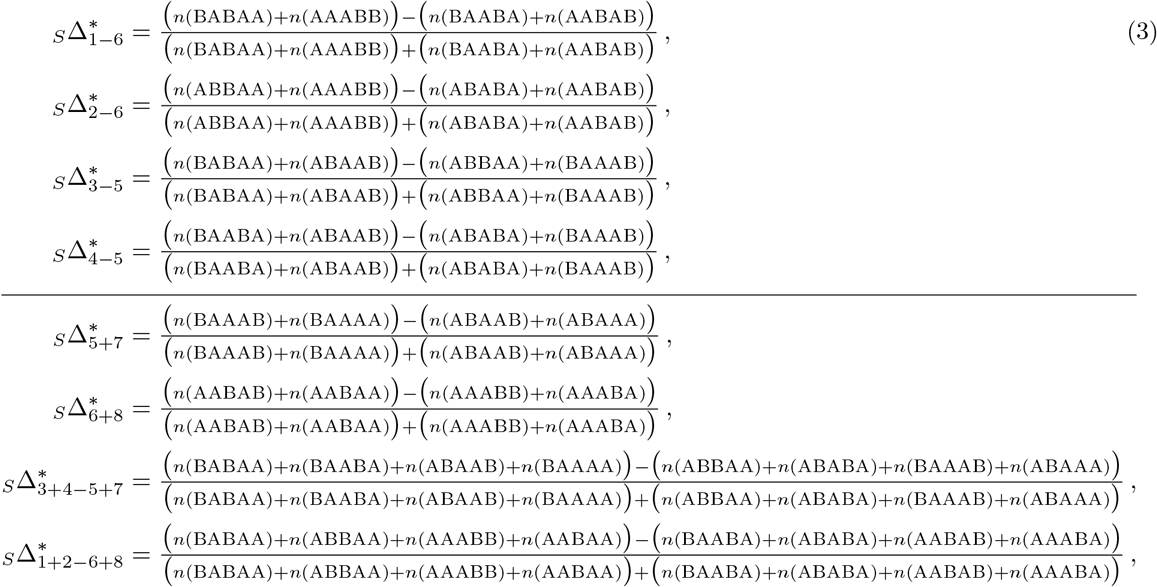

for the asymmetric tree, *A*, we define

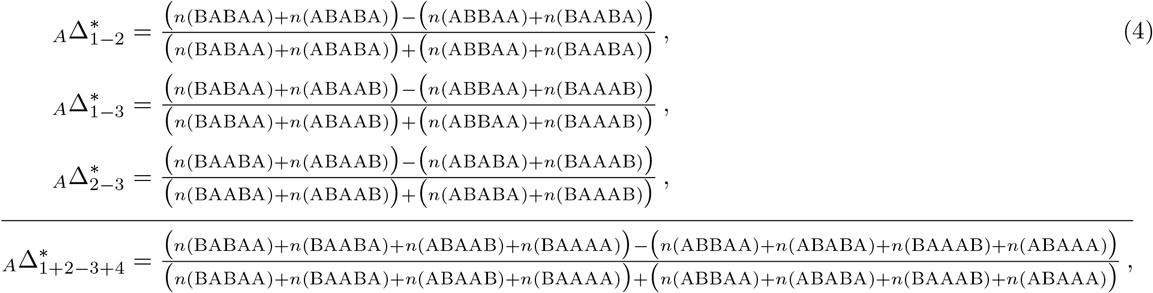

and for the quasisymmetric tree, *Q*, we define

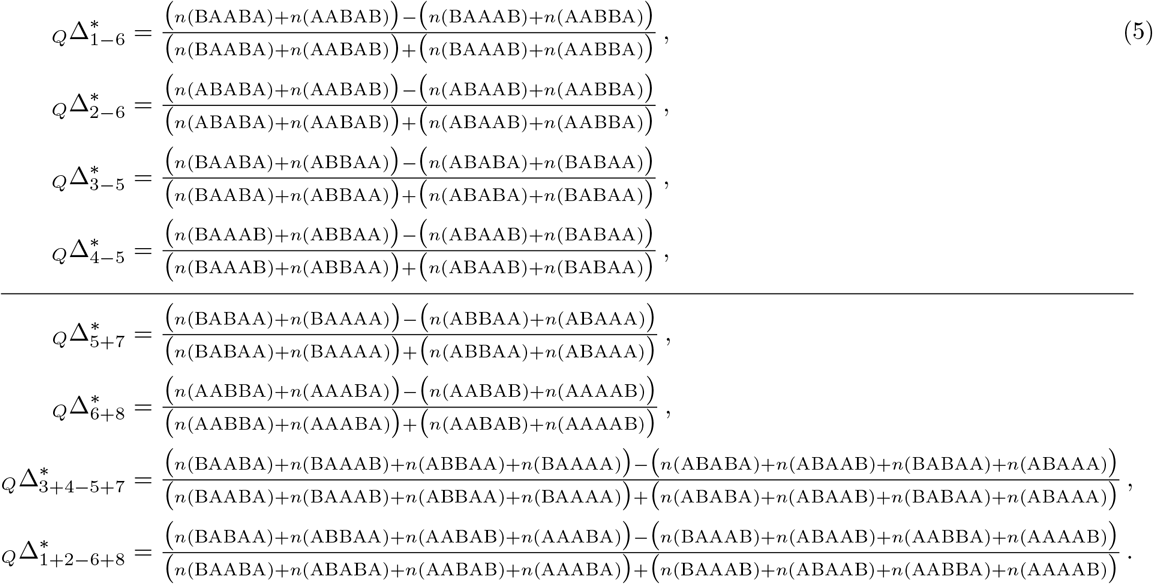

The horizontal lines remind us that the binomial statistics _*S*_Δ_7, *S*_Δ_8, *A*_Δ_4, *Q*_Δ_7_ and _*Q*_Δ_8_ require the synchronization assumption (2). See Appendix 1 for allele frequency -based formulations of (3), (4) and (5). For comparison, the classic *D*-statistic does not need to make the synchronization assumption (2) and the *D*_FOIL_-statistics do.

### Classification of expected Δ-statistics

The types of alternative hypotheses considered in this work are *unidirectional gene flow events*. The species tree becomes equipped with a new branch, making it an admixture graph. We denote, for example, 3 → 12, to indicate that the new branch stems from the terminal branch 3 and then admixes into the branch that is ancestral to populations 1 and 2, as shown in Figure 3 Panel I). From the coalescence point of view, lineages reaching the admixture node have some positive probability of entering the new branch instead. The new branch carrying the gene flow is permitted to be of any length within the constraints of the tree; 1234 → 1 in *S* cannot have zero length, for example. There is reason to think that introgression may typically be mediated by some unsampled *ghost population* (Tricou, Tannier, & de Vienne, 2022), hence we do not assume that the length of the admixture branch is zero even when it could be. The bidirectional arrow 3 ↔ 12 is used as a shorthand notation that means either 3 → 12 or 12 → 3 (although truly bidirectional gene flow will behave similarly whenever the symbol ↔ is invoked).

One difficulty is ensuring that the expected behaviour of Δ-statistics is *consistent* under admixture events of interest. Consider gene flow 3 → 2 in tree *S*. If lineage 2 is introgressed and coalesces with lineage 3 before the populations 3 and 4 merge (Figure 3 Panel II)), then we expect _*S*_Δ_4_ to be positive as the pattern ABABA cannot be observed at all. On the other hand, if lineage 2 is introgressed but does not coalesce with lineage 3 by the time populations 3 and 4 merge (Figure 3 Panel III)), pattern ABABA becomes more likely than pattern BAABA (see Lemma 6 in Appendix 2) and _*S*_Δ_4_ is expected to be negative. This is an example of inconsistency, where the expected sign depends on graph parameters (the probability of coalescence within the terminal branch 3) that we are committed to making no assumptions about. The twenty Δ-statistics (3), (4) and (5) highlighted in the previous section are chosen in such a way that the expected behaviour is always consistent and the ability to identify different gene flow events is maximized within the framework of equal probability sets and binomial statistics, while avoiding the use of singleton patterns to the extent it is possible.

The expected classifications of Δ-statistics (3), (4) and (5) under every unidirectional gene flow event on trees *S, A* and *Q* are collected in Tables 2, 3 and 4. The entry + (resp. −) signifies that the expectation is some unknown positive (resp. negative) number, and the entry 0 means it is zero. The proof can be found in Appendix 2. If a gene flow event is not listed in the table (and there are many), the expected classification of each Δ-statistic is then 0. The vertical dividers indicate that the right sides of the tables depend on the synchronization assumption (2); see Appendix 3 for collapsed tables that only use the left sides, that is, the Δ-statistics that are independent of (2). Apart from gene flow 5 → *x* and *x* → 5, *x* ∈ {1, 2, 3, 4} in tree *S* and *x* 5, *x* 1, 2 in tree *A*, the Tables 2 and 3 are also valid if the data have been filtered such that lineage 5 always carries the ancestral allele, contrary to the outgroup mutation assumption (1). For this fact we will only provide a sketch proof at the end of Appendix 2.

**Table 2:** Predicted effects on tree *S* = (((1, 2), (3, 4)), 5). The last four columns assume (2). Events 5 → *x* and *x*→ 5, where *x* ∈ {1, 2, 3, 4}, assume (1). Unidirectional gene flow events not listed in the table produce a zero signature.

| | $S\Delta_{1-6}^*$ | $S\Delta_{2-6}^*$ | $S\Delta_{3-5}^*$ | $S\Delta_{4-5}^*$ | $S\Delta_{5+7}^*$ | $S\Delta_{6+8}^*$ | $S\Delta_{3+4-5+7}^*$ | $S\Delta_{1+2-6+8}^*$ |
| --- | --- | --- | --- | --- | --- | --- | --- | --- |
| $1 \rightarrow 3$ | + | + | + | 0 | − | − | 0 | 0 |
| $3 \rightarrow 1$ | + | 0 | + | + | − | − | 0 | 0 |
| $1 \rightarrow 4$ | − | − | 0 | + | − | + | 0 | 0 |
| $4 \rightarrow 1$ | − | 0 | + | + | − | + | 0 | 0 |
| $2 \rightarrow 3$ | + | + | − | 0 | + | − | 0 | 0 |
| $3 \rightarrow 2$ | 0 | + | − | − | + | − | 0 | 0 |
| $2 \rightarrow 4$ | − | − | 0 | − | + | + | 0 | 0 |
| $4 \rightarrow 2$ | 0 | − | − | − | + | + | 0 | 0 |
| $12345 \rightarrow 1$ | 0 | 0 | − | − | + | 0 | + | 0 |
| $34 \leftrightarrow 2$<br>$1234 \rightarrow 1$ | 0 | 0 | − | − | + | 0 | 0 | 0 |
| $5 \rightarrow 1$ | 0 | 0 | − | − | + | 0 | − | 0 |
| $1 \rightarrow 5$ | 0 | 0 | − | − | 0 | 0 | − | 0 |
| $12345 \rightarrow 2$ | 0 | 0 | + | + | − | 0 | − | 0 |
| $34 \leftrightarrow 1$<br>$1234 \rightarrow 2$ | 0 | 0 | + | + | − | 0 | 0 | 0 |
| $5 \rightarrow 2$ | 0 | 0 | + | + | − | 0 | + | 0 |
| $2 \rightarrow 5$ | 0 | 0 | + | + | 0 | 0 | + | 0 |
| $12345 \rightarrow 3$ | − | − | 0 | 0 | 0 | + | 0 | + |
| $12 \leftrightarrow 4$<br>$1234 \rightarrow 3$ | − | − | 0 | 0 | 0 | + | 0 | 0 |
| $5 \rightarrow 3$ | − | − | 0 | 0 | 0 | + | 0 | − |
| $3 \rightarrow 5$ | − | − | 0 | 0 | 0 | 0 | 0 | − |
| $12345 \rightarrow 4$ | + | + | 0 | 0 | 0 | − | 0 | − |
| $12 \leftrightarrow 3$<br>$1234 \rightarrow 4$ | + | + | 0 | 0 | 0 | − | 0 | 0 |
| $5 \rightarrow 4$ | + | + | 0 | 0 | 0 | − | 0 | + |
| $4 \rightarrow 5$ | + | + | 0 | 0 | 0 | 0 | 0 | + |

**Table 3:**
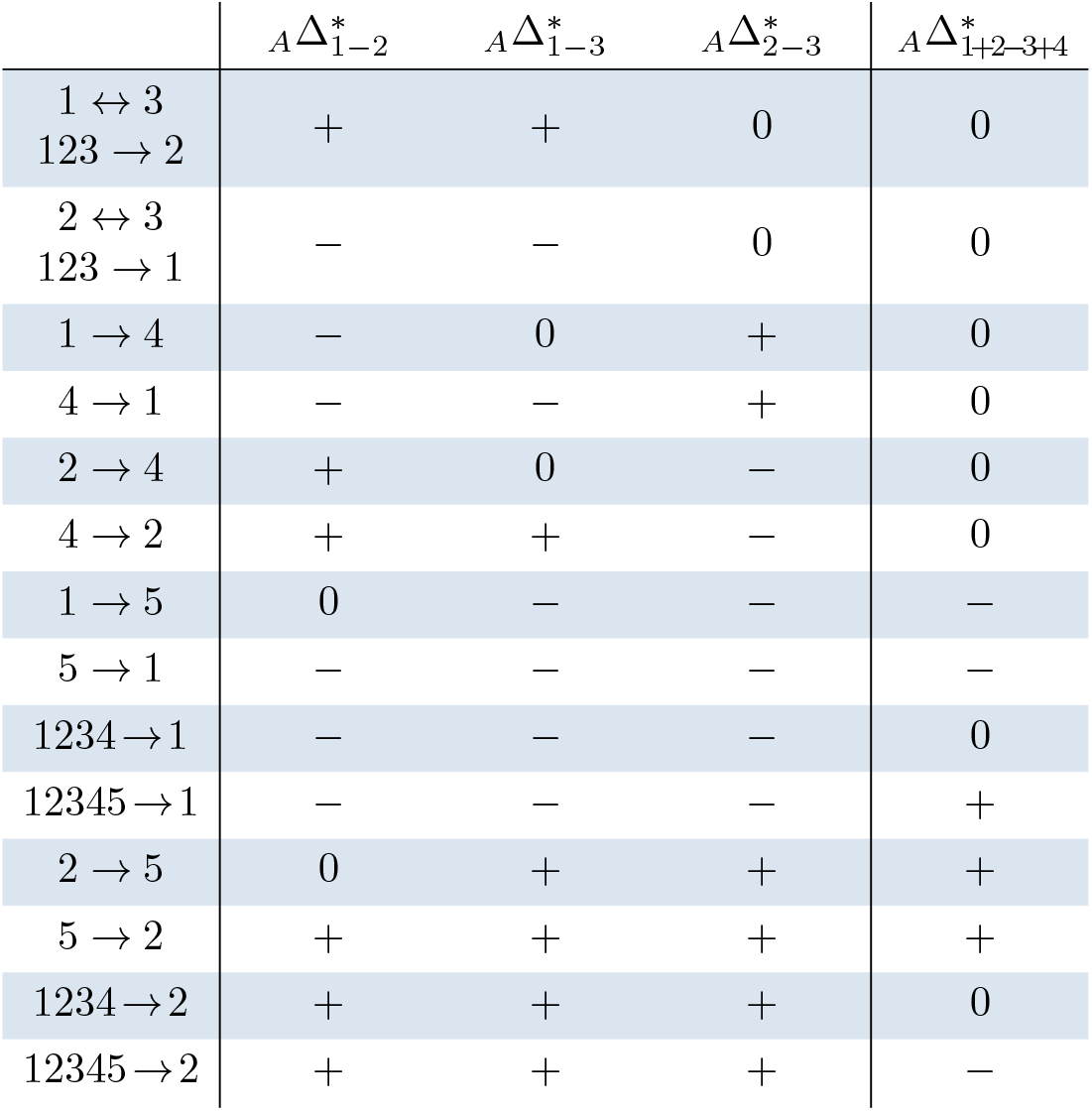
Predicted effects on tree *A* = ((((1, 2), 3), 4), 5). The last column assumes (2). Events 5 → 1 and 5 → 2 assume (1). Unidirectional gene flow events not listed in the table produce a zero signature.

| | ${}^A\Delta_{1-2}^*$ | ${}^A\Delta_{1-3}^*$ | ${}^A\Delta_{2-3}^*$ | ${}^A\Delta_{1+2-3+4}^*$ |
| --- | --- | --- | --- | --- |
| $1 \leftrightarrow 3$<br>$123 \rightarrow 2$ | + | + | 0 | 0 |
| $2 \leftrightarrow 3$<br>$123 \rightarrow 1$ | − | − | 0 | 0 |
| $1 \rightarrow 4$ | − | 0 | + | 0 |
| $4 \rightarrow 1$ | − | − | + | 0 |
| $2 \rightarrow 4$ | + | 0 | − | 0 |
| $4 \rightarrow 2$ | + | + | − | 0 |
| $1 \rightarrow 5$ | 0 | − | − | − |
| $5 \rightarrow 1$ | − | − | − | − |
| $1234 \rightarrow 1$ | − | − | − | 0 |
| $12345 \rightarrow 1$ | − | − | − | + |
| $2 \rightarrow 5$ | 0 | + | + | + |
| $5 \rightarrow 2$ | + | + | + | + |
| $1234 \rightarrow 2$ | + | + | + | 0 |
| $12345 \rightarrow 2$ | + | + | + | − |

**Table 4:** Predicted effects on tree *Q* = (((1, 2), 3), (4, 5)). The last four columns assume (2). Unidirectional gene flow events not listed in the table produce a zero signature.

| | $Q\Delta_{1-6}^*$ | $Q\Delta_{2-6}^*$ | $Q\Delta_{3-5}^*$ | $Q\Delta_{4-5}^*$ | $Q\Delta_{5+7}^*$ | $Q\Delta_{6+8}^*$ | $Q\Delta_{3+4-5+7}^*$ | $Q\Delta_{1+2-6+8}^*$ |
| --- | --- | --- | --- | --- | --- | --- | --- | --- |
| $1 \rightarrow 4$ | + | + | + | 0 | − | − | 0 | − |
| $4 \rightarrow 1$ | + | 0 | + | + | − | − | + | 0 |
| $1 \rightarrow 5$ | − | − | 0 | + | − | + | 0 | + |
| $5 \rightarrow 1$ | − | 0 | + | + | − | + | + | 0 |
| $2 \rightarrow 4$ | + | + | − | 0 | + | − | 0 | − |
| $4 \rightarrow 2$ | 0 | + | − | − | + | − | − | 0 |
| $2 \rightarrow 5$ | − | − | 0 | − | + | + | 0 | + |
| $5 \rightarrow 2$ | 0 | − | − | − | + | + | − | 0 |
| $2 \rightarrow 45$ | 0 | 0 | − | − | + | 0 | 0 | 0 |
| $45 \rightarrow 2$ | 0 | 0 | − | − | + | 0 | − | 0 |
| $1 \leftrightarrow 3$<br>$123 \rightarrow 2$ | 0 | 0 | − | − | 0 | 0 | − | 0 |
| $12345 \rightarrow 2$ | 0 | 0 | − | − | − | 0 | − | 0 |
| $1 \rightarrow 45$ | 0 | 0 | + | + | − | 0 | 0 | 0 |
| $45 \rightarrow 1$ | 0 | 0 | + | + | − | 0 | + | 0 |
| $2 \leftrightarrow 3$<br>$123 \rightarrow 1$ | 0 | 0 | + | + | 0 | 0 | + | 0 |
| $12345 \rightarrow 1$ | 0 | 0 | + | + | + | 0 | + | 0 |
| $5 \rightarrow 12$ | − | − | 0 | 0 | 0 | + | 0 | 0 |
| $12 \rightarrow 5$ | − | − | 0 | 0 | 0 | + | 0 | + |
| $4 \rightarrow 3$ | − | − | 0 | 0 | 0 | 0 | 0 | − |
| $3 \rightarrow 4$ | − | − | 0 | 0 | 0 | − | 0 | − |
| $4 \rightarrow 12$ | + | + | 0 | 0 | 0 | − | 0 | 0 |
| $12 \rightarrow 4$ | + | + | 0 | 0 | 0 | − | 0 | − |
| $5 \rightarrow 3$ | + | + | 0 | 0 | 0 | 0 | 0 | + |
| $3 \rightarrow 5$ | + | + | 0 | 0 | 0 | + | 0 | + |
| $123 \leftrightarrow 5$<br>$12345 \rightarrow 4$ | 0 | 0 | 0 | 0 | 0 | + | 0 | + |
| $123 \leftrightarrow 4$<br>$12345 \rightarrow 5$ | 0 | 0 | 0 | 0 | 0 | − | 0 | − |

### Classification of observed Δ-statistics

With the null hypothesis of no admixture events, and assuming independent loci, the number of observations in *L* follows the binomial distribution *n*(*L*) ∼ *B*(*n*(*L*) + *n*(*R*), 0.5). Using normal approximation, the number

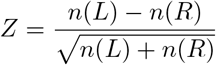

has nearly the standard Gaussian distribution. Let *α* be the maximum tolerated probability of classifications − or + when the null hypothesis holds (type I error). We classify the observed statistic as

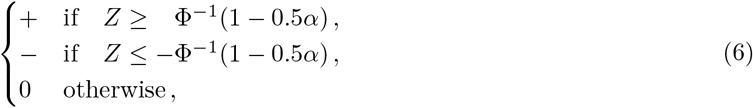

and as a convention, the case *n*(*L*) = *n*(*R*) = 0 as 0. Following *D*_FOIL_ (Pease & Hahn, 2015), we use the value *α* = 0.01 throughout this paper. A stricter value, *α* = 0.0027, could also be considered because the cutoff *Z* = ±3 is essentially a community standard for the classic *D*-statistic (Green et al., 2010). In practice, the assumption of independent loci is invalid and linkage disequilibrium will inflate the *Z*-scores in absolute value. The severity and solution of this issue are context dependent. For the *D*-statistic, a typical way to alleviate the effects of linkage is the block jackknife procedure (Busing, Meijer, & Leeden, 1999).

### Local ancestry

Introgression happens on the level of complete chromosomes, consequently broken down by recombination. By zooming into smaller windows we can prevent multiple different gene flow events from influencing an analysis. For example, bidirectional gene flow event 2↔3 in tree *S* produces the signal (+ + − − |+ − 0 0) by Table 2, rows 5 and 6. But so would the mixture of 34→ 2 and 12 →3 by rows 10 and 22. These two scenarios can be told apart by studying the local ancestry; in the first case we expect some windows with signature (+ + − 0 |+ − 0 0) and some with signature (0 + − − |+ − 0 0), whereas in the latter case we expect signatures (0 0− − |+ 0 0 0) and (+ + 0 0 | 0 − 0 0). Because the alternative hypotheses we consider are unidirectional gene flow events, the size of windows to zoom into should be so small that it is reasonable to suppose each window is affected by at most one of the events. We dub this requirement the *single event assumption*.

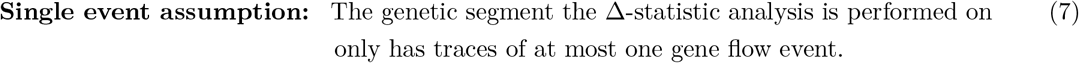

Shorter windows are more likely to satisfy the single event assumption (7), but eventually face issues with stochasticity and statistical power. The effect of linkage disequilibrium is more difficult to remove on shorter windows.

## 3 Results

### Admixture simulations

We simulated gene trees in a coalescence process, where the time until coalescence between two given lineages in the same randomly mating population follows an exponential distribution with parameter *λ*. A unit of time is then approximately 2*λN*_*e*_ generations, where *N*_*e*_ is the effective population size. Number of mutations on a (mutation rate weighted) simulated gene tree is then sampled from a Poisson distribution and the location of mutations within it from a uniform distribution, resulting in biallelic patterns. Compared to simulating genetic regions under recombination, the simple coalescent approach is a compromise because it does not inform us on the window size necessary to infer gene flow events.

We tested all unidirectional gene flow scenarios that have non-zero expected signatures in trees *S, A* and *Q*, for a total of 32, 18 and 34 simulations, respectively. The number of patterns simulated for each scenario was 1,000,000 (presented here), 100,000 or 10,000 (presented in Appendix 4). We used *λ* = 1, with the last common ancestor of all five populations living at time point 3 (for example, if generation time is 10 years and *N*_*e*_ = 10, 000, this translates into 600,000 years). For mutations, the Poisson parameter per unit of time was set to 0.0001 (corresponding to per-year-per-locus mutation rate of 0.5 × 10^−9^ in the example given above). The other nodes of the species tree were placed roughly evenly, and the admixture events were instantaneous when possible and mediated by a ghost population otherwise, with admixture proportion at 10%. For details, see documentation on our GitHub page (https://github.com/KalleLeppala/Delta-statistics).

In all but one scenarios with 1,000,000 patterns, the observed Δ-statistics matched the predictions in Tables 2, 3 and 4 (results in Tables 5, 6 and 7). With 100,000 independent samples, the Δ-statistics depending on the synchronization assumption (2) became unreliable (Appendix 4), while the left hand sides of the Tables 2, 3 and 4 (collapsed versions can be found in Appendix 3) remained accurate. With only 10,000 independent samples, the Δ-statistics no longer functioned well (Appendix 4).

**Table 5:** Simulation results for the symmetric tree *S*. For each statistic under each scenario we report the value, the 99% Wald confidence interval with radius 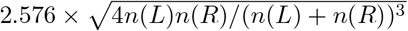, the *Z*-score and the classification. All results match the predictions in Table 2 except for the signature of 3 → 12 where (dependent) statistics _*S*_Δ_3 5_ and _*S*_Δ_4 5_ (highlighted with gray) were supposed to be zero.

| | $s\Delta_{1-6}^*$ | $s\Delta_{2-6}^*$ | $s\Delta_{3-5}^*$ | $s\Delta_{4-5}^*$ | $s\Delta_{5+7}^*$ | $s\Delta_{6+8}^*$ | $s\Delta_{3+4-5+7}^*$ | $s\Delta_{1+2-6+8}^*$ |
| --- | --- | --- | --- | --- | --- | --- | --- | --- |
| $1 \rightarrow 3$ | $0.388 \pm 0.015$<br>63.44 : + | $0.290 \pm 0.016$<br>44.82 : + | $0.157 \pm 0.018$<br>23.07 : + | $-0.003 \pm 0.020$<br>-0.406 : 0 | $-0.011 \pm 0.006$<br>-5.766 : - | $-0.058 \pm 0.005$<br>-30.48 : - | $0.001 \pm 0.005$<br>0.739 : 0 | $0.005 \pm 0.005$<br>2.507 : 0 |
| $3 \rightarrow 1$ | $0.163 \pm 0.018$<br>23.89 : + | $0.004 \pm 0.020$<br>0.547 : 0 | $0.391 \pm 0.015$<br>63.62 : + | $0.292 \pm 0.016$<br>44.94 : + | $-0.063 \pm 0.005$<br>-33.12 : - | $-0.013 \pm 0.006$<br>-6.731 : - | $-0.000 \pm 0.005$<br>-0.127 : 0 | $0.001 \pm 0.005$<br>0.266 : 0 |
| $1 \rightarrow 4$ | $-0.388 \pm 0.015$<br>-62.86 : - | $-0.286 \pm 0.017$<br>-43.90 : - | $-0.001 \pm 0.020$<br>-0.085 : 0 | $0.160 \pm 0.018$<br>23.46 : + | $-0.012 \pm 0.006$<br>-6.350 : - | $0.062 \pm 0.005$<br>32.64 : + | $0.001 \pm 0.005$<br>0.350 : 0 | $0.001 \pm 0.005$<br>0.353 : 0 |
| $4 \rightarrow 1$ | $-0.160 \pm 0.018$<br>-23.41 : - | $-0.013 \pm 0.020$<br>-1.656 : 0 | $0.295 \pm 0.016$<br>45.38 : + | $0.384 \pm 0.015$<br>62.62 : + | $-0.061 \pm 0.005$<br>-31.98 : - | $0.013 \pm 0.006$<br>6.819 : + | $0.001 \pm 0.005$<br>0.739 : 0 | $-0.001 \pm 0.005$<br>-0.314 : 0 |
| $2 \rightarrow 3$ | $0.294 \pm 0.017$<br>45.10 : + | $0.393 \pm 0.015$<br>63.68 : + | $-0.149 \pm 0.018$<br>-21.95 : - | $0.010 \pm 0.020$<br>1.303 : 0 | $0.015 \pm 0.006$<br>7.748 : + | $-0.063 \pm 0.005$<br>-33.07 : - | $0.003 \pm 0.005$<br>1.722 : 0 | $-0.000 \pm 0.005$<br>-0.125 : 0 |
| $3 \rightarrow 2$ | $-0.005 \pm 0.020$<br>-0.663 : 0 | $0.154 \pm 0.018$<br>22.45 : + | $-0.379 \pm 0.015$<br>-61.66 : - | $-0.281 \pm 0.017$<br>-43.28 : - | $0.060 \pm 0.005$<br>31.45 : + | $-0.015 \pm 0.006$<br>-7.821 : - | $-0.001 \pm 0.005$<br>-0.396 : 0 | $-0.003 \pm 0.005$<br>-1.527 : 0 |
| $2 \rightarrow 4$ | $-0.287 \pm 0.017$<br>-44.19 : - | $-0.390 \pm 0.015$<br>-63.52 : - | $0.005 \pm 0.020$<br>0.615 : 0 | $-0.161 \pm 0.018$<br>-23.66 : - | $0.014 \pm 0.006$<br>7.093 : + | $0.061 \pm 0.005$<br>32.20 : + | $0.001 \pm 0.005$<br>0.443 : 0 | $-0.001 \pm 0.005$<br>-0.545 : 0 |
| $4 \rightarrow 2$ | $-0.013 \pm 0.020$<br>-1.637 : 0 | $-0.174 \pm 0.018$<br>-25.46 : - | $-0.283 \pm 0.017$<br>-43.46 : - | $-0.385 \pm 0.015$<br>-62.58 : - | $0.063 \pm 0.005$<br>33.25 : + | $0.015 \pm 0.006$<br>7.560 : + | $0.002 \pm 0.005$<br>1.096 : 0 | $-0.000 \pm 0.005$<br>-0.146 : 0 |
| $12345 \rightarrow 1$ | $0.010 \pm 0.021$<br>1.183 : 0 | $0.006 \pm 0.021$<br>0.750 : 0 | $-0.285 \pm 0.017$<br>-43.67 : - | $-0.289 \pm 0.017$<br>-44.15 : - | $0.070 \pm 0.005$<br>37.59 : + | $0.003 \pm 0.006$<br>1.779 : 0 | $0.023 \pm 0.005$<br>12.57 : + | $0.004 \pm 0.005$<br>2.226 : 0 |
| $34 \rightarrow 2$ | $0.017 \pm 0.026$<br>1.683 : 0 | $0.021 \pm 0.024$<br>2.342 : 0 | $-0.262 \pm 0.016$<br>-43.38 : - | $-0.259 \pm 0.016$<br>-42.84 : - | $0.045 \pm 0.005$<br>26.06 : + | $-0.004 \pm 0.006$<br>-1.771 : 0 | $0.002 \pm 0.005$<br>1.229 : 0 | $-0.002 \pm 0.006$<br>-0.857 : 0 |
| $2 \rightarrow 34$ | $0.007 \pm 0.020$<br>0.948 : 0 | $0.005 \pm 0.019$<br>0.698 : 0 | $-0.089 \pm 0.017$<br>-13.86 : - | $-0.090 \pm 0.017$<br>-14.04 : - | $0.016 \pm 0.005$<br>8.804 : + | $-0.003 \pm 0.006$<br>-1.372 : 0 | $0.002 \pm 0.005$<br>1.112 : 0 | $-0.002 \pm 0.006$<br>-0.898 : 0 |
| $1234 \rightarrow 1$ | $0.003 \pm 0.021$<br>0.416 : 0 | $0.004 \pm 0.020$<br>0.584 : 0 | $-0.160 \pm 0.018$<br>-23.33 : - | $-0.160 \pm 0.018$<br>-23.20 : - | $0.023 \pm 0.005$<br>12.37 : + | $-0.003 \pm 0.006$<br>-1.386 : 0 | $-0.001 \pm 0.005$<br>-0.383 : 0 | $-0.002 \pm 0.005$<br>-1.118 : 0 |
| $5 \rightarrow 1$ | $-0.003 \pm 0.021$<br>-0.350 : 0 | $-0.011 \pm 0.020$<br>-1.364 : 0 | $-0.610 \pm 0.011$<br>-121.1 : - | $-0.613 \pm 0.011$<br>-121.8 : - | $0.049 \pm 0.005$<br>26.31 : + | $-0.001 \pm 0.006$<br>-0.604 : 0 | $-0.113 \pm 0.005$<br>-62.14 : - | $-0.002 \pm 0.005$<br>-1.018 : 0 |
| $1 \rightarrow 5$ | $0.012 \pm 0.021$<br>1.471 : 0 | $0.019 \pm 0.021$<br>2.401 : 0 | $-0.157 \pm 0.018$<br>-23.09 : - | $-0.152 \pm 0.018$<br>-22.37 : - | $-0.004 \pm 0.005$<br>-1.983 : 0 | $0.001 \pm 0.005$<br>0.416 : 0 | $-0.028 \pm 0.005$<br>-14.62 : - | $0.003 \pm 0.005$<br>1.340 : 0 |
| $12345 \rightarrow 2$ | $0.002 \pm 0.021$<br>0.250 : 0 | $-0.002 \pm 0.021$<br>-0.298 : 0 | $0.279 \pm 0.017$<br>42.07 : + | $0.276 \pm 0.017$<br>41.60 : + | $-0.069 \pm 0.005$<br>-37.43 : - | $0.001 \pm 0.006$<br>0.713 : 0 | $-0.025 \pm 0.005$<br>-13.88 : - | $0.001 \pm 0.005$<br>0.694 : 0 |
| $34 \rightarrow 1$ | $0.001 \pm 0.024$<br>0.146 : 0 | $0.009 \pm 0.026$<br>0.944 : 0 | $0.267 \pm 0.015$<br>44.52 : + | $0.270 \pm 0.015$<br>45.02 : + | $-0.044 \pm 0.005$<br>-25.36 : - | $0.001 \pm 0.006$<br>0.649 : 0 | $0.001 \pm 0.005$<br>0.557 : 0 | $0.002 \pm 0.006$<br>0.868 : 0 |
| $1 \rightarrow 34$ | $-0.004 \pm 0.019$<br>-0.569 : 0 | $-0.012 \pm 0.020$<br>-1.540 : 0 | $0.101 \pm 0.017$<br>15.79 : + | $0.097 \pm 0.017$<br>15.06 : + | $-0.013 \pm 0.005$<br>-7.549 : - | $-0.001 \pm 0.006$<br>-0.608 : 0 | $0.002 \pm 0.005$<br>0.959 : 0 | $-0.002 \pm 0.006$<br>-1.154 : 0 |
| $1234 \rightarrow 2$ | $0.003 \pm 0.020$<br>0.372 : 0 | $0.001 \pm 0.021$<br>0.166 : 0 | $0.151 \pm 0.018$<br>21.61 : + | $0.149 \pm 0.018$<br>21.37 : + | $-0.020 \pm 0.005$<br>-10.65 : - | $-0.000 \pm 0.006$<br>-0.016 : 0 | $0.002 \pm 0.005$<br>0.991 : 0 | $0.000 \pm 0.005$<br>0.117 : 0 |
| $5 \rightarrow 2$ | $0.009 \pm 0.020$<br>1.116 : 0 | $0.005 \pm 0.021$<br>0.583 : 0 | $0.606 \pm 0.011$<br>119.8 : + | $0.606 \pm 0.011$<br>119.6 : + | $-0.043 \pm 0.005$<br>-23.51 : - | $0.001 \pm 0.006$<br>0.501 : 0 | $0.115 \pm 0.005$<br>63.10 : + | $0.002 \pm 0.005$<br>0.910 : 0 |
| $2 \rightarrow 5$ | $0.006 \pm 0.021$<br>0.761 : 0 | $0.001 \pm 0.021$<br>0.095 : 0 | $0.162 \pm 0.018$<br>23.81 : + | $0.158 \pm 0.018$<br>23.28 : + | $-0.001 \pm 0.005$<br>-0.326 : 0 | $0.001 \pm 0.005$<br>0.364 : 0 | $0.024 \pm 0.005$<br>12.82 : + | $0.001 \pm 0.005$<br>0.567 : 0 |
| $12345 \rightarrow 3$ | $-0.293 \pm 0.017$<br>-44.51 : - | $-0.295 \pm 0.017$<br>-44.90 : - | $-0.003 \pm 0.021$<br>-0.377 : 0 | $-0.007 \pm 0.020$<br>-0.912 : 0 | $0.003 \pm 0.006$<br>1.378 : 0 | $0.072 \pm 0.005$<br>39.02 : + | $0.002 \pm 0.005$<br>1.039 : 0 | $0.025 \pm 0.005$<br>13.72 : + |
| $12 \rightarrow 4$ | $-0.270 \pm 0.015$<br>-44.94 : - | $-0.268 \pm 0.015$<br>-44.64 : - | $0.001 \pm 0.026$<br>0.141 : 0 | $0.006 \pm 0.024$<br>0.627 : 0 | $0.004 \pm 0.006$<br>1.706 : 0 | $0.044 \pm 0.005$<br>25.32 : + | $0.004 \pm 0.006$<br>1.851 : 0 | $-0.001 \pm 0.005$<br>-0.541 : 0 |
| $4 \rightarrow 12$ | $-0.105 \pm 0.017$<br>-16.31 : - | $-0.104 \pm 0.017$<br>-16.21 : - | $0.006 \pm 0.020$<br>0.782 : 0 | $0.006 \pm 0.019$<br>0.855 : 0 | $-0.001 \pm 0.006$<br>-0.246 : 0 | $0.014 \pm 0.005$<br>7.841 : + | $0.000 \pm 0.006$<br>0.202 : 0 | $-0.002 \pm 0.005$<br>-1.092 : 0 |
| $1234 \rightarrow 3$ | $-0.165 \pm 0.018$<br>-24.06 : - | $-0.167 \pm 0.018$<br>-24.31 : - | $0.000 \pm 0.021$<br>0.024 : 0 | $-0.002 \pm 0.021$<br>-0.264 : 0 | $-0.002 \pm 0.006$<br>-1.006 : 0 | $0.024 \pm 0.005$<br>12.65 : + | $-0.002 \pm 0.005$<br>-1.053 : 0 | $-0.001 \pm 0.005$<br>-0.591 : 0 |
| $5 \rightarrow 3$ | $-0.615 \pm 0.011$<br>-121.8 : - | $-0.614 \pm 0.011$<br>-121.9 : - | $0.000 \pm 0.021$<br>0.049 : 0 | $-0.004 \pm 0.020$<br>-0.477 : 0 | $-0.000 \pm 0.005$<br>-0.033 : 0 | $0.045 \pm 0.005$<br>24.34 : + | $-0.000 \pm 0.005$<br>-0.140 : 0 | $-0.118 \pm 0.005$<br>-64.31 : - |
| $3 \rightarrow 5$ | $-0.145 \pm 0.018$<br>-21.47 : - | $-0.151 \pm 0.018$<br>-22.37 : - | $0.011 \pm 0.021$<br>1.367 : 0 | $0.003 \pm 0.021$<br>0.331 : 0 | $-0.002 \pm 0.005$<br>-0.783 : 0 | $0.001 \pm 0.005$<br>0.723 : 0 | $-0.001 \pm 0.005$<br>-0.368 : 0 | $-0.022 \pm 0.005$<br>-11.60 : - |
| $12345 \rightarrow 4$ | $0.289 \pm 0.017$<br>44.06 : + | $0.290 \pm 0.017$<br>44.32 : + | $-0.009 \pm 0.020$<br>-1.115 : 0 | $-0.007 \pm 0.021$<br>-0.849 : 0 | $0.002 \pm 0.006$<br>1.142 : 0 | $-0.068 \pm 0.005$<br>-36.98 : - | $0.001 \pm 0.005$<br>0.641 : 0 | $-0.022 \pm 0.005$<br>-11.89 : - |
| $12 \rightarrow 3$ | $0.268 \pm 0.015$<br>44.58 : + | $0.270 \pm 0.015$<br>44.98 : + | $-0.013 \pm 0.024$<br>-1.449 : 0 | $-0.009 \pm 0.026$<br>-0.863 : 0 | $0.002 \pm 0.006$<br>0.865 : 0 | $-0.041 \pm 0.005$<br>-23.62 : - | $0.001 \pm 0.006$<br>0.342 : 0 | $0.004 \pm 0.005$<br>2.243 : 0 |
| $3 \rightarrow 12$ | $0.098 \pm 0.017$<br>15.50 : + | $0.101 \pm 0.017$<br>15.85 : + | $0.019 \pm 0.019$<br>2.685 : + | $0.025 \pm 0.020$<br>3.246 : + | $-0.003 \pm 0.006$<br>-1.591 : 0 | $-0.013 \pm 0.005$<br>-7.530 : - | $0.000 \pm 0.006$<br>0.032 : 0 | $0.002 \pm 0.005$<br>1.189 : 0 |
| $1234 \rightarrow 4$ | $0.157 \pm 0.018$<br>22.85 : + | $0.161 \pm 0.018$<br>23.34 : + | $-0.001 \pm 0.021$<br>-0.133 : 0 | $0.003 \pm 0.021$<br>0.389 : 0 | $-0.001 \pm 0.006$<br>-0.287 : 0 | $-0.025 \pm 0.005$<br>-13.10 : - | $-0.000 \pm 0.005$<br>-0.222 : 0 | $-0.001 \pm 0.005$<br>-0.489 : 0 |
| $5 \rightarrow 4$ | $0.608 \pm 0.011$<br>119.9 : + | $0.604 \pm 0.011$<br>119.3 : + | $0.015 \pm 0.020$<br>1.982 : 0 | $0.009 \pm 0.021$<br>1.098 : 0 | $-0.002 \pm 0.006$<br>-1.144 : 0 | $-0.042 \pm 0.005$<br>-22.45 : - | $-0.001 \pm 0.005$<br>-0.374 : 0 | $0.117 \pm 0.005$<br>64.13 : + |
| $4 \rightarrow 5$ | $0.161 \pm 0.018$<br>23.76 : + | $0.164 \pm 0.018$<br>24.29 : + | $-0.007 \pm 0.021$<br>-0.944 : 0 | $-0.003 \pm 0.021$<br>-0.317 : 0 | $0.001 \pm 0.005$<br>0.368 : 0 | $-0.001 \pm 0.005$<br>-0.398 : 0 | $0.000 \pm 0.005$<br>0.061 : 0 | $0.025 \pm 0.005$<br>13.08 : + |

**Table 6:** Simulation results for the asymmetric tree *A*. For each statistic under each scenario we report the value, the 99% Wald confidence interval with radius 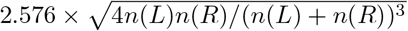, the *Z*-score and the classification. All results match the predictions in Table 3.

| | $A\Delta_{1-2}^*$ | $A\Delta_{1-3}^*$ | $A\Delta_{2-3}^*$ | $A\Delta_{1+2-3+4}^*$ |
| --- | --- | --- | --- | --- |
| $1 \rightarrow 3$ | $0.103 \pm 0.017$<br>16.02 : + | $0.103 \pm 0.017$<br>16.08 : + | $0.002 \pm 0.026$<br>0.160 : 0 | $0.001 \pm 0.006$<br>0.338 : 0 |
| $3 \rightarrow 1$ | $0.265 \pm 0.015$<br>43.99 : + | $0.267 \pm 0.015$<br>44.23 : + | $0.002 \pm 0.025$<br>0.210 : 0 | $0.001 \pm 0.006$<br>0.318 : 0 |
| $123 \rightarrow 2$ | $0.101 \pm 0.017$<br>15.83 : + | $0.101 \pm 0.017$<br>15.79 : + | $-0.001 \pm 0.024$<br>-0.083 : 0 | $0.003 \pm 0.006$<br>1.712 : 0 |
| $2 \rightarrow 3$ | $-0.089 \pm 0.017$<br>-13.87 : - | $-0.094 \pm 0.017$<br>-14.72 : - | $-0.014 \pm 0.026$<br>-1.366 : 0 | $0.002 \pm 0.006$<br>0.769 : 0 |
| $3 \rightarrow 2$ | $-0.265 \pm 0.015$<br>-44.36 : - | $-0.263 \pm 0.015$<br>-44.03 : - | $0.005 \pm 0.025$<br>0.510 : 0 | $0.003 \pm 0.006$<br>1.500 : 0 |
| $123 \rightarrow 1$ | $-0.099 \pm 0.017$<br>-15.40 : - | $-0.095 \pm 0.017$<br>-14.73 : - | $0.008 \pm 0.024$<br>0.832 : 0 | $-0.003 \pm 0.006$<br>-1.449 : 0 |
| $1 \rightarrow 4$ | $-0.096 \pm 0.017$<br>-15.11 : - | $0.005 \pm 0.018$<br>0.729 : 0 | $0.169 \pm 0.021$<br>20.47 : + | $-0.001 \pm 0.006$<br>-0.664 : 0 |
| $4 \rightarrow 1$ | $-0.471 \pm 0.012$<br>-89.68 : - | $-0.193 \pm 0.016$<br>-30.91 : - | $0.552 \pm 0.015$<br>81.83 : + | $0.002 \pm 0.006$<br>1.070 : 0 |
| $2 \rightarrow 4$ | $0.105 \pm 0.017$<br>16.48 : + | $0.008 \pm 0.018$<br>1.131 : 0 | $-0.167 \pm 0.022$<br>-20.16 : - | $0.002 \pm 0.006$<br>0.791 : 0 |
| $4 \rightarrow 2$ | $0.476 \pm 0.012$<br>90.96 : + | $0.187 \pm 0.016$<br>30.07 : + | $-0.572 \pm 0.015$<br>-84.66 : - | $-0.001 \pm 0.006$<br>-0.456 : 0 |
| $1 \rightarrow 5$ | $-0.004 \pm 0.018$<br>-0.573 : 0 | $-0.104 \pm 0.017$<br>-16.50 : - | $-0.170 \pm 0.021$<br>-20.73 : - | $-0.022 \pm 0.006$<br>-10.59 : - |
| $5 \rightarrow 1$ | $-0.190 \pm 0.016$<br>-30.57 : - | $-0.593 \pm 0.010$<br>-127.4 : - | $-0.710 \pm 0.011$<br>-126.3 : - | $-0.134 \pm 0.005$<br>-68.70 : - |
| $1234 \rightarrow 1$ | $-0.195 \pm 0.016$<br>-30.99 : - | $-0.266 \pm 0.016$<br>-43.89 : - | $-0.163 \pm 0.022$<br>-19.39 : - | $-0.000 \pm 0.006$<br>-0.111 : 0 |
| $12345 \rightarrow 1$ | $-0.198 \pm 0.016$<br>-31.28 : - | $-0.344 \pm 0.015$<br>-58.45 : - | $-0.319 \pm 0.020$<br>-39.85 : - | $0.023 \pm 0.006$<br>11.54 : + |
| $2 \rightarrow 5$ | $-0.008 \pm 0.018$<br>-1.213 : 0 | $0.089 \pm 0.017$<br>13.98 : + | $0.161 \pm 0.021$<br>19.54 : + | $0.021 \pm 0.006$<br>10.41 : + |
| $5 \rightarrow 2$ | $0.191 \pm 0.016$<br>30.84 : + | $0.590 \pm 0.010$<br>127.1 : + | $0.708 \pm 0.011$<br>125.8 : + | $0.132 \pm 0.005$<br>67.52 : + |
| $1234 \rightarrow 2$ | $0.186 \pm 0.016$<br>29.71 : + | $0.259 \pm 0.016$<br>42.83 : + | $0.163 \pm 0.022$<br>19.51 : + | $-0.001 \pm 0.006$<br>-0.512 : 0 |
| $12345 \rightarrow 2$ | $0.184 \pm 0.016$<br>29.26 : + | $0.331 \pm 0.015$<br>56.60 : + | $0.317 \pm 0.020$<br>39.89 : + | $-0.023 \pm 0.006$<br>-11.76 : - |

**Table 7:** Simulation results for the quasisymmetric tree *Q*. For each statistic under each scenario we report the value, the 99% Wald confidence interval with radius 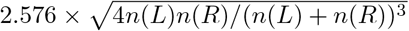, the *Z*-score and the classification. All results match the predictions in Table 4.

| | $Q\Delta_{1-6}^*$ | $Q\Delta_{2-6}^*$ | $Q\Delta_{3-5}^*$ | $Q\Delta_{4-5}^*$ | $Q\Delta_{5+7}^*$ | $Q\Delta_{6+8}^*$ | $Q\Delta_{3+4-5+7}^*$ | $Q\Delta_{1+2-6+8}^*$ |
| --- | --- | --- | --- | --- | --- | --- | --- | --- |
| $1 \rightarrow 4$ | $0.568 \pm 0.019$<br>65.18 : + | $0.435 \pm 0.022$<br>46.41 : + | $0.096 \pm 0.017$<br>14.97 : + | $-0.009 \pm 0.018$<br>-1.292 : 0 | $-0.011 \pm 0.006$<br>-5.424 : - | $-0.077 \pm 0.004$<br>-50.61 : - | $-0.002 \pm 0.006$<br>-0.869 : 0 | $-0.047 \pm 0.004$<br>-31.45 : - |
| $4 \rightarrow 1$ | $0.710 \pm 0.014$<br>93.41 : + | $0.006 \pm 0.038$<br>0.433 : 0 | $0.592 \pm 0.010$<br>126.5 : + | $0.437 \pm 0.013$<br>80.37 : + | $-0.091 \pm 0.006$<br>-44.70 : - | $-0.029 \pm 0.005$<br>-18.45 : - | $0.077 \pm 0.006$<br>38.91 : + | $0.001 \pm 0.004$<br>0.743 : 0 |
| $1 \rightarrow 5$ | $-0.563 \pm 0.019$<br>-64.46 : - | $-0.437 \pm 0.022$<br>-46.68 : - | $-0.006 \pm 0.018$<br>-0.941 : 0 | $0.092 \pm 0.017$<br>14.44 : + | $-0.009 \pm 0.006$<br>-4.240 : - | $0.074 \pm 0.004$<br>48.65 : + | $0.001 \pm 0.006$<br>0.243 : 0 | $0.045 \pm 0.004$<br>29.60 : + |
| $5 \rightarrow 1$ | $-0.701 \pm 0.014$<br>-92.27 : - | $-0.019 \pm 0.037$<br>-1.338 : 0 | $0.436 \pm 0.013$<br>80.29 : + | $0.587 \pm 0.010$<br>125.5 : + | $-0.089 \pm 0.006$<br>-44.19 : - | $0.029 \pm 0.005$<br>18.61 : + | $0.077 \pm 0.006$<br>38.95 : + | $-0.001 \pm 0.004$<br>-0.505 : 0 |
| $2 \rightarrow 4$ | $0.443 \pm 0.022$<br>47.16 : + | $0.571 \pm 0.019$<br>65.43 : + | $-0.101 \pm 0.017$<br>-15.76 : - | $0.000 \pm 0.018$<br>0.069 : 0 | $0.013 \pm 0.006$<br>6.102 : + | $-0.075 \pm 0.004$<br>-49.13 : - | $0.002 \pm 0.006$<br>0.914 : 0 | $-0.045 \pm 0.004$<br>-29.86 : - |
| $4 \rightarrow 2$ | $0.007 \pm 0.038$<br>0.476 : 0 | $0.714 \pm 0.014$<br>93.85 : + | $-0.589 \pm 0.010$<br>-125.8 : - | $-0.432 \pm 0.013$<br>-79.23 : - | $0.088 \pm 0.006$<br>43.50 : + | $-0.031 \pm 0.005$<br>-19.90 : - | $-0.077 \pm 0.006$<br>-39.21 : - | $-0.001 \pm 0.004$<br>-0.602 : 0 |
| $2 \rightarrow 5$ | $-0.434 \pm 0.022$<br>-46.83 : - | $-0.566 \pm 0.019$<br>-65.40 : - | $-0.002 \pm 0.018$<br>-0.338 : 0 | $-0.104 \pm 0.017$<br>-16.31 : - | $0.013 \pm 0.006$<br>6.235 : + | $0.075 \pm 0.004$<br>49.49 : + | $0.002 \pm 0.006$<br>0.741 : 0 | $0.045 \pm 0.004$<br>30.07 : + |
| $5 \rightarrow 2$ | $-0.030 \pm 0.038$<br>-2.085 : 0 | $-0.717 \pm 0.014$<br>-94.23 : - | $-0.431 \pm 0.013$<br>-79.23 : - | $-0.586 \pm 0.010$<br>-125.3 : - | $0.090 \pm 0.006$<br>44.59 : + | $0.031 \pm 0.005$<br>19.82 : + | $-0.075 \pm 0.006$<br>-38.07 : - | $0.000 \pm 0.004$<br>0.248 : 0 |
| $2 \rightarrow 45$ | $0.000 \pm 0.025$<br>0.029 : 0 | $0.005 \pm 0.023$<br>0.548 : 0 | $-0.105 \pm 0.017$<br>-16.58 : - | $-0.104 \pm 0.017$<br>-16.25 : - | $0.017 \pm 0.005$<br>9.583 : + | $0.002 \pm 0.006$<br>1.039 : 0 | $0.001 \pm 0.005$<br>0.489 : 0 | $0.002 \pm 0.006$<br>1.152 : 0 |
| $45 \rightarrow 2$ | $-0.031 \pm 0.054$<br>-1.489 : 0 | $-0.011 \pm 0.037$<br>-0.786 : 0 | $-0.478 \pm 0.012$<br>-91.14 : - | $-0.476 \pm 0.012$<br>-90.96 : - | $0.074 \pm 0.005$<br>43.48 : + | $0.000 \pm 0.006$<br>0.162 : 0 | $-0.027 \pm 0.005$<br>-15.70 : - | $-0.000 \pm 0.006$<br>-0.107 : 0 |
| $1 \rightarrow 3$ | $-0.009 \pm 0.039$<br>-0.620 : 0 | $-0.011 \pm 0.039$<br>-0.741 : 0 | $-0.094 \pm 0.017$<br>-14.71 : - | $-0.094 \pm 0.017$<br>-14.77 : - | $0.005 \pm 0.006$<br>2.398 : 0 | $0.001 \pm 0.004$<br>0.799 : 0 | $-0.015 \pm 0.006$<br>-7.215 : - | $0.001 \pm 0.004$<br>0.656 : 0 |
| $3 \rightarrow 1$ | $-0.025 \pm 0.039$<br>-1.679 : 0 | $-0.024 \pm 0.038$<br>-1.649 : 0 | $-0.263 \pm 0.015$<br>-43.59 : - | $-0.262 \pm 0.015$<br>-43.54 : - | $-0.001 \pm 0.006$<br>-0.718 : 0 | $-0.002 \pm 0.004$<br>-1.583 : 0 | $-0.062 \pm 0.006$<br>-30.30 : - | $-0.003 \pm 0.004$<br>-1.926 : 0 |
| $123 \rightarrow 2$ | $0.004 \pm 0.038$<br>0.294 : 0 | $0.003 \pm 0.036$<br>0.194 : 0 | $-0.203 \pm 0.016$<br>-32.22 : - | $-0.203 \pm 0.016$<br>-32.27 : - | $0.001 \pm 0.006$<br>0.432 : 0 | $0.001 \pm 0.004$<br>0.465 : 0 | $-0.041 \pm 0.006$<br>-20.21 : - | $0.001 \pm 0.004$<br>0.515 : 0 |
| $12345 \rightarrow 2$ | $-0.014 \pm 0.039$<br>-0.962 : 0 | $-0.018 \pm 0.033$<br>-1.385 : 0 | $-0.339 \pm 0.015$<br>-57.65 : - | $-0.341 \pm 0.015$<br>-57.93 : - | $-0.024 \pm 0.006$<br>-11.90 : - | $0.001 \pm 0.005$<br>0.637 : 0 | $-0.099 \pm 0.006$<br>-50.33 : - | $0.001 \pm 0.004$<br>0.364 : 0 |
| $1 \rightarrow 45$ | $-0.003 \pm 0.023$<br>-0.304 : 0 | $0.001 \pm 0.025$<br>0.086 : 0 | $0.094 \pm 0.017$<br>14.65 : + | $0.096 \pm 0.017$<br>14.91 : + | $-0.016 \pm 0.005$<br>-9.232 : - | $-0.001 \pm 0.006$<br>-0.364 : 0 | $-0.002 \pm 0.005$<br>-1.082 : 0 | $-0.001 \pm 0.006$<br>-0.409 : 0 |
| $45 \rightarrow 1$ | $-0.033 \pm 0.036$<br>-2.388 : 0 | $-0.025 \pm 0.054$<br>-1.214 : 0 | $0.477 \pm 0.012$<br>91.08 : + | $0.479 \pm 0.012$<br>91.59 : + | $-0.071 \pm 0.005$<br>-41.56 : - | $-0.000 \pm 0.006$<br>-0.133 : 0 | $0.031 \pm 0.005$<br>17.98 : + | $-0.001 \pm 0.006$<br>-0.610 : 0 |
| $2 \rightarrow 3$ | $-0.011 \pm 0.039$<br>-0.726 : 0 | $-0.008 \pm 0.039$<br>-0.522 : 0 | $0.099 \pm 0.017$<br>15.33 : + | $0.099 \pm 0.017$<br>15.43 : + | $0.002 \pm 0.006$<br>0.829 : 0 | $-0.000 \pm 0.004$<br>-0.021 : 0 | $0.022 \pm 0.006$<br>10.77 : + | $-0.000 \pm 0.004$<br>-0.150 : 0 |
| $3 \rightarrow 2$ | $-0.018 \pm 0.038$<br>-1.222 : 0 | $-0.011 \pm 0.040$<br>-0.698 : 0 | $0.280 \pm 0.015$<br>46.27 : + | $0.281 \pm 0.015$<br>46.47 : + | $-0.000 \pm 0.006$<br>-0.156 : 0 | $0.001 \pm 0.004$<br>0.559 : 0 | $0.064 \pm 0.006$<br>31.22 : + | $0.001 \pm 0.004$<br>0.360 : 0 |
| $123 \rightarrow 1$ | $-0.000 \pm 0.036$<br>-0.000 : 0 | $0.003 \pm 0.038$<br>0.174 : 0 | $0.183 \pm 0.016$<br>29.10 : + | $0.184 \pm 0.016$<br>29.16 : + | $0.002 \pm 0.006$<br>0.939 : 0 | $0.000 \pm 0.004$<br>0.136 : 0 | $0.039 \pm 0.006$<br>19.57 : + | $0.000 \pm 0.004$<br>0.154 : 0 |
| $12345 \rightarrow 1$ | $-0.012 \pm 0.033$<br>-0.921 : 0 | $-0.020 \pm 0.038$<br>-1.367 : 0 | $0.343 \pm 0.015$<br>58.41 : + | $0.343 \pm 0.015$<br>58.31 : + | $0.025 \pm 0.006$<br>12.48 : + | $0.002 \pm 0.005$<br>1.064 : 0 | $0.101 \pm 0.006$<br>51.37 : + | $0.001 \pm 0.004$<br>0.804 : 0 |
| $5 \rightarrow 12$ | $-0.574 \pm 0.019$<br>-65.77 : - | $-0.572 \pm 0.019$<br>-65.09 : - | $0.003 \pm 0.014$<br>0.590 : 0 | $0.005 \pm 0.013$<br>1.158 : 0 | $-0.001 \pm 0.006$<br>-0.538 : 0 | $0.038 \pm 0.005$<br>24.48 : + | $0.000 \pm 0.006$<br>0.173 : 0 | $0.002 \pm 0.004$<br>1.175 : 0 |
| $12 \rightarrow 5$ | $-0.250 \pm 0.026$<br>-24.80 : - | $-0.257 \pm 0.026$<br>-25.50 : - | $0.001 \pm 0.018$<br>0.159 : 0 | $-0.002 \pm 0.018$<br>-0.337 : 0 | $-0.001 \pm 0.006$<br>-0.695 : 0 | $0.060 \pm 0.004$<br>39.60 : + | $-0.002 \pm 0.006$<br>-0.743 : 0 | $0.048 \pm 0.004$<br>31.83 : + |
| $4 \rightarrow 3$ | $-0.719 \pm 0.014$<br>-94.84 : - | $-0.722 \pm 0.014$<br>-95.47 : - | $-0.007 \pm 0.019$<br>-0.984 : 0 | $-0.013 \pm 0.019$<br>-1.797 : 0 | $-0.003 \pm 0.006$<br>-1.398 : 0 | $-0.000 \pm 0.004$<br>-0.104 : 0 | $-0.005 \pm 0.006$<br>-2.190 : 0 | $-0.059 \pm 0.004$<br>-38.57 : - |
| $3 \rightarrow 4$ | $-0.566 \pm 0.019$<br>-65.45 : - | $-0.573 \pm 0.019$<br>-66.25 : - | $0.002 \pm 0.019$<br>0.258 : 0 | $-0.002 \pm 0.018$<br>-0.360 : 0 | $0.001 \pm 0.006$<br>0.460 : 0 | $-0.044 \pm 0.004$<br>-29.33 : - | $0.001 \pm 0.006$<br>0.423 : 0 | $-0.079 \pm 0.004$<br>-52.15 : - |
| $4 \rightarrow 12$ | $0.554 \pm 0.019$<br>63.62 : + | $0.571 \pm 0.019$<br>66.02 : + | $-0.001 \pm 0.013$<br>-0.289 : 0 | $0.007 \pm 0.014$<br>1.409 : 0 | $-0.001 \pm 0.006$<br>-0.355 : 0 | $-0.037 \pm 0.005$<br>-23.43 : - | $0.000 \pm 0.006$<br>0.057 : 0 | $-0.000 \pm 0.004$<br>-0.140 : 0 |
| $12 \rightarrow 4$ | $0.267 \pm 0.026$<br>26.18 : + | $0.261 \pm 0.026$<br>25.75 : + | $0.001 \pm 0.018$<br>0.081 : 0 | $-0.001 \pm 0.018$<br>-0.145 : 0 | $-0.003 \pm 0.006$<br>-1.203 : 0 | $-0.060 \pm 0.004$<br>-39.26 : - | $-0.003 \pm 0.006$<br>-1.208 : 0 | $-0.047 \pm 0.004$<br>-31.32 : - |
| $5 \rightarrow 3$ | $0.701 \pm 0.014$<br>92.21 : + | $0.702 \pm 0.014$<br>92.41 : + | $-0.009 \pm 0.019$<br>-1.219 : 0 | $-0.007 \pm 0.019$<br>-1.004 : 0 | $0.004 \pm 0.006$<br>1.746 : 0 | $-0.000 \pm 0.004$<br>-0.190 : 0 | $0.002 \pm 0.006$<br>1.104 : 0 | $0.057 \pm 0.004$<br>36.97 : + |
| $3 \rightarrow 5$ | $0.560 \pm 0.019$<br>64.41 : + | $0.564 \pm 0.019$<br>64.92 : + | $0.014 \pm 0.018$<br>1.987 : 0 | $0.018 \pm 0.019$<br>2.542 : 0 | $-0.005 \pm 0.006$<br>-2.516 : 0 | $0.047 \pm 0.004$<br>30.90 : + | $-0.002 \pm 0.006$<br>-1.146 : 0 | $0.080 \pm 0.004$<br>53.17 : + |
| $123 \rightarrow 5$ | $0.001 \pm 0.030$<br>0.092 : 0 | $-0.003 \pm 0.030$<br>-0.275 : 0 | $-0.009 \pm 0.018$<br>-1.254 : 0 | $-0.010 \pm 0.018$<br>-1.475 : 0 | $0.001 \pm 0.006$<br>0.395 : 0 | $0.034 \pm 0.004$<br>22.24 : + | $-0.001 \pm 0.006$<br>-0.430 : 0 | $0.033 \pm 0.004$<br>22.13 : + |
| $5 \rightarrow 123$ | $-0.003 \pm 0.025$<br>-0.333 : 0 | $-0.005 \pm 0.025$<br>-0.483 : 0 | $0.002 \pm 0.017$<br>0.380 : 0 | $0.002 \pm 0.017$<br>0.276 : 0 | $-0.002 \pm 0.006$<br>-0.810 : 0 | $0.011 \pm 0.004$<br>7.232 : + | $-0.001 \pm 0.006$<br>-0.588 : 0 | $0.011 \pm 0.004$<br>7.056 : + |
| $12345 \rightarrow 4$ | $0.003 \pm 0.036$<br>0.208 : 0 | $0.009 \pm 0.036$<br>0.635 : 0 | $-0.006 \pm 0.018$<br>-0.809 : 0 | $-0.004 \pm 0.018$<br>-0.594 : 0 | $-0.001 \pm 0.006$<br>-0.353 : 0 | $0.015 \pm 0.004$<br>9.775 : + | $-0.002 \pm 0.006$<br>-0.774 : 0 | $0.015 \pm 0.004$<br>9.839 : + |
| $123 \rightarrow 4$ | $0.011 \pm 0.030$<br>0.964 : 0 | $0.008 \pm 0.030$<br>0.710 : 0 | $0.000 \pm 0.018$<br>0.069 : 0 | $-0.001 \pm 0.018$<br>-0.083 : 0 | $0.004 \pm 0.006$<br>1.910 : 0 | $-0.035 \pm 0.004$<br>-23.06 : - | $0.004 \pm 0.006$<br>1.891 : 0 | $-0.034 \pm 0.004$<br>-22.74 : - |
| $4 \rightarrow 123$ | $0.009 \pm 0.025$<br>0.997 : 0 | $0.009 \pm 0.025$<br>0.952 : 0 | $0.006 \pm 0.017$<br>0.913 : 0 | $0.006 \pm 0.017$<br>0.892 : 0 | $-0.000 \pm 0.006$<br>-0.043 : 0 | $-0.013 \pm 0.004$<br>-8.313 : - | $0.001 \pm 0.006$<br>0.544 : 0 | $-0.012 \pm 0.004$<br>-7.947 : - |
| $12345 \rightarrow 5$ | $0.013 \pm 0.036$<br>0.939 : 0 | $0.000 \pm 0.036$<br>0.014 : 0 | $-0.009 \pm 0.018$<br>-1.304 : 0 | $-0.012 \pm 0.018$<br>-1.773 : 0 | $0.003 \pm 0.006$<br>1.373 : 0 | $-0.014 \pm 0.004$<br>-9.337 : - | $0.001 \pm 0.006$<br>0.438 : 0 | $-0.014 \pm 0.004$<br>-9.205 : - |

Compared with the *D*_FOIL_ method applied to the same simulated patterns on the symmetric tree *S* (Appendix 5), the Δ-statistics had similar accuracy with all three sample sizes, while the Δ-statistics limited to Table S1 in Appendix 3 had considerable advantages in power. We also show by simulations (Appendix 5) that the dependence of *D*_FOIL_ on the synchronization assumption (2) biases results on ancient data, that this problem is not caused by the outgroup mutation assumption (1), and that it cannot be alleviated by simply disregarding the singleton patterns.

### An empirical example: Brown bear – polar bear gene flow

Several independent population genomic studies of polar bears (*Ursus maritimus*) and brown bears (*U. arctos*) by different groups of investigators have in recent years reported on considerable ancient introgressive hybridization between the two bear lineages (Miller et al., 2012; Liu et al., 2014; Cahill et al., 2015; Lan et al., 2022; Wang et al., 2022). However, employing different approaches, findings on the directionality of this admixture have been contradictory. Hence, we sought here to reevaluate the hypotheses previously proposed by us and other groups concerning brown bear – polar bear gene flow using Δ-statistics.

For example, using five- and six-leaf admixture graphs and data from an ancient 120,000-year-old polar bear from Svalbard (which we dubbed APB – “ancient polar bear”), our own results strongly supported bidirectional admixture between these bear species, but with gene flow in the direction from brown bear to polar bear being the predominant pattern. Almost concurrently, another investigator group published a genome of their own ancient polar bear sample (which they named “Bruno”) and examined admixture between polar and brown bears using the *D*_FOIL_ method on tree *S* (Wang et al., 2022). Their results, which in fact principally provided an indecisive result when it came to gene flow direction (see Appendix 6), were interpreted to principally support unidirectional gene flow from polar bears into brown bears. Importantly, we note that the population history of brown and polar bears is complex and cannot alone be explained by models of admixture (see also Lan et al., 2022). Therefore, inferences of demographic histories of any set of populations and species would ideally incorporate multiple lines of evidence (see also Maier et al., 2023). Nevertheless, we find the reassessment of directionality of brown bear – polar bear gene flow and our application of the Δ-statistics appropriate, particularly given considerable previous attention to the subject and the importance of understanding polar bear evolutionary history, especially regarding impacts of climate change and the potential roles hybridization may have played (Lan et al., 2022).

We conducted Δ-statistics analyses using bear SNP data from our previous work (Lan et al., 2022), adding to it the data for the Bruno sample, to directly reassess the previously published hypotheses on gene flow directionality. Based on our previous results (Lan et al., 2022) (and initial analyses of PCA and TreeMix with our new dataset that included the Bruno sample (see Appendix 6)), we performed Δ-statistics analyses using the following populations: brown bears from Admiralty, Baranof, and Chichagof Islands (ABC), European brown bears (EBB), North American mainland brown bears (BB), modern polar bears (PB from Svalbard and AK from Alaska), the ancient individuals (APB and Bruno), and black bears (BLK). The following combinations were tested:

- Symmetric: *S* = (((1 = ABC, 2 = EBB/BB), (3 = APB/AK/Bruno, 4 = PB)), 5 = BLK)
- Asymmetric: *A* = ((((1 = ABC, 2 = BB), 3 = EBB), 4 = APB/Bruno/PB), 5 = BLK)
- Quasisymmetric: *Q* = (((1 = ABC, 2 = BB), 3 = EBB), (4 = APB/AK/Bruno, 5 = PB))

These combinations can be seen as various five-leaf perspectives that together provide a better picture of the full phylogenetic tree ((((ABC, BB), EBB), (APB/Bruno, (AK, PB))), BLK) (see also Figure S15 in Lan et al., 2022). Our Δ-statistics analyses involved non-overlapping windows of varying sizes (100 kb, 500 kb, 1 Mb, 5 Mb, and 10 Mb, Supplementary Tables S19–S24) to replicate *D*_FOIL_ analyses performed in Wang et al., 2022 (window sizes spanning 100 kb to 500 kb), and also to explore the impact of using window sizes several times larger than the extent of linkage disequilibrium (*>* 200 kb, see Liu et al., 2014). The sign of each Δ-statistic in each signature was determined using the threshold *α* = 0.01. Here, we present and discuss the results most relevant to the previously published hypotheses, using APB as representative for the ancient polar bear lineage (Figures 4 and 5), but for results from combinations not presented in Figures 4 and 5, see Appendix 6 (Figures S5–S7) and Supplementary Tables S19–S24.

**Figure 4:**
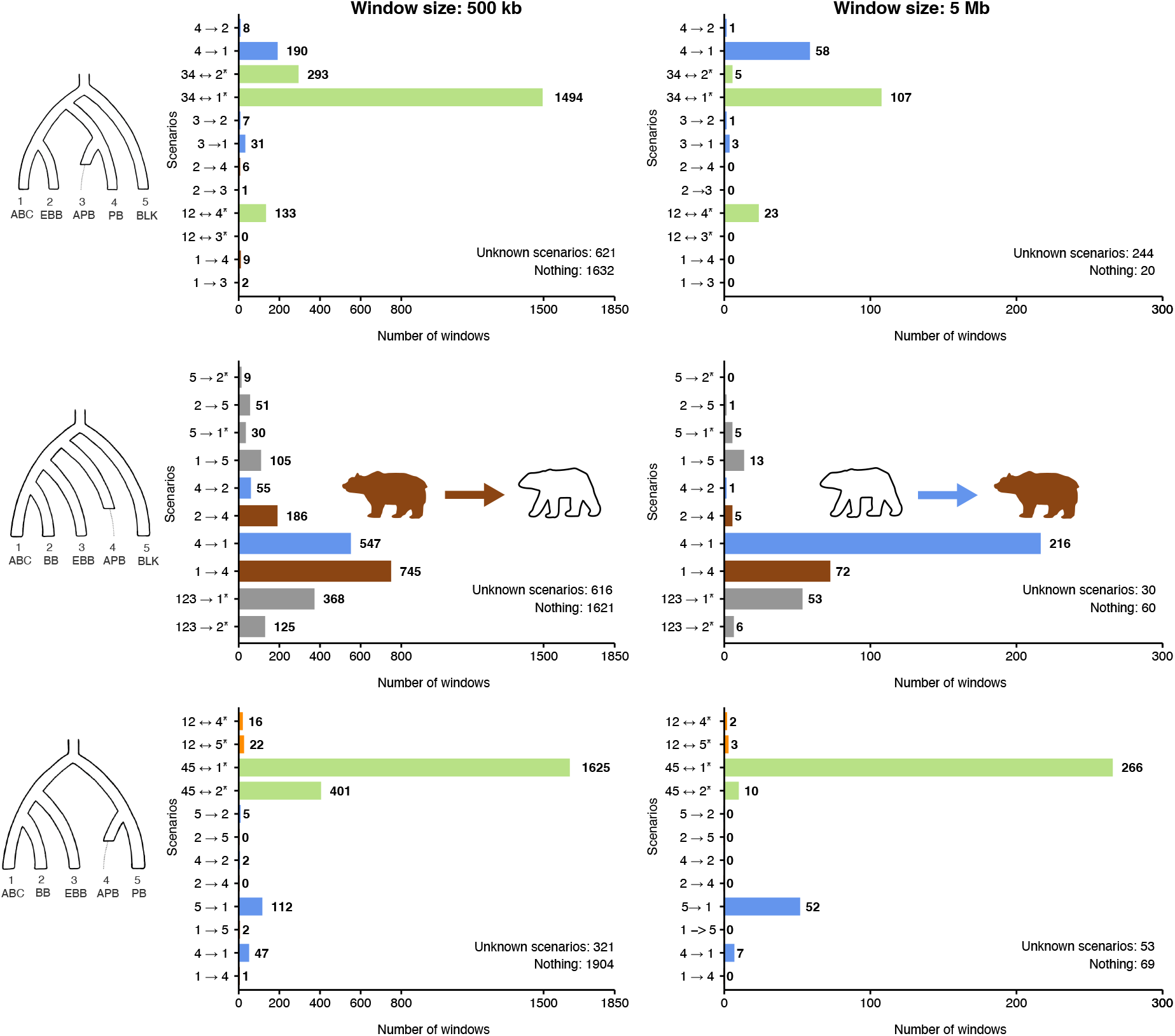
Bar plots of various Δ-statistic analyses of brown bear – polar bear gene flow incorporating a modern and an ancient polar bear (APB). Because APB is an ancient sample, we cannot make the synchronization assumption (2), so we therefore exclude singletons and use the collapsed Tables S1–S3 in Appendix 3. The asterisk means there are additional gene flow events consistent with the signature to the representative written, for example 34 ↔ 2^*^ in tree *S* stands for 12345 → 1, 34 ↔ 2, 1234 → 1 or 1 ↔ 5, according to Table S1 in Appendix 3. The colours of the bars signify the following: Brown — the signature supports only gene flow from brown bears into polar bears; Blue — the signature supports only gene flow from polar bears into brown bears; Orange — the signature supports gene flow in either direction between brown bears and polar bears; Gray — the signature supports some gene flow event that has nothing to do with the brown bear – polar bear question; Green — the signature supports gene flow of either or neither direction between brown bears and polar bears. For example, 34 ↔ 2^*^ is Green in tree *S* because it’s consistent with gene flow from brown bears into polar bears (2 → 34), with gene flow from polar bears into brown bears (34 → 2), and with gene flow that has nothing to do with the brown bear – polar bear question (such as 1234 → 1). The windows with zero signature are counted under “Nothing”, and windows with non-zero signature are discordant with any row in Tables S1–S3 in Appendix 3 under “Unknown scenarios”. See the main text for description of the populations and interpretation.

**Figure 5:**
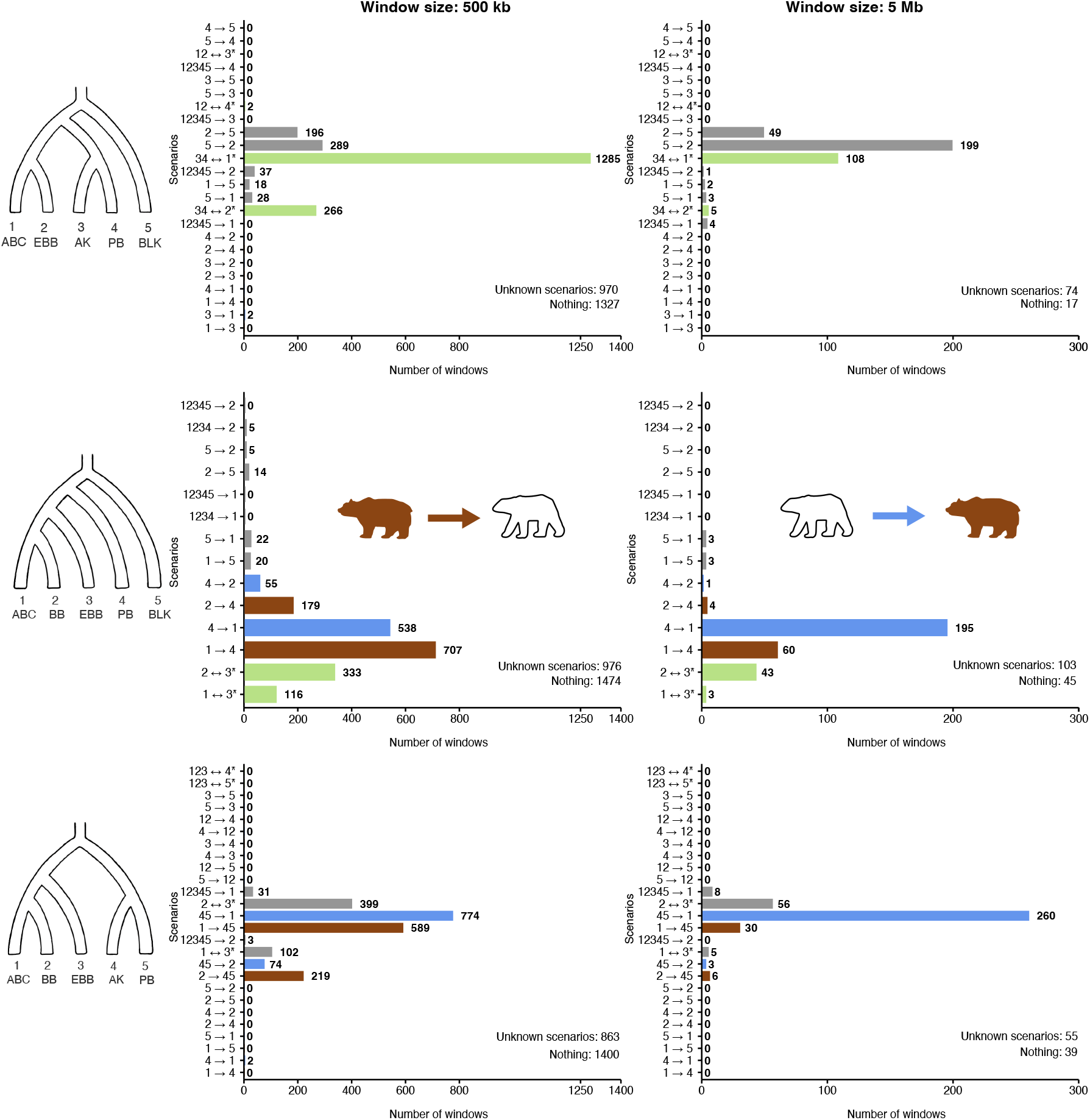
Bar plots of various Δ-statistic analyses of brown bear – polar bear gene flow incorporating two modern polar bears. Because all the samples are modern, we commit to the synchronization assumption (2) (although the generation times between populations likely vary a little), and therefore we use the complete Tables 2–4 in the main text. The asterisk means there are additional gene flow events consistent with the signature to the representative written, for example 34 ↔ 2^*^ in tree *S* stands for 1234 → 1 or 34 ↔ 2, according to Table 2 in the main text. The colours of the bars are as in Figure 4. The windows with zero signature are counted under “Nothing”, and windows with non-zero signature are discordant with any row in Tables S1–S3 in Appendix 3 under “Unknown scenarios”. See the main text for description of the populations and interpretation.

Starting with the symmetric tree, *S* = (((1 = ABC, 2 = EBB), (3 = APB/AK, 4 = PB)), 5 = BLK) (Figures 4 and 5, top rows), we discover that the greatest number of windows support events 34 ↔ 1^*^ (the asterisk emphasizing that there are other possible explanations aside from the chosen representative 34 ↔1; see Tables S1 and 2). Some of this signal in Figure 4 actually represents gene flow between EBB and BLK (Kumar et al., 2017), which separates into bars 2 →5 and 5 → 2 in Figure 5. The rest is either ancient admixture between brown bears and polar bears, or ghost relatives of all brown bears and polar bears (such as cave bears (Barlow et al., 2018)) introgressing into EBB (1234 → 2). There is clear modern gene flow from PB into ABC when 3 = APB (Figure 4), however, this signal dissolves when 3 is replaced with a modern polar bear individual (Figure 5), presumably because the split time between 3 and 4 is pulled towards the present. We also wish to address the apparent conversion of 34 ↔ 1^*^ into 5 → 2 as the window size is increased in Figure 5. We interpret this as a violation of the single event assumption (7). As it happens, the signature of 34 ↔ 1^*^ is (0 0 + + | − 0 0 0) and the signature of 5 → 2 is (0 0 + +| − 0 + 0). A window containing segments from both events will produce the mixture signature (0 0 + + 0 | − + 0) and be classified as 5 → 2. On the same note, the mixture of signals 4 → 1 and 34 ↔ 1^*^ in Figure 4 is dominated by 4 → 1, as can be observed comparing the two window sizes. Overall, the symmetric tree *S* provides evidence for modern polar bear → brown bear gene flow, and for admixture between European brown bears and black bears, but the strongest signal remains ambiguous in nature. This result is consistent when replacing APB with Bruno (Figure S5 in Appendix 6), and with *D*_FOIL_ results achieved by both Wang et al., 2022 and ourselves (Figures S8, S9 and discussion in Appendix 6). The difficulty in interpreting the strongest signal is in the tree *S* itself; even if we are looking at brown bear – polar bear admixture, the direction of gene flow can only be inferred if it is recent enough to affect only one of the two polar bear samples. As we currently do not have demonstrably unadmixed ancient polar bear specimens available, we overcome this methodological problem by studying the asymmetric tree *A*, which does not require an unadmixed sister sample.

The asymmetric tree, *A* = ((((1 = ABC, 2 = BB), 3 = EBB), 4 = APB/PB), 5 = BLK) (Figures 4 and 5 middle rows), provides evidence for gene flow in both directions between ABC brown bears and polar bears. Under bidirectional gene flow, the mixture of signatures (− − + | 0) for 4 → 1 and (− 0 +| 0) for 1 → 4 looks like the former; this dominance also applies without the singleton patterns. Thus, even tree *A* can be biased towards polar bear → brown bear gene flow, which contributes to the difference between results using 500 kb versus 5 Mb windows. Another contributing factor is that larger window sizes will favour more recent gene flow events whose traces are less uniformly distributed across the genome. Hence, similar to our own 2022 findings, the bidirectional brown – polar bear admixture appears to have a temporal element wherein brown bear → polar bear admixture, evidenced by genomic blocks substantially broken up by longterm recombination, is older than polar bear → brown bear gene flow, where larger admixed blocks might still predominate given fewer generations since gene flow. Because tree *A* cannot even detect admixture between EBB and BLK (3 ↔ 5) or the ghost relatives of all brown bears and polar bears (1234 → 3), the strong ambiguous signal from Wang et al., 2022 and our own symmetric tree *S* analyses were indeed largely due to ancient bidirectional gene flow between polar bears and ABC brown bears. Additionally, we detect gene flow or ILS within the brown bear clade (Lan et al., 2022).

We also experimented with the quasisymmetric tree, *Q* = (((1 = ABC, 2 = BB), 3 = EBB), (4 = APB/AK, 5 = PB)) (Figures 4 and 5 bottom rows). Using the ancient sample APB without singleton patterns, the results are very similar to those obtained with the symmetric tree *S*: a strong but ambiguous 45 ↔ 1^*^ signal and some evidence for recent polar bear → brown bear admixture. Using two modern polar bear samples and incorporating singleton patterns (Figure 5) the results resemble those obtained with the asymmetric tree *A*.

Our empirical example using Δ-statistics on bear data provides clear evidence for ancient bidirectional gene flow between polar bears and brown bears. This admixture appears to have a temporal element where the proportion of polar bear → brown bear gene flow increases over time relative to brown bear → polar bear gene flow, and a clear signature of modern polar bear brown → bear gene flow in trees *S* and *Q*. We also find signs of admixture within brown bears, whereas the two modern polar bear populations do not separate from each other at all, likely as a result of their recent split. Finally, we find signals of gene flow consistent with scenarios involving black bears, or possibly ghost relatives of all brown bears and polar bears. This example also highlights the importance of the synchronization assumption (2) using *D*_FOIL_ and some of the new Δ-statistics. The signature of gene flow from Bruno into North American brown bears detected by Wang et al., 2022 (gray box plots in their Figure 3) was several times more prevalent than the signature of gene flow from modern polar bears into North American brown bears (ochre box plots in their Figure 3). Such results can be explained simply by noting that as an ancient sample, Bruno has fewer private mutations than the modern polar bear sample, thereby biasing *D*_FOIL_. In contrast, Δ-statistics without singleton patterns on comparable trees (top left panel Figure 4 using APB and Figure S5 in Appendix 6 using Bruno) detect substantially more windows (190 windows versus 31, and 118 windows versus 61) supporting gene flow from the modern sample than from the ancient one. This result is what we would expect under occasional hybridization continuing to this day. We also tested what would have happened if we had used the full Tables 2–4 with the ancient sample APB; the dramatically biased results (Figure S6 and S7) can be found in Appendix 6.

Finally, to confirm that our dataset shows admixture signals comparable to Wang et al., 2022, and to compare our results from Δ-statistics with *D*_FOIL_, we ran *D*_FOIL_ using our expanded *D*_FOIL_ prediction table (Appendix 5). We tested the symmetric tree *S* = (((1 = ABC, 2 = EBB), (3 = X, 4 = PB)), 5 = BLK), where three individuals each for populations ABC, EBB, and PB were incorporated in independent runs of all combinations, and *x* could represent either APB, Bruno, a modern polar bear AK34, or three different modern polar bears from the AK group. The same window sizes were used as in our Δ-statistic analyses (100 kb, 500 kb, 1 Mb, 5 Mb, 10 Mb). Our *D*_FOIL_ analyses yielded results consistent with Wang et al., 2022 regardless of whether *x* represented a single polar bear (AK34) or three polar bears, except a wider range was observed in the box plots when more than one modern polar bear individual was in the third position (Appendix 6, Figure S8 and S9). Across all combinations, bidirectional admixture scenarios between polar and brown bears received the strongest support. Notably, there was a discrepancy between modern and ancient polar bears in terms of the number of windows supporting most scenarios, which is consistent with results from (Wang et al., 2022).

## 4 Discussion

We have developed a new method for gene flow detection — Δ-statistics — that can be seen as a natural generalization of the classic *D*-statistic to five-leaf trees. Additionally, Δ-statistics provide opportunities to discern admixture directions. Like the *D*-statistic, we worked with minimal assumptions and avoided parametrizing any unnecessary features of the species tree. Instead, we leveraged symmetry that resides in all trees, even the “asymmetric” tree *A*, to construct difference statistics whose expected signs are informative about admixture direction. We point out that while the good behaviour of the statistics under the null hypotheses relies on *symmetry*, their good behaviour under various gene flow events to some degree relies on *asymmetry* as well; for example, the direction of gene flow 12 ↔ 4 can be worked out in tree *Q* but not in tree *S*, ultimately because the position of the root in *Q* is less symmetric. The insight that we are drawing on the interplay between symmetry and asymmetry can be compared to the qpAdm method (part of the ADMIXTOOLS package (Patterson et al., 2012), see (Harney, Patterson, Reich, & Wakeley, 2021)), where the *reference* populations must be asymmetrically aligned with respect to the *source* and *target* populations. We believe ours are the best possible collections of Δ-statistics that can be constructed from binomial statistics using the equal probability sets as the starting point. Still, it should be pointed out that in tree *A* the statistic *n*(BAABA) + *n*(ABABA) + 2*n*(AABAB) − *n*(BAAAB) − *n*(ABAAB) − 2*n*(AABBA) mentioned in the Methods section is capable of indicating additional gene flow events such as 12 ↔ 3 and 12↔ 4. Pattern counting in the style of the *D*-statistic is straightforward to understand and perform, but this simple approach is only one among many in the wider population genetics setting. One complicating factor is the assumption of independence between loci, a context dependent problem for which we have chosen not to offer any general guidelines. Another potential issue lies in the discretizing of populations. Natural populations often form a dynamic continuum rather than distinct clusters connected by a small number of gene flow events, and approximating such continua by increasing this number can conflict with the single event assumption (7).

To compare our method to our inspiration, *D*_FOIL_ (Pease & Hahn, 2015), we must stress that *D*_FOIL_ makes the synchronization assumption (2), whereas the classic *D* does not, and we can either choose to make it or not with Δ-statistics. Without the synchronization assumption (2), our method is applicable to ancient samples and populations with unequal generation times, and Δ-statistics have more statistical power than *D*_FOIL_ does. For the capacity to infer gene flow events we should compare Table S1 in Appendix 3 to Table S4 in Appendix 5, because the original *D*_FOIL_ paper considered several gene flow events as being outside the realm of possibilities. The only difference in the tables is that *D*_FOIL_ is blind to gene flow into 5 from other terminal branches, while Δ-statistics can detect it but not distinguish it from a number of other gene flow events — and which is better is context dependent. On the other hand, if we make the synchronization assumption (2), our ability to distinguish different gene flow events is superior (full Table 2 in the main text), while statistical power remains comparable to *D*_FOIL_. The original *D*_FOIL_ article (Pease & Hahn, 2015) suggests that the dependence on the synchronization assumption (2) could be lifted by simply omitting all singleton patterns from both the numerator and the denominator of the statistics, in other words, using the statistics 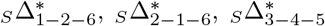 and 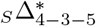 instead. This idea does not work, however, as the predicted signs change and even become inconsistent; see Appendix 5. Nonetheless, as an anonymous reviewer noticed, by removing not only the singletons but all the patterns Pease and Hahn call “inverse”, the *D*_FOIL_ statistics become (up to the outgroup mutation assumption (1)) our 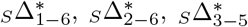 and 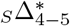 and are thereby free of (2).

We next discuss errors first discovered by Pease & Hahn, 2015 in another attempt to discern 3 → 2 and 2 → 3 in tree *S* — the Partitioned *D* method (Eaton & Ree, 2013). The proposed statistics are 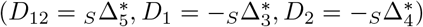, of which *D*_12_ is purportedly informative about gene flow direction. Simulations presented in their Table 3 result in signatures (±*±* 0) for 3 → 2 (tests 4.1–4.6) and (0 ± 0) for 2 → 3 (tests 6.1–6.6), where ± means a significantly non-zero statistic. However, as we show in Appendix 2, the actual expected signatures are (+ + ?) for 3 → 2 and (+ + −) for 2 → 3, where the question mark stands for an inconsistent statistic whose sign depends on graph parameters. Hence, *D*_12_ is not informative about gene flow direction, and the ability of the Partitioned *D* method to infer direction relies on benevolent graph parameters and insufficient power to detect the correct signature. The Eaton & Ree paper juxtaposes the method against an alternative idea of comparing several classic *D*-statistics calculated on a four-population subtree, which would in fact be a valid technique. Gene flow into the unsampled population cannot be detected, however, whereas gene flow from the unsampled population is effectively the same as gene flow from its sister population. This is why tests 2.4–2.7 in their Table 2 are consistent with the null *D* = 0, but tests 3.4–3.7 are not. In fact, by the law of total probability, ignoring one of the populations 1–4 in tree *S* and computing the classic *D* among the rest is equivalent to using the Δ-statistics 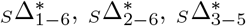 and 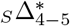.

One more point of comparison to Δ-statistics is the *D*_GEN_ method (Elworth, Allen, Benedict, Dulworth, & Nakhleh, 2018). The *D*_GEN_-statistic is defined using equal probability sets, as the Δ-statistics are. However, the user is expected to know the actual probabilities of each allelic pattern in order to construct the statistic as a sum of differences expected to be zero under the null hypothesis, and in turn positive under the single alternative hypothesis. This makes the *D*_GEN_ method sensitive to graph parameters such as branch lengths and coalescence rates within branches of the species tree (recall that it is in general ambiguous which of the patterns BAABA and ABABA become more common under gene flow 3 → 2 in tree *S*), so *D*_GEN_ cannot be considered a true generalization of the classic *D*-statistic.

Finally, using an empirical example of brown and polar bear admixture that has been thoroughly explored over the last 10 years, we find support for our own principal conclusions (Lan et al., 2022) that admixture between the two species has been most evidently bidirectional following their initial split into distinct species. Furthermore, using Δ-statistics with window sizes varying by a factor of 10, we uncovered what appear to be at least two episodes of gene flow that differed in their directional biases, only the most recent bias favoring the frequently cited hypothesis of polar bear into ABC brown bear introgression.

## Acknowledgements

We are grateful for useful feedback from Dr. Jeffery Ross-Ibarra. The authors acknowledge financial support from U.S. National Science Foundation awards 2221988 and 2139311 to C.L. and 2030871 to V.A.A. K.L. was supported by Academy of Finland Center of Excellence in Tree Biology (TreeBio AoF CoE 346139).

