## Appendix for "Five-leaf generalizations of the *D*-statistic reveal the directionality of admixture": Appendix.pdf

### Appendix 1: allele frequency based formulae

We list with only a sketch proof the allele frequency based formulae of the 20 preferred (scaled) $\Delta$-statistics, in the style $D$- and $D_{\mathrm{FOIL}}$-statistics are presented in Durand, Patterson, Reich, & Slatkin, 2011 and Harris & DeGiorgio, 2017. Let $a_j, b_j, c_j, d_j$ and $e_j$ denote the allele frequencies in populations $1, 2, 3, 4$ and $5$, respectively, of the variant $j \in \{1, 2, \dots, J\}$. We make a slight approximation

$$\mathbb{E}\left(\frac{n(L) - n(R)}{n(L) + n(R)}\right) \approx \frac{\mathbb{E}\big(n(L) - n(R)\big)}{\mathbb{E}\big(n(L) + n(R)\big)},$$

where $L$ and $R$ are sets of allelic patterns. Then, for example,

$$\mathbb{E}(n(\mathrm{ABBAB})) = \sum_{j=1}^{J} \Big( a_j(1-b_j)(1-c_j)d_j(1-e_j) + (1-a_j)b_jc_j(1-d_j)e_j \Big).$$

Using this observation to the sets $L$ and $R$ of definitions (3), (4) and (5) and simplifying yields:

$$
\begin{aligned}
{}_S\Delta^*_{1-6} &= \frac{\sum_{j=1}^{J}(c_j - d_j)(a_j - e_j)}{\sum_{j=1}^{J}(c_j + d_j - 2c_jd_j)(a_j + e_j - 2a_je_j)}, \\
{}_S\Delta^*_{2-6} &= \frac{\sum_{j=1}^{J}(c_j - d_j)(b_j - e_j)}{\sum_{j=1}^{J}(c_j + d_j - 2c_jd_j)(b_j + e_j - 2b_je_j)}, \\
{}_S\Delta^*_{3-5} &= \frac{\sum_{j=1}^{J}(a_j - b_j)(c_j - e_j)}{\sum_{j=1}^{J}(a_j + b_j - 2a_jb_j)(c_j + e_j - 2c_je_j)}, \\
{}_S\Delta^*_{4-5} &= \frac{\sum_{j=1}^{J}(a_j - b_j)(d_j - e_j)}{\sum_{j=1}^{J}(a_j + b_j - 2a_jb_j)(d_j + e_j - 2d_je_j)}, \\
{}_S\Delta^*_{5+7} &= \frac{\sum_{j=1}^{J}(a_j - b_j)(1 - c_j - d_j)}{\sum_{j=1}^{J}(a_j + b_j - 2a_jb_j)(1 - c_j - d_j + 2c_jd_j)}, \\
{}_S\Delta^*_{6+8} &= \frac{\sum_{j=1}^{J}(c_j - d_j)(1 - a_j - b_j)}{\sum_{j=1}^{J}(c_j + d_j - 2c_jd_j)(1 - a_j - b_j + 2a_jb_j)}, \\
{}_S\Delta^*_{3+4-5+7} &= \frac{\sum_{j=1}^{J}(a_j - b_j)(1 - 2e_j)}{\sum_{j=1}^{J}(a_j + b_j - 2a_jb_j)}, \\
{}_S\Delta^*_{1+2-6+8} &= \frac{\sum_{j=1}^{J}(c_j - d_j)(1 - 2e_j)}{\sum_{j=1}^{J}(c_j + d_j - 2c_jd_j)}, \\
{}_A\Delta^*_{1-2} &= \frac{\sum_{j=1}^{J}(a_j - b_j)(c_j - d_j)}{\sum_{j=1}^{J}(a_j + b_j - 2a_jb_j)(c_j + d_j - 2c_jd_j)}, \\
{}_A\Delta^*_{1-3} &= \frac{\sum_{j=1}^{J}(a_j - b_j)(c_j - e_j)}{\sum_{j=1}^{J}(a_j + b_j - 2a_jb_j)(c_j + e_j - 2c_je_j)}, \\
{}_A\Delta^*_{2-3} &= \frac{\sum_{j=1}^{J}(a_j - b_j)(d_j - e_j)}{\sum_{j=1}^{J}(a_j + b_j - 2a_jb_j)(d_j + e_j - 2d_je_j)}, \\
{}_A\Delta^*_{1+2-3+4} &= \frac{\sum_{j=1}^{J}(a_j - b_j)(1 - 2e_j)}{\sum_{j=1}^{J}(a_j + b_j - 2a_jb_j)},
\end{aligned}
$$

$$ {}_Q\Delta^*_{1-6} = \frac{\sum_{j=1}^J (d_j - e_j)(a_j - c_j)}{\sum_{j=1}^J (d_j + e_j - 2d_j e_j)(a_j + c_j - 2a_j c_j)} , $$

$$ {}_Q\Delta^*_{2-6} = \frac{\sum_{j=1}^J (d_j - e_j)(b_j - c_j)}{\sum_{j=1}^J (d_j + e_j - 2d_j e_j)(b_j + c_j - 2b_j c_j)} , $$

$$ {}_Q\Delta^*_{3-5} = \frac{\sum_{j=1}^J (a_j - b_j)(d_j - c_j)}{\sum_{j=1}^J (a_j + b_j - 2a_j b_j)(d_j + c_j - 2d_j c_j)} , $$

$$ {}_Q\Delta^*_{4-5} = \frac{\sum_{j=1}^J (a_j - b_j)(e_j - c_j)}{\sum_{j=1}^J (a_j + b_j - 2a_j b_j)(e_j + c_j - 2e_j c_j)} , $$

$$ {}_Q\Delta^*_{5+7} = \frac{\sum_{j=1}^J (a_j - b_j)(1 - d_j - e_j)}{\sum_{j=1}^J (a_j + b_j - 2a_j b_j)(1 - d_j - e_j + 2d_j e_j)} , $$

$$ {}_Q\Delta^*_{6+8} = \frac{\sum_{j=1}^J (d_j - e_j)(1 - a_j - b_j)}{\sum_{j=1}^J (d_j + e_j - 2d_j e_j)(1 - a_j - b_j + 2a_j b_j)} , $$

$$ {}_Q\Delta^*_{3+4-5+7} = \frac{\sum_{j=1}^J (a_j - b_j)(1 - 2c_j)}{\sum_{j=1}^J (a_j + b_j - 2a_j b_j)} , $$

$$ {}_Q\Delta^*_{1+2-6+8} = \frac{\sum_{j=1}^J (d_j - e_j)(1 - 2c_j)}{\sum_{j=1}^J (d_j + e_j - 2d_j e_j)} , $$

These formulae make it very noticeable how the statistics ${}_S\Delta^*_{1-6}, {}_S\Delta^*_{2-6}, {}_S\Delta^*_{3-5}$ and ${}_S\Delta^*_{4-5}$ are in fact just classic $D$-statistics where one of the populations 1–4 is ignored, and how the statistics ${}_S\Delta^*_{3+4-5+7}$ and ${}_S\Delta^*_{1+2-6+8}$ are formally equivalent to $D_{\text{FOIL}}$-statistics where population 5 has assumed the role of one of the populations 1–4. We stress that while the allele frequency based formulae encourage to use the perspective of drift instead of the perspective of coalescence, simply choosing to use them does not make the synchronization assumption (2) true. For instance, from the perspective of drift alone, we might get the wrong impression that the expectation of ${}_S\Delta^*_{3+4-5+7}$ is zero even when 2 is an ancient sample, as in Figure 2 Panel III) of the main paper. The terminal branch leading to 1 is longer than that leading to 2, but we still we expect $a_j - b_j$ to be zero and independent of $1 - 2e_j$. But this is only because the drift perspective smuggled in an assumption that the locus was already polymorphic at the root, ignoring the possibility of a recent mutation on the terminal branches altogether. Assuming that all the mutations happened after the root is a special case of making the synchronization assumption (2), as at that moment all the lineages are in the same population.

#### Appendix 2: derivation of the expected Δ-statistics

We adapt a Newick notation where only leaf nodes are named, and their names are sets consisting of one or more (genetic) lineages. For example, by $(\{1,2\},\{3\})$ we mean a situation where lineages 1 and 2 are sampled from one population, and lineage 3 from another.

In this example, the allelic patterns BAA and ABA have the same probability by symmetry, and provided that the synchronization assumption (2) is valid, the probability of AAB is higher. Indeed, if lineages 1 and 2 coalesce before the two populations merge, and if the mutation happened after this coalescence event, then BAA and ABA are impossible while AAB is not; otherwise BAA and ABA have the same probability that under (2) equals the probability of AAB. To denote the use of the synchronization assumption (2) we shall use the asterisk, in this case:

$$0 < \mathbb{P}(\mathrm{AAB}) \overset{*}{>} \mathbb{P}(\mathrm{BAA}) = \mathbb{P}(\mathrm{ABA}) > 0\,.$$

In the next section we expand the logic of this toy example into eight lemmas. The toy example is not codified as a lemma of its own, but it will be repeatedly used so the reader might want to think of it as "Lemma 0". The statements of the lemmas are somewhat stronger than is necessary for this work, for future reference.

The full proof of the validity of Tables 2, 3 and 4 is long. The predictions are verified using the eight lemmas in the subsequent three sections. Most cases are reduced to figures, equations and tables — only at the start we give full explanations. Cases that can be obtained from others by permutations of the leaves have been omitted.

#### Lemmas

**Lemma 1.** *Suppose* $(\{1,2,3,4\})$ *and denote*

$$\begin{cases} \mathbb{P}(\text{BAAA}) = \mathbb{P}(\text{ABAA}) = \mathbb{P}(\text{AABA}) = \mathbb{P}(\text{AAAB}) = Y\,, \\ \mathbb{P}(\text{BBAA}) = \mathbb{P}(\text{BABA}) = \mathbb{P}(\text{BAAB}) = Z\,. \end{cases}$$

*Then* $Y > 1.5Z > 0$.

*Proof.* We first think of the coalescent process at a locus conditional on exactly one mutation somewhere in the gene tree. No matter what the gene tree happens to be exactly (the lineages coalescing, number of generations between coalescence events), there are 17 other gene trees that had the exact same probability of occurring (see figure). Assuming the gene tree is one of these 18, let $a$ be the probability that the mutation happened before the first coalescence event, $b$ the probability it happened between the first and the second coalescence events, and $c$ the probability it happened between the second and the last coalescence events. As shown in the figure, $Y$ is then one eighteenth times $18a + 9b + 6c$, while $Z$ is one eighteenth times $6b + 4c$. For all positive values of $a$, $b$ and $c$ (i.e., for whichever class of 18 trees considered) it is always true that $Y > 1.5Z > 0$. Note that since all the lineages are in the same population, the synchronization assumption (2) is automatically valid — the mutation rate can freely vary along the time-axis of the figure. ☐

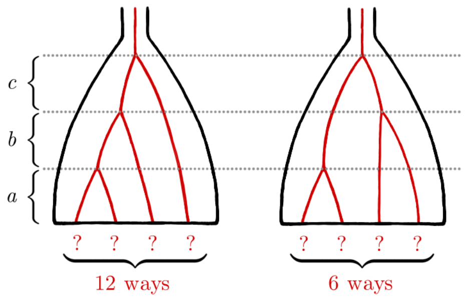


$$\begin{cases} Y = & 6a{+}3(a{+}b) \;{+}3(a{+}b{+}2c) & {+}3a{+}3(a{+}b) & = 18a{+}9b{+}6c \\ Z = & 4b & {+}2(b{+}2c) & = 6b{+}4c \end{cases}$$

**Lemma 2.** *Suppose* $(\{1,2,3,4,5\})$ *and denote*

$$\begin{cases} \mathbb{P}(\mathrm{BAAAA}) = \mathbb{P}(\mathrm{ABAAA}) = \mathbb{P}(\mathrm{AABAA}) = \mathbb{P}(\mathrm{AAABA}) = \mathbb{P}(\mathrm{AAAAB}) = Y\,, \\ \mathbb{P}(\mathrm{BBAAA}) = \mathbb{P}(\mathrm{BABAA}) = \mathbb{P}(\mathrm{BAABA}) = \mathbb{P}(\mathrm{BAAAB}) = \mathbb{P}(\mathrm{ABBAA}) = \mathbb{P}(\mathrm{ABABA}) \\ \qquad = \mathbb{P}(\mathrm{ABAAB}) = \mathbb{P}(\mathrm{AABBA}) = \mathbb{P}(\mathrm{AABAB}) = \mathbb{P}(\mathrm{AAABB}) = Z\,. \end{cases}$$

*Then* $Y > 2Z > 0$.

*Proof.* The proof is analogous to the proof of Lemma 1. ☐

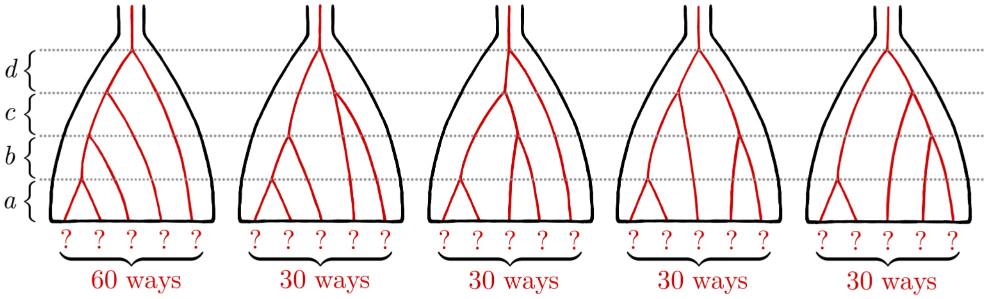


$$\begin{cases} Y = \begin{array}{lllll} 24a+12(a+b) & +12a & +12a & +12a & +12a \\ +12(a+b+c) & +6(a+b) & +12(a+b) & +6(a+b+c) & +6(a+b+c) \\ +12(a+b+c+2d) & +12(a+b+c) & +6(a+b+c+2d) & +12(a+b) & +12(a+b) \end{array} = \begin{array}{l} 180a+108b \\ +54c+36d \end{array} \\ \\ Z = \begin{array}{lllll} 6b & +3b & +3(b+c) & +3(b+c) & +3(b+c+2d) \\ +6c & +3(c+2d) & +3c & +3(c+2d) & +3c \end{array} = \begin{array}{l} 18b+27c \\ +18d \end{array} \end{cases}$$

**Lemma 3.** *Suppose* $(\{1,2,3\},\{4\})$ *and denote*

$$\begin{cases} \mathbb{P}(\mathrm{AAAB}) = X\,, \\ \mathbb{P}(\mathrm{BAAA}) = \mathbb{P}(\mathrm{ABAA}) = \mathbb{P}(\mathrm{AABA}) = Y\,, \\ \mathbb{P}(\mathrm{BBAA}) = \mathbb{P}(\mathrm{BABA}) = \mathbb{P}(\mathrm{BAAB}) = Z\,. \end{cases}$$

*Then* $X > 0 < Z < Y$ *and* $Y \overset{*}{<} X \overset{*}{>} 1.5Z$.
*The inequalities marked with an asterisk require the synchronization assumption (2).*

*Proof.* Define six distinct events denoted by Roman numerals I) – VI) by the number of coalescence events happening before the merging of the two populations, and by whether or not the only mutation that occurs happens before or after this merging (see figure). For example, event I) means that no coalescence events nor the mutation happened until the merging, so that Lemma 1 applies. The probabilities of events I) – VI) are unknown and positive. What is known about the numbers $X$, $Y$ and $Z$ conditional on each event I) – VI) is shown in the figure. In events V) and VI) we use the logic applied in Lemma 1: for each gene tree there are two others that have the exact same probability, and these three are dealt with jointly. Note that we now have two distinct populations until the merging, so we need the synchronization assumption (2) to compare $X$ to other numbers $Y$ and $Z$. The steps that use (2) are marked with an asterisk.

In each event I) – VI), whatever the positive numbers $a - e$ are, we have $X > 0 \leq Z \leq Y$ even without using the synchronization assumption (2). Moreover, $Z > 0$ in events I), II), V) and VI), and since all events have strictly positive probabilities, $0 < Z$ in general. Likewise, $Z < Y$ in events I), IV), V) and VI), so $Z < Y$ in general.

Similarly, to show that $Y \overset{*}{<} X \overset{*}{>} 1.5Z$, we again check event-by-event, but this time allowing the use of equations marked with an asterisk. ☐

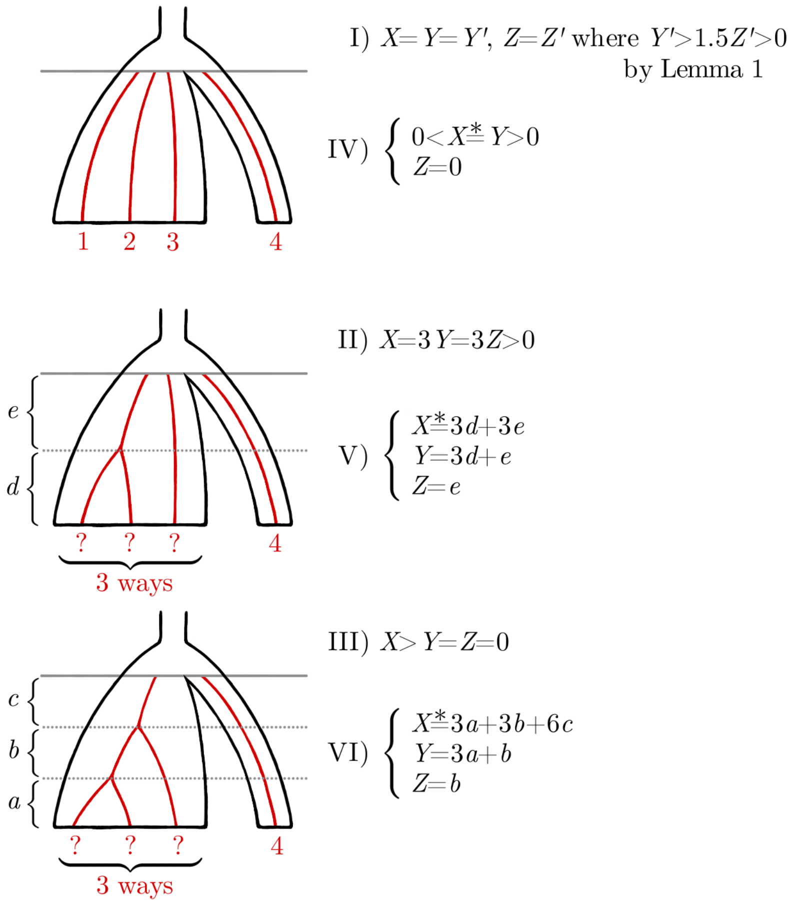

**Lemma 4.** *Suppose* $(\{1,2,3,4\},\{5\})$ *and denote*

$$\begin{cases}\mathbb{P}(\text{AAAAB}) = W\,,\\ \mathbb{P}(\text{BAAAA}) = \mathbb{P}(\text{ABAAA}) = \mathbb{P}(\text{AABAA}) = \mathbb{P}(\text{AAABA}) = X\,,\\ \mathbb{P}(\text{BAAAB}) = \mathbb{P}(\text{ABAAB}) = \mathbb{P}(\text{AABAB}) = \mathbb{P}(\text{AAABB}) = Y\,,\\ \mathbb{P}(\text{BBAAA}) = \mathbb{P}(\text{BABAA}) = \mathbb{P}(\text{BAABA}) = \mathbb{P}(\text{ABBAA}) = \mathbb{P}(\text{ABABA}) = \mathbb{P}(\text{AABBA}) = Z\,.\end{cases}$$

*Then* $W > 0 < Y < X > 1.5Z > 0$, $X \overset{*}{<} W \overset{*}{>} 2Y$ *and* $W + Y \overset{*}{>} X + Z$.
*The inequalities marked with an asterisk require the synchronization assumption (2).*

*Proof.* The proof is analogous to the proof of Lemma 3. □

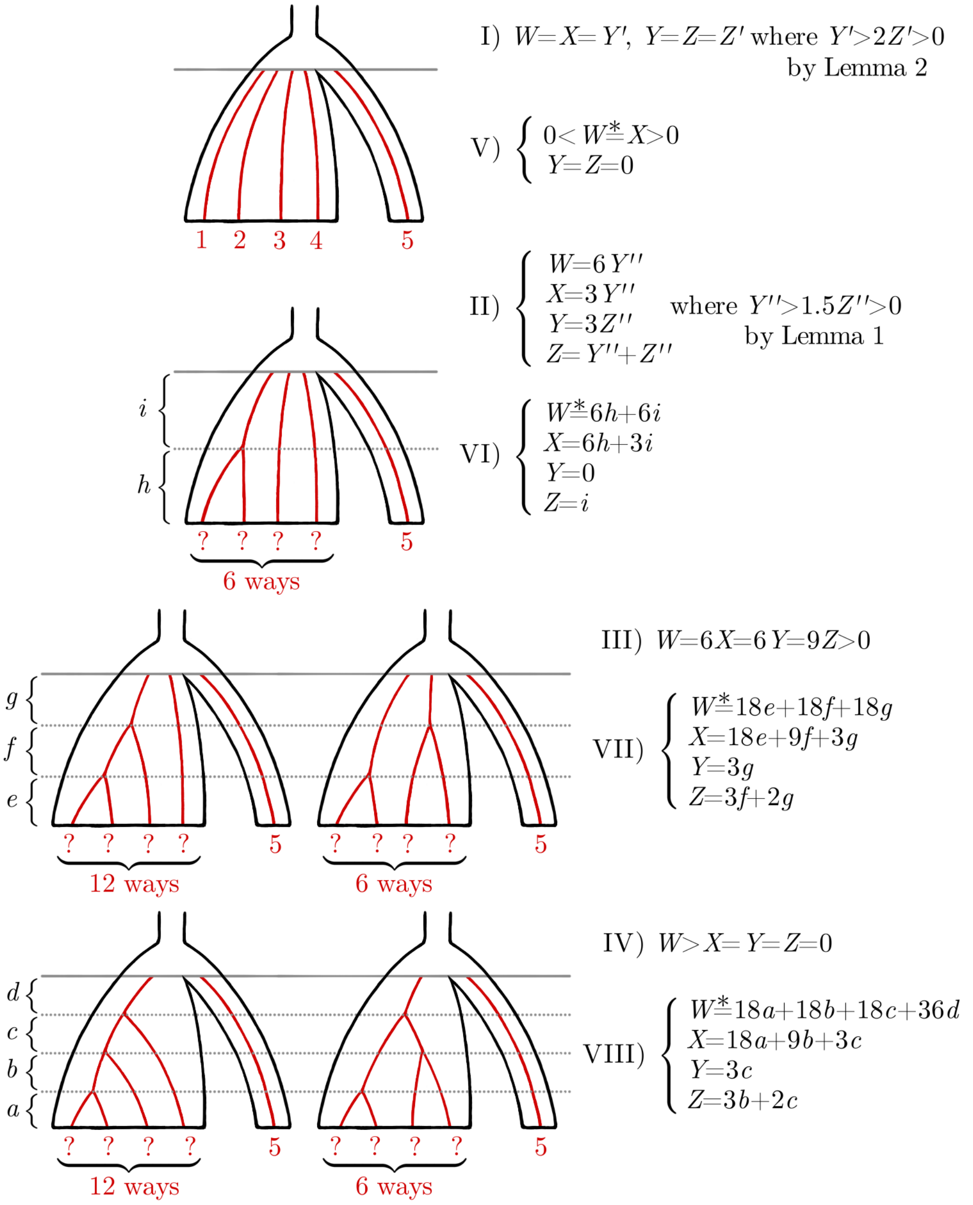

**Lemma 5.** *Suppose* $((\{1,2\},\{3\}),\{4\})$ *and denote*

$$\begin{cases} \mathbb{P}(\mathrm{AAAB}) = V\,, \\ \mathbb{P}(\mathrm{AABA}) = W\,, \\ \mathbb{P}(\mathrm{BAAA}) = \mathbb{P}(\mathrm{ABAA}) = X\,, \\ \mathbb{P}(\mathrm{BBAA}) = Y\,, \\ \mathbb{P}(\mathrm{ABBA}) = \mathbb{P}(\mathrm{BABA}) = Z\,. \end{cases}$$

*Then*

$$\begin{cases} V > Z < W \\ X > Z < Y \\ Z > 0 \end{cases} \qquad \textit{and} \qquad \begin{cases} W \overset{*}{<} V \overset{*}{>} 1.5Z \\ X \overset{*}{<} W \overset{*}{>} Y \\ W + Z \overset{*}{=} X + Y \end{cases} .$$

*The relations marked with an asterisk require the synchronization assumption (2).*

*Proof.* The proof is analogous to the proof of Lemma 3. □

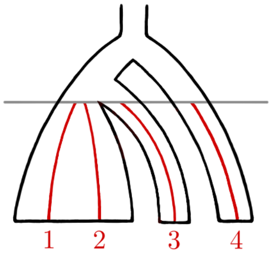


I) $\begin{cases} V = X' \\ W = X = Y' \\ Y = Z = Z' \end{cases}$ where $\begin{cases} X' > 0 < Z' < Y' \\ Y' \overset{*}{<} X' \overset{*}{>} 1.5Z' \end{cases}$ by Lemma 3

III) $\begin{cases} 0 < V \overset{*}{=} W \overset{*}{=} X > 0,\ W > 0 \\ Y = Z = 0 \end{cases}$

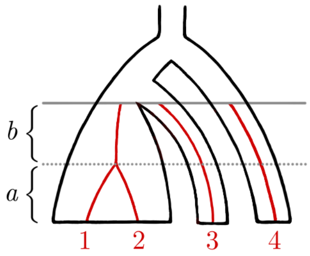


II) $0 < V \overset{*}{>} W = Y > X = Z = 0$

IV) $\begin{cases} V \overset{*}{=} W \overset{*}{=} a + b \\ X = a \\ Y = b \\ Z = 0 \end{cases}$

**Lemma 6.** *Suppose* $((\{1,2,3\},\{4\}),\{5\})$ *and denote*

$$\begin{cases} \mathbb{P}(\mathrm{AAAAB}) = T\,, \\ \mathbb{P}(\mathrm{AAABA}) = U\,, \\ \mathbb{P}(\mathrm{BAAAA}) = \mathbb{P}(\mathrm{ABAAA}) = \mathbb{P}(\mathrm{AABAA}) = V\,, \\ \mathbb{P}(\mathrm{AAABB}) = W\,, \\ \mathbb{P}(\mathrm{BAAAB}) = \mathbb{P}(\mathrm{ABAAB}) = \mathbb{P}(\mathrm{AABAB}) = X\,, \\ \mathbb{P}(\mathrm{BBAAA}) = \mathbb{P}(\mathrm{BABAA}) = \mathbb{P}(\mathrm{ABBAA}) = Y\,, \\ \mathbb{P}(\mathrm{BAABA}) = \mathbb{P}(\mathrm{ABABA}) = \mathbb{P}(\mathrm{AABBA}) = Z\,. \end{cases}$$

*Then*

$$\begin{cases} U > 0 < X < W \\ X < V > Y > Z > 0 \end{cases} \qquad \textit{and} \qquad \begin{cases} 1.5Y \overset{*}{<} U \overset{*}{<} T \overset{*}{>} 2X \\ W - X \overset{*}{<} U - V \overset{*}{>} Y - Z \\ T + X \overset{*}{>} U + Z \\ T + W \overset{*}{>} V + Z \\ U + X + 2Z \overset{*}{=} V + W + 2Y \overset{*}{<} T + Z + 2X \end{cases}.$$

*The relations marked with an asterisk require the synchronization assumption (2).*

*Proof.* The proof is analogous to the proof of Lemma 3. □

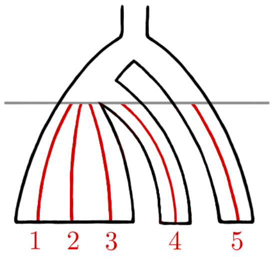


$$\text{I)} \begin{cases} T = W' \\ U = V = X' \\ W = X = Y' \\ Y = Z = Z' \end{cases} \quad \text{where} \quad \begin{cases} W' > 0 < Y' < X' > 1.5Z' > 0 \\ X' \overset{*}{<} W' \overset{*}{>} 2Y' \\ W' + Y' \overset{*}{>} X' + Z' \end{cases} \text{ by Lemma 4}$$

$$\text{IV)} \begin{cases} 0 < T \overset{*}{=} U \overset{*}{=} V > 0, \ U > 0 \\ W = X = Y = Z = 0 \end{cases}$$

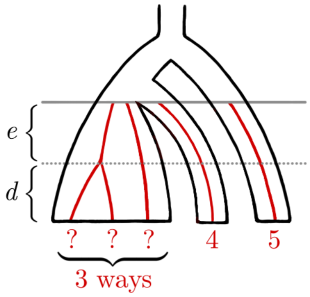


$$\text{II)} \begin{cases} T = 3X'' \\ U = 3Y'' \\ V = Y = Y'' \\ W = 3Z'' \\ X = Z = Z'' \end{cases} \quad \text{where} \quad \begin{cases} X'' > 0 < Z'' < Y'' \\ Y'' \overset{*}{<} X'' \overset{*}{>} 1.5Z'' \end{cases} \text{ by Lemma 3}$$

$$\text{V)} \begin{cases} T \overset{*}{=} U \overset{*}{=} 3d + 3e \\ V = 3d + e \\ W = X = Z = 0 \\ Y = e \end{cases}$$

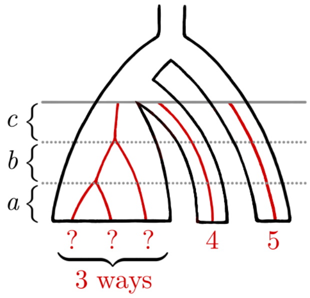


$$\text{III)} \ 0 < T \overset{*}{>} U = W > V = X = Y = Z = 0$$

$$\text{VI)} \begin{cases} T \overset{*}{=} U \overset{*}{=} 3a + 3b + 3c \\ V = 3a + b \\ W = 3c \\ X = Z = 0 \\ Y = b \end{cases}$$

**Lemma 7.** *Suppose* $(\{1,2\},\{3,4\})$ *and denote*

$$\begin{cases} \mathbb{P}(\mathrm{AABA}) = \mathbb{P}(\mathrm{AAAB}) = W\,, \\ \mathbb{P}(\mathrm{BAAA}) = \mathbb{P}(\mathrm{ABAA}) = X\,, \\ \mathbb{P}(\mathrm{BBAA}) = Y\,, \\ \mathbb{P}(\mathrm{BABA}) = \mathbb{P}(\mathrm{BAAB}) = Z\,. \end{cases}$$

*Then* $W > 1.5Z < X$, $Y > Z > 0$ *and* $W - X \overset{*}{<} Y - Z \overset{*}{>} X - W$.
*The inequalities marked with an asterisk require the synchronization assumption (2).*

*Proof.* The proof is analogous to the proof of Lemma 3. □

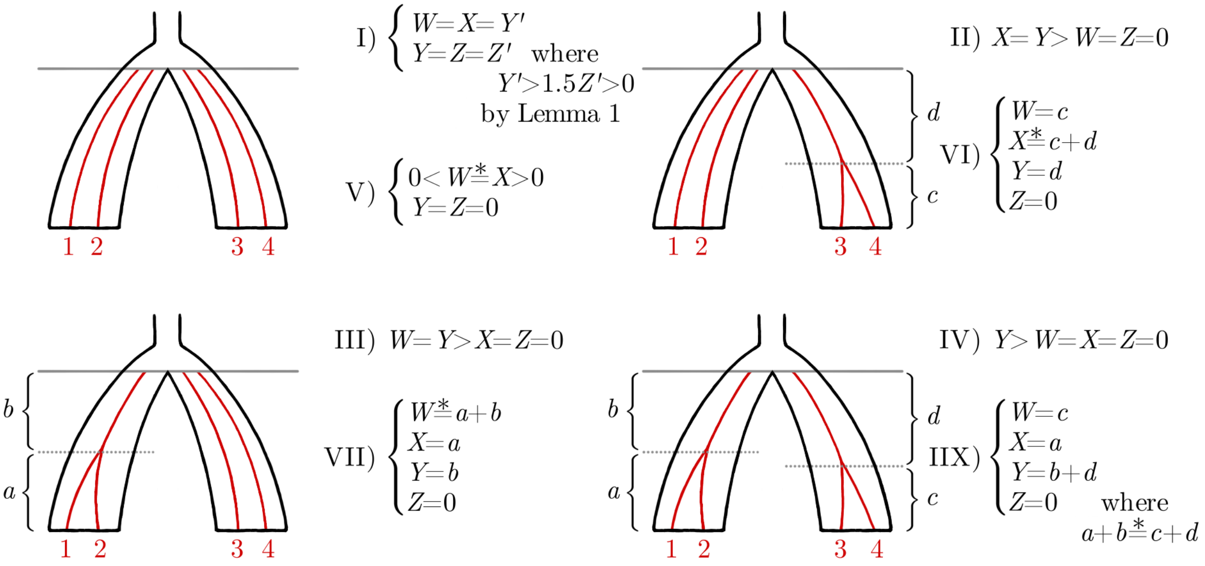

**Lemma 8.** *Suppose* $(\{1,2,3\},\{4,5\})$ *and denote*

$$\begin{cases} \mathbb{P}(\mathrm{AAABA}) = \mathbb{P}(\mathrm{AAAAB}) = V\,, \\ \mathbb{P}(\mathrm{BAAAA}) = \mathbb{P}(\mathrm{ABAAA}) = \mathbb{P}(\mathrm{AABAA}) = W\,, \\ \mathbb{P}(\mathrm{AAABB}) = X\,, \\ \mathbb{P}(\mathrm{BBAAA}) = \mathbb{P}(\mathrm{BABAA}) = \mathbb{P}(\mathrm{ABBAA}) = Y\,, \\ \mathbb{P}(\mathrm{BAABA}) = \mathbb{P}(\mathrm{BAAAB}) = \mathbb{P}(\mathrm{ABABA}) = \mathbb{P}(\mathrm{ABAAB}) = \mathbb{P}(\mathrm{AABBA}) = \mathbb{P}(\mathrm{AABAB}) = Z\,. \end{cases}$$

*Then*

$$\begin{cases} 3V > 6Z < 4W \\ X > Z < Y \\ Z > 0 \end{cases} \quad \text{and} \quad \begin{cases} V + X \overset{*}{>} W + Z \\ V + 3Z \overset{*}{<} W + X + 2Y \end{cases}.$$

*The inequalities marked with an asterisk require the synchronization assumption (2).*

*Proof.* The proof is analogous to the proof of Lemma 3. □

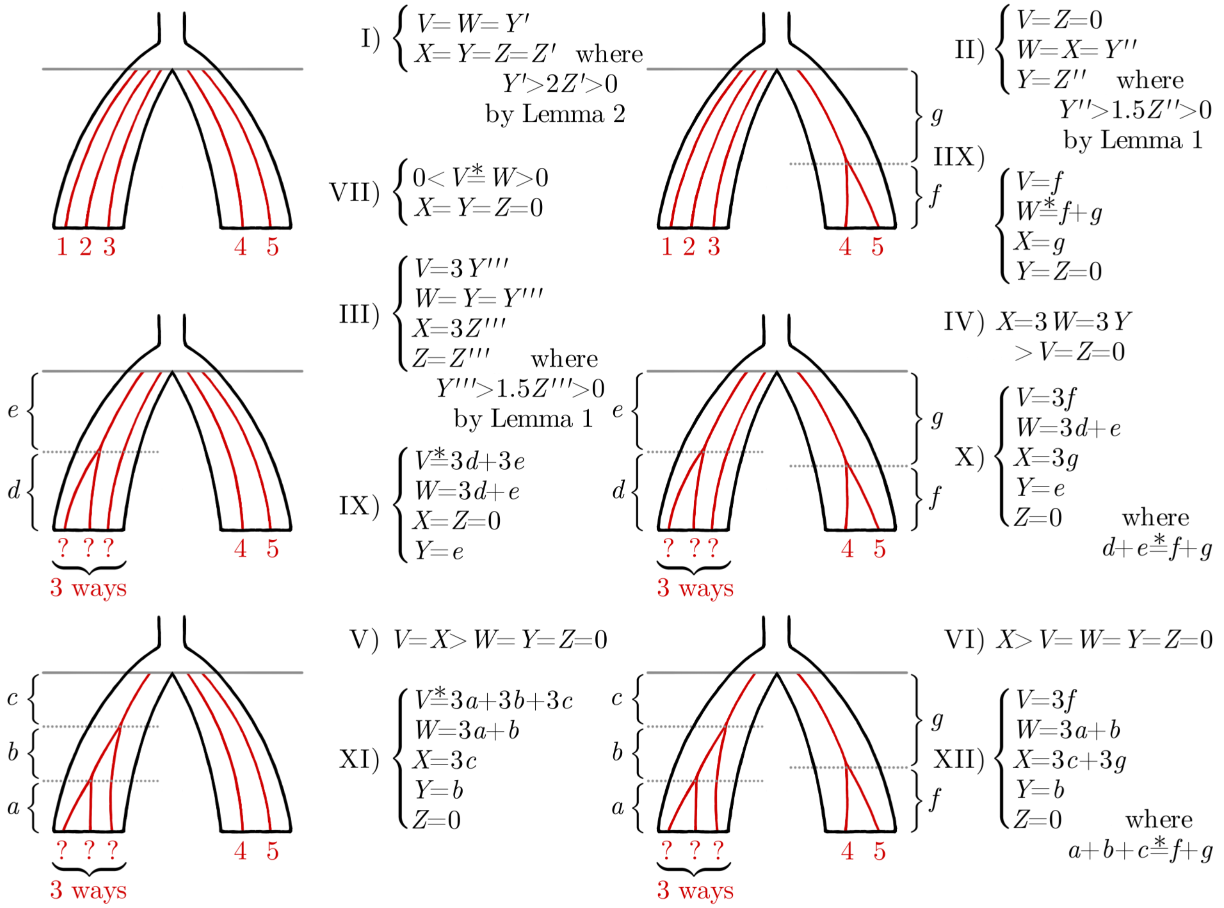

#### Symmetric tree

As shown in the main text, all the binomial $\Delta$-statistics are expected to equal zero when there are no admixture events, but for ${}_S\Delta_7$ and ${}_S\Delta_8$ we need to assume (2). We mark this relation down as equation ($S0$) — keeping in mind that an asterisk stands for the synchronization assumption (2).

$$(S0) \quad \begin{cases} {}_S\Delta_7 \overset{*}{=} {}_S\Delta_8 \overset{*}{=} {}_S\Delta_1 = {}_S\Delta_2 = {}_S\Delta_3 = {}_S\Delta_4 = {}_S\Delta_5 = {}_S\Delta_6 = 0 \end{cases}$$

**3 → 2.**
Assume that lineage 2 is introgressed at the locus; otherwise ($S0$) applies. Define four distinct events I) – IV) of unknown probabilities by whether lineages 2 and 3 coalesce and whether the mutation occurs before the merging of the populations that include lineages 3 and 4. Relevant relations between the binomial $\Delta$-statistics ${}_S\Delta_1$ – ${}_S\Delta_8$ in each event I) – IV) are written down in relations ($S0$) – ($S3$). Note that the sign of ${}_S\Delta_4$ is not consistent: in event I) ${}_S\Delta_4 < 0$ by ($S1$) and in event II) ${}_S\Delta_4 > 0$ by ($S2$), otherwise ${}_S\Delta_4 = 0$.

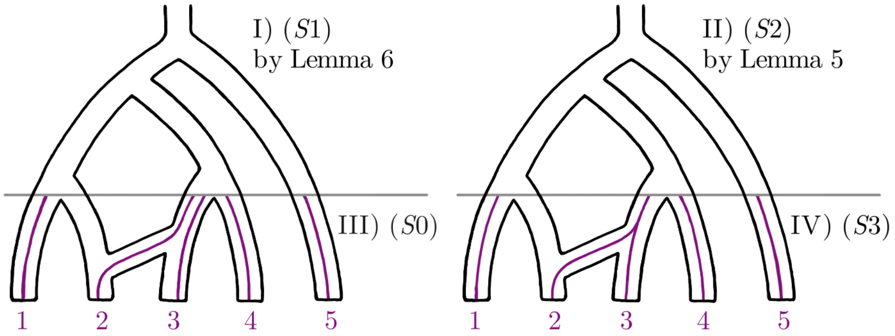


$$(S1) \quad \begin{cases} {}_S\Delta_1 = {}_S\Delta_2 = {}_S\Delta_6 = {}_S\Delta_8 = 0 \\ {}_S\Delta_3 = {}_S\Delta_4 = Z - Y \\ {}_S\Delta_5 = W - X \\ {}_S\Delta_7 = U - V \end{cases} \quad \text{where} \quad \begin{cases} U \overset{*}{>} V \\ W > X \\ Y > Z \\ U + X + 2Z \overset{*}{=} V + W + 2Y \end{cases}$$

$$(S2) \quad \begin{cases} -{}_S\Delta_1 = {}_S\Delta_4 = -{}_S\Delta_6 = Z \\ {}_S\Delta_2 = -{}_S\Delta_3 = -{}_S\Delta_8 = X \\ {}_S\Delta_5 = Y \\ {}_S\Delta_7 = W \end{cases} \quad \text{where} \quad \begin{cases} 0 < W \overset{*}{>} X > 0 \\ Y > Z > 0 \\ W + Z \overset{*}{=} X + Y \end{cases}$$

$$(S3) \quad \begin{cases} {}_S\Delta_1 = {}_S\Delta_4 = {}_S\Delta_5 = {}_S\Delta_6 = 0 \\ {}_S\Delta_7 \overset{*}{=} -{}_S\Delta_8 \overset{*}{=} {}_S\Delta_2 = -{}_S\Delta_3 > 0 \end{cases}$$

We then tabulate the signs of our eight preferred (unscaled) $\Delta$-statistics under all the relations ($S0$) – ($S3$), marking down an asterisk when the result requires assumption (2). Because the probabilities of events I) – IV) are positive, the expected sign will be zero if it's always (in all ($S0$) – ($S3$)) zero, positive if it's positive at least sometimes and never negative, and negative when it's negative at least sometimes and never positive. Notice that the behaviour of ${}_S\Delta_{1-6}$, ${}_S\Delta_{2-6}$, ${}_S\Delta_{3-5}$ and ${}_S\Delta_{4-5}$ is known without having to resort to the synchronization assumption (2).

| | ${}_S\Delta_{1-6}$ | ${}_S\Delta_{2-6}$ | ${}_S\Delta_{3-5}$ | ${}_S\Delta_{4-5}$ | ${}_S\Delta_{5+7}$ | ${}_S\Delta_{6+8}$ | ${}_S\Delta_{3+4-5+7}$ | ${}_S\Delta_{1+2-6+8}$ |
|---|---|---|---|---|---|---|---|---|
| ($S0$) | 0 | 0 | 0 | 0 | $0^*$ | $0^*$ | $0^*$ | $0^*$ |
| ($S1$) | 0 | 0 | − | − | $+^*$ | 0 | $0^*$ | 0 |
| ($S2$) | 0 | + | − | − | + | − | $0^*$ | 0 |
| ($S3$) | 0 | + | − | 0 | $+^*$ | $-^*$ | $0^*$ | $0^*$ |
| 3 → 2 | 0 | + | − | − | $+^*$ | $-^*$ | $0^*$ | $0^*$ |

**12345 → 4.**
The proof is analogous to the previous proof.

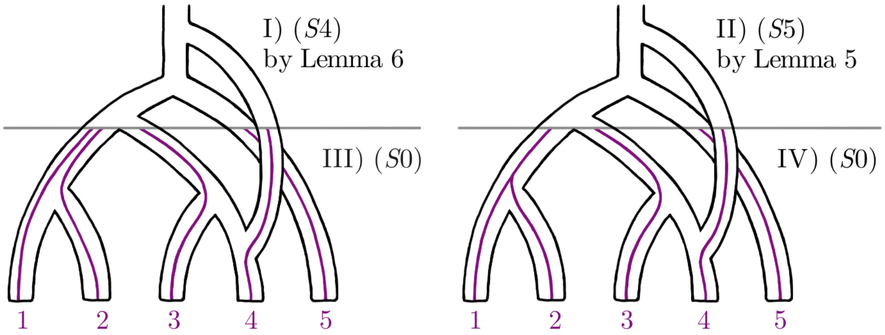


$$(S4)\quad \begin{cases} {}_S\Delta_1 = {}_S\Delta_2 = Y - X \\ {}_S\Delta_3 = {}_S\Delta_4 = {}_S\Delta_5 = {}_S\Delta_7 = 0 \\ {}_S\Delta_6 = Z - W \\ {}_S\Delta_8 = V - T \end{cases} \quad \text{where} \quad \begin{cases} W > X \\ Y > Z \\ T + W \overset{*}{>} V + Z \\ T + Z + 2X \overset{*}{>} W + V + 2Y \end{cases}$$

$$(S5)\quad \begin{cases} {}_S\Delta_1 = {}_S\Delta_2 = {}_S\Delta_3 = {}_S\Delta_4 = {}_S\Delta_5 = {}_S\Delta_7 = 0 \\ {}_S\Delta_6 = Z - Y \\ {}_S\Delta_8 = X - V \end{cases} \quad \text{where} \quad \begin{cases} V \overset{*}{>} X \\ Y > Z \\ V + Z \overset{*}{>} X + Y \end{cases}$$

| | ${}_S\Delta_{1-6}$ | ${}_S\Delta_{2-6}$ | ${}_S\Delta_{3-5}$ | ${}_S\Delta_{4-5}$ | ${}_S\Delta_{5+7}$ | ${}_S\Delta_{6+8}$ | ${}_S\Delta_{3+4-5+7}$ | ${}_S\Delta_{1+2-6+8}$ |
|---|---|---|---|---|---|---|---|---|
| $(S0)$ | 0 | 0 | 0 | 0 | $0^*$ | $0^*$ | $0^*$ | $0^*$ |
| $(S4), (S5)$ | + | + | 0 | 0 | 0 | $-^*$ | 0 | $-^*$ |
| $12345 \to 4$ | + | + | 0 | 0 | $0^*$ | $-^*$ | $0^*$ | $-^*$ |

**3 → 12.**
First assume that only one of the lineages 1 and 2 is introgressed (again, if neither is introgressed, we have ($S0$)). Define four distinct events I) – IV) of positive probability by whether the introgressed lineage coalesces with lineage 3 and whether the single mutation happens before the merging of the populations that include lineages 3 and 4. Because both lineages 1 and 2 have the same chance of being the introgressed one, we deal with gene trees two at a time as was done in Lemmas 1 – 8. This method of summing may warrant clarification, so as an example we look more closely at how ($S7$) follows in event II).

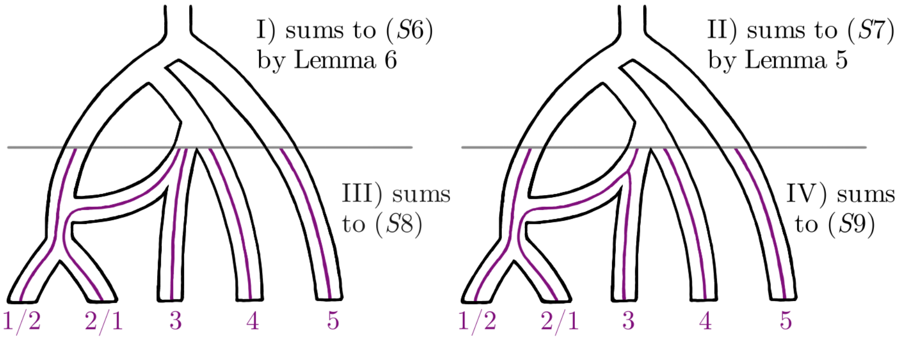


If lineage 2 is the introgressed one, we have

$$\begin{cases} {}_S\Delta_1 = -{}_S\Delta_4 = {}_S\Delta_6 = -Z \\ -{}_S\Delta_2 = {}_S\Delta_3 = {}_S\Delta_8 = -X \\ {}_S\Delta_5 = Y \\ {}_S\Delta_7 = W \end{cases}$$

by Lemma 5. On the other hand, if lineage 1 is the introgressed one, we have

$$\begin{cases} {}_S\Delta_1 = {}_S\Delta_3 = -{}_S\Delta_8 = X \\ {}_S\Delta_2 = {}_S\Delta_4 = {}_S\Delta_6 = -Z \\ {}_S\Delta_5 = -Y \\ {}_S\Delta_7 = -W \end{cases}$$

again by Lemma 5. Both of these sub-events have the same probability, and therefore we obtain ($S7$) as their sum. Relations ($S6$), ($S8$) and ($S9$) are obtained similarly.

$$(S6) \quad \begin{cases} {}_S\Delta_1 = {}_S\Delta_2 = {}_S\Delta_3 = {}_S\Delta_4 = {}_S\Delta_5 = {}_S\Delta_6 = {}_S\Delta_7 = {}_S\Delta_8 = 0 \end{cases}$$

$$(S7) \quad \begin{cases} {}_S\Delta_1 = {}_S\Delta_2 = X - Z \\ {}_S\Delta_3 = {}_S\Delta_4 = {}_S\Delta_5 = {}_S\Delta_7 = 0 \\ {}_S\Delta_6 = -2Z \\ {}_S\Delta_8 = -2X \end{cases} \quad \text{where} \quad X > 0 < Z$$

$$(S8) \quad \begin{cases} {}_S\Delta_8 \overset{*}{=} {}_S\Delta_1 = {}_S\Delta_2 = {}_S\Delta_3 = {}_S\Delta_4 = {}_S\Delta_5 = {}_S\Delta_6 = {}_S\Delta_7 = 0 \end{cases}$$

$$(S9) \quad \begin{cases} -{}_S\Delta_8 \overset{*}{=} 2{}_S\Delta_1 = 2{}_S\Delta_2 > 0 \\ {}_S\Delta_3 = {}_S\Delta_4 = {}_S\Delta_5 = {}_S\Delta_6 = {}_S\Delta_7 = 0 \end{cases}$$

Then assume that both the lineages 1 and 2 are introgressed. The rest of the proof is analogous to earlier proofs. We point out that the cases $12 \to 3$ and $1234 \to 4$ reduce to this second half of the proof, i.e. relations $(S0)$, $(S10)$ and $(S11)$.

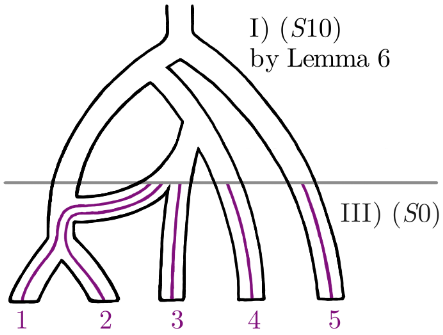


$$(S10) \quad \begin{cases} {}_S\Delta_1 = {}_S\Delta_2 = Y - Z \\ {}_S\Delta_3 = {}_S\Delta_4 = {}_S\Delta_5 = {}_S\Delta_7 = 0 \\ {}_S\Delta_6 = X - W \\ {}_S\Delta_8 = V - U \end{cases} \quad \text{where} \quad \begin{cases} U \overset{*}{>} V \\ W > X \\ Y > Z \\ U + X + 2Z \overset{*}{=} V + W + 2Y \end{cases}$$

$$(S11) \quad \begin{cases} {}_S\Delta_1 = {}_S\Delta_2 = {}_S\Delta_3 = {}_S\Delta_4 = {}_S\Delta_5 = {}_S\Delta_7 = 0 \\ {}_S\Delta_6 = Z - Y \\ {}_S\Delta_8 = X - W \end{cases} \quad \text{where} \quad \begin{cases} W \overset{*}{>} X \\ Y > Z \\ W + Z \overset{*}{=} X + Y \end{cases}$$

| | ${}_S\Delta_{1-6}$ | ${}_S\Delta_{2-6}$ | ${}_S\Delta_{3-5}$ | ${}_S\Delta_{4-5}$ | ${}_S\Delta_{5+7}$ | ${}_S\Delta_{6+8}$ | ${}_S\Delta_{3+4-5+7}$ | ${}_S\Delta_{1+2-6+8}$ |
|---|---|---|---|---|---|---|---|---|
| $(S0)$ | 0 | 0 | 0 | 0 | $0^*$ | $0^*$ | $0^*$ | $0^*$ |
| $(S6)$ | 0 | 0 | 0 | 0 | 0 | 0 | 0 | 0 |
| $(S7)$ | $+$ | $+$ | 0 | 0 | 0 | $-$ | 0 | 0 |
| $(S8)$ | 0 | 0 | 0 | 0 | 0 | $0^*$ | 0 | $0^*$ |
| $(S9)$ | $+$ | $+$ | 0 | 0 | $0^*$ | $-^*$ | $0^*$ | $0^*$ |
| $(S10), (S11)$ | $+$ | $+$ | 0 | 0 | 0 | $-^*$ | 0 | $0^*$ |
| $3 \to 12$ | $+$ | $+$ | 0 | 0 | $0^*$ | $-^*$ | $0^*$ | $0^*$ |

**5 → 4.**

This time we have three binary conditions that define eight distinct events I) – IIX). Depending on the admixture graph, the source of gene flow might actually be after (higher up than) the merging, rendering events II), IV), VI) and IIX) impossible, so we need to make sure the predicted behaviour of 5 → 4 also holds without ($S13$), ($S15$) and ($S16$), and that is the case.

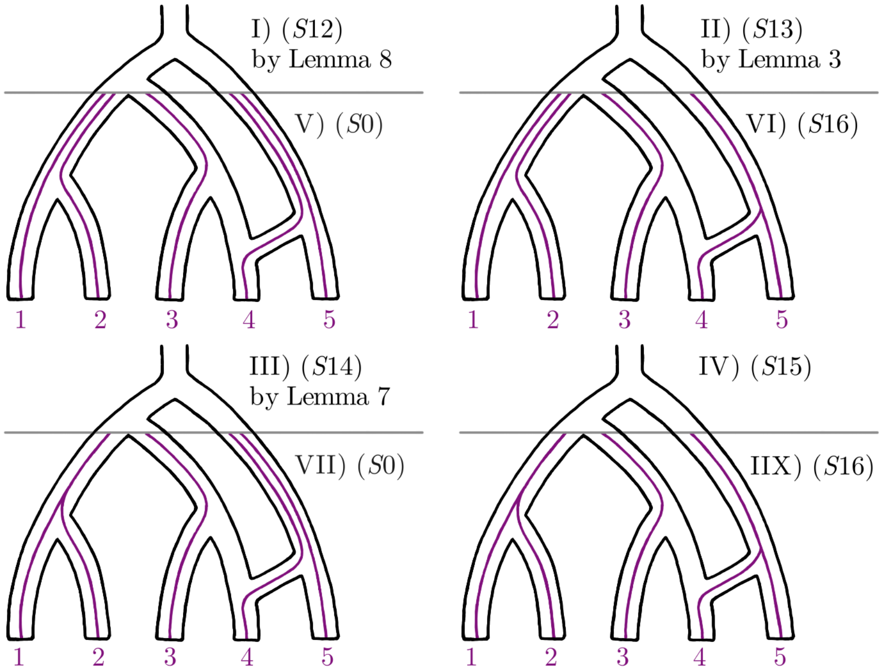


$$(S12) \quad \begin{cases} {}_S\Delta_1 = {}_S\Delta_2 = Y - Z \\ {}_S\Delta_3 = {}_S\Delta_4 = {}_S\Delta_5 = {}_S\Delta_7 = 0 \\ {}_S\Delta_6 = Z - X \\ {}_S\Delta_8 = W - V \end{cases} \quad \text{where} \quad \begin{cases} X > Z < Y \\ V + X \overset{*}{>} W + Z \\ V + 3Z \overset{*}{<} W + X + 2Y \end{cases}$$

$$(S13) \quad \begin{cases} {}_S\Delta_1 = {}_S\Delta_2 = Z \\ {}_S\Delta_3 = {}_S\Delta_4 = {}_S\Delta_5 = {}_S\Delta_7 = 0 \\ {}_S\Delta_6 = -X \\ {}_S\Delta_8 = Y \end{cases} \quad \text{where} \quad 0 < X \overset{*}{>} Y > 0 < Z$$

$$(S14) \quad \begin{cases} {}_S\Delta_1 = {}_S\Delta_2 = {}_S\Delta_3 = {}_S\Delta_4 = {}_S\Delta_5 = {}_S\Delta_7 = 0 \\ {}_S\Delta_6 = Z - Y \\ {}_S\Delta_8 = X - W \end{cases} \quad \text{where} \quad \begin{cases} Y > Z \\ W - X \overset{*}{<} Y - Z \overset{*}{>} X - W \end{cases}$$

$$(S15) \quad \begin{cases} {}_S\Delta_1 = {}_S\Delta_2 = {}_S\Delta_3 = {}_S\Delta_4 = {}_S\Delta_5 = {}_S\Delta_7 = 0 \\ 0 < -{}_S\Delta_6 \overset{*}{>} {}_S\Delta_8 > 0 \end{cases}$$

$$(S16) \quad \begin{cases} {}_S\Delta_7 \overset{*}{=} {}_S\Delta_1 = {}_S\Delta_2 = {}_S\Delta_3 = {}_S\Delta_4 = {}_S\Delta_5 = 0 \\ {}_S\Delta_8 \overset{*}{=} -{}_S\Delta_6 > 0 \end{cases}$$

| | ${}_S\Delta_{1-6}$ | ${}_S\Delta_{2-6}$ | ${}_S\Delta_{3-5}$ | ${}_S\Delta_{4-5}$ | ${}_S\Delta_{5+7}$ | ${}_S\Delta_{6+8}$ | ${}_S\Delta_{3+4-5+7}$ | ${}_S\Delta_{1+2-6+8}$ |
|---|---|---|---|---|---|---|---|---|
| ($S0$) | 0 | 0 | 0 | 0 | $0^*$ | $0^*$ | $0^*$ | $0^*$ |
| ($S12$), ($S14$) | + | + | 0 | 0 | 0 | $-^*$ | 0 | $+^*$ |
| ($S13$), ($S15$) | + | + | 0 | 0 | 0 | $-^*$ | 0 | + |
| ($S16$) | + | + | 0 | 0 | $0^*$ | $0^*$ | $0^*$ | $+^*$ |
| 5 → 4 | + | + | 0 | 0 | $0^*$ | $-^*$ | $0^*$ | $+^*$ |

**4 → 5.**
The proof is analogous to earlier proofs. We must keep in mind the possibility that events II) and IV) might have probability zero.

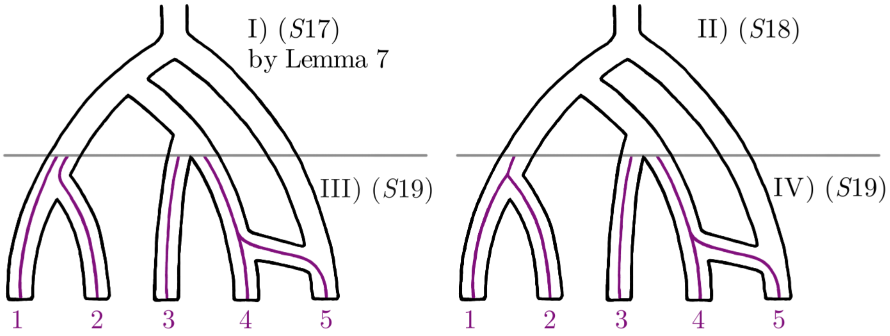


$$(S17)\quad \begin{cases} {}_S\Delta_1 = {}_S\Delta_2 = Z \\ {}_S\Delta_3 = {}_S\Delta_4 = {}_S\Delta_5 = {}_S\Delta_7 = 0 \qquad \text{where} \quad W > 0 < Z \\ -{}_S\Delta_6 = {}_S\Delta_8 = W \end{cases}$$

$$(S18)\quad \begin{cases} {}_S\Delta_1 = {}_S\Delta_2 = {}_S\Delta_3 = {}_S\Delta_4 = {}_S\Delta_5 = {}_S\Delta_7 = 0 \\ -{}_S\Delta_6 = {}_S\Delta_8 > 0 \end{cases}$$

$$(S19)\quad \begin{cases} {}_S\Delta_7 \overset{*}{=} {}_S\Delta_1 = {}_S\Delta_2 = {}_S\Delta_3 = {}_S\Delta_4 = {}_S\Delta_5 = 0 \\ {}_S\Delta_8 \overset{*}{=} -{}_S\Delta_6 > 0 \end{cases}$$

| | ${}_S\Delta_{1-6}$ | ${}_S\Delta_{2-6}$ | ${}_S\Delta_{3-5}$ | ${}_S\Delta_{4-5}$ | ${}_S\Delta_{5+7}$ | ${}_S\Delta_{6+8}$ | ${}_S\Delta_{3+4-5+7}$ | ${}_S\Delta_{1+2-6+8}$ |
|---|---|---|---|---|---|---|---|---|
| $(S0)$ | 0 | 0 | 0 | 0 | $0^*$ | $0^*$ | $0^*$ | $0^*$ |
| $(S17), (S18)$ | + | + | 0 | 0 | 0 | 0 | 0 | + |
| $(S19)$ | + | + | 0 | 0 | $0^*$ | $0^*$ | $0^*$ | $+^*$ |
| $4 \to 5$ | + | + | 0 | 0 | $0^*$ | $0^*$ | $0^*$ | $+^*$ |

#### Asymmetric tree

As shown in the main text, all binomial $\Delta$-statistics are expected to be zero when there are no admixture events, but for ${}_A\Delta_4$ we need to assume (2). We mark this relation down as equation ($A0$).

$$(A0)\quad \begin{cases} {}_A\Delta_4 \overset{*}{=} {}_A\Delta_1 = {}_A\Delta_2 = {}_A\Delta_3 = 0 \end{cases}$$

**3 → 2.**
The proof is analogous to earlier proofs. Note that the cases $2 \to 3$ and $123 \to 1$ are essentially the same as this.

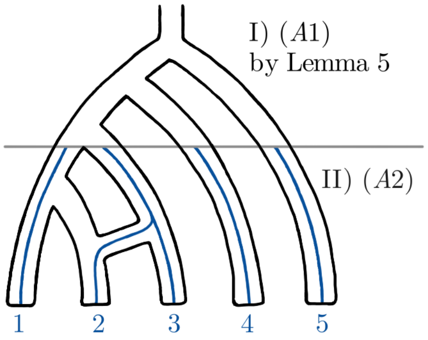


$$(A1)\quad \begin{cases} -{}_A\Delta_1 = {}_A\Delta_4 = X \\ {}_A\Delta_2 = {}_A\Delta_3 = Z \end{cases} \qquad \text{where} \quad X > 0 < Z$$

$$(A2)\quad \begin{cases} {}_A\Delta_4 \overset{*}{=} -{}_A\Delta_1 > 0 \\ {}_A\Delta_2 = {}_A\Delta_3 = 0 \end{cases}$$

| | ${}_A\Delta_{1-2}$ | ${}_A\Delta_{1-3}$ | ${}_A\Delta_{2-3}$ | ${}_A\Delta_{1+2-3+4}$ |
|---|---|---|---|---|
| ($A0$) | 0 | 0 | 0 | $0^*$ |
| ($A1$) | – | – | 0 | 0 |
| ($A2$) | – | – | 0 | $0^*$ |
| $3 \to 2$ | – | – | 0 | $0^*$ |

**2 → 4.**
The proof is analogous to earlier proofs.

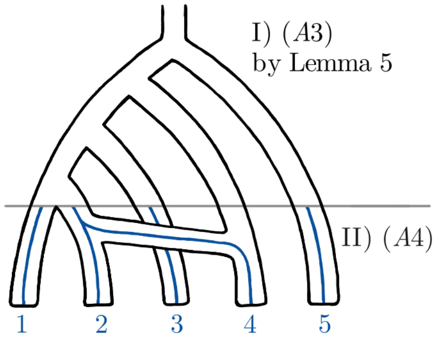


$$(A3)\quad \begin{cases} {}_A\Delta_1 = {}_A\Delta_3 = Z \\ -{}_A\Delta_2 = {}_A\Delta_4 = X \end{cases} \qquad \text{where} \quad X > 0 < Z$$

$$(A4)\quad \begin{cases} {}_A\Delta_1 = {}_A\Delta_3 = 0 \\ {}_A\Delta_4 \overset{*}{=} -{}_A\Delta_2 > 0 \end{cases}$$

| | ${}_A\Delta_{1-2}$ | ${}_A\Delta_{1-3}$ | ${}_A\Delta_{2-3}$ | ${}_A\Delta_{1+2-3+4}$ |
|---|---|---|---|---|
| $(A0)$ | 0 | 0 | 0 | $0^*$ |
| $(A3)$ | + | 0 | − | 0 |
| $(A4)$ | + | 0 | − | $0^*$ |
| $2 \to 4$ | + | 0 | − | $0^*$ |

**4 → 2.**
The proof is analogous to earlier proofs.

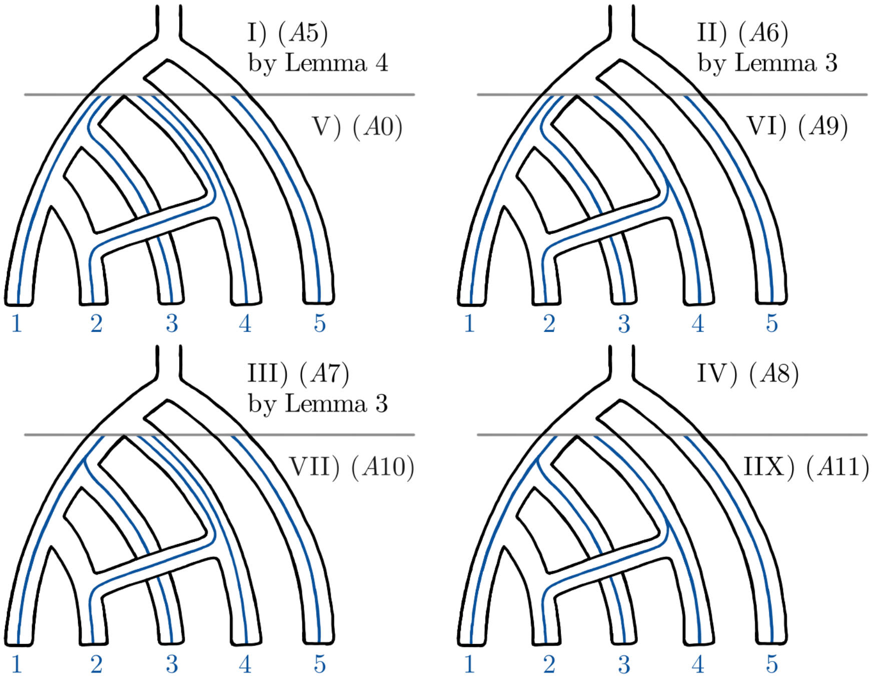


$$(A5)\qquad \begin{cases} {}_A\Delta_1 = {}_A\Delta_2 = {}_A\Delta_3 = {}_A\Delta_4 = 0 \end{cases}$$

$$(A6)\qquad \begin{cases} {}_A\Delta_1 = {}_A\Delta_3 = Z \\ -{}_A\Delta_2 = {}_A\Delta_4 = Y \end{cases} \qquad \text{where} \quad Y > 0 < Z$$

$$(A7)\qquad \begin{cases} {}_A\Delta_1 = -{}_A\Delta_4 = Y \\ {}_A\Delta_2 = {}_A\Delta_3 = -Z \end{cases} \qquad \text{where} \quad Y > 0 < Z$$

$$(A8)\qquad \begin{cases} {}_A\Delta_1 = -{}_A\Delta_2 > 0 \\ {}_A\Delta_3 = {}_A\Delta_4 = 0 \end{cases}$$

$$(A9)\qquad \begin{cases} {}_A\Delta_1 = {}_A\Delta_3 = 0 \\ {}_A\Delta_4 \overset{*}{=} -{}_A\Delta_2 > 0 \end{cases}$$

$$(A10)\qquad \begin{cases} -{}_A\Delta_4 \overset{*}{=} {}_A\Delta_1 > 0 \\ {}_A\Delta_2 = {}_A\Delta_3 = 0 \end{cases}$$

$$(A11)\qquad \begin{cases} {}_A\Delta_1 > 0 \\ {}_A\Delta_2 < 0 \\ {}_A\Delta_3 = 0 \\ {}_A\Delta_4 \overset{*}{=} -{}_A\Delta_1 - {}_A\Delta_2 \end{cases}$$

| | ${}_A\Delta_{1-2}$ | ${}_A\Delta_{1-3}$ | ${}_A\Delta_{2-3}$ | ${}_A\Delta_{1+2-3+4}$ |
|---|---|---|---|---|
| $(A0)$ | 0 | 0 | 0 | $0^*$ |
| $(A5)$ | 0 | 0 | 0 | 0 |
| $(A6)$ | + | 0 | − | 0 |
| $(A7)$ | + | + | 0 | 0 |
| $(A8)$ | + | + | − | 0 |
| $(A9)$ | + | 0 | − | $0^*$ |
| $(A10)$ | + | + | 0 | $0^*$ |
| $(A11)$ | + | + | − | $0^*$ |
| $4 \to 2$ | + | + | − | $0^*$ |

**2 → 5.**
The proof is analogous to earlier proofs.

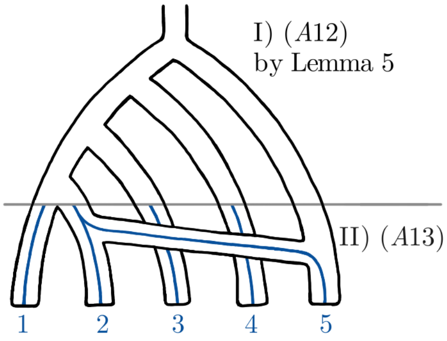


$$(A12) \quad \begin{cases} {}_A\Delta_1 = {}_A\Delta_2 = Z \\ -{}_A\Delta_3 = {}_A\Delta_4 = X \end{cases} \qquad \text{where} \quad X > 0 < Z$$

$$(A13) \quad \begin{cases} {}_A\Delta_1 = {}_A\Delta_2 = 0 \\ {}_A\Delta_4 \overset{*}{=} -{}_A\Delta_3 > 0 \end{cases}$$

| | ${}_A\Delta_{1-2}$ | ${}_A\Delta_{1-3}$ | ${}_A\Delta_{2-3}$ | ${}_A\Delta_{1+2-3+4}$ |
|---|---|---|---|---|
| $(A0)$ | 0 | 0 | 0 | $0^*$ |
| $(A12)$ | 0 | + | + | + |
| $(A13)$ | 0 | + | + | $+^*$ |
| $2 \to 5$ | 0 | + | + | $+^*$ |

**5 → 2.**
The proof is analogous to earlier proofs. We must keep in mind the possibility that events II), IV), VI) and IIX) might have probability zero.

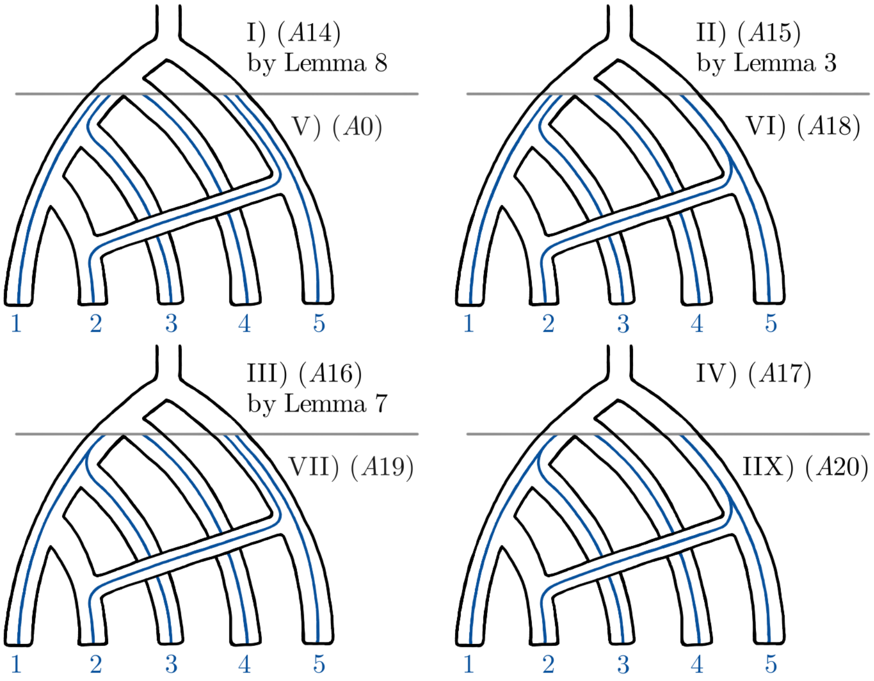


$$(A14) \quad \begin{cases} {}_A\Delta_1 = {}_A\Delta_2 = Y - Z \\ {}_A\Delta_3 = Z - X \\ {}_A\Delta_4 = W - V \end{cases} \quad \text{where} \quad \begin{cases} X > Z < Y \\ V + 3Z \overset{*}{<} W + X + 2Y \end{cases}$$

$$(A15) \quad \begin{cases} {}_A\Delta_1 = {}_A\Delta_2 = Z \\ {}_A\Delta_3 = -X \\ {}_A\Delta_4 = Y \end{cases} \quad \text{where} \quad \begin{cases} X > 0 < Y \\ Z > 0 \end{cases}$$

$$(A16) \quad \begin{cases} {}_A\Delta_1 = X \\ {}_A\Delta_2 = -Z \\ {}_A\Delta_3 = -Y \\ {}_A\Delta_4 = -W \end{cases} \quad \text{where} \quad \begin{cases} Y > Z > 0 < X \\ Y - Z \overset{*}{>} W - X \end{cases}$$

$$(A17) \quad \begin{cases} {}_A\Delta_1 > 0 \\ {}_A\Delta_2 = {}_A\Delta_4 = 0 \\ {}_A\Delta_3 < 0 \end{cases}$$

$$(A18) \quad \begin{cases} {}_A\Delta_1 = {}_A\Delta_2 = 0 \\ {}_A\Delta_4 \overset{*}{=} -{}_A\Delta_3 > 0 \end{cases}$$

$$(A19) \quad \begin{cases} -{}_A\Delta_4 \overset{*}{=} {}_A\Delta_1 > 0 \\ {}_A\Delta_2 = {}_A\Delta_3 = 0 \end{cases}$$

$$(A20) \quad \begin{cases} {}_A\Delta_1 > 0 \\ {}_A\Delta_2 = 0 \\ {}_A\Delta_3 < 0 \\ {}_A\Delta_4 \overset{*}{=} -{}_A\Delta_1 - {}_A\Delta_3 \end{cases}$$

| | ${}_A\Delta_{1-2}$ | ${}_A\Delta_{1-3}$ | ${}_A\Delta_{2-3}$ | ${}_A\Delta_{1+2-3+4}$ |
|---|---|---|---|---|
| $(A0)$ | 0 | 0 | 0 | $0^*$ |
| $(A14), (A18)$ | 0 | + | + | $+^*$ |
| $(A15)$ | 0 | + | + | + |
| $(A16), (A20)$ | + | + | + | $+^*$ |
| $(A17)$ | + | + | + | + |
| $(A19)$ | + | + | 0 | $0^*$ |
| $5 \to 2$ | + | + | + | $+^*$ |

**1234 → 2.**
The proof is analogous to earlier proofs.

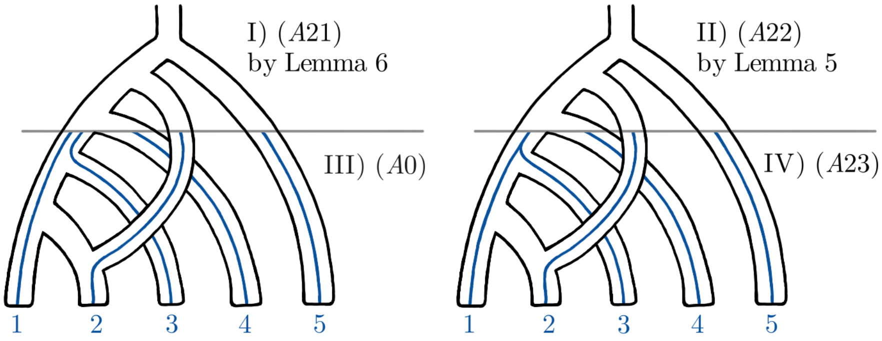


$$(A21)\quad \begin{cases} {}_A\Delta_1 = {}_A\Delta_2 = Y - Z \\ {}_A\Delta_3 = X - W \\ {}_A\Delta_4 = V - U \end{cases} \quad \text{where} \quad \begin{cases} Y > Z \\ W > X \\ U + X + 2Z \overset{*}{=} V + W + 2Y \end{cases}$$

$$(A22)\quad \begin{cases} {}_A\Delta_1 = X \\ {}_A\Delta_2 = -Z \\ {}_A\Delta_3 = -Y \\ {}_A\Delta_4 = -W \end{cases} \quad \text{where} \quad \begin{cases} Y > Z > 0 < X \\ W + Z \overset{*}{=} X + Y \end{cases}$$

$$(A23)\quad \begin{cases} -{}_A\Delta_4 \overset{*}{=} {}_A\Delta_1 > 0 \\ {}_A\Delta_2 = {}_A\Delta_3 = 0 \end{cases}$$

| | ${}_A\Delta_{1-2}$ | ${}_A\Delta_{1-3}$ | ${}_A\Delta_{2-3}$ | ${}_A\Delta_{1+2-3+4}$ |
|---|---|---|---|---|
| (A0) | 0 | 0 | 0 | $0^*$ |
| (A21) | 0 | + | + | $0^*$ |
| (A22) | + | + | + | $0^*$ |
| (A23) | + | + | 0 | $0^*$ |
| 1234 → 2 | + | + | + | $0^*$ |

**12345 → 2.**
The proof is analogous to earlier proofs.

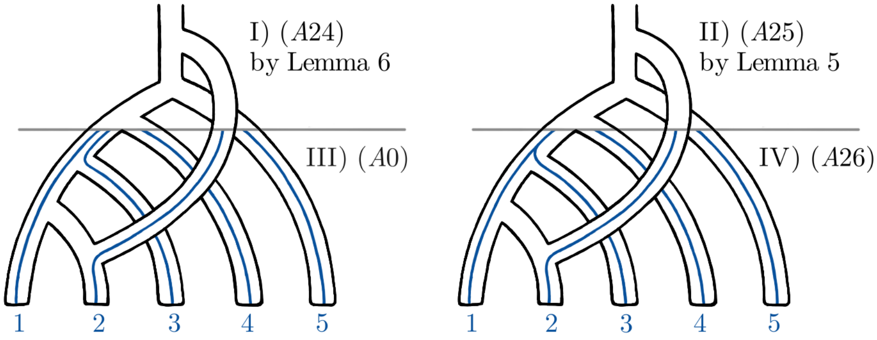


$$(A24) \quad \begin{cases} {}_A\Delta_1 = {}_A\Delta_2 = Y - X \\ {}_A\Delta_3 = Z - W \\ {}_A\Delta_4 = V - T \end{cases} \quad \text{where} \quad \begin{cases} W > X \\ Y > Z \\ T + Z + 2X \overset{*}{>} V + W + 2Y \end{cases}$$

$$(A25) \quad \begin{cases} {}_A\Delta_1 = X \\ {}_A\Delta_2 = -Z \\ {}_A\Delta_3 = -Y \\ {}_A\Delta_4 = -V \end{cases} \quad \text{where} \quad \begin{cases} Y > Z > 0 < X \\ V + Z \overset{*}{>} X + Y \end{cases}$$

$$(A26) \quad \begin{cases} -{}_A\Delta_4 \overset{*}{=} {}_A\Delta_1 > 0 \\ {}_A\Delta_2 = {}_A\Delta_3 = 0 \end{cases}$$

| | ${}_A\Delta_{1-2}$ | ${}_A\Delta_{1-3}$ | ${}_A\Delta_{2-3}$ | ${}_A\Delta_{1+2-3+4}$ |
|---|---|---|---|---|
| $(A0)$ | 0 | 0 | 0 | $0^*$ |
| $(A24)$ | 0 | + | + | $-^*$ |
| $(A25)$ | + | + | + | $-^*$ |
| $(A26)$ | + | + | 0 | $0^*$ |
| $12345 \to 2$ | + | + | + | $-^*$ |

##### Quasisymmetric tree

As shown in the main text, all binomial Δ-statistics are expected to be zero when there are no admixture events, but for ${}_Q\Delta_7$ and ${}_Q\Delta_8$ we need to assume (2). We mark this relation down as equation ($Q0$).

$$(Q0) \quad \begin{cases} {}_Q\Delta_7 \overset{*}{=} {}_Q\Delta_8 \overset{*}{=} {}_Q\Delta_1 = {}_Q\Delta_2 = {}_Q\Delta_3 = {}_Q\Delta_4 = {}_Q\Delta_5 = {}_Q\Delta_6 = 0 \end{cases}$$

**2 → 4.**
The proof is analogous to earlier proofs.

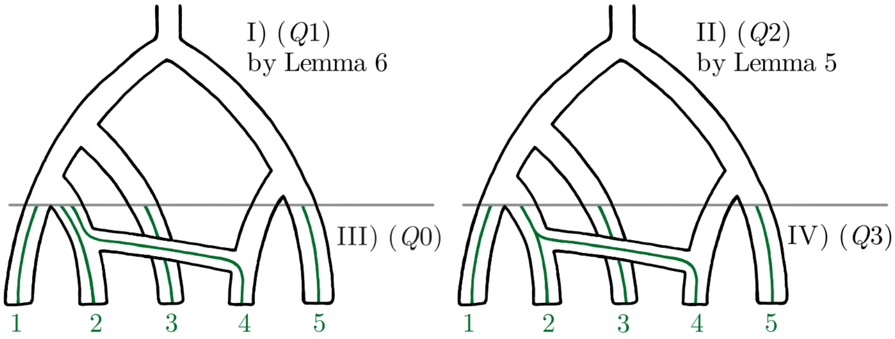


$$(Q1) \quad \begin{cases} {}_Q\Delta_1 = {}_Q\Delta_2 = Y - X \\ {}_Q\Delta_3 = {}_Q\Delta_4 = {}_Q\Delta_5 = {}_Q\Delta_7 = 0 \\ {}_Q\Delta_6 = Z - W \\ {}_Q\Delta_8 = V - T \end{cases} \quad \text{where} \quad \begin{cases} W > X > 0 \\ Y > Z > 0 \\ T + W \overset{*}{>} V + Z \\ T + Z + 2X \overset{*}{>} V + W + 2Y \end{cases}$$

$$(Q2) \quad \begin{cases} -{}_Q\Delta_1 = {}_Q\Delta_4 = {}_Q\Delta_5 = Z \\ {}_Q\Delta_2 = -{}_Q\Delta_3 = {}_Q\Delta_7 = X \\ {}_Q\Delta_6 = -Y \\ {}_Q\Delta_8 = -V \end{cases} \quad \text{where} \quad \begin{cases} V > 0 < X \\ Y > Z > 0 \\ V + Z \overset{*}{>} X + Y \end{cases}$$

$$(Q3) \quad \begin{cases} {}_Q\Delta_1 = {}_Q\Delta_4 = {}_Q\Delta_5 = {}_Q\Delta_6 = 0 \\ {}_Q\Delta_7 \overset{*}{=} -{}_Q\Delta_8 \overset{*}{=} {}_Q\Delta_2 = -{}_Q\Delta_3 > 0 \end{cases}$$

| | ${}_Q\Delta_{1-6}$ | ${}_Q\Delta_{2-6}$ | ${}_Q\Delta_{3-5}$ | ${}_Q\Delta_{4-5}$ | ${}_Q\Delta_{5+7}$ | ${}_Q\Delta_{6+8}$ | ${}_Q\Delta_{3+4-5+7}$ | ${}_Q\Delta_{1+2-6+8}$ |
|---|---|---|---|---|---|---|---|---|
| ($Q0$) | 0 | 0 | 0 | 0 | $0^*$ | $0^*$ | $0^*$ | $0^*$ |
| ($Q1$) | + | + | 0 | 0 | 0 | $-^*$ | 0 | $-^*$ |
| ($Q2$) | + | + | − | 0 | + | − | 0 | $-^*$ |
| ($Q3$) | 0 | + | − | 0 | $+^*$ | $-^*$ | $0^*$ | $0^*$ |
| 2 → 4 | + | + | − | 0 | $+^*$ | $-^*$ | $0^*$ | $-^*$ |

**4 → 2.**
The proof is analogous to earlier proofs. We must keep in mind the possibility that events III), IV), VII) and IIX) might have probability zero.

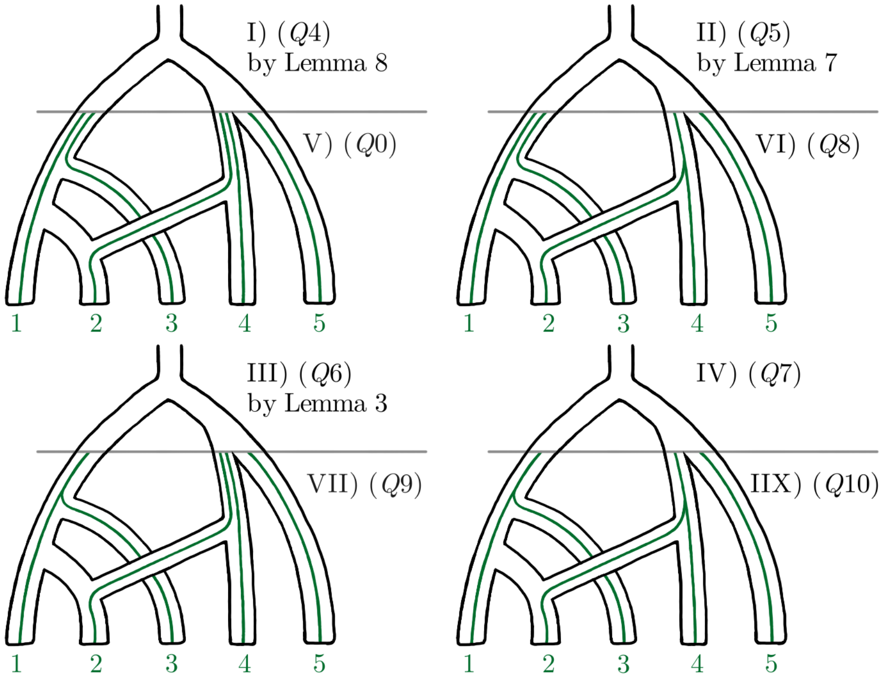


$$(Q4)\quad \begin{cases} {}_Q\Delta_1 = {}_Q\Delta_2 = {}_Q\Delta_6 = {}_Q\Delta_8 = 0 \\ {}_Q\Delta_3 = {}_Q\Delta_4 = Z - Y \\ {}_Q\Delta_5 = X - Z \\ {}_Q\Delta_7 = V - W \end{cases} \quad \text{where} \quad \begin{cases} X > Z < Y \\ V + X \overset{*}{>} W + Z \\ V + 3Z \overset{*}{<} W + X + 2Y \end{cases}$$

$$(Q5)\quad \begin{cases} -{}_Q\Delta_1 = {}_Q\Delta_4 = -{}_Q\Delta_6 = Z \\ {}_Q\Delta_2 = -{}_Q\Delta_3 = -{}_Q\Delta_8 = X \\ {}_Q\Delta_5 = Y \\ {}_Q\Delta_7 = W \end{cases} \quad \text{where} \quad \begin{cases} W > 0 < X \\ Y > Z > 0 \\ W - X \overset{*}{<} Y - Z \end{cases}$$

$$(Q6)\quad \begin{cases} {}_Q\Delta_1 = {}_Q\Delta_2 = {}_Q\Delta_6 = {}_Q\Delta_8 = 0 \\ {}_Q\Delta_3 = {}_Q\Delta_4 = -Z \\ {}_Q\Delta_5 = X \\ {}_Q\Delta_7 = -Y \end{cases} \quad \text{where} \quad 0 < X \overset{*}{>} Y > 0 < Z$$

$$(Q7)\quad \begin{cases} {}_Q\Delta_1 = {}_Q\Delta_4 = {}_Q\Delta_6 = {}_Q\Delta_7 = 0 \\ {}_Q\Delta_2 = -{}_Q\Delta_3 = -{}_Q\Delta_8 > 0 \\ {}_Q\Delta_5 > 0 \end{cases}$$

$$(Q8)\quad \begin{cases} {}_Q\Delta_1 = {}_Q\Delta_4 = {}_Q\Delta_5 = {}_Q\Delta_6 = 0 \\ {}_Q\Delta_7 \overset{*}{=} -{}_Q\Delta_8 \overset{*}{=} {}_Q\Delta_2 = -{}_Q\Delta_3 > 0 \end{cases}$$

$$(Q9)\quad \begin{cases} {}_Q\Delta_8 \overset{*}{=} {}_Q\Delta_1 = {}_Q\Delta_2 = {}_Q\Delta_3 = {}_Q\Delta_4 = {}_Q\Delta_6 = 0 \\ -{}_Q\Delta_7 \overset{*}{=} {}_Q\Delta_5 > 0 \end{cases}$$

$$(Q10)\quad \begin{cases} {}_Q\Delta_1 = {}_Q\Delta_4 = {}_Q\Delta_6 = 0 \\ -{}_Q\Delta_8 \overset{*}{=} {}_Q\Delta_2 = -{}_Q\Delta_3 > 0 \\ {}_Q\Delta_5 > 0 \\ {}_Q\Delta_7 \overset{*}{=} {}_Q\Delta_2 - {}_Q\Delta_5 \end{cases}$$

| | $_Q\Delta_{1-6}$ | $_Q\Delta_{2-6}$ | $_Q\Delta_{3-5}$ | $_Q\Delta_{4-5}$ | $_Q\Delta_{5+7}$ | $_Q\Delta_{6+8}$ | $_Q\Delta_{3+4-5+7}$ | $_Q\Delta_{1+2-6+8}$ |
|---|---|---|---|---|---|---|---|---|
| $(Q0)$ | 0 | 0 | 0 | 0 | $0^*$ | $0^*$ | $0^*$ | $0^*$ |
| $(Q4)$ | 0 | 0 | $-$ | $-$ | $+^*$ | 0 | $-^*$ | 0 |
| $(Q5)$ | 0 | $+$ | $-$ | $-$ | $+$ | $-$ | $-^*$ | 0 |
| $(Q6)$ | 0 | 0 | $-$ | $-$ | $+^*$ | 0 | $-$ | 0 |
| $(Q7)$ | 0 | $+$ | $-$ | $-$ | $+$ | $-$ | $-$ | 0 |
| $(Q8)$ | 0 | $+$ | $-$ | 0 | $+^*$ | $-^*$ | $0^*$ | $0^*$ |
| $(Q9)$ | 0 | 0 | $-$ | $-$ | $0^*$ | $0^*$ | $-^*$ | $0^*$ |
| $(Q10)$ | 0 | $+$ | $-$ | $-$ | $+^*$ | $-^*$ | $-^*$ | $0^*$ |
| $4 \to 2$ | 0 | $+$ | $-$ | $-$ | $+^*$ | $-^*$ | $-^*$ | $0^*$ |

**2 → 45.**
The proof is analogous to earlier proofs.

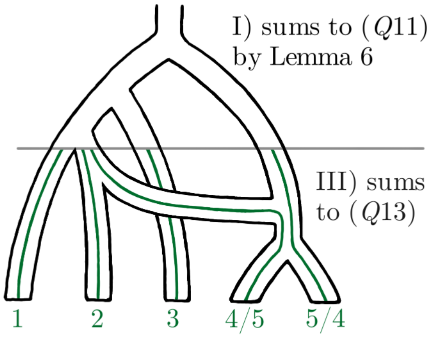


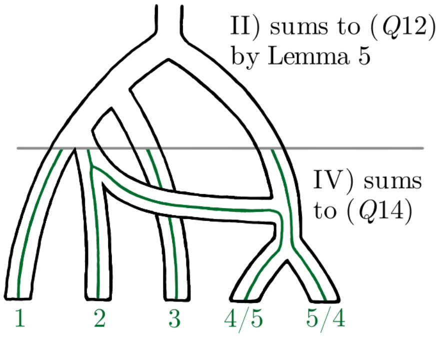


$$(Q11) \quad \begin{cases} {}_Q\Delta_1 = {}_Q\Delta_2 = {}_Q\Delta_3 = {}_Q\Delta_4 = {}_Q\Delta_5 = {}_Q\Delta_6 = {}_Q\Delta_7 = {}_Q\Delta_8 = 0 \end{cases}$$

$$(Q12) \quad \begin{cases} {}_Q\Delta_1 = {}_Q\Delta_2 = {}_Q\Delta_6 = {}_Q\Delta_8 = 0 \\ {}_Q\Delta_3 = {}_Q\Delta_4 = Z - X \\ {}_Q\Delta_5 = 2Z \\ {}_Q\Delta_7 = 2X \end{cases} \qquad \text{where} \quad X > 0 < Z$$

$$(Q13) \quad \begin{cases} {}_Q\Delta_7 \overset{*}{=} {}_Q\Delta_1 = {}_Q\Delta_2 = {}_Q\Delta_3 = {}_Q\Delta_4 = {}_Q\Delta_5 = {}_Q\Delta_6 = {}_Q\Delta_8 = 0 \end{cases}$$

$$(Q14) \quad \begin{cases} {}_Q\Delta_1 = {}_Q\Delta_2 = {}_Q\Delta_5 = {}_Q\Delta_6 = {}_Q\Delta_8 = 0 \\ {}_Q\Delta_7 \overset{*}{=} -2{}_Q\Delta_3 = -2{}_Q\Delta_4 > 0 \end{cases}$$

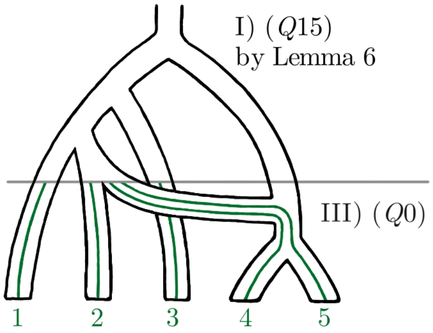


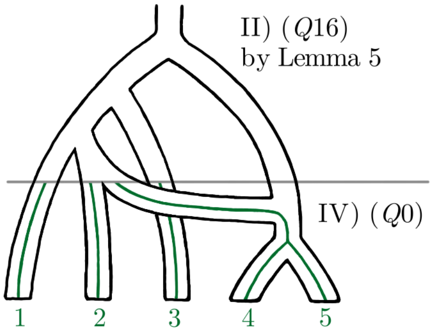


$$(Q15) \quad \begin{cases} {}_Q\Delta_1 = {}_Q\Delta_2 = {}_Q\Delta_6 = {}_Q\Delta_8 = 0 \\ {}_Q\Delta_3 = {}_Q\Delta_4 = Z - Y \\ {}_Q\Delta_5 = W - X \\ {}_Q\Delta_7 = U - V \end{cases} \qquad \text{where} \quad \begin{cases} U \overset{*}{>} V \\ W > X \\ Y > Z \\ U + X + 2Z \overset{*}{=} V + W + 2Y \end{cases}$$

$$(Q16) \quad \begin{cases} {}_Q\Delta_1 = {}_Q\Delta_2 = {}_Q\Delta_3 = {}_Q\Delta_4 = {}_Q\Delta_6 = {}_Q\Delta_8 = 0 \\ {}_Q\Delta_5 = Y - Z \\ {}_Q\Delta_7 = W - X \end{cases} \qquad \text{where} \quad \begin{cases} W \overset{*}{>} X \\ Y > Z \\ W + Z \overset{*}{=} X + Y \end{cases}$$

| | ${}_Q\Delta_{1-6}$ | ${}_Q\Delta_{2-6}$ | ${}_Q\Delta_{3-5}$ | ${}_Q\Delta_{4-5}$ | ${}_Q\Delta_{5+7}$ | ${}_Q\Delta_{6+8}$ | ${}_Q\Delta_{3+4-5+7}$ | ${}_Q\Delta_{1+2-6+8}$ |
|---|---|---|---|---|---|---|---|---|
| $(Q0)$ | 0 | 0 | 0 | 0 | $0^*$ | $0^*$ | $0^*$ | $0^*$ |
| $(Q11)$ | 0 | 0 | 0 | 0 | 0 | 0 | 0 | 0 |
| $(Q12)$ | 0 | 0 | $-$ | $-$ | $+$ | 0 | 0 | 0 |
| $(Q13)$ | 0 | 0 | 0 | 0 | $0^*$ | 0 | $0^*$ | 0 |
| $(Q14), (Q15), (Q16)$ | 0 | 0 | $-$ | $-$ | $+^*$ | 0 | $0^*$ | 0 |
| $2 \to 45$ | 0 | 0 | $-$ | $-$ | $+^*$ | $0^*$ | $0^*$ | $0^*$ |

**45 → 2.**

The proof is analogous to earlier proofs. We must keep in mind the possibility that events III), IV), VII) and IIX) might have probability zero.

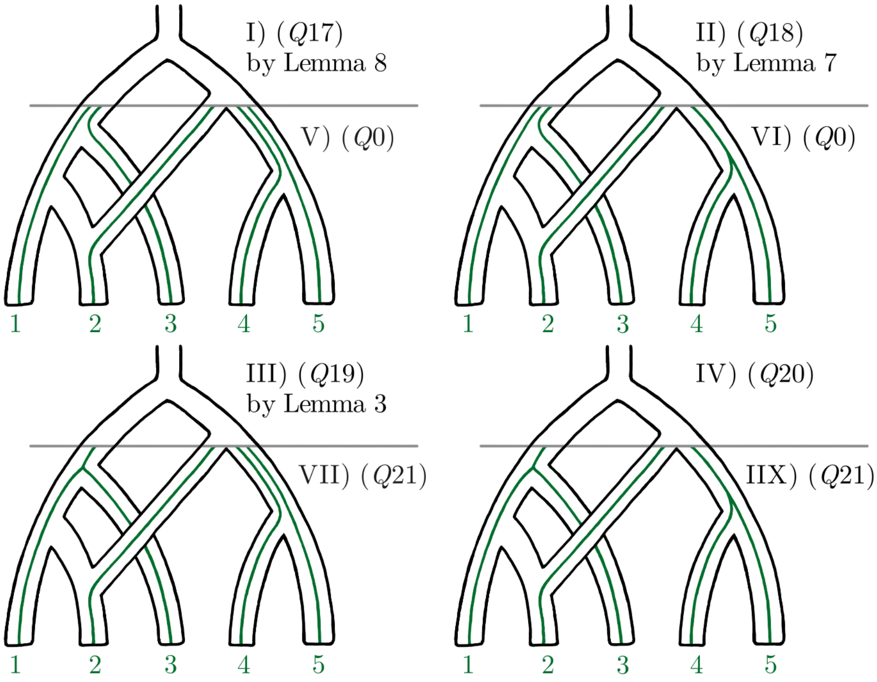


$$(Q17)\quad \begin{cases} {}_Q\Delta_1 = {}_Q\Delta_2 = {}_Q\Delta_6 = {}_Q\Delta_8 = 0 \\ {}_Q\Delta_3 = {}_Q\Delta_4 = Z - Y \\ {}_Q\Delta_5 = X - Z \\ {}_Q\Delta_7 = V - W \end{cases} \quad \text{where} \quad \begin{cases} X > Z < Y \\ V + X \overset{*}{>} W + Z \\ V + 3Z \overset{*}{<} W + X + 2Y \end{cases}$$

$$(Q18)\quad \begin{cases} {}_Q\Delta_1 = {}_Q\Delta_2 = {}_Q\Delta_3 = {}_Q\Delta_4 = {}_Q\Delta_6 = {}_Q\Delta_8 = 0 \\ {}_Q\Delta_5 = Y - Z \\ {}_Q\Delta_7 = W - X \end{cases} \quad \text{where} \quad \begin{cases} Y > Z \\ W - X \overset{*}{<} Y - Z \overset{*}{>} X - W \end{cases}$$

$$(Q19)\quad \begin{cases} {}_Q\Delta_1 = {}_Q\Delta_2 = {}_Q\Delta_6 = {}_Q\Delta_8 = 0 \\ {}_Q\Delta_3 = {}_Q\Delta_4 = -Z \\ {}_Q\Delta_5 = X \\ {}_Q\Delta_7 = -Y \end{cases} \quad \text{where} \quad Y \overset{*}{>} X > 0 < Z$$

$$(Q20)\quad \begin{cases} {}_Q\Delta_1 = {}_Q\Delta_2 = {}_Q\Delta_3 = {}_Q\Delta_4 = {}_Q\Delta_6 = {}_Q\Delta_8 = 0 \\ 0 < {}_Q\Delta_5 \overset{*}{>} -{}_Q\Delta_7 > 0 \end{cases}$$

$$(Q21)\quad \begin{cases} {}_Q\Delta_8 \overset{*}{=} {}_Q\Delta_1 = {}_Q\Delta_2 = {}_Q\Delta_3 = {}_Q\Delta_4 = {}_Q\Delta_6 = 0 \\ -{}_Q\Delta_7 \overset{*}{=} {}_Q\Delta_5 > 0 \end{cases}$$

| | ${}_Q\Delta_{1-6}$ | ${}_Q\Delta_{2-6}$ | ${}_Q\Delta_{3-5}$ | ${}_Q\Delta_{4-5}$ | ${}_Q\Delta_{5+7}$ | ${}_Q\Delta_{6+8}$ | ${}_Q\Delta_{3+4-5+7}$ | ${}_Q\Delta_{1+2-6+8}$ |
|---|---|---|---|---|---|---|---|---|
| $(Q0)$ | 0 | 0 | 0 | 0 | $0^*$ | $0^*$ | $0^*$ | $0^*$ |
| $(Q17), (Q18), (Q19)$ | 0 | 0 | − | − | $+^*$ | 0 | $-^*$ | 0 |
| $(Q20)$ | 0 | 0 | − | − | $+^*$ | 0 | − | 0 |
| $(Q21)$ | 0 | 0 | − | − | $0^*$ | $0^*$ | $-^*$ | $0^*$ |
| 45 → 2 | 0 | 0 | − | − | $+^*$ | $0^*$ | $-^*$ | $0^*$ |

**123 → 2.**
The proof is analogous to earlier proofs. Note that the cases 1 ↔ 3 are essentially the same as this.

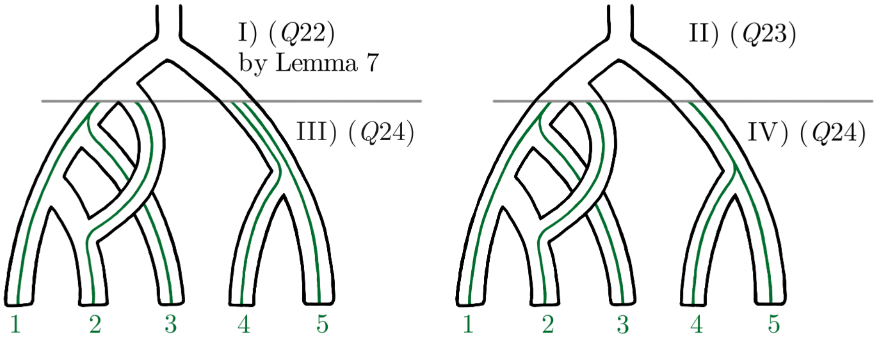


$$(Q22)\quad \begin{cases} {}_Q\Delta_1 = {}_Q\Delta_2 = {}_Q\Delta_6 = {}_Q\Delta_8 = 0 \\ {}_Q\Delta_3 = {}_Q\Delta_4 = -Z \\ {}_Q\Delta_5 = -{}_Q\Delta_7 = W \end{cases} \qquad \text{where} \quad W > 0 < Z$$

$$(Q23)\quad \begin{cases} {}_Q\Delta_1 = {}_Q\Delta_2 = {}_Q\Delta_3 = {}_Q\Delta_4 = {}_Q\Delta_6 = {}_Q\Delta_8 = 0 \\ {}_Q\Delta_5 = -{}_Q\Delta_7 > 0 \end{cases}$$

$$(Q24)\quad \begin{cases} {}_Q\Delta_8 \overset{*}{=} {}_Q\Delta_1 = {}_Q\Delta_2 = {}_Q\Delta_3 = {}_Q\Delta_4 = {}_Q\Delta_6 = 0 \\ -{}_Q\Delta_7 \overset{*}{=} {}_Q\Delta_5 > 0 \end{cases}$$

| | ${}_Q\Delta_{1-6}$ | ${}_Q\Delta_{2-6}$ | ${}_Q\Delta_{3-5}$ | ${}_Q\Delta_{4-5}$ | ${}_Q\Delta_{5+7}$ | ${}_Q\Delta_{6+8}$ | ${}_Q\Delta_{3+4-5+7}$ | ${}_Q\Delta_{1+2-6+8}$ |
|---|---|---|---|---|---|---|---|---|
| $(Q0)$ | 0 | 0 | 0 | 0 | $0^*$ | $0^*$ | $0^*$ | $0^*$ |
| $(Q22), (Q23)$ | 0 | 0 | − | − | 0 | 0 | − | 0 |
| $(Q24)$ | 0 | 0 | − | − | $0^*$ | $0^*$ | $-^*$ | $0^*$ |
| $123 \to 2$ | 0 | 0 | − | − | $0^*$ | $0^*$ | $-^*$ | $0^*$ |

**12345 → 2.**
The proof is analogous to earlier proofs.

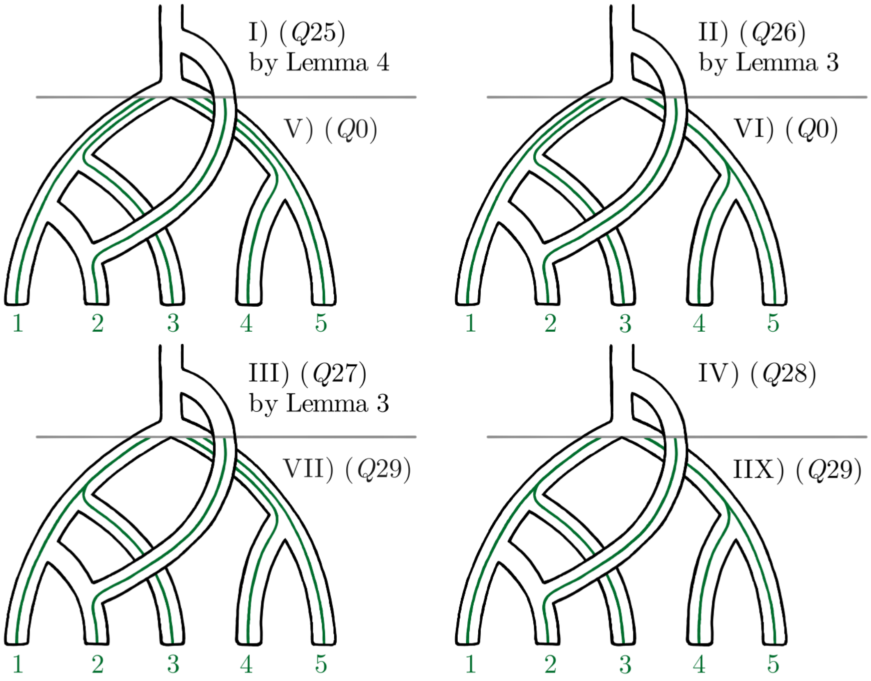


$$(Q25)\quad \begin{cases} {}_Q\Delta_1 = {}_Q\Delta_2 = {}_Q\Delta_6 = {}_Q\Delta_8 = 0 \\ {}_Q\Delta_3 = {}_Q\Delta_4 = {}_Q\Delta_5 = Z - Y \\ {}_Q\Delta_7 = X - W \end{cases} \quad \text{where} \quad W + Y \overset{*}{>} X + Z$$

$$(Q26)\quad \begin{cases} {}_Q\Delta_1 = {}_Q\Delta_2 = {}_Q\Delta_3 = {}_Q\Delta_4 = {}_Q\Delta_5 = {}_Q\Delta_6 = {}_Q\Delta_8 = 0 \\ {}_Q\Delta_7 = Y - X \end{cases} \quad \text{where} \quad X \overset{*}{>} Y$$

$$(Q27)\quad \begin{cases} {}_Q\Delta_1 = {}_Q\Delta_2 = {}_Q\Delta_3 = {}_Q\Delta_4 = {}_Q\Delta_6 = {}_Q\Delta_8 = 0 \\ {}_Q\Delta_5 = Y \\ {}_Q\Delta_7 = -X \end{cases} \quad \text{where} \quad 0 < X \overset{*}{>} Y > 0$$

$$(Q28)\quad \begin{cases} {}_Q\Delta_1 = {}_Q\Delta_2 = {}_Q\Delta_3 = {}_Q\Delta_4 = {}_Q\Delta_6 = {}_Q\Delta_8 = 0 \\ 0 < -{}_Q\Delta_7 \overset{*}{>} {}_Q\Delta_5 > 0 \end{cases}$$

$$(Q29)\quad \begin{cases} {}_Q\Delta_8 \overset{*}{=} {}_Q\Delta_1 = {}_Q\Delta_2 = {}_Q\Delta_3 = {}_Q\Delta_4 = {}_Q\Delta_6 = 0 \\ -{}_Q\Delta_7 \overset{*}{=} {}_Q\Delta_5 > 0 \end{cases}$$

| | ${}_Q\Delta_{1-6}$ | ${}_Q\Delta_{2-6}$ | ${}_Q\Delta_{3-5}$ | ${}_Q\Delta_{4-5}$ | ${}_Q\Delta_{5+7}$ | ${}_Q\Delta_{6+8}$ | ${}_Q\Delta_{3+4-5+7}$ | ${}_Q\Delta_{1+2-6+8}$ |
|---|---|---|---|---|---|---|---|---|
| ($Q0$) | 0 | 0 | 0 | 0 | $0^*$ | $0^*$ | $0^*$ | $0^*$ |
| ($Q25$), ($Q26$) | 0 | 0 | 0 | 0 | $-^*$ | 0 | $-^*$ | 0 |
| ($Q27$), ($Q28$) | 0 | 0 | $-$ | $-$ | $-^*$ | 0 | $-$ | 0 |
| ($Q29$) | 0 | 0 | $-$ | $-$ | $0^*$ | $0^*$ | $-^*$ | $0^*$ |
| 12345 → 2 | 0 | 0 | $-$ | $-$ | $-^*$ | $0^*$ | $-^*$ | $0^*$ |

**4 → 12.**

The proof is analogous to earlier proofs. We must keep in mind the possibility that events I) and III) in the first half of the proof might have probability zero.

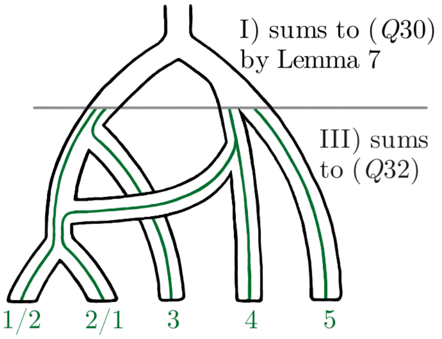


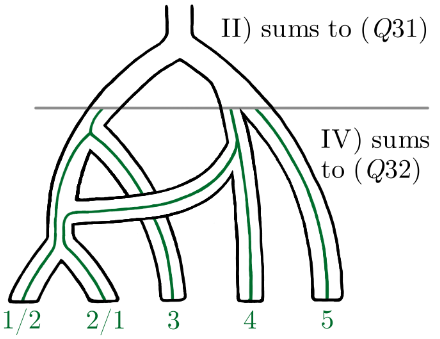


$$(Q30)\quad \begin{cases} {}_Q\Delta_1 = {}_Q\Delta_2 = X - Z \\ {}_Q\Delta_3 = {}_Q\Delta_4 = {}_Q\Delta_5 = {}_Q\Delta_7 = 0 \\ {}_Q\Delta_6 = -2Z \\ {}_Q\Delta_8 = -2X \end{cases} \quad \text{where} \quad X > 0 < Z$$

$$(Q31)\quad \begin{cases} 2{}_Q\Delta_1 = 2{}_Q\Delta_2 = -{}_Q\Delta_8 > 0 \\ {}_Q\Delta_3 = {}_Q\Delta_4 = {}_Q\Delta_5 = {}_Q\Delta_6 = {}_Q\Delta_7 = 0 \end{cases}$$

$$(Q32)\quad \begin{cases} -{}_Q\Delta_8 \overset{*}{=} 2{}_Q\Delta_1 = 2{}_Q\Delta_2 > 0 \\ {}_Q\Delta_3 = {}_Q\Delta_4 = {}_Q\Delta_5 = {}_Q\Delta_6 = {}_Q\Delta_7 = 0 \end{cases}$$

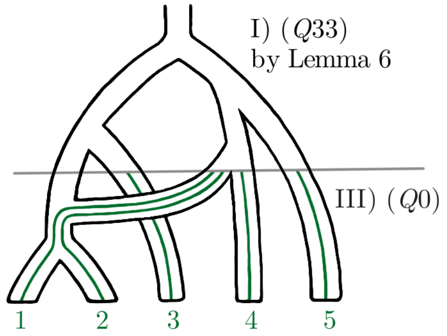


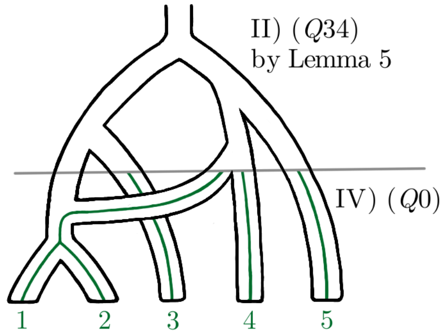


$$(Q33)\quad \begin{cases} {}_Q\Delta_1 = {}_Q\Delta_2 = Y - Z \\ {}_Q\Delta_3 = {}_Q\Delta_4 = {}_Q\Delta_5 = {}_Q\Delta_7 = 0 \\ {}_Q\Delta_6 = X - W \\ {}_Q\Delta_8 = V - U \end{cases} \quad \text{where} \quad \begin{cases} U \overset{*}{>} V \\ W > X \\ Y > Z \\ U + X + 2Z \overset{*}{=} V + W + 2Y \end{cases}$$

$$(Q34)\quad \begin{cases} {}_Q\Delta_1 = {}_Q\Delta_2 = {}_Q\Delta_3 = {}_Q\Delta_4 = {}_Q\Delta_5 = {}_Q\Delta_7 = 0 \\ {}_Q\Delta_6 = Z - Y \\ {}_Q\Delta_8 = X - W \end{cases} \quad \text{where} \quad \begin{cases} W \overset{*}{>} X \\ Y > Z \\ W + Z \overset{*}{=} X + Y \end{cases}$$

| | ${}_Q\Delta_{1-6}$ | ${}_Q\Delta_{2-6}$ | ${}_Q\Delta_{3-5}$ | ${}_Q\Delta_{4-5}$ | ${}_Q\Delta_{5+7}$ | ${}_Q\Delta_{6+8}$ | ${}_Q\Delta_{3+4-5+7}$ | ${}_Q\Delta_{1+2-6+8}$ |
|---|---|---|---|---|---|---|---|---|
| $(Q0)$ | 0 | 0 | 0 | 0 | $0^*$ | $0^*$ | $0^*$ | $0^*$ |
| $(Q30), (Q31)$ | + | + | 0 | 0 | 0 | − | 0 | 0 |
| $(Q32), (Q33), (Q34)$ | + | + | 0 | 0 | 0 | $-^*$ | 0 | $0^*$ |
| $4 \to 12$ | + | + | 0 | 0 | $0^*$ | $-^*$ | $0^*$ | $0^*$ |

**12 → 4.**
The proof is analogous to earlier proofs.

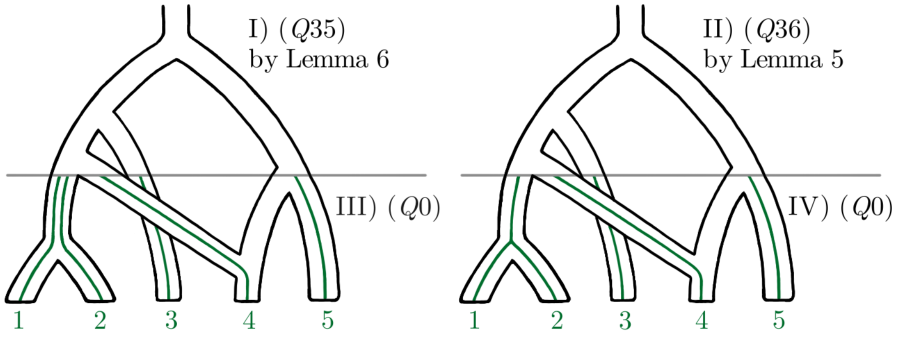


$$(Q35)\quad \begin{cases} {}_Q\Delta_1 = {}_Q\Delta_2 = Y - X \\ {}_Q\Delta_3 = {}_Q\Delta_4 = {}_Q\Delta_5 = {}_Q\Delta_7 = 0 \\ {}_Q\Delta_6 = Z - W \\ {}_Q\Delta_8 = V - T \end{cases} \quad \text{where} \quad \begin{cases} W > X \\ Y > Z \\ T + W \overset{*}{>} V + Z \\ T + Z + 2X \overset{*}{>} V + W + 2Y \end{cases}$$

$$(Q36)\quad \begin{cases} {}_Q\Delta_1 = {}_Q\Delta_2 = {}_Q\Delta_3 = {}_Q\Delta_4 = {}_Q\Delta_5 = {}_Q\Delta_7 = 0 \\ {}_Q\Delta_6 = Z - Y \\ {}_Q\Delta_8 = X - V \end{cases} \quad \text{where} \quad \begin{cases} V \overset{*}{>} X \\ Y > Z \\ V + Z \overset{*}{>} X + Y \end{cases}$$

| | ${}_Q\Delta_{1-6}$ | ${}_Q\Delta_{2-6}$ | ${}_Q\Delta_{3-5}$ | ${}_Q\Delta_{4-5}$ | ${}_Q\Delta_{5+7}$ | ${}_Q\Delta_{6+8}$ | ${}_Q\Delta_{3+4-5+7}$ | ${}_Q\Delta_{1+2-6+8}$ |
|---|---|---|---|---|---|---|---|---|
| $(Q0)$ | 0 | 0 | 0 | 0 | $0^*$ | $0^*$ | $0^*$ | $0^*$ |
| $(Q35), (Q36)$ | $+$ | $+$ | 0 | 0 | 0 | $-^*$ | 0 | $-^*$ |
| $12 \to 4$ | $+$ | $+$ | 0 | 0 | $0^*$ | $-^*$ | $0^*$ | $-^*$ |

**5 → 3.**
The proof is analogous to earlier proofs. We must keep in mind the possibility that events II) and IV) might have probability zero.

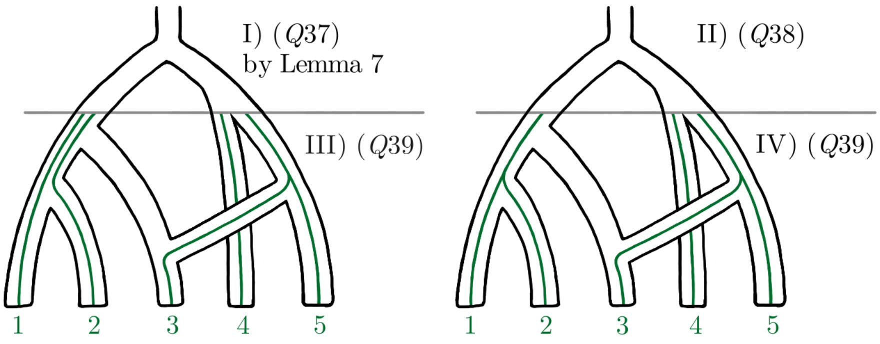


$$(Q37)\quad \begin{cases} {}_Q\Delta_1 = {}_Q\Delta_2 = Z \\ {}_Q\Delta_3 = {}_Q\Delta_4 = {}_Q\Delta_5 = {}_Q\Delta_7 = 0 \\ {}_Q\Delta_6 = -W \\ {}_Q\Delta_8 = W \end{cases} \qquad \text{where} \quad W > 0 < Z$$

$$(Q38)\quad \begin{cases} {}_Q\Delta_1 = {}_Q\Delta_2 = {}_Q\Delta_3 = {}_Q\Delta_4 = {}_Q\Delta_5 = {}_Q\Delta_7 = 0 \\ -{}_Q\Delta_6 = {}_Q\Delta_8 > 0 \end{cases}$$

$$(Q39)\quad \begin{cases} {}_Q\Delta_7 \overset{*}{=} {}_Q\Delta_1 = {}_Q\Delta_2 = {}_Q\Delta_3 = {}_Q\Delta_4 = {}_Q\Delta_5 = 0 \\ {}_Q\Delta_8 \overset{*}{=} -{}_Q\Delta_6 > 0 \end{cases}$$

| | ${}_Q\Delta_{1-6}$ | ${}_Q\Delta_{2-6}$ | ${}_Q\Delta_{3-5}$ | ${}_Q\Delta_{4-5}$ | ${}_Q\Delta_{5+7}$ | ${}_Q\Delta_{6+8}$ | ${}_Q\Delta_{3+4-5+7}$ | ${}_Q\Delta_{1+2-6+8}$ |
|---|---|---|---|---|---|---|---|---|
| $(Q0)$ | 0 | 0 | 0 | 0 | $0^*$ | $0^*$ | $0^*$ | $0^*$ |
| $(Q37), (Q38)$ | + | + | 0 | 0 | 0 | 0 | 0 | + |
| $(Q39)$ | + | + | 0 | 0 | $0^*$ | $0^*$ | $0^*$ | $+^*$ |
| $5 \to 3$ | + | + | 0 | 0 | $0^*$ | $0^*$ | $0^*$ | $+^*$ |

**3 → 5.**

The proof is analogous to earlier proofs.

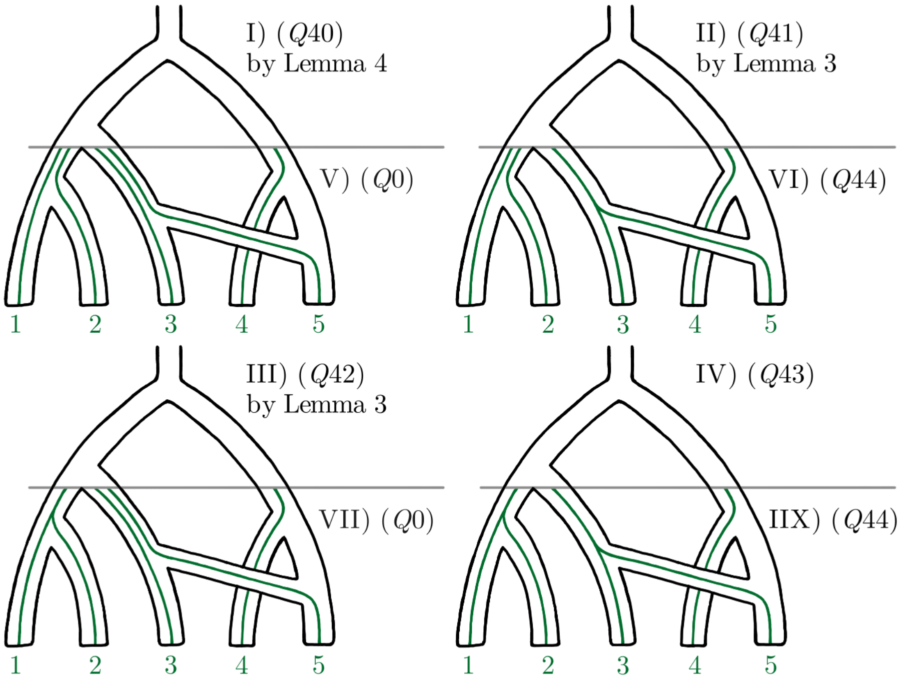


$$(Q40) \quad \begin{cases} {}_Q\Delta_1 = {}_Q\Delta_2 = {}_Q\Delta_6 = Y - Z \\ {}_Q\Delta_3 = {}_Q\Delta_4 = {}_Q\Delta_5 = {}_Q\Delta_7 = 0 \\ {}_Q\Delta_8 = W - X \end{cases} \qquad \text{where} \quad W + Y \overset{*}{>} X + Z$$

$$(Q41) \quad \begin{cases} {}_Q\Delta_1 = {}_Q\Delta_2 = Z \\ {}_Q\Delta_3 = {}_Q\Delta_4 = {}_Q\Delta_5 = {}_Q\Delta_7 = 0 \\ {}_Q\Delta_6 = -Y \\ {}_Q\Delta_8 = X \end{cases} \qquad \text{where} \quad 0 < X \overset{*}{>} Y > 0 < Z$$

$$(Q42) \quad \begin{cases} {}_Q\Delta_1 = {}_Q\Delta_2 = {}_Q\Delta_3 = {}_Q\Delta_4 = {}_Q\Delta_5 = {}_Q\Delta_6 = {}_Q\Delta_7 = 0 \\ {}_Q\Delta_8 = X - Y \end{cases} \qquad \text{where} \quad X \overset{*}{>} Y$$

$$(Q43) \quad \begin{cases} {}_Q\Delta_1 = {}_Q\Delta_2 = {}_Q\Delta_3 = {}_Q\Delta_4 = {}_Q\Delta_5 = {}_Q\Delta_7 = 0 \\ 0 < {}_Q\Delta_8 \overset{*}{>} -{}_Q\Delta_6 > 0 \end{cases}$$

$$(Q44) \quad \begin{cases} {}_Q\Delta_7 \overset{*}{=} {}_Q\Delta_1 = {}_Q\Delta_2 = {}_Q\Delta_3 = {}_Q\Delta_4 = {}_Q\Delta_5 = 0 \\ {}_Q\Delta_8 \overset{*}{=} -{}_Q\Delta_6 > 0 \end{cases}$$

| | ${}_Q\Delta_{1-6}$ | ${}_Q\Delta_{2-6}$ | ${}_Q\Delta_{3-5}$ | ${}_Q\Delta_{4-5}$ | ${}_Q\Delta_{5+7}$ | ${}_Q\Delta_{6+8}$ | ${}_Q\Delta_{3+4-5+7}$ | ${}_Q\Delta_{1+2-6+8}$ |
|---|---|---|---|---|---|---|---|---|
| $(Q0)$ | 0 | 0 | 0 | 0 | $0^*$ | $0^*$ | $0^*$ | $0^*$ |
| $(Q40), (Q42)$ | 0 | 0 | 0 | 0 | 0 | $+^*$ | 0 | $+^*$ |
| $(Q41), (Q43)$ | + | + | 0 | 0 | 0 | $+^*$ | 0 | + |
| $(Q44)$ | + | + | 0 | 0 | $0^*$ | $0^*$ | $0^*$ | $+^*$ |
| $3 \to 5$ | + | + | 0 | 0 | $0^*$ | $+^*$ | $0^*$ | $+^*$ |

**123 → 4.**
The proof is analogous to earlier proofs. Note that the cases $4 \to 123$ and $12345 \to 5$ are essentially the same as this.

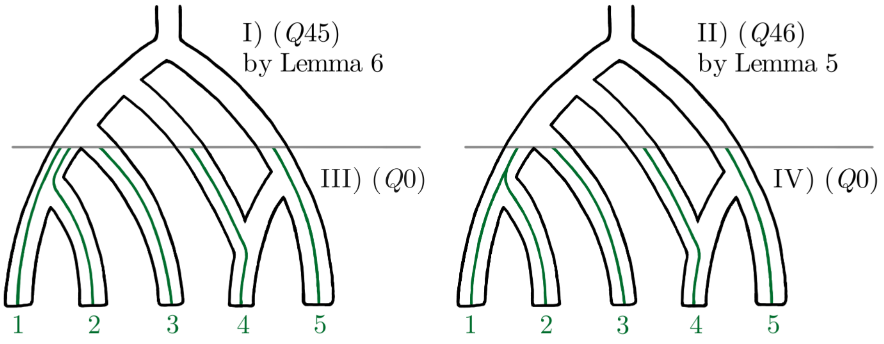


$$(Q45) \quad \begin{cases} {}_Q\Delta_1 = {}_Q\Delta_2 = {}_Q\Delta_6 = Z - X \\ {}_Q\Delta_3 = {}_Q\Delta_4 = {}_Q\Delta_5 = {}_Q\Delta_7 = 0 \\ {}_Q\Delta_8 = U - T \end{cases} \quad \text{where} \quad T + X \overset{*}{>} U + Z$$

$$(Q46) \quad \begin{cases} {}_Q\Delta_1 = {}_Q\Delta_2 = {}_Q\Delta_3 = {}_Q\Delta_4 = {}_Q\Delta_5 = {}_Q\Delta_6 = {}_Q\Delta_7 = 0 \\ {}_Q\Delta_8 = W - V \end{cases} \quad \text{where} \quad V \overset{*}{>} W$$

| | ${}_Q\Delta_{1-6}$ | ${}_Q\Delta_{2-6}$ | ${}_Q\Delta_{3-5}$ | ${}_Q\Delta_{4-5}$ | ${}_Q\Delta_{5+7}$ | ${}_Q\Delta_{6+8}$ | ${}_Q\Delta_{3+4-5+7}$ | ${}_Q\Delta_{1+2-6+8}$ |
|---|---|---|---|---|---|---|---|---|
| $(Q0)$ | 0 | 0 | 0 | 0 | $0^*$ | $0^*$ | $0^*$ | $0^*$ |
| $(Q45), (Q46)$ | 0 | 0 | 0 | 0 | 0 | $-^*$ | 0 | $-^*$ |
| $1234 \to 4$ | 0 | 0 | 0 | 0 | $0^*$ | $-^*$ | $0^*$ | $-^*$ |

#### Omitting the outgroup mutation assumption (1)

We operate under the outgroup mutation assumption (1), but earlier methods like the $D$-statistic or the $D_{\text{FOIL}}$ do not. That is, we treat all populations the same even in in trees $S$ or $A$ instead of filtering the data so that lineage 5 would always carry the ancestral allele (it's unclear how the tree $Q$ without a potential outgroup should be viewed then). Pease & Hahn, 2015 also point out that mathematically the outgroup is not needed. We stress that the filtering doesn't protect against varying generation times and mutation rates between populations, in other words against situations where the synchronization assumption (2) is not valid. Nor do Pease and Hahn claim that it would, but we believe such misconception nevertheless exists. No protection is even necessary for those $\Delta$-statistics that don't use singleton patterns; this is also true for the classic $D$-statistics. And when the singleton patterns are in use, the filtering wouldn't help — for example the predictions of the $D_{\text{FOIL}}$-statistics malfunction with ancient data whether the filtering is done or not (see Appendix 5). (Note that ancient samples can be viewed as extreme cases of violating the synchronization assumption (2)). As the filtering procedure is either unnecessary or insufficient, depending on whether one avoids the singleton patterns or uses the full set of $\Delta$-statistics, we have chosen to not recommend it and formulated the outgroup mutation assumption (1) instead.

We still recognize that in practice a preprocessing step of filtering is sometimes reality, and it's therefore necessary to explore its expected consequences. First, the derived alleles private to lineage 5 are not used for anything is trees $S$ and $A$, so removing them changes nothing. Now suppose that 5 is a "true outgroup" in the sense that it's not involved in any gene flow events. Because the population 5 has the deepest divergence time, non-singleton patterns arising due to ILS where 5 carries the derived allele are relatively uncommon. That is not to say they aren't informative about gene flow events between other populations (for example in main text Figure 3 Panel II) the pattern AABAB is impossible while the pattern AAABB can still occur whether B is the derived or the ancestral allele), only that the harm caused by disregarding them is limited. Real issues are encountered when 5 is not a "true outgroup". For example in tree $S$ the event $5 \to 4$ would result in large number of patterns AAABB where B is the derived allele: the filtering discards key information. Worse, it turns out that some predictions in Tables 2 and 3 are no longer valid. More precisely, the filtering affects the statistic ${}_S\Delta_{5+7}$ under $1 \leftrightarrow 5$ or $2 \leftrightarrow 5$, the statistic ${}_S\Delta_{6+8}$ under $2 \leftrightarrow 5$ or $3 \leftrightarrow 5$, and the statistic ${}_A\Delta_{1-2}$ under $1 \to 5$ or $2 \to 5$. Other cases are unaffected — when 5 is a "true outgroup" the filtering does no harm. While not giving a full formal proof, we will next illuminate these claims. We present thirteen specialized versions of Lemmas 1–8, where one lineage is assumed to carry the ancestral allele. Using these Lemmas, it's straightforward if laborious to go through the proofs on symmetric and asymmetric trees again, this time assuming that 5 can't be downstream of a mutation. When the task is complete, the reader will appreciate how the outgroup mutation assumption (1) naturally simplifies the mathematics.

**Lemma 1** (4 ancestral)**.** *Suppose* $(\{1,2,3,4\})$*, assume that the allele in lineage 4 is ancestral and denote*

$$\begin{cases} \mathbb{P}(\text{BAAA}) = \mathbb{P}(\text{ABAA}) = \mathbb{P}(\text{AABA}) = Y\,, \\ \mathbb{P}(\text{AAAB}) = \overline{Y}\,, \\ \mathbb{P}(\text{BBAA}) = \mathbb{P}(\text{BABA}) = \mathbb{P}(\text{BAAB}) = Z\,. \end{cases}$$

*Then* $0 < \overline{Y} < Y > 1.5Z > 0$*.*

**Lemma 2** (5 ancestral)**.** *Suppose* $(\{1,2,3,4,5\})$*, assume that the allele in lineage 5 is ancestral and denote*

$$\begin{cases} \mathbb{P}(\text{BAAAA}) = \mathbb{P}(\text{ABAAA}) = \mathbb{P}(\text{AABAA}) = \mathbb{P}(\text{AAABA}) = Y\,, \\ \mathbb{P}(\text{AAAAB}) = \overline{Y}\,, \\ \mathbb{P}(\text{ABBAA}) = \mathbb{P}(\text{ABABA}) = \mathbb{P}(\text{ABAAB}) = \mathbb{P}(\text{AABBA}) = \mathbb{P}(\text{AABAB}) = \mathbb{P}(\text{AAABB}) = Z\,, \\ \mathbb{P}(\text{BBAAA}) = \mathbb{P}(\text{BABAA}) = \mathbb{P}(\text{BAABA}) = \mathbb{P}(\text{BAAAB}) = \overline{Z}\,. \end{cases}$$

*Then* $0 < \overline{Y} < Y > 2Z > 2\overline{Z} > 0$*.*

**Lemma 3** (4 ancestral)**.** *Suppose* $(\{1,2,3\},\{4\})$*, assume that the allele in lineage 4 is ancestral and denote*

$$\begin{cases} \mathbb{P}(\text{AAAB}) = X\,, \\ \mathbb{P}(\text{BAAA}) = \mathbb{P}(\text{ABAA}) = \mathbb{P}(\text{AABA}) = Y\,, \\ \mathbb{P}(\text{BBAA}) = \mathbb{P}(\text{BABA}) = \mathbb{P}(\text{BAAB}) = Z\,. \end{cases}$$

*Then* $X > 0 < Z < Y$*.*

**Lemma 3** (3 ancestral)**.** *Suppose* $(\{1,2,3\},\{4\})$*, assume that the allele in lineage 3 is ancestral and denote*

$$\begin{cases} \mathbb{P}(\mathrm{AAAB}) = X\,, \\ \mathbb{P}(\mathrm{BAAA}) = \mathbb{P}(\mathrm{ABAA}) = Y\,, \\ \mathbb{P}(\mathrm{AABA}) = \overline{Y} \\ \mathbb{P}(\mathrm{BAAB}) = \mathbb{P}(\mathrm{BABA}) = Z\,, \\ \mathbb{P}(\mathrm{BBAA}) = \overline{Z}\,. \end{cases}$$

*Then* $Y > \overline{Z} > Z > 0$*,* $Y > \overline{Y} > Z$*,* $\overline{Y} < X > 1.5Z$*,* $Y \overset{*}{<} X \overset{*}{>} 1.5\overline{Z}$ *and* $X + Z \overset{*}{>} Y + \overline{Z}$*.*
*The inequalities marked with an asterisk require the synchronization assumption (2).*

**Lemma 4** (5 ancestral)**.** *Suppose* $(\{1,2,3,4\},\{5\})$*, assume that the allele in lineage 5 is ancestral and denote*

$$\begin{cases} \mathbb{P}(\mathrm{AAAAB}) = W\,, \\ \mathbb{P}(\mathrm{BAAAA}) = \mathbb{P}(\mathrm{ABAAA}) = \mathbb{P}(\mathrm{AABAA}) = \mathbb{P}(\mathrm{AAABA}) = X\,, \\ \mathbb{P}(\mathrm{BAAAB}) = \mathbb{P}(\mathrm{ABAAB}) = \mathbb{P}(\mathrm{AABAB}) = \mathbb{P}(\mathrm{AAABB}) = Y\,, \\ \mathbb{P}(\mathrm{BBAAA}) = \mathbb{P}(\mathrm{BABAA}) = \mathbb{P}(\mathrm{BAABA}) = \mathbb{P}(\mathrm{ABBAA}) = \mathbb{P}(\mathrm{ABABA}) = \mathbb{P}(\mathrm{AABBA}) = Z\,. \end{cases}$$

*Then* $W > 0 < Y < X > 1.5Z > 0$*.*

**Lemma 4** (4 ancestral)**.** *Suppose* $(\{1,2,3,4\},\{5\})$*, assume that the allele in lineage 4 is ancestral and denote*

$$\begin{cases} \mathbb{P}(\mathrm{AAAAB}) = W\,, \\ \mathbb{P}(\mathrm{BAAAA}) = \mathbb{P}(\mathrm{ABAAA}) = \mathbb{P}(\mathrm{AABAA}) = X\,, \\ \mathbb{P}(\mathrm{AAABA}) = \overline{X}\,, \\ \mathbb{P}(\mathrm{BAAAB}) = \mathbb{P}(\mathrm{ABAAB}) = \mathbb{P}(\mathrm{AABAB}) = Y\,, \\ \mathbb{P}(\mathrm{AAABB}) = \overline{Y}\,, \\ \mathbb{P}(\mathrm{BBAAA}) = \mathbb{P}(\mathrm{BABAA}) = \mathbb{P}(\mathrm{ABBAA}) = Z\,, \\ \mathbb{P}(\mathrm{BAABA}) = \mathbb{P}(\mathrm{ABABA}) = \mathbb{P}(\mathrm{AABBA}) = \overline{Z}\,. \end{cases}$$

*Then* $0 < Y < X > 1.5Z > 1.5\overline{Z} > 0$*,* $\overline{Y} > \overline{Z}$*,* $X > \overline{X} > 0 < W$*,* $\overline{Y} + Z > Y + \overline{Z}$*,* $W \overset{*}{>} X$ *and* $W + \overline{Z} + 2Y \overset{*}{>} X + \overline{Y} + 2Z$*.*
*The inequality marked with an asterisk requires the synchronization assumption (2).*

**Lemma 5** (4 ancestral)**.** *Suppose* $((\{1,2\},\{3\}),\{4\})$*, assume that the allele in lineage 4 is ancestral and denote*

$$\begin{cases} \mathbb{P}(\mathrm{AAAB}) = V\,, \\ \mathbb{P}(\mathrm{AABA}) = W\,, \\ \mathbb{P}(\mathrm{BAAA}) = \mathbb{P}(\mathrm{ABAA}) = X\,, \\ \mathbb{P}(\mathrm{BBAA}) = Y\,, \\ \mathbb{P}(\mathrm{ABBA}) = \mathbb{P}(\mathrm{BABA}) = Z\,. \end{cases}$$

*Then* $V > 0 < Z < W$*,* $X > Z < Y$*,* $X \overset{*}{<} W \overset{*}{>} Y$ *and* $W + Z \overset{*}{=} X + Y$*.*
*The relations marked with an asterisk require the synchronization assumption (2).*

**Lemma 5** (3 ancestral)**.** *Suppose* $((\{1,2\},\{3\}),\{4\})$*, assume that the allele in lineage 3 is ancestral and denote*

$$\begin{cases} \mathbb{P}(\mathrm{AAAB}) = V\,, \\ \mathbb{P}(\mathrm{AABA}) = W\,, \\ \mathbb{P}(\mathrm{BAAA}) = \mathbb{P}(\mathrm{ABAA}) = X\,, \\ \mathbb{P}(\mathrm{BBAA}) = Y\,, \\ \mathbb{P}(\mathrm{ABBA}) = \mathbb{P}(\mathrm{BABA}) = Z\,. \end{cases}$$

*Then* $V > 1.5Z < 1.5W$*,* $X > Z < Y$*,* $Z > 0$*,* $X \overset{*}{<} V \overset{*}{>} Y$ *and* $V + Z \overset{*}{>} X + Y \overset{*}{>} W + Z$*.*
*The inequalities marked with an asterisk require the synchronization assumption (2).*

**Lemma 5** (2 ancestral)**.** *Suppose* $((\{1,2\},\{3\}),\{4\})$*, assume that the allele in lineage 2 is ancestral and denote*

$$\begin{cases} \mathbb{P}(\mathrm{AAAB}) = V\,, \\ \mathbb{P}(\mathrm{AABA}) = W\,, \\ \mathbb{P}(\mathrm{BAAA}) = X\,, \\ \mathbb{P}(\mathrm{ABAA}) = \overline{X}\,, \\ \mathbb{P}(\mathrm{BBAA}) = Y\,, \\ \mathbb{P}(\mathrm{ABBA}) = Z \\ \mathbb{P}(\mathrm{BABA}) = \overline{Z}\,. \end{cases}$$

*Then* $0 < 1.5Z < V > \overline{X} < W$, $V > \overline{Z} < W$, $X > \overline{X} > Z < Y$, $X > \overline{Z} > Z$, $W \overset{*}{<} V \overset{*}{>} 1.5\overline{Z}$, $X \overset{*}{<} W \overset{*}{>} Y$ *and* $W + Z \overset{*}{>} X + Y$.
*The inequalities marked with an asterisk require the synchronization assumption (2).*

**Lemma 6** (5 ancestral)**.** *Suppose* $((\{1,2,3\},\{4\}),\{5\})$*, assume that the allele in lineage 5 is ancestral and denote*

$$\begin{cases} \mathbb{P}(\mathrm{AAAAB}) = T\,, \\ \mathbb{P}(\mathrm{AAABA}) = U\,, \\ \mathbb{P}(\mathrm{BAAAA}) = \mathbb{P}(\mathrm{ABAAA}) = \mathbb{P}(\mathrm{AABAA}) = V\,, \\ \mathbb{P}(\mathrm{AAABB}) = W\,, \\ \mathbb{P}(\mathrm{BAAAB}) = \mathbb{P}(\mathrm{ABAAB}) = \mathbb{P}(\mathrm{AABAB}) = X\,, \\ \mathbb{P}(\mathrm{BBAAA}) = \mathbb{P}(\mathrm{BABAA}) = \mathbb{P}(\mathrm{ABBAA}) = Y\,, \\ \mathbb{P}(\mathrm{BAABA}) = \mathbb{P}(\mathrm{ABABA}) = \mathbb{P}(\mathrm{AABBA}) = Z\,. \end{cases}$$

*Then* $V > Y > Z < X < W$, $U > Z > 0 < T$, $U \overset{*}{>} Y \overset{*}{<} V$, $W - X \overset{*}{<} U - V \overset{*}{>} Y - Z$ *and* $U + X + 2Z \overset{*}{=} V + W + 2Y$.
*The relations marked with an asterisk require the synchronization assumption (2).*

**Lemma 6** (4 ancestral)**.** *Suppose* $((\{1,2,3\},\{4\}),\{5\})$*, assume that the allele in lineage 4 is ancestral and denote*

$$\begin{cases} \mathbb{P}(\mathrm{AAAAB}) = T\,, \\ \mathbb{P}(\mathrm{AAABA}) = U\,, \\ \mathbb{P}(\mathrm{BAAAA}) = \mathbb{P}(\mathrm{ABAAA}) = \mathbb{P}(\mathrm{AABAA}) = V\,, \\ \mathbb{P}(\mathrm{AAABB}) = W\,, \\ \mathbb{P}(\mathrm{BAAAB}) = \mathbb{P}(\mathrm{ABAAB}) = \mathbb{P}(\mathrm{AABAB}) = X\,, \\ \mathbb{P}(\mathrm{BBAAA}) = \mathbb{P}(\mathrm{BABAA}) = \mathbb{P}(\mathrm{ABBAA}) = Y\,, \\ \mathbb{P}(\mathrm{BAABA}) = \mathbb{P}(\mathrm{ABABA}) = \mathbb{P}(\mathrm{AABBA}) = Z\,. \end{cases}$$

*Then* $0 < X < V > Y > Z > 0$, $W > Z$, $U > 0 < T$, $W + Y > X + Z$, $T + Z + 2X \overset{*}{>} V + W + 2Y \overset{*}{>} U + X + 2Z$ *and* $V \overset{*}{<} T \overset{*}{>} U$.
*The inequalities marked with an asterisk require the synchronization assumption (2).*

**Lemma 7** (4 ancestral)**.** *Suppose* $(\{1,2\},\{3,4\})$*, assume that the allele in lineage 4 is ancestral and denote*

$$\begin{cases} \mathbb{P}(\mathrm{AABA}) = W\,, \\ \mathbb{P}(\mathrm{AAAB}) = \overline{W}\,, \\ \mathbb{P}(\mathrm{BAAA}) = \mathbb{P}(\mathrm{ABAA}) = X\,, \\ \mathbb{P}(\mathrm{BBAA}) = Y\,, \\ \mathbb{P}(\mathrm{BABA}) = \mathbb{P}(\mathrm{BAAB}) = Z\,. \end{cases}$$

*Then* $0 < \overline{W} < W > 1.5Z < X$, $Y > Z > 0$ *and* $Y - Z \overset{*}{>} W - X$.
*The inequality marked with an asterisk requires the synchronization assumption (2).*

**Lemma 8** (5 ancestral)**.** *Suppose* $(\{1,2,3\},\{4,5\})$*, assume that the allele in lineage 5 is ancestral and denote*

$$\begin{cases} \mathbb{P}(\mathrm{AAABA}) = V\,, \\ \mathbb{P}(\mathrm{AAAAB}) = \overline{V}\,, \\ \mathbb{P}(\mathrm{BAAAA}) = \mathbb{P}(\mathrm{ABAAA}) = \mathbb{P}(\mathrm{AABAA}) = W\,, \\ \mathbb{P}(\mathrm{AAABB}) = X\,, \\ \mathbb{P}(\mathrm{BBAAA}) = \mathbb{P}(\mathrm{BABAA}) = \mathbb{P}(\mathrm{ABBAA}) = Y\,, \\ \mathbb{P}(\mathrm{BAABA}) = \mathbb{P}(\mathrm{ABABA}) = \mathbb{P}(\mathrm{AABBA}) = Z\,, \\ \mathbb{P}(\mathrm{BAAAB}) = \mathbb{P}(\mathrm{ABAAB}) = \mathbb{P}(\mathrm{AABAB}) = \overline{Z}\,. \end{cases}$$

*Then* $0 < 3\overline{V} < 3V > 6Z < 4W$*,* $X > \overline{Z} < Z < Y$*,* $\overline{Z} > 0$ *and* $W + X + 2Y \overset{*}{>} V + \overline{Z} + 2Z$*.*
*The inequality marked with an asterisk requires the synchronization assumption (2).*

While the equations $(S0)$–$(S19)$ and $(A0)$–$(A26)$ will be a little different under the condition that 5 carries the ancestral allele, only in three instances the final predicted table is affected. In the case $5 \to 4$ in tree $S$ the signature becomes $(+\,+\,0\,0\,|\,0\,?\,0\,+)$, in the case $4 \to 5$ in tree $S$ the signature becomes $(+\,+\,0\,0\,|\,0\,+\,0\,+)$, and in the case $2 \to 5$ in tree $A$ the signature becomes $(-\,+\,+\,|\,+)$. The question mark stands for an inconsistent statistic whose behaviour depends on graph parameters.

#### Appendix 3: predicted effects of $\Delta$-statistics without the synchronization assumption (2)

Table S1: Predicted effects on tree $S = (((1,2),(3,4)),5)$ without the synchronization assumption (2). Unidirectional gene flow events not listed in the table produce a zero signature.

| | ${}_S\Delta^*_{1-6}$ | ${}_S\Delta^*_{2-6}$ | ${}_S\Delta^*_{3-5}$ | ${}_S\Delta^*_{4-5}$ |
|---|---|---|---|---|
| $1 \to 3$ | + | + | + | 0 |
| $3 \to 1$ | + | 0 | + | + |
| $1 \to 4$ | − | − | 0 | + |
| $4 \to 1$ | − | 0 | + | + |
| $2 \to 3$ | + | + | − | 0 |
| $3 \to 2$ | 0 | + | − | − |
| $2 \to 4$ | − | − | 0 | − |
| $4 \to 2$ | 0 | − | − | − |
| $12345 \to 1$<br>$34 \leftrightarrow 2$<br>$1234 \to 1$<br>$1 \leftrightarrow 5$ | 0 | 0 | − | − |
| $12345 \to 2$<br>$34 \leftrightarrow 1$<br>$1234 \to 2$<br>$2 \leftrightarrow 5$ | 0 | 0 | + | + |
| $12345 \to 3$<br>$12 \leftrightarrow 4$<br>$1234 \to 3$<br>$3 \leftrightarrow 5$ | − | − | 0 | 0 |
| $12345 \to 4$<br>$12 \leftrightarrow 3$<br>$1234 \to 4$<br>$4 \leftrightarrow 5$ | + | + | 0 | 0 |

Table S2: Predicted effects on tree $A = ((((1,2),3),4),5)$ without the synchronization assumption (2). Unidirectional gene flow events not listed in the table produce a zero signature.

| | ${}_A\Delta^*_{1-2}$ | ${}_A\Delta^*_{1-3}$ | ${}_A\Delta^*_{2-3}$ |
|---|---|---|---|
| $1 \leftrightarrow 3$<br>$123 \to 2$ | + | + | 0 |
| $2 \leftrightarrow 3$<br>$123 \to 1$ | − | − | 0 |
| $1 \to 4$ | − | 0 | + |
| $4 \to 1$ | − | − | + |
| $2 \to 4$ | + | 0 | − |
| $4 \to 2$ | + | + | − |
| $1 \to 5$ | 0 | − | − |
| $5 \to 1$<br>$1234 \to 1$<br>$12345 \to 1$ | − | − | − |
| $2 \to 5$ | 0 | + | + |
| $5 \to 2$<br>$1234 \to 2$<br>$12345 \to 2$ | + | + | + |

Table S3: Predicted effects on tree $Q = (((1,2),3),(4,5))$ without the synchronization assumption (2). Unidirectional gene flow events not listed in the table produce a zero signature.

| | ${}_Q\Delta^*_{1-6}$ | ${}_Q\Delta^*_{2-6}$ | ${}_Q\Delta^*_{3-5}$ | ${}_Q\Delta^*_{4-5}$ |
|---|---|---|---|---|
| $1 \to 4$ | + | + | + | 0 |
| $4 \to 1$ | + | 0 | + | + |
| $1 \to 5$ | − | − | 0 | + |
| $5 \to 1$ | − | 0 | + | + |
| $2 \to 4$ | + | + | − | 0 |
| $4 \to 2$ | 0 | + | − | − |
| $2 \to 5$ | − | − | 0 | − |
| $5 \to 2$ | 0 | − | − | − |
| $2 \leftrightarrow 45$<br>$1 \leftrightarrow 3$<br>$123 \to 2$<br>$12345 \to 2$ | 0 | 0 | − | − |
| $1 \leftrightarrow 45$<br>$2 \leftrightarrow 3$<br>$123 \to 1$<br>$12345 \to 1$ | 0 | 0 | + | + |
| $5 \leftrightarrow 12$<br>$4 \leftrightarrow 3$ | − | − | 0 | 0 |
| $4 \leftrightarrow 12$<br>$5 \leftrightarrow 3$ | + | + | 0 | 0 |

#### Appendix 4: Simulations with fewer independent patterns

In Tables S4 – S9 (over following three pages) we present the results from simulations using 100,000 or 10,000 independent allelic patterns.

We observe that the statistics using the singleton patterns and synchronization assumption (2) start to deteriorate already at sample size 100,000. In particular, ${}_S\Delta^*_{5+7}$, ${}_S\Delta^*_{6+8}$, ${}_Q\Delta^*_{5+7}$ and ${}_Q\Delta^*_{6+8}$ that have proportionally the highest usage of singleton patterns suffer. The ability to distinguish gene flow events is impaired. Disregarding those statistics, in other words restricting to Tables S1, S2 and S3 in Appendix 3, means there is less resolution among gene flow events that can be identified, but as can be seen from the simulations, those predictions can still be done accurately.

At sample size 10,000, the $\Delta$-statistic method does not work well anymore. Restricting to Tables S1, S2 and S3 in Appendix 3 brings a notable increase in power, but still only about half of the gene flow events were recognized.

Comparable analyses for the $D_{\mathrm{FOIL}}$ method can be found in Appendix 5.

Table S4: Simulation results for the symmetric tree $S$ using 100,000 independent patterns. For each statistic under each scenario we report the value, the 99% Wald confidence interval with radius $2.576 \times \sqrt{4n(L)n(R)/(n(L)+n(R))^3}$, the $Z$-score and the classification. Misclassifications are highlighted in gray. Compared to Table 2 in the main text, thirteen scenarios were misclassified. Compared to Table S1 in Appendix 3, everything was correct.

| | $_S\Delta^*_{1-6}$ | $_S\Delta^*_{2-6}$ | $_S\Delta^*_{3-5}$ | $_S\Delta^*_{4-5}$ | $_S\Delta^*_{5+7}$ | $_S\Delta^*_{6+8}$ | $_S\Delta^*_{3+4-5+7}$ | $_S\Delta^*_{1+2-6+8}$ |
|---|---|---|---|---|---|---|---|---|
| $1 \to 3$ | 0.370±0.046<br>19.37 : + | 0.266±0.050<br>13.27 : + | 0.149±0.056<br>6.853 : + | −0.023±0.063<br>−0.925 : 0 | −0.016±0.017<br>−2.605 : − | −0.064±0.016<br>−10.57 : − | −0.005±0.016<br>−0.857 : 0 | −0.003±0.016<br>−0.460 : 0 |
| $3 \to 1$ | 0.155±0.055<br>7.154 : + | 0.018±0.063<br>0.731 : 0 | 0.396±0.047<br>20.37 : + | 0.316±0.051<br>15.36 : + | −0.064±0.016<br>−10.61 : − | −0.001±0.016<br>−0.198 : 0 | 0.001±0.016<br>0.189 : 0 | 0.012±0.016<br>1.993 : 0 |
| $1 \to 4$ | −0.390±0.046<br>−20.28 : − | −0.288±0.051<br>−14.18 : − | 0.005±0.062<br>0.214 : 0 | 0.165±0.054<br>7.752 : + | −0.012±0.016<br>−1.969 : 0 | 0.063±0.016<br>10.33 : + | 0.002±0.016<br>0.332 : 0 | −0.002±0.016<br>−0.267 : 0 |
| $4 \to 1$ | −0.166±0.056<br>−7.652 : − | 0.009±0.065<br>0.348 : 0 | 0.294±0.051<br>14.37 : + | 0.397±0.046<br>20.58 : + | −0.055±0.016<br>−9.096 : − | 0.015±0.016<br>2.382 : 0 | 0.009±0.016<br>1.549 : 0 | 0.002±0.016<br>0.284 : 0 |
| $2 \to 3$ | 0.298±0.051<br>14.56 : + | 0.406±0.046<br>21.02 : + | −0.174±0.055<br>−8.118 : − | −0.002±0.063<br>−0.073 : 0 | 0.015±0.016<br>2.443 : 0 | −0.050±0.016<br>−8.359 : − | 0.000±0.016<br>0.079 : 0 | 0.014±0.016<br>2.447 : 0 |
| $3 \to 2$ | −0.030±0.062<br>−1.227 : 0 | 0.141±0.055<br>6.538 : + | −0.383±0.047<br>−19.62 : − | −0.276±0.052<br>−13.39 : − | 0.069±0.016<br>11.40 : + | −0.008±0.016<br>−1.330 : 0 | 0.008±0.016<br>1.379 : 0 | 0.001±0.016<br>0.230 : 0 |
| $2 \to 4$ | −0.278±0.051<br>−13.75 : − | −0.380±0.047<br>−19.67 : − | 0.018±0.062<br>0.743 : 0 | −0.142±0.055<br>−6.615 : − | 0.014±0.016<br>2.211 : 0 | 0.061±0.016<br>10.17 : + | 0.003±0.016<br>0.489 : 0 | −0.001±0.016<br>−0.089 : 0 |
| $4 \to 2$ | 0.018±0.063<br>0.748 : 0 | −0.161±0.055<br>−7.570 : − | −0.261±0.051<br>−12.75 : − | −0.382±0.047<br>−19.62 : − | 0.066±0.016<br>10.97 : + | 0.023±0.017<br>3.685 : + | 0.006±0.016<br>1.088 : 0 | 0.010±0.016<br>1.633 : 0 |
| $12345 \to 1$ | 0.026±0.066<br>0.999 : 0 | 0.053±0.065<br>2.119 : 0 | −0.275±0.052<br>−13.15 : − | −0.253±0.052<br>−12.14 : − | 0.067±0.016<br>11.50 : + | 0.007±0.017<br>1.109 : 0 | 0.025±0.015<br>4.364 : + | 0.011±0.016<br>1.856 : 0 |
| $34 \to 2$ | 0.002±0.081<br>0.063 : 0 | −0.000±0.074<br>−0.000 : 0 | −0.273±0.047<br>−14.45 : − | −0.275±0.047<br>−14.52 : − | 0.051±0.015<br>9.365 : + | −0.010±0.018<br>−1.461 : 0 | 0.005±0.014<br>0.971 : 0 | −0.009±0.017<br>−1.425 : 0 |
| $2 \to 34$ | −0.023±0.064<br>−0.955 : 0 | −0.009±0.059<br>−0.408 : 0 | −0.105±0.052<br>−5.267 : − | −0.099±0.052<br>−4.883 : − | 0.013±0.015<br>2.395 : 0 | 0.012±0.017<br>1.784 : 0 | −0.002±0.015<br>−0.430 : 0 | 0.009±0.017<br>1.383 : 0 |
| $1234 \to 1$ | 0.037±0.065<br>1.473 : 0 | 0.024±0.063<br>0.978 : 0 | −0.164±0.055<br>−7.613 : − | −0.175±0.055<br>−8.080 : − | 0.028±0.016<br>4.628 : + | −0.003±0.016<br>−0.540 : 0 | 0.002±0.016<br>0.294 : 0 | 0.000±0.016<br>0.073 : 0 |
| $5 \to 1$ | −0.054±0.067<br>−2.081 : 0 | −0.031±0.064<br>−1.238 : 0 | −0.602±0.034<br>−37.17 : − | −0.589±0.034<br>−36.52 : − | 0.041±0.016<br>6.919 : + | 0.001±0.016<br>0.178 : 0 | −0.113±0.015<br>−19.58 : − | −0.004±0.016<br>−0.613 : 0 |
| $1 \to 5$ | −0.012±0.065<br>−0.499 : 0 | −0.012±0.064<br>−0.472 : 0 | −0.202±0.055<br>−9.411 : − | −0.199±0.054<br>−9.323 : − | −0.005±0.016<br>−0.758 : 0 | −0.010±0.016<br>−1.677 : 0 | −0.036±0.016<br>−6.001 : − | −0.011±0.016<br>−1.893 : 0 |
| $12345 \to 2$ | 0.032±0.064<br>1.300 : 0 | 0.009±0.065<br>0.377 : 0 | 0.257±0.052<br>12.34 : + | 0.242±0.053<br>11.60 : + | −0.069±0.016<br>−11.86 : − | −0.002±0.016<br>−0.384 : 0 | −0.029±0.015<br>−5.064 : − | 0.000±0.016<br>0.037 : 0 |
| $34 \to 1$ | 0.029±0.075<br>0.987 : 0 | 0.006±0.083<br>0.192 : 0 | 0.278±0.048<br>14.47 : + | 0.271±0.048<br>14.03 : + | −0.040±0.015<br>−7.283 : − | 0.001±0.018<br>0.201 : 0 | 0.005±0.014<br>0.830 : 0 | 0.003±0.017<br>0.461 : 0 |
| $1 \to 34$ | −0.013±0.059<br>−0.570 : 0 | −0.039±0.062<br>−1.610 : 0 | 0.084±0.052<br>4.176 : + | 0.067±0.052<br>3.336 : + | −0.008±0.015<br>−1.432 : 0 | −0.001±0.017<br>−0.149 : 0 | 0.004±0.015<br>0.638 : 0 | −0.005±0.017<br>−0.730 : 0 |
| $1234 \to 2$ | −0.019±0.063<br>−0.799 : 0 | −0.032±0.064<br>−1.282 : 0 | 0.183±0.055<br>8.454 : + | 0.172±0.055<br>7.997 : + | −0.036±0.016<br>−5.997 : − | 0.002±0.016<br>0.333 : 0 | −0.008±0.016<br>−1.435 : 0 | −0.001±0.016<br>−0.188 : 0 |
| $5 \to 2$ | 0.012±0.063<br>0.487 : 0 | 0.011±0.067<br>0.410 : 0 | 0.602±0.033<br>37.69 : + | 0.603±0.033<br>37.71 : + | −0.047±0.016<br>−8.070 : − | −0.005±0.016<br>−0.831 : 0 | 0.111±0.015<br>19.27 : + | −0.004±0.016<br>−0.602 : 0 |
| $2 \to 5$ | −0.015±0.064<br>−0.595 : 0 | −0.004±0.066<br>−0.152 : 0 | 0.192±0.055<br>8.983 : + | 0.202±0.055<br>9.411 : + | −0.000±0.016<br>−0.024 : 0 | 0.003±0.016<br>0.419 : 0 | 0.031±0.016<br>5.111 : + | 0.001±0.016<br>0.234 : 0 |
| $12345 \to 3$ | −0.282±0.052<br>−13.50 : − | −0.274±0.052<br>−13.07 : − | 0.032±0.067<br>1.256 : 0 | 0.046±0.065<br>1.823 : 0 | −0.004±0.017<br>−0.722 : 0 | 0.070±0.016<br>11.96 : + | 0.000±0.016<br>0.037 : 0 | 0.026±0.015<br>4.486 : + |
| $12 \to 4$ | −0.229±0.048<br>−12.02 : − | −0.218±0.048<br>−11.47 : − | −0.078±0.083<br>−2.419 : 0 | −0.043±0.077<br>−1.430 : 0 | −0.001±0.018<br>−0.215 : 0 | 0.047±0.015<br>8.552 : + | −0.007±0.018<br>−1.026 : 0 | 0.010±0.015<br>1.757 : 0 |
| $4 \to 12$ | −0.088±0.053<br>−4.280 : − | −0.089±0.053<br>−4.347 : − | −0.026±0.064<br>−1.060 : 0 | −0.025±0.060<br>−1.085 : 0 | −0.002±0.017<br>−0.350 : 0 | 0.008±0.015<br>1.517 : 0 | −0.006±0.017<br>−0.914 : 0 | −0.005±0.015<br>−0.821 : 0 |
| $1234 \to 3$ | −0.164±0.055<br>−7.673 : − | −0.179±0.055<br>−8.313 : − | 0.017±0.065<br>0.698 : 0 | −0.000±0.063<br>−0.000 : 0 | −0.002±0.016<br>−0.321 : 0 | 0.023±0.016<br>3.909 : + | −0.001±0.016<br>−0.146 : 0 | −0.003±0.016<br>−0.536 : 0 |
| $5 \to 3$ | −0.616±0.033<br>−39.01 : − | −0.620±0.032<br>−39.25 : − | −0.030±0.066<br>−1.152 : 0 | −0.036±0.063<br>−1.487 : 0 | 0.022±0.016<br>3.487 : + | 0.054±0.015<br>9.212 : + | 0.017±0.016<br>2.795 : + | −0.112±0.015<br>−19.40 : − |
| $3 \to 5$ | −0.169±0.055<br>−7.905 : − | −0.181±0.055<br>−8.391 : − | −0.000±0.065<br>−0.000 : 0 | −0.013±0.064<br>−0.514 : 0 | 0.009±0.016<br>1.557 : 0 | −0.010±0.016<br>−1.700 : 0 | 0.008±0.016<br>1.414 : 0 | −0.037±0.016<br>−6.238 : − |
| $12345 \to 4$ | 0.295±0.052<br>14.07 : + | 0.299±0.052<br>14.26 : + | 0.019±0.063<br>0.781 : 0 | 0.026±0.065<br>1.023 : 0 | −0.012±0.016<br>−1.891 : 0 | −0.055±0.016<br>−9.413 : − | −0.009±0.016<br>−1.419 : 0 | −0.009±0.015<br>−1.481 : 0 |
| $12 \to 3$ | 0.270±0.048<br>14.00 : + | 0.254±0.048<br>13.30 : + | −0.031±0.075<br>−1.065 : 0 | −0.069±0.082<br>−2.185 : 0 | −0.001±0.018<br>−0.160 : 0 | −0.045±0.015<br>−8.231 : − | −0.006±0.017<br>−0.854 : 0 | −0.002±0.014<br>−0.429 : 0 |
| $3 \to 12$ | 0.130±0.053<br>6.358 : + | 0.116±0.053<br>5.689 : + | 0.022±0.058<br>0.985 : 0 | 0.007±0.063<br>0.289 : 0 | 0.005±0.017<br>0.743 : 0 | −0.010±0.015<br>−1.843 : 0 | 0.007±0.017<br>1.081 : 0 | 0.008±0.015<br>1.447 : 0 |
| $1234 \to 4$ | 0.171±0.055<br>7.936 : + | 0.175±0.056<br>8.022 : + | 0.014±0.064<br>0.566 : 0 | 0.014±0.064<br>0.546 : 0 | 0.007±0.016<br>1.133 : 0 | −0.017±0.016<br>−2.857 : − | 0.008±0.016<br>1.391 : 0 | 0.009±0.016<br>1.508 : 0 |
| $5 \to 4$ | 0.595±0.034<br>37.00 : + | 0.590±0.034<br>36.66 : + | 0.009±0.064<br>0.370 : 0 | −0.005±0.067<br>−0.180 : 0 | −0.003±0.016<br>−0.474 : 0 | −0.041±0.016<br>−7.086 : − | −0.003±0.016<br>−0.419 : 0 | 0.112±0.015<br>19.41 : + |
| $4 \to 5$ | 0.164±0.056<br>7.472 : + | 0.145±0.056<br>6.636 : + | 0.057±0.065<br>2.245 : 0 | 0.033±0.066<br>1.288 : 0 | 0.001±0.016<br>0.085 : 0 | −0.010±0.016<br>−1.630 : 0 | 0.006±0.016<br>0.922 : 0 | 0.014±0.016<br>2.261 : 0 |

Table S5: Simulation results for the symmetric tree $S$ using 10,000 independent patterns. For each statistic under each scenario we report the value, the 99% Wald confidence interval with radius $2.576 \times \sqrt{4n(L)n(R)/(n(L)+n(R))^3}$, the $Z$-score and the classification. Misclassifications are highlighted in gray. Compared to Table 2 in the main text, only five scenarios were correctly identified. Compared to Table S1 in Appendix 3, nineteen of the admixture scenarios were correct.

| | ${}_S\Delta^*_{1-6}$ | ${}_S\Delta^*_{2-6}$ | ${}_S\Delta^*_{3-5}$ | ${}_S\Delta^*_{4-5}$ | ${}_S\Delta^*_{5+7}$ | ${}_S\Delta^*_{6+8}$ | ${}_S\Delta^*_{3+4-5+7}$ | ${}_S\Delta^*_{1+2-6+8}$ |
|---|---|---|---|---|---|---|---|---|
| 1 → 3 | 0.357±0.145<br>5.948 : + | 0.283±0.156<br>4.481 : + | 0.187±0.173<br>2.734 : + | 0.067±0.192<br>0.894 : 0 | −0.043±0.051<br>−2.197 : 0 | −0.072±0.049<br>−3.802 : − | −0.022±0.050<br>−1.153 : 0 | −0.011±0.048<br>−0.575 : 0 |
| 3 → 1 | 0.191±0.173<br>2.796 : + | 0.085±0.201<br>1.093 : 0 | 0.333±0.154<br>5.260 : + | 0.243±0.165<br>3.693 : + | −0.042±0.049<br>−2.242 : 0 | −0.010±0.051<br>−0.508 : 0 | 0.007±0.048<br>0.370 : 0 | 0.011±0.050<br>0.556 : 0 |
| 1 → 4 | −0.363±0.152<br>−5.744 : − | −0.285±0.162<br>−4.371 : − | −0.107±0.198<br>−1.389 : 0 | 0.029±0.180<br>0.418 : 0 | 0.010±0.052<br>0.520 : 0 | 0.049±0.049<br>2.597 : + | 0.005±0.051<br>0.273 : 0 | −0.007±0.048<br>−0.389 : 0 |
| 4 → 1 | −0.105±0.186<br>−1.451 : 0 | −0.049±0.203<br>−0.629 : 0 | 0.441±0.151<br>6.770 : + | 0.464±0.145<br>7.336 : + | −0.087±0.050<br>−4.552 : − | 0.016±0.051<br>0.806 : 0 | −0.006±0.049<br>−0.320 : 0 | 0.005±0.050<br>0.251 : 0 |
| 2 → 3 | 0.209±0.156<br>3.391 : + | 0.356±0.142<br>6.059 : + | −0.205±0.169<br>−3.074 : − | 0.012±0.198<br>0.153 : 0 | 0.018±0.052<br>0.895 : 0 | −0.063±0.050<br>−3.309 : − | 0.000±0.050<br>0.019 : 0 | −0.005±0.049<br>−0.280 : 0 |
| 3 → 2 | −0.011±0.196<br>−0.152 : 0 | 0.092±0.166<br>1.420 : 0 | −0.439±0.143<br>−7.139 : − | −0.380±0.154<br>−5.914 : − | 0.068±0.050<br>3.493 : + | −0.015±0.051<br>−0.742 : 0 | −0.010±0.049<br>−0.530 : 0 | −0.007±0.050<br>−0.343 : 0 |
| 2 → 4 | −0.288±0.161<br>−4.426 : − | −0.371±0.143<br>−6.215 : − | −0.048±0.214<br>−0.581 : 0 | −0.221±0.180<br>−3.079 : − | −0.000±0.051<br>−0.000 : 0 | 0.082±0.049<br>4.353 : + | −0.019±0.050<br>−0.964 : 0 | 0.020±0.048<br>1.074 : 0 |
| 4 → 2 | 0.026±0.188<br>0.364 : 0 | −0.231±0.166<br>−3.502 : − | −0.246±0.163<br>−3.775 : − | −0.400±0.139<br>−6.812 : − | 0.035±0.049<br>1.849 : 0 | 0.033±0.051<br>1.691 : 0 | −0.026±0.048<br>−1.402 : 0 | 0.014±0.050<br>0.729 : 0 |
| 12345 → 1 | 0.055±0.213<br>0.662 : 0 | 0.110±0.200<br>1.406 : 0 | −0.308±0.152<br>−4.995 : − | −0.285±0.157<br>−4.499 : − | 0.095±0.048<br>5.104 : + | 0.007±0.051<br>0.336 : 0 | 0.041±0.048<br>2.254 : 0 | 0.016±0.051<br>0.838 : 0 |
| 34 → 2 | 0.010±0.252<br>0.098 : 0 | −0.017±0.236<br>−0.183 : 0 | −0.311±0.149<br>−5.112 : − | −0.319±0.148<br>−5.265 : − | 0.048±0.046<br>2.755 : + | −0.035±0.055<br>−1.648 : 0 | −0.004±0.045<br>−0.243 : 0 | −0.034±0.054<br>−1.643 : 0 |
| 2 → 34 | 0.041±0.197<br>0.535 : 0 | 0.118±0.177<br>1.721 : 0 | −0.076±0.163<br>−1.204 : 0 | −0.004±0.171<br>−0.066 : 0 | 0.015±0.046<br>0.858 : 0 | 0.026±0.053<br>1.255 : 0 | 0.009±0.045<br>0.500 : 0 | 0.038±0.052<br>1.874 : 0 |
| 1234 → 1 | −0.032±0.206<br>−0.399 : 0 | 0.006±0.197<br>0.076 : 0 | −0.188±0.168<br>−2.842 : − | −0.164±0.170<br>−2.467 : 0 | 0.014±0.049<br>0.751 : 0 | −0.003±0.051<br>−0.177 : 0 | −0.014±0.048<br>−0.740 : 0 | −0.005±0.050<br>−0.252 : 0 |
| 5 → 1 | −0.085±0.216<br>−1.007 : 0 | −0.082±0.197<br>−1.074 : 0 | −0.645±0.103<br>−12.44 : − | −0.647±0.102<br>−12.51 : − | 0.033±0.048<br>1.802 : 0 | 0.012±0.051<br>0.625 : 0 | −0.130±0.047<br>−7.075 : − | 0.002±0.050<br>0.116 : 0 |
| 1 → 5 | 0.127±0.204<br>1.591 : 0 | 0.145±0.207<br>1.784 : 0 | −0.196±0.177<br>−2.801 : − | −0.186±0.178<br>−2.661 : − | 0.022±0.050<br>1.141 : 0 | −0.003±0.050<br>−0.173 : 0 | −0.007±0.050<br>−0.363 : 0 | 0.012±0.049<br>0.627 : 0 |
| 12345 → 2 | 0.018±0.202<br>0.235 : 0 | −0.085±0.201<br>−1.093 : 0 | 0.368±0.158<br>5.593 : + | 0.298±0.163<br>4.503 : + | −0.075±0.047<br>−4.151 : − | −0.015±0.052<br>−0.740 : 0 | −0.024±0.047<br>−1.360 : 0 | −0.019±0.051<br>−0.945 : 0 |
| 34 → 1 | 0.141±0.226<br>1.591 : 0 | 0.105±0.240<br>1.124 : 0 | 0.284±0.147<br>4.798 : + | 0.265±0.148<br>4.458 : + | −0.034±0.045<br>−1.965 : 0 | 0.014±0.055<br>0.657 : 0 | 0.013±0.045<br>0.740 : 0 | 0.026±0.054<br>1.271 : 0 |
| 1 → 34 | −0.042±0.187<br>−0.580 : 0 | 0.056±0.203<br>0.709 : 0 | 0.063±0.162<br>1.008 : 0 | 0.128±0.160<br>2.058 : 0 | −0.041±0.046<br>−2.349 : 0 | −0.012±0.054<br>−0.563 : 0 | −0.025±0.045<br>−1.459 : 0 | −0.011±0.053<br>−0.530 : 0 |
| 1234 → 2 | −0.006±0.201<br>−0.078 : 0 | 0.053±0.211<br>0.653 : 0 | 0.076±0.178<br>1.104 : 0 | 0.114±0.173<br>1.689 : 0 | −0.009±0.049<br>−0.505 : 0 | −0.036±0.051<br>−1.821 : 0 | 0.005±0.048<br>0.258 : 0 | −0.032±0.050<br>−1.661 : 0 |
| 5 → 2 | 0.032±0.187<br>0.435 : 0 | −0.032±0.206<br>−0.399 : 0 | 0.662±0.098<br>13.10 : + | 0.656±0.100<br>12.76 : + | −0.059±0.048<br>−3.220 : − | 0.013±0.051<br>0.661 : 0 | 0.110±0.047<br>6.037 : + | 0.013±0.050<br>0.672 : 0 |
| 2 → 5 | 0.006±0.195<br>0.076 : 0 | 0.035±0.197<br>0.457 : 0 | 0.231±0.162<br>3.600 : + | 0.260±0.163<br>3.979 : + | −0.000±0.049<br>−0.000 : 0 | 0.026±0.050<br>1.319 : 0 | 0.041±0.049<br>2.188 : 0 | 0.028±0.050<br>1.436 : 0 |
| 12345 → 3 | −0.373±0.150<br>−5.949 : − | −0.344±0.152<br>−5.500 : − | 0.074±0.202<br>0.943 : 0 | 0.104±0.190<br>1.405 : 0 | 0.002±0.051<br>0.117 : 0 | 0.065±0.048<br>3.553 : + | 0.014±0.050<br>0.709 : 0 | 0.003±0.047<br>0.182 : 0 |
| 12 → 4 | −0.290±0.157<br>−4.572 : − | −0.325±0.153<br>−5.198 : − | −0.085±0.265<br>−0.825 : 0 | −0.188±0.252<br>−1.891 : 0 | 0.000±0.055<br>0.021 : 0 | 0.058±0.045<br>3.375 : + | −0.011±0.054<br>−0.544 : 0 | 0.012±0.045<br>0.686 : 0 |
| 4 → 12 | −0.121±0.163<br>−1.905 : 0 | −0.151±0.163<br>−2.364 : 0 | 0.169±0.188<br>2.292 : 0 | 0.115±0.178<br>1.664 : 0 | 0.016±0.053<br>0.771 : 0 | 0.031±0.046<br>1.732 : 0 | 0.036±0.051<br>1.841 : 0 | 0.009±0.046<br>0.526 : 0 |
| 1234 → 3 | −0.153±0.166<br>−2.343 : 0 | −0.099±0.168<br>−1.507 : 0 | 0.061±0.193<br>0.822 : 0 | 0.143±0.197<br>1.852 : 0 | −0.020±0.051<br>−1.035 : 0 | 0.049±0.049<br>2.587 : + | −0.007±0.050<br>−0.346 : 0 | 0.027±0.048<br>1.450 : 0 |
| 5 → 3 | −0.585±0.103<br>−11.89 : − | −0.589±0.102<br>−12.03 : − | 0.118±0.220<br>1.372 : 0 | 0.074±0.202<br>0.943 : 0 | −0.041±0.051<br>−2.075 : 0 | 0.066±0.048<br>3.576 : + | −0.029±0.050<br>−1.503 : 0 | −0.096±0.047<br>−5.301 : − |
| 3 → 5 | −0.212±0.177<br>−3.018 : − | −0.178±0.179<br>−2.533 : 0 | −0.078±0.199<br>−1.006 : 0 | −0.035±0.197<br>−0.457 : 0 | 0.002±0.050<br>0.096 : 0 | 0.015±0.050<br>0.782 : 0 | −0.005±0.049<br>−0.265 : 0 | −0.014±0.049<br>−0.717 : 0 |
| 12345 → 4 | 0.268±0.166<br>4.009 : + | 0.292±0.164<br>4.390 : + | −0.078±0.207<br>−0.967 : 0 | −0.043±0.220<br>−0.511 : 0 | 0.036±0.051<br>1.838 : 0 | −0.071±0.048<br>−3.846 : − | 0.028±0.051<br>1.461 : 0 | −0.027±0.048<br>−1.486 : 0 |
| 12 → 3 | 0.359±0.142<br>6.107 : + | 0.353±0.138<br>6.174 : + | −0.130±0.239<br>−1.399 : 0 | −0.111±0.258<br>−1.106 : 0 | −0.010±0.055<br>−0.490 : 0 | −0.072±0.045<br>−4.108 : − | −0.022±0.054<br>−1.027 : 0 | −0.007±0.045<br>−0.398 : 0 |
| 3 → 12 | 0.138±0.159<br>2.233 : 0 | 0.134±0.161<br>2.133 : 0 | 0.004±0.171<br>0.066 : 0 | −0.006±0.193<br>−0.075 : 0 | 0.022±0.053<br>1.082 : 0 | −0.039±0.046<br>−2.190 : 0 | 0.021±0.052<br>1.055 : 0 | −0.016±0.045<br>−0.926 : 0 |
| 1234 → 4 | 0.101±0.174<br>1.490 : 0 | 0.106±0.178<br>1.525 : 0 | 0.065±0.219<br>0.763 : 0 | 0.060±0.211<br>0.737 : 0 | −0.010±0.051<br>−0.526 : 0 | −0.024±0.049<br>−1.236 : 0 | −0.003±0.050<br>−0.173 : 0 | −0.007±0.049<br>−0.395 : 0 |
| 5 → 4 | 0.568±0.107<br>11.33 : + | 0.568±0.106<br>11.45 : + | −0.054±0.190<br>−0.737 : 0 | −0.031±0.202<br>−0.392 : 0 | 0.029±0.051<br>1.463 : 0 | −0.019±0.048<br>−1.003 : 0 | 0.022±0.050<br>1.152 : 0 | 0.135±0.047<br>7.379 : + |
| 4 → 5 | 0.149±0.169<br>2.252 : 0 | 0.109±0.170<br>1.652 : 0 | 0.023±0.196<br>0.303 : 0 | −0.032±0.206<br>−0.399 : 0 | 0.022±0.049<br>1.138 : 0 | −0.031±0.050<br>−1.589 : 0 | 0.021±0.049<br>1.103 : 0 | −0.008±0.050<br>−0.439 : 0 |

Table S6: Simulation results for the asymmetric tree $A$ using 100,000 independent patterns. For each statistic under each scenario we report the value, the 99% Wald confidence interval with radius $2.576 \times \sqrt{4n(L)n(R)/(n(L)+n(R))^3}$, the $Z$-score and the classification. Misclassification is highlighted in gray. Compared to Table 3 in the main text, only the admixture scenario $2 \to 5$ was misclassified. Compared to Table S2 in Appendix 3, everything was correct.

| | ${}_A\Delta^*_{1-2}$ | ${}_A\Delta^*_{1-3}$ | ${}_A\Delta^*_{2-3}$ | ${}_A\Delta^*_{1+2-3+4}$ |
|---|---|---|---|---|
| $1 \to 3$ | 0.087±0.052<br>4.362 : + | 0.088±0.052<br>4.424 : + | 0.004±0.082<br>0.126 : 0 | −0.003±0.017<br>−0.413 : 0 |
| $3 \to 1$ | 0.281±0.048<br>14.58 : + | 0.263±0.048<br>13.75 : + | −0.033±0.077<br>−1.125 : 0 | 0.002±0.017<br>0.384 : 0 |
| $123 \to 2$ | 0.115±0.054<br>5.520 : + | 0.132±0.054<br>6.314 : + | 0.031±0.077<br>1.063 : 0 | −0.002±0.017<br>−0.282 : 0 |
| $2 \to 3$ | −0.099±0.052<br>−4.901 : − | −0.098±0.052<br>−4.870 : − | 0.001±0.082<br>0.032 : 0 | 0.005±0.017<br>0.758 : 0 |
| $3 \to 2$ | −0.270±0.048<br>−14.21 : − | −0.265±0.048<br>−13.83 : − | 0.028±0.081<br>0.878 : 0 | −0.002±0.017<br>−0.238 : 0 |
| $123 \to 1$ | −0.100±0.053<br>−4.849 : − | −0.090±0.053<br>−4.388 : − | 0.017±0.074<br>0.600 : 0 | −0.000±0.017<br>−0.006 : 0 |
| $1 \to 4$ | −0.067±0.052<br>−3.317 : − | 0.025±0.056<br>1.171 : 0 | 0.148±0.067<br>5.691 : + | 0.001±0.017<br>0.177 : 0 |
| $4 \to 1$ | −0.481±0.038<br>−28.91 : − | −0.181±0.051<br>−9.098 : − | 0.572±0.045<br>27.04 : + | 0.008±0.017<br>1.336 : 0 |
| $2 \to 4$ | 0.099±0.052<br>4.925 : + | 0.013±0.057<br>0.612 : 0 | −0.153±0.068<br>−5.765 : − | 0.006±0.017<br>0.840 : 0 |
| $4 \to 2$ | 0.457±0.038<br>27.87 : + | 0.184±0.050<br>9.487 : + | −0.543±0.046<br>−25.64 : − | −0.005±0.017<br>−0.763 : 0 |
| $1 \to 5$ | 0.013±0.056<br>0.622 : 0 | −0.093±0.051<br>−4.723 : − | −0.174±0.065<br>−6.835 : − | −0.020±0.017<br>−3.049 : − |
| $5 \to 1$ | −0.205±0.050<br>−10.34 : − | −0.594±0.031<br>−40.43 : − | −0.728±0.032<br>−40.27 : − | −0.131±0.016<br>−21.14 : − |
| $1234 \to 1$ | −0.175±0.050<br>−8.899 : − | −0.234±0.048<br>−12.45 : − | −0.146±0.068<br>−5.525 : − | −0.001±0.017<br>−0.125 : 0 |
| $12345 \to 1$ | −0.208±0.050<br>−10.50 : − | −0.324±0.045<br>−17.80 : − | −0.276±0.062<br>−11.11 : − | 0.028±0.016<br>4.487 : + |
| $2 \to 5$ | 0.015±0.056<br>0.692 : 0 | 0.123±0.051<br>6.180 : + | 0.187±0.066<br>7.207 : + | 0.016±0.017<br>2.523 : 0 |
| $5 \to 2$ | 0.174±0.050<br>8.908 : + | 0.583±0.031<br>39.62 : + | 0.705±0.033<br>39.71 : + | 0.131±0.016<br>21.06 : + |
| $1234 \to 2$ | 0.209±0.050<br>10.58 : + | 0.268±0.049<br>13.80 : + | 0.120±0.068<br>4.588 : + | 0.000±0.017<br>0.006 : 0 |
| $12345 \to 2$ | 0.186±0.051<br>9.426 : + | 0.330±0.045<br>17.93 : + | 0.318±0.062<br>12.58 : + | −0.018±0.016<br>−2.899 : − |

Table S7: Simulation results for the asymmetric tree $A$ using 10,000 independent patterns. For each statistic under each scenario we report the value, the 99% Wald confidence interval with radius $2.576 \times \sqrt{4n(L)n(R)/(n(L)+n(R))^3}$, the $Z$-score and the classification. Misclassifications are highlighted in gray. Compared to Table 3 in the main text, only five scenarios were correctly identified. Compared to Table S2 in Appendix 3, also $12345 \to 1$ is correct.

| | ${}_A\Delta^*_{1-2}$ | ${}_A\Delta^*_{1-3}$ | ${}_A\Delta^*_{2-3}$ | ${}_A\Delta^*_{1+2-3+4}$ |
|---|---|---|---|---|
| $1 \to 3$ | 0.024±0.164<br>0.381 : 0 | 0.064±0.168<br>0.978 : 0 | 0.077±0.238<br>0.832 : 0 | 0.011±0.054<br>0.517 : 0 |
| $3 \to 1$ | 0.157±0.148<br>2.718 : + | 0.169±0.151<br>2.848 : + | 0.008±0.237<br>0.092 : 0 | −0.013±0.053<br>−0.630 : 0 |
| $123 \to 2$ | 0.049±0.158<br>0.796 : 0 | 0.152±0.162<br>2.403 : 0 | 0.194±0.223<br>2.201 : 0 | 0.018±0.053<br>0.890 : 0 |
| $2 \to 3$ | −0.145±0.165<br>−2.255 : 0 | −0.195±0.167<br>−2.961 : − | −0.098±0.254<br>−0.990 : 0 | 0.005±0.054<br>0.230 : 0 |
| $3 \to 2$ | −0.306±0.149<br>−5.042 : − | −0.298±0.152<br>−4.853 : − | 0.038±0.251<br>0.389 : 0 | 0.005±0.053<br>0.223 : 0 |
| $123 \to 1$ | −0.135±0.157<br>−2.207 : 0 | −0.122±0.161<br>−1.941 : 0 | 0.038±0.225<br>0.437 : 0 | −0.005±0.053<br>−0.264 : 0 |
| $1 \to 4$ | −0.094±0.164<br>−1.469 : 0 | −0.048±0.171<br>−0.727 : 0 | 0.080±0.210<br>0.980 : 0 | −0.017±0.054<br>−0.833 : 0 |
| $4 \to 1$ | −0.452±0.116<br>−8.968 : − | −0.177±0.156<br>−2.887 : − | 0.544±0.140<br>8.438 : + | −0.008±0.052<br>−0.401 : 0 |
| $2 \to 4$ | 0.188±0.168<br>2.842 : + | −0.021±0.185<br>−0.287 : 0 | −0.333±0.205<br>−3.958 : − | −0.020±0.054<br>−0.978 : 0 |
| $4 \to 2$ | 0.396±0.124<br>7.548 : + | 0.121±0.155<br>1.997 : 0 | −0.481±0.149<br>−7.303 : − | −0.032±0.052<br>−1.629 : 0 |
| $1 \to 5$ | −0.046±0.184<br>−0.641 : 0 | −0.119±0.167<br>−1.823 : 0 | −0.135±0.215<br>−1.600 : 0 | 0.003±0.054<br>0.146 : 0 |
| $5 \to 1$ | −0.111±0.151<br>−1.886 : 0 | −0.563±0.095<br>−12.65 : − | −0.783±0.090<br>−14.04 : − | −0.140±0.050<br>−7.171 : − |
| $1234 \to 1$ | −0.169±0.156<br>−2.754 : − | −0.277±0.151<br>−4.556 : − | −0.214±0.213<br>−2.535 : 0 | −0.029±0.051<br>−1.463 : 0 |
| $12345 \to 1$ | −0.242±0.154<br>−3.939 : − | −0.366±0.134<br>−6.576 : − | −0.338±0.192<br>−4.269 : − | 0.042±0.050<br>2.201 : 0 |
| $2 \to 5$ | −0.005±0.182<br>−0.071 : 0 | 0.111±0.165<br>1.732 : 0 | 0.177±0.202<br>2.228 : 0 | 0.017±0.052<br>0.848 : 0 |
| $5 \to 2$ | 0.178±0.164<br>2.770 : + | 0.575±0.099<br>12.34 : + | 0.698±0.104<br>12.45 : + | 0.143±0.050<br>7.336 : + |
| $1234 \to 2$ | 0.167±0.160<br>2.646 : + | 0.236±0.150<br>3.944 : + | 0.167±0.212<br>2.000 : 0 | −0.035±0.051<br>−1.768 : 0 |
| $12345 \to 2$ | 0.147±0.161<br>2.335 : 0 | 0.299±0.144<br>5.132 : + | 0.313±0.192<br>3.995 : + | −0.086±0.050<br>−4.441 : − |

Table S8: Simulation results for the quasisymmetric tree $Q$ using 100,000 independent patterns. For each statistic under each scenario we report the value, the 99% Wald confidence interval with radius $2.576 \times \sqrt{4n(L)n(R)/(n(L)+n(R))^3}$, the $Z$-score and the classification. Misclassifications are highlighted in gray. Compared to Table 4 in the main text, six scenarios were misclassified. Compared to Table S3 in Appendix 3, everything was correct.

| | $_Q\Delta^*_{1-6}$ | $_Q\Delta^*_{2-6}$ | $_Q\Delta^*_{3-5}$ | $_Q\Delta^*_{4-5}$ | $_Q\Delta^*_{5+7}$ | $_Q\Delta^*_{6+8}$ | $_Q\Delta^*_{3+4-5+7}$ | $_Q\Delta^*_{1+2-6+8}$ |
|---|---|---|---|---|---|---|---|---|
| $1 \to 4$ | 0.522±0.062<br>18.69 : + | 0.400±0.070<br>13.67 : + | 0.087±0.052<br>4.323 : + | 0.006±0.056<br>0.260 : 0 | −0.024±0.018<br>−3.621 : − | −0.074±0.013<br>−15.31 : − | −0.014±0.017<br>−2.077 : 0 | −0.047±0.013<br>−9.767 : − |
| $4 \to 1$ | 0.725±0.044<br>29.76 : + | 0.060±0.117<br>1.317 : 0 | 0.581±0.031<br>39.47 : + | 0.433±0.040<br>25.40 : + | −0.083±0.017<br>−13.03 : − | −0.028±0.013<br>−5.754 : − | 0.081±0.016<br>13.15 : + | 0.002±0.013<br>0.428 : 0 |
| $1 \to 5$ | −0.587±0.057<br>−21.64 : − | −0.464±0.067<br>−15.90 : − | −0.011±0.057<br>−0.484 : 0 | 0.095±0.052<br>4.683 : + | −0.022±0.018<br>−3.245 : − | 0.073±0.013<br>15.19 : + | −0.012±0.017<br>−1.821 : 0 | 0.041±0.013<br>8.664 : + |
| $5 \to 1$ | −0.695±0.045<br>−29.04 : − | −0.023±0.119<br>−0.508 : 0 | 0.434±0.041<br>25.08 : + | 0.585±0.032<br>39.39 : + | −0.090±0.017<br>−14.03 : − | 0.026±0.013<br>5.297 : + | 0.074±0.016<br>11.91 : + | −0.004±0.013<br>−0.737 : 0 |
| $2 \to 4$ | 0.453±0.071<br>14.84 : + | 0.592±0.058<br>21.50 : + | −0.100±0.052<br>−5.019 : − | 0.021±0.056<br>0.957 : 0 | 0.007±0.018<br>1.045 : 0 | −0.076±0.013<br>−15.94 : − | −0.002±0.017<br>−0.336 : 0 | −0.047±0.013<br>−9.773 : − |
| $4 \to 2$ | 0.070±0.117<br>1.542 : 0 | 0.694±0.045<br>28.68 : + | −0.584±0.031<br>−39.59 : − | −0.449±0.040<br>−26.24 : − | 0.095±0.017<br>14.81 : + | −0.036±0.013<br>−7.281 : − | −0.073±0.016<br>−11.76 : − | −0.006±0.013<br>−1.221 : 0 |
| $2 \to 5$ | −0.452±0.067<br>−15.61 : − | −0.564±0.058<br>−20.79 : − | −0.013±0.057<br>−0.611 : 0 | −0.104±0.052<br>−5.144 : − | 0.012±0.018<br>1.731 : 0 | 0.077±0.013<br>16.07 : + | −0.001±0.017<br>−0.158 : 0 | 0.046±0.013<br>9.745 : + |
| $5 \to 2$ | 0.040±0.118<br>0.868 : 0 | −0.710±0.045<br>−29.13 : − | −0.428±0.040<br>−25.12 : − | −0.585±0.031<br>−39.66 : − | 0.082±0.017<br>12.86 : + | 0.028±0.013<br>5.621 : + | −0.083±0.016<br>−13.41 : − | −0.001±0.013<br>−0.169 : 0 |
| $2 \to 45$ | −0.011±0.080<br>−0.370 : 0 | −0.008±0.071<br>−0.303 : 0 | −0.113±0.052<br>−5.624 : − | −0.112±0.052<br>−5.580 : − | 0.003±0.015<br>0.562 : 0 | 0.003±0.017<br>0.432 : 0 | −0.014±0.015<br>−2.508 : 0 | 0.002±0.016<br>0.283 : 0 |
| $45 \to 2$ | −0.052±0.164<br>−0.824 : 0 | 0.043±0.112<br>0.993 : 0 | −0.471±0.038<br>−28.67 : − | −0.464±0.038<br>−28.17 : − | 0.073±0.014<br>13.53 : + | 0.006±0.018<br>0.919 : 0 | −0.027±0.014<br>−5.108 : − | 0.006±0.018<br>0.977 : 0 |
| $1 \to 3$ | 0.012±0.118<br>0.273 : 0 | 0.019±0.118<br>0.412 : 0 | −0.120±0.052<br>−5.933 : − | −0.119±0.052<br>−5.866 : − | 0.004±0.018<br>0.667 : 0 | −0.006±0.013<br>−1.303 : 0 | −0.021±0.017<br>−3.168 : − | −0.006±0.013<br>−1.227 : 0 |
| $3 \to 1$ | 0.014±0.123<br>0.286 : 0 | −0.029±0.118<br>−0.639 : 0 | −0.286±0.047<br>−15.03 : − | −0.294±0.047<br>−15.42 : − | 0.012±0.017<br>1.827 : 0 | −0.004±0.013<br>−0.841 : 0 | −0.055±0.017<br>−8.498 : − | −0.004±0.013<br>−0.878 : 0 |
| $123 \to 2$ | −0.017±0.121<br>−0.373 : 0 | 0.026±0.116<br>0.583 : 0 | −0.216±0.051<br>−10.82 : − | −0.206±0.051<br>−10.35 : − | 0.002±0.017<br>0.308 : 0 | −0.010±0.013<br>−2.031 : 0 | −0.041±0.017<br>−6.462 : − | −0.010±0.013<br>−1.999 : 0 |
| $12345 \to 2$ | 0.010±0.117<br>0.225 : 0 | −0.015±0.102<br>−0.393 : 0 | −0.313±0.046<br>−17.02 : − | −0.320±0.046<br>−17.33 : − | −0.016±0.017<br>−2.618 : − | −0.001±0.013<br>−0.236 : 0 | −0.088±0.016<br>−14.16 : − | −0.001±0.013<br>−0.260 : 0 |
| $1 \to 45$ | −0.010±0.072<br>−0.361 : 0 | −0.017±0.080<br>−0.557 : 0 | 0.124±0.051<br>6.234 : + | 0.121±0.051<br>6.095 : + | −0.013±0.015<br>−2.373 : 0 | −0.004±0.017<br>−0.636 : 0 | 0.006±0.015<br>1.065 : 0 | −0.005±0.016<br>−0.818 : 0 |
| $45 \to 1$ | 0.030±0.116<br>0.671 : 0 | 0.039±0.180<br>0.557 : 0 | 0.466±0.038<br>28.40 : + | 0.464±0.038<br>28.29 : + | −0.076±0.014<br>−14.00 : − | 0.008±0.018<br>1.176 : 0 | 0.025±0.014<br>4.626 : + | 0.009±0.018<br>1.318 : 0 |
| $2 \to 3$ | 0.013±0.122<br>0.283 : 0 | 0.018±0.122<br>0.376 : 0 | 0.124±0.053<br>6.024 : + | 0.124±0.053<br>6.047 : + | −0.006±0.018<br>−0.855 : 0 | −0.001±0.013<br>−0.179 : 0 | 0.020±0.017<br>3.014 : + | −0.001±0.013<br>−0.111 : 0 |
| $3 \to 2$ | −0.013±0.122<br>−0.283 : 0 | −0.045±0.129<br>−0.900 : 0 | 0.273±0.048<br>14.36 : + | 0.268±0.048<br>14.13 : + | 0.004±0.017<br>0.572 : 0 | 0.003±0.013<br>0.709 : 0 | 0.066±0.017<br>10.27 : + | 0.003±0.013<br>0.590 : 0 |
| $123 \to 1$ | 0.027±0.113<br>0.614 : 0 | 0.042±0.119<br>0.919 : 0 | 0.198±0.050<br>10.02 : + | 0.201±0.050<br>10.15 : + | −0.008±0.017<br>−1.296 : 0 | −0.005±0.013<br>−1.050 : 0 | 0.033±0.017<br>5.225 : + | −0.004±0.013<br>−0.879 : 0 |
| $12345 \to 1$ | −0.033±0.103<br>−0.833 : 0 | −0.036±0.115<br>−0.800 : 0 | 0.349±0.046<br>18.65 : + | 0.349±0.046<br>18.68 : + | 0.027±0.017<br>4.305 : + | −0.005±0.013<br>−1.096 : 0 | 0.103±0.016<br>16.65 : + | −0.006±0.013<br>−1.282 : 0 |
| $5 \to 12$ | −0.543±0.061<br>−19.34 : − | −0.582±0.059<br>−20.71 : − | 0.016±0.043<br>0.962 : 0 | 0.002±0.039<br>0.134 : 0 | −0.004±0.017<br>−0.632 : 0 | 0.028±0.013<br>5.701 : + | −0.001±0.016<br>−0.203 : 0 | −0.006±0.013<br>−1.318 : 0 |
| $12 \to 5$ | −0.257±0.081<br>−7.956 : − | −0.232±0.080<br>−7.357 : − | −0.015±0.057<br>−0.678 : 0 | −0.008±0.056<br>−0.385 : 0 | 0.000±0.018<br>0.066 : 0 | 0.054±0.013<br>11.23 : + | −0.002±0.017<br>−0.256 : 0 | 0.042±0.013<br>8.885 : + |
| $4 \to 3$ | −0.706±0.044<br>−29.77 : − | −0.716±0.043<br>−30.11 : − | 0.012±0.060<br>0.531 : 0 | 0.007±0.060<br>0.299 : 0 | 0.008±0.018<br>1.168 : 0 | 0.012±0.013<br>2.509 : 0 | 0.009±0.018<br>1.399 : 0 | −0.047±0.013<br>−9.681 : − |
| $3 \to 4$ | −0.598±0.058<br>−21.39 : − | −0.612±0.057<br>−21.95 : − | 0.032±0.059<br>1.387 : 0 | 0.019±0.057<br>0.877 : 0 | −0.002±0.018<br>−0.252 : 0 | −0.045±0.013<br>−9.412 : − | 0.003±0.017<br>0.414 : 0 | −0.080±0.013<br>−16.78 : − |
| $4 \to 12$ | 0.562±0.060<br>20.04 : + | 0.572±0.060<br>20.32 : + | −0.022±0.039<br>−1.463 : 0 | −0.025±0.044<br>−1.505 : 0 | −0.003±0.017<br>−0.498 : 0 | −0.029±0.013<br>−5.878 : − | −0.010±0.017<br>−1.642 : 0 | 0.006±0.013<br>1.167 : 0 |
| $12 \to 4$ | 0.241±0.079<br>7.663 : + | 0.289±0.078<br>9.161 : + | −0.020±0.056<br>−0.921 : 0 | 0.001±0.057<br>0.065 : 0 | 0.003±0.018<br>0.519 : 0 | −0.062±0.013<br>−13.00 : − | 0.002±0.017<br>0.250 : 0 | −0.050±0.013<br>−10.37 : − |
| $5 \to 3$ | 0.691±0.045<br>28.97 : + | 0.691±0.045<br>29.02 : + | 0.003±0.061<br>0.140 : 0 | 0.005±0.061<br>0.210 : 0 | −0.012±0.018<br>−1.842 : 0 | −0.005±0.013<br>−1.045 : 0 | −0.011±0.017<br>−1.733 : 0 | 0.052±0.013<br>10.70 : + |
| $3 \to 5$ | 0.553±0.059<br>20.30 : + | 0.573±0.059<br>20.74 : + | −0.033±0.057<br>−1.507 : 0 | −0.033±0.059<br>−1.472 : 0 | 0.002±0.018<br>0.344 : 0 | 0.044±0.013<br>9.267 : + | −0.004±0.017<br>−0.538 : 0 | 0.078±0.013<br>16.34 : + |
| $123 \to 5$ | 0.017±0.095<br>0.477 : 0 | 0.024±0.095<br>0.661 : 0 | 0.022±0.057<br>1.013 : 0 | 0.024±0.057<br>1.115 : 0 | −0.001±0.018<br>−0.220 : 0 | 0.032±0.013<br>6.772 : + | 0.003±0.018<br>0.424 : 0 | 0.033±0.013<br>6.894 : + |
| $5 \to 123$ | −0.021±0.077<br>−0.712 : 0 | −0.001±0.078<br>−0.030 : 0 | 0.017±0.053<br>0.851 : 0 | 0.026±0.052<br>1.301 : 0 | 0.001±0.017<br>0.216 : 0 | 0.012±0.013<br>2.376 : 0 | 0.006±0.017<br>0.907 : 0 | 0.011±0.013<br>2.240 : 0 |
| $12345 \to 4$ | −0.064±0.111<br>−1.502 : 0 | −0.024±0.111<br>−0.559 : 0 | −0.020±0.057<br>−0.899 : 0 | −0.009±0.057<br>−0.419 : 0 | −0.007±0.018<br>−1.101 : 0 | 0.016±0.013<br>3.295 : + | −0.010±0.018<br>−1.492 : 0 | 0.015±0.013<br>3.056 : + |
| $123 \to 4$ | −0.018±0.097<br>−0.485 : 0 | −0.008±0.097<br>−0.226 : 0 | −0.001±0.056<br>−0.065 : 0 | 0.002±0.056<br>0.087 : 0 | 0.000±0.018<br>0.027 : 0 | −0.028±0.013<br>−5.750 : − | 0.000±0.017<br>0.033 : 0 | −0.028±0.013<br>−5.818 : − |
| $4 \to 123$ | −0.017±0.080<br>−0.553 : 0 | −0.027±0.079<br>−0.881 : 0 | 0.019±0.052<br>0.927 : 0 | 0.014±0.053<br>0.707 : 0 | −0.001±0.017<br>−0.099 : 0 | −0.012±0.013<br>−2.400 : 0 | 0.003±0.017<br>0.429 : 0 | −0.013±0.013<br>−2.612 : − |
| $12345 \to 5$ | −0.039±0.114<br>−0.880 : 0 | 0.016±0.115<br>0.354 : 0 | −0.015±0.057<br>−0.684 : 0 | −0.001±0.057<br>−0.066 : 0 | 0.016±0.018<br>2.367 : 0 | −0.015±0.013<br>−3.027 : − | 0.014±0.018<br>2.129 : 0 | −0.015±0.013<br>−3.076 : − |

Table S9: Simulation results for the quasisymmetric tree $Q$ using 10,000 independent patterns. For each statistic under each scenario we report the value, the 99% Wald confidence interval with radius $2.576 \times \sqrt{4n(L)n(R)/(n(L)+n(R))^3}$, the $Z$-score and the classification. Misclassifications are highlighted in gray. Compared to Table 4 in the main text, only six admixture scenarios were correctly identified. Compared to Table S3 in Appendix 3, twenty-three of the admixture scenarios were correctly identified, including the six scenarios that have a zero signature.

| | $_Q\Delta^*_{1-6}$ | $_Q\Delta^*_{2-6}$ | $_Q\Delta^*_{3-5}$ | $_Q\Delta^*_{4-5}$ | $_Q\Delta^*_{5+7}$ | $_Q\Delta^*_{6+8}$ | $_Q\Delta^*_{3+4-5+7}$ | $_Q\Delta^*_{1+2-6+8}$ |
|---|---|---|---|---|---|---|---|---|
| 1 → 4 | 0.417±0.208<br>4.703 : + | 0.315±0.236<br>3.272 : + | 0.041±0.165<br>0.640 : 0 | –0.041±0.174<br>–0.605 : 0 | –0.000±0.055<br>–0.000 : 0 | –0.072±0.040<br>–4.707 : – | 0.000±0.054<br>0.021 : 0 | –0.051±0.039<br>–3.348 : – |
| 4 → 1 | 0.624±0.155<br>8.130 : + | –0.228±0.333<br>–1.722 : 0 | 0.603±0.100<br>12.40 : + | 0.444±0.132<br>7.775 : + | –0.109±0.053<br>–5.350 : – | –0.005±0.041<br>–0.344 : 0 | 0.050±0.051<br>2.526 : 0 | 0.017±0.040<br>1.090 : 0 |
| 1 → 5 | –0.460±0.204<br>–5.167 : – | –0.410±0.230<br>–4.196 : – | –0.004±0.169<br>–0.065 : 0 | 0.056±0.163<br>0.882 : 0 | –0.018±0.055<br>–0.861 : 0 | 0.075±0.039<br>4.939 : + | –0.012±0.054<br>–0.580 : 0 | 0.051±0.039<br>3.389 : + |
| 5 → 1 | –0.718±0.138<br>–9.357 : – | –0.000±0.389<br>–0.000 : 0 | 0.465±0.123<br>8.627 : + | 0.613±0.095<br>13.15 : + | –0.134±0.053<br>–6.586 : – | 0.016±0.041<br>1.005 : 0 | 0.047±0.052<br>2.359 : 0 | –0.014±0.040<br>–0.896 : 0 |
| 2 → 4 | 0.345±0.225<br>3.714 : + | 0.576±0.184<br>6.615 : + | –0.079±0.166<br>–1.224 : 0 | 0.081±0.178<br>1.176 : 0 | –0.010±0.055<br>–0.488 : 0 | –0.070±0.039<br>–4.608 : – | –0.011±0.054<br>–0.522 : 0 | –0.043±0.039<br>–2.828 : – |
| 4 → 2 | –0.088±0.340<br>–0.662 : 0 | 0.743±0.129<br>9.941 : + | –0.611±0.095<br>–13.15 : – | –0.425±0.127<br>–7.852 : – | 0.105±0.053<br>5.125 : + | –0.032±0.041<br>–2.068 : 0 | –0.070±0.052<br>–3.533 : – | –0.001±0.040<br>–0.076 : 0 |
| 2 → 5 | –0.487±0.208<br>–5.270 : – | –0.517±0.181<br>–6.308 : – | –0.009±0.175<br>–0.135 : 0 | –0.087±0.162<br>–1.380 : 0 | –0.008±0.055<br>–0.377 : 0 | 0.065±0.039<br>4.319 : + | –0.018±0.054<br>–0.866 : 0 | 0.034±0.039<br>2.275 : 0 |
| 5 → 2 | 0.111±0.427<br>0.667 : 0 | –0.720±0.140<br>–9.214 : – | –0.357±0.136<br>–6.321 : – | –0.547±0.105<br>–11.31 : – | 0.051±0.053<br>2.507 : 0 | 0.006±0.041<br>0.362 : 0 | –0.086±0.051<br>–4.362 : – | –0.022±0.040<br>–1.408 : 0 |
| 2 → 45 | –0.119±0.245<br>–1.245 : 0 | –0.068±0.223<br>–0.780 : 0 | –0.093±0.167<br>–1.432 : 0 | –0.075±0.166<br>–1.162 : 0 | –0.023±0.046<br>–1.325 : 0 | 0.020±0.051<br>1.003 : 0 | –0.035±0.045<br>–2.006 : 0 | 0.011±0.050<br>0.562 : 0 |
| 45 → 2 | 0.474±0.521<br>2.065 : 0 | 0.115±0.355<br>0.832 : 0 | –0.468±0.120<br>–8.951 : – | –0.478±0.119<br>–9.120 : – | 0.085±0.044<br>5.006 : + | 0.026±0.054<br>1.233 : 0 | –0.014±0.044<br>–0.844 : 0 | 0.032±0.054<br>1.534 : 0 |
| 1 → 3 | 0.277±0.362<br>1.896 : 0 | 0.208±0.364<br>1.443 : 0 | –0.171±0.162<br>–2.678 : – | –0.187±0.164<br>–2.899 : – | 0.039±0.054<br>1.867 : 0 | 0.000±0.040<br>0.015 : 0 | 0.001±0.054<br>0.042 : 0 | 0.006±0.040<br>0.366 : 0 |
| 3 → 1 | –0.190±0.391<br>–1.234 : 0 | –0.211±0.409<br>–1.298 : 0 | –0.220±0.150<br>–3.692 : – | –0.214±0.148<br>–3.641 : – | –0.021±0.053<br>–1.043 : 0 | –0.003±0.040<br>–0.169 : 0 | –0.072±0.053<br>–3.554 : – | –0.006±0.040<br>–0.413 : 0 |
| 123 → 2 | 0.023±0.393<br>0.152 : 0 | –0.064±0.375<br>–0.438 : 0 | –0.133±0.165<br>–2.066 : 0 | –0.146±0.163<br>–2.295 : 0 | 0.002±0.052<br>0.100 : 0 | –0.006±0.041<br>–0.376 : 0 | –0.025±0.052<br>–1.251 : 0 | –0.006±0.041<br>–0.406 : 0 |
| 12345 → 2 | –0.055±0.347<br>–0.405 : 0 | –0.143±0.322<br>–1.134 : 0 | –0.359±0.144<br>–6.025 : – | –0.392±0.144<br>–6.476 : – | 0.010±0.052<br>0.482 : 0 | 0.014±0.041<br>0.922 : 0 | –0.073±0.052<br>–3.665 : – | 0.011±0.041<br>0.731 : 0 |
| 1 → 45 | 0.006±0.205<br>0.079 : 0 | –0.022±0.221<br>–0.256 : 0 | 0.114±0.155<br>1.876 : 0 | 0.095±0.153<br>1.605 : 0 | 0.012±0.046<br>0.666 : 0 | 0.018±0.051<br>0.891 : 0 | 0.029±0.045<br>1.662 : 0 | 0.016±0.051<br>0.838 : 0 |
| 45 → 1 | –0.037±0.351<br>–0.272 : 0 | 0.391±0.495<br>1.877 : 0 | 0.445±0.123<br>8.415 : + | 0.467±0.120<br>8.910 : + | –0.077±0.045<br>–4.468 : – | 0.005±0.054<br>0.229 : 0 | 0.020±0.045<br>1.179 : 0 | 0.008±0.054<br>0.371 : 0 |
| 2 → 3 | –0.106±0.374<br>–0.729 : 0 | –0.080±0.364<br>–0.566 : 0 | 0.114±0.158<br>1.846 : 0 | 0.118±0.158<br>1.912 : 0 | –0.001±0.054<br>–0.062 : 0 | 0.011±0.040<br>0.738 : 0 | 0.025±0.054<br>1.194 : 0 | 0.009±0.040<br>0.597 : 0 |
| 3 → 2 | 0.143±0.365<br>1.000 : 0 | 0.115±0.355<br>0.832 : 0 | 0.303±0.145<br>5.135 : + | 0.301±0.146<br>5.085 : + | 0.024±0.052<br>1.188 : 0 | 0.002±0.041<br>0.109 : 0 | 0.093±0.052<br>4.653 : + | 0.005±0.040<br>0.310 : 0 |
| 123 → 1 | –0.016±0.330<br>–0.128 : 0 | –0.016±0.330<br>–0.128 : 0 | 0.162±0.167<br>2.484 : 0 | 0.168±0.169<br>2.528 : 0 | 0.014±0.053<br>0.688 : 0 | 0.009±0.040<br>0.596 : 0 | 0.045±0.052<br>2.216 : 0 | 0.009±0.040<br>0.564 : 0 |
| 12345 → 1 | –0.057±0.354<br>–0.412 : 0 | –0.067±0.384<br>–0.447 : 0 | 0.428±0.141<br>7.103 : + | 0.418±0.140<br>7.027 : + | –0.009±0.051<br>–0.475 : 0 | –0.004±0.041<br>–0.264 : 0 | 0.082±0.051<br>4.164 : + | –0.006±0.040<br>–0.356 : 0 |
| 5 → 12 | –0.431±0.210<br>–4.779 : – | –0.603±0.184<br>–6.771 : – | 0.030±0.135<br>0.573 : 0 | –0.027±0.122<br>–0.568 : 0 | –0.036±0.053<br>–1.779 : 0 | 0.023±0.041<br>1.450 : 0 | –0.035±0.051<br>–1.762 : 0 | –0.009±0.040<br>–0.554 : 0 |
| 12 → 5 | –0.216±0.256<br>–2.132 : 0 | –0.124±0.250<br>–1.269 : 0 | –0.030±0.183<br>–0.426 : 0 | 0.010±0.178<br>0.138 : 0 | 0.044±0.055<br>2.086 : 0 | 0.048±0.039<br>3.137 : + | 0.041±0.054<br>1.977 : 0 | 0.039±0.039<br>2.605 : + |
| 4 → 3 | –0.665±0.149<br>–8.589 : – | –0.637±0.152<br>–8.335 : – | –0.054±0.173<br>–0.805 : 0 | –0.045±0.173<br>–0.671 : 0 | 0.004±0.054<br>0.207 : 0 | –0.021±0.040<br>–1.350 : 0 | –0.005±0.053<br>–0.247 : 0 | –0.073±0.040<br>–4.748 : – |
| 3 → 4 | –0.543±0.192<br>–6.123 : – | –0.583±0.196<br>–6.248 : – | 0.021±0.186<br>0.289 : 0 | 0.031±0.184<br>0.429 : 0 | 0.010±0.055<br>0.462 : 0 | –0.044±0.039<br>–2.901 : – | 0.014±0.054<br>0.666 : 0 | –0.074±0.039<br>–4.932 : – |
| 4 → 12 | 0.603±0.191<br>6.499 : + | 0.469±0.202<br>5.303 : + | 0.004±0.119<br>0.092 : 0 | –0.021±0.131<br>–0.405 : 0 | –0.041±0.052<br>–2.024 : 0 | –0.037±0.041<br>–2.374 : 0 | –0.041±0.051<br>–2.094 : 0 | –0.005±0.040<br>–0.339 : 0 |
| 12 → 4 | 0.186±0.273<br>1.725 : 0 | 0.290±0.256<br>2.800 : + | –0.099±0.173<br>–1.477 : 0 | –0.053±0.178<br>–0.761 : 0 | 0.036±0.055<br>1.699 : 0 | –0.070±0.039<br>–4.641 : – | 0.021±0.055<br>0.988 : 0 | –0.060±0.039<br>–3.971 : – |
| 5 → 3 | 0.709±0.136<br>9.492 : + | 0.658±0.142<br>8.995 : + | 0.172±0.196<br>2.231 : 0 | 0.145±0.194<br>1.901 : 0 | –0.027±0.054<br>–1.291 : 0 | –0.011±0.040<br>–0.710 : 0 | –0.003±0.054<br>–0.166 : 0 | 0.048±0.040<br>3.142 : + |
| 3 → 5 | 0.471±0.195<br>5.488 : + | 0.447±0.195<br>5.306 : + | 0.034±0.179<br>0.487 : 0 | 0.031±0.186<br>0.433 : 0 | –0.017±0.055<br>–0.798 : 0 | 0.057±0.039<br>3.765 : + | –0.011±0.054<br>–0.520 : 0 | 0.085±0.039<br>5.650 : + |
| 123 → 5 | –0.014±0.302<br>–0.117 : 0 | –0.061±0.317<br>–0.492 : 0 | 0.038±0.177<br>0.549 : 0 | 0.024±0.179<br>0.348 : 0 | –0.003±0.055<br>–0.148 : 0 | 0.017±0.040<br>1.092 : 0 | 0.003±0.055<br>0.126 : 0 | 0.015±0.039<br>1.012 : 0 |
| 5 → 123 | 0.030±0.257<br>0.299 : 0 | –0.067±0.251<br>–0.683 : 0 | 0.102±0.158<br>1.659 : 0 | 0.061±0.154<br>1.018 : 0 | 0.017±0.054<br>0.830 : 0 | 0.003±0.040<br>0.184 : 0 | 0.035±0.053<br>1.722 : 0 | 0.002±0.040<br>0.122 : 0 |
| 12345 → 4 | 0.280±0.350<br>1.980 : 0 | –0.022±0.384<br>–0.149 : 0 | 0.051±0.184<br>0.714 : 0 | –0.027±0.189<br>–0.366 : 0 | 0.012±0.055<br>0.570 : 0 | 0.017±0.040<br>1.136 : 0 | 0.014±0.055<br>0.672 : 0 | 0.020±0.039<br>1.329 : 0 |
| 123 → 4 | –0.000±0.300<br>–0.000 : 0 | –0.000±0.304<br>–0.000 : 0 | 0.166±0.193<br>2.192 : 0 | 0.164±0.191<br>2.180 : 0 | –0.035±0.054<br>–1.691 : 0 | –0.021±0.040<br>–1.403 : 0 | –0.010±0.054<br>–0.477 : 0 | –0.021±0.040<br>–1.397 : 0 |
| 4 → 123 | 0.121±0.248<br>1.257 : 0 | 0.171±0.241<br>1.803 : 0 | –0.000±0.165<br>–0.000 : 0 | 0.026±0.170<br>0.396 : 0 | 0.012±0.055<br>0.553 : 0 | –0.013±0.040<br>–0.859 : 0 | 0.014±0.055<br>0.673 : 0 | –0.006±0.040<br>–0.366 : 0 |
| 12345 → 5 | 0.193±0.335<br>1.457 : 0 | 0.018±0.348<br>0.135 : 0 | 0.121±0.172<br>1.808 : 0 | 0.078±0.174<br>1.149 : 0 | –0.035±0.055<br>–1.673 : 0 | –0.027±0.040<br>–1.767 : 0 | –0.016±0.055<br>–0.737 : 0 | –0.024±0.040<br>–1.580 : 0 |

### Appendix 5: Comparison to the $D_{\mathbf{FOIL}}$ method

#### Classification of expected $D_{\mathbf{FOIL}}$-statistics

In contrast to the original $D_{\text{FOIL}}$-paper (Pease & Hahn, 2015), we have considered gene flow events in tree $S$ that involve population 5, as well as gene flow from ancient populations 1234 or 12345 mediated by a ghost population. While Pease and Hahn assumed that 5 always carries the ancestral allele, consistent prediction under scenario $5 \to x$, $x \in \{1,2,3,4\}$ requires the outgroup mutation assumption (1), and the filtering would also alter the signals of $1 \to 5$, $2 \to 5$, $3 \to 5$ and $4 \to 5$ from zero signatures into $(0\,0++)$, $(0\,0--)$, $(++0\,0)$ and $(--0\,0)$, respectively. We begin this section by expanding Table 1 from Pease & Hahn, 2015 into Table S10. Note that while the table otherwise resembles Table S1 from Appendix 3, gene flow from other terminal branches into 5 produces a zero signature.

Table S10: Expanded $D_{\text{FOIL}}$-predictions. Every table element requires (2). Events $5 \to x$ and $x \to 5$, where $x \in \{1,2,3,4\}$, assume (1). Unidirectional gene flow events not listed in the table (notably $x \to 5$, $x \in \{1,2,3,4\}$) produce a zero signature.

| | $D_{\text{FO}}$ | $D_{\text{IL}}$ | $D_{\text{FI}}$ | $D_{\text{OL}}$ |
|---|---|---|---|---|
| $1 \to 3$ | + | + | + | 0 |
| $3 \to 1$ | + | 0 | + | + |
| $1 \to 4$ | − | − | 0 | + |
| $4 \to 1$ | − | 0 | + | + |
| $2 \to 3$ | + | + | − | 0 |
| $3 \to 2$ | 0 | + | − | − |
| $2 \to 4$ | − | − | 0 | − |
| $4 \to 2$ | 0 | − | − | − |
| $12345 \to 1$<br>$34 \leftrightarrow 2$<br>$1234 \to 1$<br>$5 \to 1$ | 0 | 0 | − | − |
| $12345 \to 2$<br>$34 \leftrightarrow 1$<br>$1234 \to 2$<br>$5 \to 2$ | 0 | 0 | + | + |
| $12345 \to 3$<br>$12 \leftrightarrow 4$<br>$1234 \to 3$<br>$5 \to 3$ | − | − | 0 | 0 |
| $12345 \to 4$<br>$12 \leftrightarrow 3$<br>$1234 \to 4$<br>$5 \to 4$ | + | + | 0 | 0 |

We will now quickly outline the proof of Table S10. Using the identities

$$\begin{cases} D_{\text{FO}} = {}_S\Delta^*_{1-2-6-8} \\ D_{\text{IL}} \;= {}_S\Delta^*_{2-1-6-8} \\ D_{\text{FI}} \;= {}_S\Delta^*_{3-4-5-7} \\ D_{\text{OL}} = {}_S\Delta^*_{4-3-5-7} \end{cases}$$

we can follow the proof of Table 2 in the main text, presented in Appendix 2. The figures given for the five essentially different admixture scenarios on $S$ still apply, as do equations $(S0)$ – $(S19)$. If population 5 is assumed to always carry the ancestral allele, relations depending on Lemmas 1–9 must now justified using the thirteen specializes versions of the Lemmas at the end of Appendix 2; this produces an inconsistent result in case $5 \to 4$ and $(--0\,0)$ in case $4 \to 5$. The necessary tables are written in the next page. As before, an asterisk means relying on the synchronization assumption (2).

**$3 \to 2$.**

| | ${}_S\Delta_{1-2-6-8}$ | ${}_S\Delta_{2-1-6-8}$ | ${}_S\Delta_{3-4-5-7}$ | ${}_S\Delta_{4-3-5-7}$ |
|---|---|---|---|---|
| $(S0)$ | $0^*$ | $0^*$ | $0^*$ | $0^*$ |
| $(S1)$ | $0$ | $0$ | $-^*$ | $-^*$ |
| $(S2)$ | $0$ | $+$ | $-$ | $-^*$ |
| $(S3)$ | $0^*$ | $+^*$ | $-^*$ | $0^*$ |
| $3 \to 2$ | $0^*$ | $+^*$ | $-^*$ | $-^*$ |

**$12345 \to 4$.**

| | ${}_S\Delta_{1-2-6-8}$ | ${}_S\Delta_{2-1-6-8}$ | ${}_S\Delta_{3-4-5-7}$ | ${}_S\Delta_{4-3-5-7}$ |
|---|---|---|---|---|
| $(S0)$ | $0^*$ | $0^*$ | $0^*$ | $0^*$ |
| $(S4), (S5)$ | $+^*$ | $+^*$ | $0$ | $0$ |
| $12345 \to 4$ | $+^*$ | $+^*$ | $0^*$ | $0^*$ |

**$3 \to 12$.**

| | ${}_S\Delta_{1-2-6-8}$ | ${}_S\Delta_{2-1-6-8}$ | ${}_S\Delta_{3-4-5-7}$ | ${}_S\Delta_{4-3-5-7}$ |
|---|---|---|---|---|
| $(S0)$ | $0^*$ | $0^*$ | $0^*$ | $0^*$ |
| $(S6)$ | $0$ | $0$ | $0$ | $0$ |
| $(S7)$ | $+$ | $+$ | $0$ | $0$ |
| $(S8)$ | $0^*$ | $0^*$ | $0$ | $0$ |
| $(S9), (S10),$ $(S11)$ | $+^*$ | $+^*$ | $0$ | $0$ |
| $3 \to 12$ | $+^*$ | $+^*$ | $0^*$ | $0^*$ |

**$5 \to 4$.**

| | ${}_S\Delta_{1-2-6-8}$ | ${}_S\Delta_{2-1-6-8}$ | ${}_S\Delta_{3-4-5-7}$ | ${}_S\Delta_{4-3-5-7}$ |
|---|---|---|---|---|
| $(S0), (S16)$ | $0^*$ | $0^*$ | $0^*$ | $0^*$ |
| $(S12), (S13)$ $(S14), (S15)$ | $+^*$ | $+^*$ | $0$ | $0$ |
| $5 \to 4$ | $+^*$ | $+^*$ | $0^*$ | $0^*$ |

**$4 \to 5$.**

| | ${}_S\Delta_{1-2-6-8}$ | ${}_S\Delta_{2-1-6-8}$ | ${}_S\Delta_{3-4-5-7}$ | ${}_S\Delta_{4-3-5-7}$ |
|---|---|---|---|---|
| $(S0), (S19)$ | $0^*$ | $0^*$ | $0^*$ | $0^*$ |
| $(S17), (S18)$ | $0$ | $0$ | $0$ | $0$ |
| $4 \to 5$ | $0^*$ | $0^*$ | $0^*$ | $0^*$ |

#### Simulations

We tested the $D_{\text{FOIL}}$-statistics on the same simulated data sets that were used for $\Delta$-statistics on $S$, with 1,000,000 (presented in the main text), 100,000 and 10,000 (presented in Appendix 4) independent allelic patterns. For details of the simulations, consult the main document and our GitHub page. Results are shown in Tables S11, S12 and S13.

As with $\Delta$-statistics, the ability to identify admixture scenarios correctly suffered at 100,000 independent patterns and became very weak at 10,000 patterns. The $\Delta$-statistics on tree $S$ restricted to Table S1 in Appendix 3 not only are free from using the synchronization assumption (2), but also have a notable power advantage over $D_{\text{FOIL}}$.

Table S11: Simulation results using the data set with 1,000,000 independent patterns. For each statistic under each scenario we report the value, the 99% Wald confidence interval with radius $2.576 \times \sqrt{4n(L)n(R)/(n(L)+n(R))^3}$, the $Z$-score and the classification. All results are in concordance with Table S10. Recall that the $D_{\mathrm{FOIL}}$ method is blind to events $1 \to 5$, $2 \to 5$, $3 \to 5$ and $4 \to 5$ and the signatures were supposed to be zero.

| | $D_{\mathrm{FO}}$ | $D_{\mathrm{IL}}$ | $D_{\mathrm{FI}}$ | $D_{\mathrm{OL}}$ |
|---|---|---|---|---|
| $1 \to 3$ | 0.068 ± 0.005<br>36.26 : + | 0.044 ± 0.005<br>23.41 : + | 0.023 ± 0.005<br>12.22 : + | −0.002 ± 0.005<br>−0.941 : 0 |
| $3 \to 1$ | 0.025 ± 0.005<br>13.16 : + | 0.000 ± 0.005<br>0.006 : 0 | 0.073 ± 0.005<br>38.85 : + | 0.049 ± 0.005<br>26.01 : + |
| $1 \to 4$ | −0.072 ± 0.005<br>−38.41 : − | −0.048 ± 0.005<br>−25.51 : − | −0.001 ± 0.005<br>−0.393 : 0 | 0.025 ± 0.005<br>12.81 : + |
| $4 \to 1$ | −0.025 ± 0.005<br>−12.83 : − | −0.001 ± 0.005<br>−0.514 : 0 | 0.047 ± 0.005<br>25.33 : + | 0.070 ± 0.005<br>37.31 : + |
| $2 \to 3$ | 0.049 ± 0.005<br>26.03 : + | 0.072 ± 0.005<br>38.73 : + | −0.027 ± 0.005<br>−14.08 : − | −0.002 ± 0.005<br>−1.071 : 0 |
| $3 \to 2$ | 0.002 ± 0.005<br>1.197 : 0 | 0.027 ± 0.005<br>14.11 : + | −0.069 ± 0.005<br>−37.09 : − | −0.046 ± 0.005<br>−24.52 : − |
| $2 \to 4$ | −0.047 ± 0.005<br>−24.90 : − | −0.071 ± 0.005<br>−38.15 : − | −0.000 ± 0.005<br>−0.136 : 0 | −0.026 ± 0.005<br>−13.74 : − |
| $4 \to 2$ | −0.001 ± 0.005<br>−0.674 : 0 | −0.027 ± 0.005<br>−14.12 : − | −0.049 ± 0.005<br>−26.02 : − | −0.073 ± 0.005<br>−39.12 : − |
| $12345 \to 1$ | −0.003 ± 0.005<br>−1.656 : 0 | −0.004 ± 0.005<br>−1.854 : 0 | −0.068 ± 0.005<br>−37.04 : − | −0.068 ± 0.005<br>−37.23 : − |
| $34 \to 2$ | 0.003 ± 0.006<br>1.557 : 0 | 0.004 ± 0.006<br>1.928 : 0 | −0.045 ± 0.005<br>−25.93 : − | −0.044 ± 0.005<br>−25.62 : − |
| $2 \to 34$ | 0.003 ± 0.006<br>1.395 : 0 | 0.003 ± 0.006<br>1.287 : 0 | −0.015 ± 0.005<br>−8.610 : − | −0.015 ± 0.005<br>−8.704 : − |
| $1234 \to 1$ | 0.003 ± 0.005<br>1.323 : 0 | 0.003 ± 0.005<br>1.411 : 0 | −0.023 ± 0.005<br>−12.26 : − | −0.023 ± 0.005<br>−12.17 : − |
| $5 \to 1$ | 0.002 ± 0.005<br>0.853 : 0 | 0.001 ± 0.005<br>0.339 : 0 | −0.047 ± 0.005<br>−25.74 : − | −0.048 ± 0.005<br>−26.23 : − |
| $1 \to 5$ | −0.001 ± 0.005<br>−0.639 : 0 | −0.000 ± 0.005<br>−0.184 : 0 | 0.003 ± 0.005<br>1.733 : 0 | 0.004 ± 0.005<br>2.188 : 0 |
| $12345 \to 2$ | −0.001 ± 0.005<br>−0.570 : 0 | −0.002 ± 0.005<br>−0.837 : 0 | 0.068 ± 0.005<br>37.11 : + | 0.067 ± 0.005<br>36.86 : + |
| $34 \to 1$ | −0.002 ± 0.006<br>−0.801 : 0 | −0.001 ± 0.006<br>−0.477 : 0 | 0.043 ± 0.005<br>24.96 : + | 0.043 ± 0.005<br>25.23 : + |
| $1 \to 34$ | 0.002 ± 0.006<br>0.837 : 0 | 0.001 ± 0.006<br>0.351 : 0 | 0.013 ± 0.005<br>7.633 : + | 0.013 ± 0.005<br>7.212 : + |
| $1234 \to 2$ | 0.000 ± 0.005<br>0.067 : 0 | −0.000 ± 0.005<br>−0.037 : 0 | 0.020 ± 0.005<br>10.57 : + | 0.019 ± 0.005<br>10.47 : + |
| $5 \to 2$ | −0.001 ± 0.005<br>−0.355 : 0 | −0.001 ± 0.005<br>−0.634 : 0 | 0.043 ± 0.005<br>23.36 : + | 0.042 ± 0.005<br>23.09 : + |
| $2 \to 5$ | −0.000 ± 0.005<br>−0.199 : 0 | −0.001 ± 0.005<br>−0.521 : 0 | 0.001 ± 0.005<br>0.483 : 0 | 0.000 ± 0.005<br>0.161 : 0 |
| $12345 \to 3$ | −0.070 ± 0.005<br>−38.41 : − | −0.071 ± 0.005<br>−38.67 : − | −0.002 ± 0.005<br>−1.221 : 0 | −0.003 ± 0.005<br>−1.496 : 0 |
| $12 \to 4$ | −0.043 ± 0.005<br>−25.15 : − | −0.043 ± 0.005<br>−24.96 : − | −0.004 ± 0.006<br>−1.792 : 0 | −0.003 ± 0.006<br>−1.567 : 0 |
| $4 \to 12$ | −0.013 ± 0.005<br>−7.740 : − | −0.013 ± 0.005<br>−7.681 : − | 0.000 ± 0.006<br>0.206 : 0 | 0.001 ± 0.006<br>0.275 : 0 |
| $1234 \to 3$ | −0.023 ± 0.005<br>−12.42 : − | −0.023 ± 0.005<br>−12.56 : − | 0.002 ± 0.005<br>1.064 : 0 | 0.002 ± 0.005<br>0.921 : 0 |
| $5 \to 3$ | −0.044 ± 0.005<br>−23.91 : − | −0.044 ± 0.005<br>−24.16 : − | 0.000 ± 0.005<br>0.163 : 0 | −0.000 ± 0.005<br>−0.098 : 0 |
| $3 \to 5$ | −0.001 ± 0.005<br>−0.468 : 0 | −0.002 ± 0.005<br>−0.961 : 0 | 0.002 ± 0.005<br>1.021 : 0 | 0.001 ± 0.005<br>0.527 : 0 |
| $12345 \to 4$ | 0.067 ± 0.005<br>36.46 : + | 0.067 ± 0.005<br>36.60 : + | −0.002 ± 0.005<br>−1.200 : 0 | −0.002 ± 0.005<br>−1.052 : 0 |
| $12 \to 3$ | 0.040 ± 0.005<br>23.24 : + | 0.040 ± 0.005<br>23.49 : + | −0.002 ± 0.006<br>−1.002 : 0 | −0.001 ± 0.006<br>−0.701 : 0 |
| $3 \to 12$ | 0.013 ± 0.005<br>7.316 : + | 0.013 ± 0.005<br>7.491 : + | 0.003 ± 0.006<br>1.456 : 0 | 0.003 ± 0.006<br>1.656 : 0 |
| $1234 \to 4$ | 0.024 ± 0.005<br>12.82 : + | 0.024 ± 0.005<br>13.07 : + | 0.000 ± 0.005<br>0.156 : 0 | 0.001 ± 0.005<br>0.411 : 0 |
| $5 \to 4$ | 0.041 ± 0.005<br>22.40 : + | 0.040 ± 0.005<br>21.96 : + | 0.003 ± 0.005<br>1.362 : 0 | 0.002 ± 0.005<br>0.895 : 0 |
| $4 \to 5$ | 0.000 ± 0.005<br>0.241 : 0 | 0.001 ± 0.005<br>0.545 : 0 | −0.001 ± 0.005<br>−0.515 : 0 | −0.000 ± 0.005<br>−0.212 : 0 |

Table S12: Simulation results using the data set with 100,000 independent patterns. For each statistic under each scenario we report the value, the 99% Wald confidence interval with radius $2.576 \times \sqrt{4n(L)n(R)/(n(L)+n(R))^3}$, the $Z$-score and the classification. Misclassifications are highlighted in gray. Compared to Table S10 six scenarios are misclassifications.

| | $D_{\text{FO}}$ | $D_{\text{IL}}$ | $D_{\text{FI}}$ | $D_{\text{OL}}$ |
|---|---|---|---|---|
| $1 \to 3$ | 0.073±0.016<br>12.43 : + | 0.049±0.016<br>8.258 : + | 0.029±0.016<br>4.698 : + | 0.002±0.016<br>0.395 : 0 |
| $3 \to 1$ | 0.012±0.016<br>2.018 : 0 | −0.010±0.016<br>−1.630 : 0 | 0.072±0.016<br>12.16 : + | 0.051±0.016<br>8.612 : + |
| $1 \to 4$ | −0.072±0.016<br>−12.22 : − | −0.047±0.016<br>−8.001 : − | −0.001±0.016<br>−0.223 : 0 | 0.025±0.016<br>4.073 : + |
| $4 \to 1$ | −0.027±0.016<br>−4.543 : − | −0.001±0.016<br>−0.115 : 0 | 0.040±0.016<br>6.739 : + | 0.065±0.016<br>11.07 : + |
| $2 \to 3$ | 0.035±0.016<br>5.971 : + | 0.062±0.016<br>10.40 : + | −0.028±0.016<br>−4.664 : − | −0.001±0.016<br>−0.115 : 0 |
| $3 \to 2$ | −0.005±0.016<br>−0.847 : 0 | 0.021±0.016<br>3.448 : + | −0.079±0.016<br>−13.26 : − | −0.054±0.016<br>−9.062 : − |
| $2 \to 4$ | −0.047±0.016<br>−7.948 : − | −0.071±0.016<br>−11.95 : − | −0.001±0.016<br>−0.115 : 0 | −0.025±0.016<br>−4.208 : − |
| $4 \to 2$ | −0.008±0.016<br>−1.256 : 0 | −0.036±0.016<br>−5.943 : − | −0.050±0.016<br>−8.445 : − | −0.077±0.016<br>−13.01 : − |
| $12345 \to 1$ | −0.008±0.016<br>−1.377 : 0 | −0.005±0.016<br>−0.811 : 0 | −0.067±0.015<br>−11.62 : − | −0.064±0.015<br>−11.09 : − |
| $34 \to 2$ | 0.010±0.017<br>1.452 : 0 | 0.009±0.017<br>1.425 : 0 | −0.050±0.014<br>−9.258 : − | −0.050±0.014<br>−9.279 : − |
| $2 \to 34$ | −0.012±0.017<br>−1.878 : 0 | −0.010±0.017<br>−1.611 : 0 | −0.014±0.015<br>−2.471 : 0 | −0.012±0.015<br>−2.240 : 0 |
| $1234 \to 1$ | 0.004±0.016<br>0.649 : 0 | 0.003±0.016<br>0.416 : 0 | −0.026±0.016<br>−4.458 : − | −0.028±0.016<br>−4.682 : − |
| $5 \to 1$ | −0.002±0.016<br>−0.358 : 0 | 0.000±0.016<br>0.006 : 0 | −0.041±0.015<br>−7.013 : − | −0.039±0.015<br>−6.666 : − |
| $1 \to 5$ | 0.010±0.016<br>1.652 : 0 | 0.010±0.016<br>1.664 : 0 | 0.004±0.016<br>0.744 : 0 | 0.005±0.016<br>0.756 : 0 |
| $12345 \to 2$ | 0.004±0.016<br>0.611 : 0 | 0.001±0.016<br>0.147 : 0 | 0.069±0.015<br>11.93 : + | 0.066±0.015<br>11.49 : + |
| $34 \to 1$ | −0.000±0.017<br>−0.013 : 0 | −0.003±0.017<br>−0.382 : 0 | 0.040±0.014<br>7.355 : + | 0.038±0.014<br>7.051 : + |
| $1 \to 34$ | 0.003±0.017<br>0.413 : 0 | −0.001±0.017<br>−0.121 : 0 | 0.009±0.015<br>1.639 : 0 | 0.006±0.015<br>1.177 : 0 |
| $1234 \to 2$ | −0.001±0.016<br>−0.213 : 0 | −0.003±0.016<br>−0.444 : 0 | 0.035±0.016<br>6.034 : + | 0.034±0.016<br>5.811 : + |
| $5 \to 2$ | 0.005±0.016<br>0.845 : 0 | 0.005±0.016<br>0.796 : 0 | 0.046±0.015<br>7.996 : + | 0.046±0.015<br>7.950 : + |
| $2 \to 5$ | −0.003±0.016<br>−0.521 : 0 | −0.002±0.016<br>−0.306 : 0 | −0.001±0.016<br>−0.084 : 0 | 0.001±0.016<br>0.132 : 0 |
| $12345 \to 3$ | −0.069±0.015<br>−11.96 : − | −0.067±0.015<br>−11.68 : − | 0.003±0.016<br>0.565 : 0 | 0.005±0.016<br>0.860 : 0 |
| $12 \to 4$ | −0.047±0.014<br>−8.610 : − | −0.045±0.014<br>−8.317 : − | 0.000±0.018<br>0.033 : 0 | 0.003±0.018<br>0.391 : 0 |
| $4 \to 12$ | −0.008±0.015<br>−1.471 : 0 | −0.008±0.015<br>−1.515 : 0 | 0.002±0.017<br>0.368 : 0 | 0.002±0.017<br>0.317 : 0 |
| $1234 \to 3$ | −0.022±0.016<br>−3.694 : − | −0.024±0.016<br>−4.024 : − | 0.003±0.016<br>0.487 : 0 | 0.001±0.016<br>0.146 : 0 |
| $5 \to 3$ | −0.052±0.015<br>−9.013 : − | −0.053±0.015<br>−9.197 : − | −0.020±0.016<br>−3.344 : − | −0.022±0.016<br>−3.539 : − |
| $3 \to 5$ | 0.011±0.016<br>1.807 : 0 | 0.009±0.016<br>1.555 : 0 | −0.008±0.016<br>−1.414 : 0 | −0.010±0.016<br>−1.665 : 0 |
| $12345 \to 4$ | 0.054±0.015<br>9.247 : + | 0.054±0.015<br>9.351 : + | 0.011±0.016<br>1.810 : 0 | 0.012±0.016<br>1.921 : 0 |
| $12 \to 3$ | 0.045±0.014<br>8.321 : + | 0.043±0.014<br>7.974 : + | 0.002±0.017<br>0.368 : 0 | −0.000±0.017<br>−0.053 : 0 |
| $3 \to 12$ | 0.011±0.015<br>1.988 : 0 | 0.009±0.015<br>1.635 : 0 | −0.003±0.017<br>−0.525 : 0 | −0.006±0.017<br>−0.929 : 0 |
| $1234 \to 4$ | 0.017±0.016<br>2.827 : + | 0.017±0.016<br>2.815 : + | −0.007±0.016<br>−1.112 : 0 | −0.007±0.016<br>−1.124 : 0 |
| $5 \to 4$ | 0.041±0.015<br>7.119 : + | 0.040±0.015<br>6.866 : + | 0.004±0.016<br>0.600 : 0 | 0.002±0.016<br>0.334 : 0 |
| $4 \to 5$ | 0.011±0.016<br>1.840 : 0 | 0.008±0.016<br>1.383 : 0 | 0.001±0.016<br>0.144 : 0 | −0.002±0.016<br>−0.311 : 0 |

Table S13: Simulation results using the data set with 10,000 independent patterns. For each statistic under each scenario we report the value, the 99% Wald confidence interval with radius $2.576 \times \sqrt{4n(L)n(R)/(n(L)+n(R))^3}$, the $Z$-score and the classification. Misclassifications are highlighted in gray. Compared to Table S10, only fourteen scenarios were correctly identified (including the four zero signatures).

| | $D_{\mathrm{FO}}$ | $D_{\mathrm{IL}}$ | $D_{\mathrm{FI}}$ | $D_{\mathrm{OL}}$ |
|---|---|---|---|---|
| $1 \to 3$ | 0.079±0.048<br>4.247 : + | 0.060±0.048<br>3.209 : + | 0.052±0.050<br>2.689 : + | 0.031±0.050<br>1.614 : 0 |
| $3 \to 1$ | 0.019±0.050<br>1.016 : 0 | −0.000±0.050<br>−0.019 : 0 | 0.050±0.048<br>2.704 : + | 0.032±0.048<br>1.704 : 0 |
| $1 \to 4$ | −0.055±0.048<br>−2.985 : − | −0.039±0.048<br>−2.095 : 0 | −0.019±0.051<br>−0.976 : 0 | −0.001±0.051<br>−0.039 : 0 |
| $4 \to 1$ | −0.020±0.050<br>−1.021 : 0 | −0.011±0.050<br>−0.559 : 0 | 0.080±0.049<br>4.239 : + | 0.088±0.049<br>4.691 : + |
| $2 \to 3$ | 0.044±0.049<br>2.335 : 0 | 0.077±0.048<br>4.129 : + | −0.035±0.050<br>−1.804 : 0 | 0.001±0.050<br>0.058 : 0 |
| $3 \to 2$ | 0.005±0.050<br>0.267 : 0 | 0.023±0.050<br>1.182 : 0 | −0.073±0.049<br>−3.864 : − | −0.056±0.049<br>−2.954 : − |
| $2 \to 4$ | −0.067±0.048<br>−3.594 : − | −0.091±0.048<br>−4.928 : − | 0.013±0.050<br>0.694 : 0 | −0.013±0.050<br>−0.694 : 0 |
| $4 \to 2$ | −0.010±0.050<br>−0.537 : 0 | −0.053±0.050<br>−2.764 : − | −0.014±0.048<br>−0.738 : 0 | −0.053±0.048<br>−2.878 : − |
| $12345 \to 1$ | −0.010±0.051<br>−0.526 : 0 | −0.003±0.051<br>−0.136 : 0 | −0.096±0.047<br>−5.223 : − | −0.089±0.048<br>−4.857 : − |
| $34 \to 2$ | 0.035±0.054<br>1.685 : 0 | 0.032±0.054<br>1.560 : 0 | −0.046±0.045<br>−2.674 : − | −0.048±0.045<br>−2.778 : − |
| $2 \to 34$ | −0.032±0.052<br>−1.592 : 0 | −0.017±0.052<br>−0.866 : 0 | −0.020±0.045<br>−1.155 : 0 | −0.009±0.045<br>−0.534 : 0 |
| $1234 \to 1$ | 0.001±0.050<br>0.058 : 0 | 0.006±0.050<br>0.290 : 0 | −0.016±0.048<br>−0.851 : 0 | −0.012±0.048<br>−0.629 : 0 |
| $5 \to 1$ | −0.011±0.050<br>−0.579 : 0 | −0.013±0.050<br>−0.656 : 0 | −0.032±0.048<br>−1.746 : 0 | −0.033±0.048<br>−1.819 : 0 |
| $1 \to 5$ | 0.003±0.049<br>0.133 : 0 | 0.004±0.049<br>0.209 : 0 | −0.022±0.050<br>−1.164 : 0 | −0.021±0.050<br>−1.088 : 0 |
| $12345 \to 2$ | 0.021±0.051<br>1.064 : 0 | 0.008±0.051<br>0.394 : 0 | 0.079±0.046<br>4.401 : + | 0.068±0.046<br>3.793 : + |
| $34 \to 1$ | −0.011±0.054<br>−0.521 : 0 | −0.016±0.054<br>−0.771 : 0 | 0.035±0.045<br>2.047 : 0 | 0.032±0.045<br>1.840 : 0 |
| $1 \to 34$ | 0.004±0.053<br>0.204 : 0 | 0.018±0.053<br>0.897 : 0 | 0.035±0.045<br>2.014 : 0 | 0.045±0.045<br>2.605 : + |
| $1234 \to 2$ | 0.031±0.050<br>1.622 : 0 | 0.038±0.050<br>1.970 : 0 | 0.006±0.048<br>0.332 : 0 | 0.012±0.048<br>0.664 : 0 |
| $5 \to 2$ | −0.008±0.050<br>−0.441 : 0 | −0.017±0.050<br>−0.864 : 0 | 0.062±0.047<br>3.382 : + | 0.054±0.047<br>2.982 : + |
| $2 \to 5$ | −0.027±0.050<br>−1.398 : 0 | −0.023±0.050<br>−1.206 : 0 | −0.002±0.049<br>−0.094 : 0 | 0.002±0.049<br>0.094 : 0 |
| $12345 \to 3$ | −0.066±0.047<br>−3.635 : − | −0.061±0.047<br>−3.380 : − | −0.005±0.050<br>−0.249 : 0 | 0.000±0.050<br>0.019 : 0 |
| $12 \to 4$ | −0.054±0.045<br>−3.155 : − | −0.061±0.045<br>−3.532 : − | 0.004±0.054<br>0.209 : 0 | −0.005±0.054<br>−0.251 : 0 |
| $4 \to 12$ | −0.028±0.046<br>−1.578 : 0 | −0.032±0.046<br>−1.823 : 0 | −0.012±0.051<br>−0.614 : 0 | −0.018±0.051<br>−0.891 : 0 |
| $1234 \to 3$ | −0.052±0.048<br>−2.789 : − | −0.043±0.048<br>−2.306 : 0 | 0.015±0.050<br>0.768 : 0 | 0.024±0.050<br>1.268 : 0 |
| $5 \to 3$ | −0.063±0.047<br>−3.456 : − | −0.065±0.047<br>−3.600 : − | 0.041±0.050<br>2.120 : 0 | 0.038±0.050<br>1.966 : 0 |
| $3 \to 5$ | −0.017±0.049<br>−0.905 : 0 | −0.012±0.049<br>−0.641 : 0 | −0.004±0.049<br>−0.227 : 0 | 0.001±0.049<br>0.038 : 0 |
| $12345 \to 4$ | 0.068±0.048<br>3.686 : + | 0.072±0.048<br>3.906 : + | −0.038±0.051<br>−1.929 : 0 | −0.033±0.051<br>−1.695 : 0 |
| $12 \to 3$ | 0.069±0.045<br>3.994 : + | 0.071±0.045<br>4.132 : + | 0.008±0.055<br>0.398 : 0 | 0.012±0.055<br>0.566 : 0 |
| $3 \to 12$ | 0.038±0.045<br>2.183 : 0 | 0.037±0.045<br>2.113 : 0 | −0.020±0.052<br>−1.015 : 0 | −0.022±0.052<br>−1.095 : 0 |
| $1234 \to 4$ | 0.023±0.049<br>1.222 : 0 | 0.023±0.049<br>1.222 : 0 | 0.010±0.050<br>0.520 : 0 | 0.010±0.050<br>0.520 : 0 |
| $5 \to 4$ | 0.016±0.048<br>0.897 : 0 | 0.020±0.048<br>1.080 : 0 | −0.029±0.050<br>−1.536 : 0 | −0.026±0.050<br>−1.344 : 0 |
| $4 \to 5$ | 0.033±0.050<br>1.738 : 0 | 0.027±0.050<br>1.394 : 0 | −0.018±0.049<br>−0.953 : 0 | −0.024±0.049<br>−1.290 : 0 |

#### Dependence on the synchronization assumption (2)

To show on a concrete level that $D_{\mathrm{FOIL}}$-statistics and Table 2 in the main text rely on the synchronization assumption (2) while Table S1 in Appendix 3 does not, we performed two simulation studies.

First, we attempted (and failed) to create circumstances where making the synchronization assumption (2) despite using ancient samples is justified (other than having an ancient sample from population 5 in tree $S$, populations 3, 4 or 5 in tree $A$ or population 3 in tree $Q$ — in those cases the singleton patterns of the ancient sample are not even used). We expect the singleton patterns to cause less problems when mutations tend to be old; if all the mutations happened after the root the sychronization assumption (2) would in fact be valid. Coalescence events still need to happen before the root, or all statistics tend towards zero. To achieve both ends, we simulate collapsing populations where the coalescence rate slows down with time. The default tree is $S = (((1,2),(3,4)),5)$ with the gene flow event $4 \to 5$, defined using the same parameters as in other simulations except that sample 2 is ancient data from time point 0.25. Five variations of the default have the effective population size scaled by factor $\beta$ in the time interval $[3,\infty]$ and by the factor $1/\beta$ in the time interval $[0,2]$, where $\beta \in \{1, 10, 100, 1000, 10000\}$. A total of 1,000,000 independent allelic patterns were simulated for each graph. The statistics tested were:

- $D_{\mathrm{FI}}$ to demonstrate how $D_{\mathrm{FOIL}}$ is not generally compatible with ancient data. Note that the $D_{\mathrm{FOIL}}$-statistics are blind to the event $4 \to 5$, so sign should be zero unless the ancient sample causes problems.
- ${}_S\Delta_{5+7}$ to demonstrate how those $\Delta$-statistics that contain the binomial statistics ${}_S\Delta_7$ are also sensitive to 2 being an ancient sample. The sign should be zero according to Table 2 in the main paper, unless the ancient sample causes problems.
- ${}_S\Delta_{3-5}$ to show how to fix the issue using only the $\Delta$-statistics that don't rely on the synchronization assumption (2). Recall that ${}_S\Delta_{3-5}$ corresponds to $D_{\mathrm{FI}}$ so the sign should be zero unconditionally.
- ${}_S\Delta_{1-6}$ as a control. According to Table 2 in the main paper, $4 \to 5$ should produce the sign + but slowing coalescence rates carry the risk that all statistics lose power.

In circumstances where the ancient sample is not an issue but we still retain statistical power, these four statistics would produce the signature (0 0 0 +). The observed $Z$-scores are reported below (sign in parenthesis). Power was lost before the ancient sample stopped causing problems, which point was never reached. We cannot conclusively proof that the use of $D_{\mathrm{FOIL}}$ with ancient samples is never excused, but this was our best attempt to find a benevolent scenario.

| | $D_{\mathrm{FI}}$ | ${}_S\Delta_{5+7}$ | ${}_S\Delta_{3-5}$ | ${}_S\Delta_{1-6}$ |
|---|---|---|---|---|
| $\beta = 1$ | −32.715 (−) | 33.242 (+) | 0.383 (0) | 24.560 (+) |
| $\beta = 10$ | −30.404 (−) | 30.407 (+) | 0.402 (0) | 37.013 (+) |
| $\beta = 100$ | −12.185 (−) | 12.174 (+) | 1.246 (0) | 11.773 (+) |
| $\beta = 1000$ | −4.736 (−) | 4.736 (+) | −0.177 (0) | 2.176 (0) |
| $\beta = 10{,}000$ | −2.682 (−) | 2.682 (+) | −1.994 (0) | 1.038 (0) |

Second, continued simulations on the tree $S$ with where lineage 2 is an ancient sample, this time adding all the 32 admixture scenarios in time and simulating 1,000,000 independent allelic patterns on each. We tested both $D_{\mathrm{FOIL}}$ and the $\Delta$-statistics without singletons (Table S1 in Appendix 3) on the data, results are in Tables S14 and S15. As expected, the $D_{\mathrm{FOIL}}$ is heavily biased while the $\Delta$-statistics still operate correctly. In Tables S16 – S18 we demonstrate the futility of three alternative attempts to deal with the violation of the synchronization assumption (2). First S16 we filtered the data so that 5 always carries the ancestral allele, because precisely speaking this is how the original $D_{\mathrm{FOIL}}$ paper defined the statistics. We denote the filtering with the use of lower case letters. Next S17 we leave out the singleton patterns i.e. only use the parsimony informative patterns in statistics $S\Delta^*_{1-2-6}$, $S\Delta^*_{2-1-6}$, $S\Delta^*_{3-4-5}$ and $S\Delta^*_{4-3-5}$. This procedure has no theoretical reason to help (or in fact to work even when the synchronization assumption (2) is valid), but it was recommended for this purpose in Pease & Hahn, 2015. Note that by leaving out not only the singleton patterns, but all the patterns Pease and Hahn call "inverse", we would end up with our own $S\Delta^*_{1-6}$, $S\Delta^*_{1-6}$, $S\Delta^*_{1-6}$ and $S\Delta^*_{1-6}$ and be free of (2). Finally in Table S18 we tested the combination of the two attempts.

Table S14: Simulation with ancient sample, analyzed with $D_{\mathrm{FOIL}}$. For each statistic under each scenario we report the value, the 99% Wald confidence interval with radius $2.576 \times \sqrt{4n(L)n(R)/(n(L)+n(R))^3}$, the $Z$-score and the classification. Misclassifications are highlighted in gray. Compared to Table S10, only eight scenarios were correctly identified. The ancient sample pulls $D_{\mathrm{FI}}$ and $D_{\mathrm{OL}}$ downwards; in seven of the successful scenarios these two statistics were expected to be negative anyways. Also the four zero signature scenarios ($1 \to 5$, $2 \to 5$, $3 \to 5$ and $4 \to 5$) were wrong, as indeed even the null hypothesis is not expected to produce a zero signature.

| | $D_{\mathrm{FO}}$ | $D_{\mathrm{IL}}$ | $D_{\mathrm{FI}}$ | $D_{\mathrm{OL}}$ |
|---|---|---|---|---|
| $1 \to 3$ | 0.071±0.005<br>38.41 : + | 0.048±0.005<br>25.84 : + | −0.041±0.006<br>−21.11 : − | −0.068±0.006<br>−34.42 : − |
| $3 \to 1$ | 0.025±0.005<br>13.00 : + | −0.001±0.005<br>−0.564 : 0 | 0.011±0.005<br>5.643 : + | −0.015±0.005<br>−7.994 : − |
| $1 \to 4$ | −0.069±0.005<br>−37.29 : − | −0.046±0.005<br>−24.71 : − | −0.064±0.006<br>−32.77 : − | −0.038±0.006<br>−19.51 : − |
| $4 \to 1$ | −0.025±0.005<br>−13.07 : − | −0.000±0.005<br>−0.114 : 0 | −0.012±0.005<br>−6.375 : − | 0.013±0.005<br>6.650 : + |
| $2 \to 3$ | 0.048±0.005<br>26.17 : + | 0.072±0.005<br>38.83 : + | −0.090±0.006<br>−45.74 : − | −0.063±0.006<br>−32.33 : − |
| $3 \to 2$ | 0.001±0.005<br>0.421 : 0 | 0.027±0.005<br>13.97 : + | −0.140±0.005<br>−73.38 : − | −0.114±0.005<br>−59.76 : − |
| $2 \to 4$ | −0.046±0.005<br>−25.06 : − | −0.071±0.005<br>−38.14 : − | −0.069±0.006<br>−35.03 : − | −0.096±0.006<br>−48.85 : − |
| $4 \to 2$ | −0.002±0.005<br>−1.219 : 0 | −0.027±0.005<br>−14.47 : − | −0.116±0.005<br>−60.50 : − | −0.141±0.005<br>−73.83 : − |
| $12345 \to 1$ | 0.001±0.005<br>0.277 : 0 | 0.000±0.005<br>0.177 : 0 | −0.132±0.005<br>−70.61 : − | −0.132±0.005<br>−70.71 : − |
| $34 \to 2$ | −0.001±0.006<br>−0.424 : 0 | −0.001±0.006<br>−0.531 : 0 | −0.095±0.005<br>−54.39 : − | −0.095±0.005<br>−54.48 : − |
| $2 \to 34$ | 0.002±0.006<br>1.210 : 0 | 0.002±0.006<br>0.772 : 0 | −0.067±0.005<br>−37.59 : − | −0.067±0.005<br>−37.98 : − |
| $1234 \to 1$ | −0.001±0.005<br>−0.455 : 0 | −0.001±0.005<br>−0.741 : 0 | −0.086±0.005<br>−45.04 : − | −0.086±0.005<br>−45.32 : − |
| $5 \to 1$ | 0.002±0.005<br>1.147 : 0 | 0.002±0.005<br>1.174 : 0 | −0.110±0.005<br>−58.97 : − | −0.110±0.005<br>−58.94 : − |
| $1 \to 5$ | −0.001±0.005<br>−0.357 : 0 | −0.001±0.005<br>−0.541 : 0 | −0.070±0.005<br>−35.88 : − | −0.070±0.005<br>−36.07 : − |
| $12345 \to 2$ | 0.000±0.005<br>0.244 : 0 | 0.001±0.005<br>0.313 : 0 | 0.012±0.005<br>6.559 : + | 0.012±0.005<br>6.626 : + |
| $34 \to 1$ | 0.002±0.006<br>1.043 : 0 | 0.001±0.006<br>0.720 : 0 | −0.006±0.005<br>−3.296 : − | −0.006±0.005<br>−3.569 : − |
| $1 \to 34$ | 0.001±0.006<br>0.310 : 0 | 0.001±0.006<br>0.456 : 0 | −0.036±0.005<br>−20.16 : − | −0.035±0.005<br>−20.03 : − |
| $1234 \to 2$ | −0.003±0.005<br>−1.808 : 0 | −0.003±0.005<br>−1.736 : 0 | −0.038±0.005<br>−20.19 : − | −0.038±0.005<br>−20.11 : − |
| $5 \to 2$ | 0.000±0.005<br>0.112 : 0 | −0.000±0.005<br>−0.044 : 0 | −0.013±0.005<br>−6.761 : − | −0.013±0.005<br>−6.914 : − |
| $2 \to 5$ | 0.001±0.005<br>0.305 : 0 | −0.000±0.005<br>−0.173 : 0 | −0.066±0.005<br>−33.94 : − | −0.067±0.005<br>−34.43 : − |
| $12345 \to 3$ | −0.067±0.005<br>−36.92 : − | −0.066±0.005<br>−36.66 : − | −0.068±0.006<br>−34.01 : − | −0.067±0.006<br>−33.73 : − |
| $12 \to 4$ | −0.044±0.005<br>−26.10 : − | −0.044±0.005<br>−26.02 : − | −0.079±0.006<br>−36.80 : − | −0.079±0.006<br>−36.70 : − |
| $4 \to 12$ | −0.015±0.005<br>−8.688 : − | −0.015±0.005<br>−8.947 : − | −0.072±0.006<br>−35.00 : − | −0.073±0.006<br>−35.31 : − |
| $1234 \to 3$ | −0.023±0.005<br>−12.26 : − | −0.023±0.005<br>−12.71 : − | −0.064±0.006<br>−32.48 : − | −0.065±0.006<br>−32.96 : − |
| $5 \to 3$ | −0.046±0.005<br>−25.29 : − | −0.045±0.005<br>−24.80 : − | −0.063±0.006<br>−32.40 : − | −0.062±0.006<br>−31.87 : − |
| $3 \to 5$ | −0.000±0.005<br>−0.113 : 0 | 0.000±0.005<br>0.041 : 0 | −0.064±0.005<br>−33.06 : − | −0.064±0.005<br>−32.90 : − |
| $12345 \to 4$ | 0.067±0.005<br>37.13 : + | 0.066±0.005<br>36.62 : + | −0.067±0.006<br>−33.95 : − | −0.069±0.006<br>−34.52 : − |
| $12 \to 3$ | 0.041±0.005<br>24.32 : + | 0.041±0.005<br>24.35 : + | −0.078±0.006<br>−36.32 : − | −0.078±0.006<br>−36.29 : − |
| $3 \to 12$ | 0.013±0.005<br>7.517 : + | 0.013±0.005<br>7.479 : + | −0.070±0.006<br>−33.82 : − | −0.070±0.006<br>−33.87 : − |
| $1234 \to 4$ | 0.025±0.005<br>13.73 : + | 0.023±0.005<br>12.64 : + | −0.065±0.006<br>−32.78 : − | −0.067±0.006<br>−33.96 : − |
| $5 \to 4$ | 0.044±0.005<br>24.49 : + | 0.044±0.005<br>24.36 : + | −0.066±0.006<br>−33.81 : − | −0.067±0.006<br>−33.95 : − |
| $4 \to 5$ | −0.001±0.005<br>−0.363 : 0 | −0.000±0.005<br>−0.051 : 0 | −0.064±0.005<br>−32.95 : − | −0.063±0.005<br>−32.63 : − |

Table S15: Simulation with ancient sample, analyzed with the first four Δ-statistics. For each statistic under each scenario we report the value, the 99% Wald confidence interval with radius $2.576 \times \sqrt{4n(L)n(R)/(n(L)+n(R))^3}$, the $Z$-score and the classification. Everything was correctly identified because the synchronization assumption (2) was not needed.

| | $_S\Delta^*_{1-6}$ | $_S\Delta^*_{2-6}$ | $_S\Delta^*_{3-5}$ | $_S\Delta^*_{4-5}$ |
|---|---|---|---|---|
| 1 → 3 | 0.389±0.015<br>63.71 : + | 0.291±0.016<br>45.29 : + | 0.158±0.018<br>23.22 : + | 0.001±0.020<br>0.168 : 0 |
| 3 → 1 | 0.161±0.018<br>23.71 : + | −0.005±0.020<br>−0.649 : 0 | 0.390±0.015<br>63.89 : + | 0.286±0.016<br>44.39 : + |
| 1 → 4 | −0.388±0.015<br>−63.38 : − | −0.289±0.016<br>−44.90 : − | 0.007±0.020<br>0.919 : 0 | 0.161±0.018<br>23.78 : + |
| 4 → 1 | −0.159±0.018<br>−23.39 : − | −0.002±0.020<br>−0.281 : 0 | 0.287±0.016<br>44.59 : + | 0.383±0.015<br>62.96 : + |
| 2 → 3 | 0.288±0.016<br>44.78 : + | 0.384±0.015<br>63.20 : + | −0.147±0.018<br>−21.66 : − | 0.013±0.020<br>1.674 : 0 |
| 3 → 2 | 0.004±0.020<br>0.571 : 0 | 0.168±0.018<br>24.71 : + | −0.387±0.015<br>−63.59 : − | −0.285±0.016<br>−44.23 : − |
| 2 → 4 | −0.292±0.016<br>−45.39 : − | −0.395±0.015<br>−64.63 : − | 0.003±0.020<br>0.457 : 0 | −0.159±0.018<br>−23.48 : − |
| 4 → 2 | 0.002±0.020<br>0.321 : 0 | −0.159±0.018<br>−23.42 : − | −0.285±0.016<br>−44.28 : − | −0.387±0.015<br>−63.36 : − |
| 12345 → 1 | −0.002±0.021<br>−0.222 : 0 | −0.003±0.020<br>−0.416 : 0 | −0.292±0.017<br>−44.95 : − | −0.293±0.017<br>−45.10 : − |
| 34 → 2 | 0.001±0.026<br>0.090 : 0 | −0.001±0.024<br>−0.154 : 0 | −0.263±0.015<br>−44.17 : − | −0.265±0.015<br>−44.35 : − |
| 2 → 34 | 0.020±0.020<br>2.572 : 0 | 0.012±0.019<br>1.628 : 0 | −0.091±0.017<br>−14.25 : − | −0.095±0.017<br>−14.95 : − |
| 1234 → 1 | −0.002±0.021<br>−0.195 : 0 | −0.006±0.020<br>−0.768 : 0 | −0.155±0.018<br>−22.68 : − | −0.159±0.018<br>−23.18 : − |
| 5 → 1 | 0.012±0.021<br>1.560 : 0 | 0.012±0.020<br>1.539 : 0 | −0.607±0.011<br>−121.0 : − | −0.607±0.011<br>−121.0 : − |
| 1 → 5 | 0.011±0.021<br>1.447 : 0 | 0.008±0.021<br>1.050 : 0 | −0.163±0.018<br>−24.36 : − | −0.167±0.018<br>−24.79 : − |
| 12345 → 2 | −0.001±0.020<br>−0.139 : 0 | −0.000±0.021<br>−0.000 : 0 | 0.287±0.017<br>44.28 : + | 0.290±0.017<br>44.53 : + |
| 34 → 1 | 0.012±0.024<br>1.308 : 0 | 0.007±0.026<br>0.666 : 0 | 0.260±0.015<br>43.46 : + | 0.258±0.015<br>43.04 : + |
| 1 → 34 | −0.001±0.019<br>−0.156 : 0 | 0.001±0.020<br>0.114 : 0 | 0.120±0.017<br>18.78 : + | 0.121±0.017<br>19.00 : + |
| 1234 → 2 | 0.001±0.020<br>0.100 : 0 | 0.002±0.021<br>0.252 : 0 | 0.144±0.018<br>21.07 : + | 0.146±0.018<br>21.25 : + |
| 5 → 2 | −0.001±0.020<br>−0.168 : 0 | −0.004±0.021<br>−0.506 : 0 | 0.613±0.011<br>122.7 : + | 0.612±0.011<br>122.4 : + |
| 2 → 5 | 0.007±0.021<br>0.874 : 0 | −0.001±0.021<br>−0.118 : 0 | 0.157±0.018<br>23.30 : + | 0.151±0.018<br>22.42 : + |
| 12345 → 3 | −0.276±0.017<br>−42.24 : − | −0.273±0.017<br>−41.77 : − | 0.005±0.021<br>0.562 : 0 | 0.009±0.021<br>1.106 : 0 |
| 12 → 4 | −0.264±0.015<br>−44.34 : − | −0.264±0.015<br>−44.28 : − | 0.007±0.026<br>0.737 : 0 | 0.008±0.024<br>0.893 : 0 |
| 4 → 12 | −0.104±0.017<br>−16.41 : − | −0.108±0.017<br>−16.92 : − | 0.007±0.020<br>0.918 : 0 | 0.002±0.019<br>0.322 : 0 |
| 1234 → 3 | −0.151±0.018<br>−22.08 : − | −0.158±0.018<br>−22.98 : − | 0.011±0.021<br>1.365 : 0 | 0.003±0.020<br>0.398 : 0 |
| 5 → 3 | −0.614±0.011<br>−122.1 : − | −0.610±0.011<br>−121.4 : − | −0.007±0.021<br>−0.849 : 0 | 0.002±0.020<br>0.228 : 0 |
| 3 → 5 | −0.157±0.017<br>−23.56 : − | −0.155±0.017<br>−23.28 : − | 0.003±0.021<br>0.408 : 0 | 0.006±0.021<br>0.724 : 0 |
| 12345 → 4 | 0.296±0.017<br>45.37 : + | 0.289±0.017<br>44.41 : + | 0.010±0.020<br>1.277 : 0 | 0.001±0.021<br>0.176 : 0 |
| 12 → 3 | 0.262±0.015<br>44.07 : + | 0.263±0.015<br>44.13 : + | 0.007±0.024<br>0.796 : 0 | 0.010±0.026<br>0.965 : 0 |
| 3 → 12 | 0.108±0.017<br>16.86 : + | 0.108±0.017<br>16.81 : + | −0.004±0.019<br>−0.504 : 0 | −0.005±0.020<br>−0.622 : 0 |
| 1234 → 4 | 0.169±0.018<br>24.85 : + | 0.155±0.018<br>22.74 : + | 0.002±0.020<br>0.270 : 0 | −0.016±0.021<br>−2.059 : 0 |
| 5 → 4 | 0.614±0.011<br>122.4 : + | 0.613±0.011<br>122.3 : + | −0.001±0.020<br>−0.076 : 0 | −0.003±0.021<br>−0.377 : 0 |
| 4 → 5 | 0.163±0.018<br>24.33 : + | 0.167±0.018<br>24.91 : + | 0.005±0.021<br>0.701 : 0 | 0.011±0.021<br>1.365 : 0 |

Table S16: Simulation with ancient sample, analyzed with $D_{\mathrm{FOIL}}$ after requiring that sample 5 carries the ancestral allele (denoted with the use of lower case $d$). For each statistic under each scenario we report the value, the 99% Wald confidence interval with radius $2.576 \times \sqrt{4n(L)n(R)/(n(L)+n(R))^3}$, the $Z$-score and the classification. Misclassifications are highlighted in gray. Compared to Table S10, only seven scenarios were correctly identifies. Results resemble Table S14, as 5 typically carries the ancestral allele anyway. Discarding data (patterns where 5 carries the derived allele are also informative) led to reduction of power and the prediction of 12345 → 2 now additionally fails. Arguably without (1) comparisons should be made to an accordingly modified version Table S10, so that successful prediction of 5 → 1 is disqualified as there is no consistent expected behaviour for the event, and 2 → 5 is counted as a success. Regardless, we see that making or not making the outgroup mutation assumption (1) has nothing to do with violations of the synchronization assumption (2).

| | $d_{\mathrm{FO}}$ | $d_{\mathrm{IL}}$ | $d_{\mathrm{FI}}$ | $d_{\mathrm{OL}}$ |
|---|---|---|---|---|
| 1 → 3 | 0.069±0.005<br>36.60 : + | 0.045±0.005<br>24.23 : + | −0.043±0.006<br>−21.80 : − | −0.069±0.006<br>−34.89 : − |
| 3 → 1 | 0.024±0.005<br>12.59 : + | −0.001±0.005<br>−0.470 : 0 | 0.006±0.005<br>2.900 : + | −0.020±0.005<br>−10.26 : − |
| 1 → 4 | −0.066±0.005<br>−35.08 : − | −0.043±0.005<br>−22.72 : − | −0.065±0.006<br>−32.98 : − | −0.040±0.006<br>−19.95 : − |
| 4 → 1 | −0.024±0.005<br>−12.62 : − | −0.000±0.005<br>−0.019 : 0 | −0.016±0.005<br>−8.459 : − | 0.008±0.005<br>4.237 : + |
| 2 → 3 | 0.046±0.005<br>24.48 : + | 0.069±0.005<br>36.86 : + | −0.091±0.006<br>−45.91 : − | −0.065±0.006<br>−32.80 : − |
| 3 → 2 | 0.001±0.005<br>0.330 : 0 | 0.026±0.005<br>13.55 : + | −0.139±0.005<br>−71.75 : − | −0.113±0.005<br>−58.43 : − |
| 2 → 4 | −0.044±0.005<br>−23.38 : − | −0.068±0.005<br>−36.12 : − | −0.070±0.006<br>−35.37 : − | −0.097±0.006<br>−48.83 : − |
| 4 → 2 | −0.002±0.005<br>−1.131 : 0 | −0.027±0.005<br>−14.07 : − | −0.115±0.005<br>−59.15 : − | −0.140±0.005<br>−72.20 : − |
| 12345→1 | 0.001±0.006<br>0.315 : 0 | 0.000±0.006<br>0.206 : 0 | −0.121±0.005<br>−63.58 : − | −0.121±0.005<br>−63.68 : − |
| 34 → 2 | −0.001±0.006<br>−0.374 : 0 | −0.001±0.006<br>−0.637 : 0 | −0.094±0.005<br>−52.87 : − | −0.094±0.005<br>−53.09 : − |
| 2 → 34 | 0.002±0.006<br>1.149 : 0 | 0.002±0.006<br>0.830 : 0 | −0.068±0.005<br>−37.98 : − | −0.069±0.005<br>−38.26 : − |
| 1234→1 | −0.001±0.005<br>−0.532 : 0 | −0.001±0.005<br>−0.586 : 0 | −0.085±0.005<br>−43.91 : − | −0.085±0.005<br>−43.97 : − |
| 5 → 1 | 0.002±0.005<br>1.126 : 0 | 0.002±0.005<br>1.065 : 0 | −0.059±0.006<br>−30.42 : − | −0.059±0.006<br>−30.48 : − |
| 1 → 5 | −0.001±0.005<br>−0.436 : 0 | −0.001±0.005<br>−0.520 : 0 | −0.062±0.006<br>−30.91 : − | −0.062±0.006<br>−31.00 : − |
| 12345→2 | −0.000±0.006<br>−0.243 : 0 | −0.000±0.006<br>−0.064 : 0 | −0.003±0.005<br>−1.746 : 0 | −0.003±0.005<br>−1.571 : 0 |
| 34 → 1 | 0.002±0.006<br>0.849 : 0 | 0.001±0.006<br>0.640 : 0 | −0.011±0.005<br>−6.068 : − | −0.011±0.005<br>−6.246 : − |
| 1 → 34 | 0.001±0.006<br>0.267 : 0 | 0.001±0.006<br>0.485 : 0 | −0.038±0.005<br>−21.09 : − | −0.038±0.005<br>−20.90 : − |
| 1234→2 | −0.003±0.005<br>−1.769 : 0 | −0.003±0.005<br>−1.730 : 0 | −0.043±0.005<br>−22.51 : − | −0.043±0.005<br>−22.47 : − |
| 5 → 2 | 0.000±0.005<br>0.160 : 0 | 0.000±0.005<br>0.063 : 0 | −0.075±0.006<br>−38.44 : − | −0.075±0.006<br>−38.54 : − |
| 2 → 5 | 0.000±0.005<br>0.246 : 0 | −0.000±0.005<br>−0.166 : 0 | −0.080±0.006<br>−40.35 : − | −0.081±0.006<br>−40.78 : − |
| 12345→3 | −0.055±0.005<br>−29.67 : − | −0.054±0.005<br>−29.31 : − | −0.069±0.006<br>−34.45 : − | −0.068±0.006<br>−34.06 : − |
| 12 → 4 | −0.042±0.005<br>−24.21 : − | −0.042±0.005<br>−24.01 : − | −0.081±0.006<br>−37.21 : − | −0.080±0.006<br>−36.95 : − |
| 4 → 12 | −0.014±0.005<br>−7.999 : − | −0.015±0.005<br>−8.367 : − | −0.074±0.006<br>−35.43 : − | −0.075±0.006<br>−35.87 : − |
| 1234→3 | −0.020±0.005<br>−10.68 : − | −0.020±0.005<br>−10.96 : − | −0.066±0.006<br>−33.17 : − | −0.067±0.006<br>−33.47 : − |
| 5 → 3 | 0.007±0.005<br>3.713 : + | 0.008±0.005<br>4.022 : + | −0.065±0.006<br>−32.73 : − | −0.064±0.006<br>−32.41 : − |
| 3 → 5 | 0.011±0.005<br>5.690 : + | 0.011±0.005<br>5.755 : + | −0.066±0.006<br>−33.59 : − | −0.066±0.006<br>−33.52 : − |
| 12345→4 | 0.055±0.005<br>29.84 : + | 0.054±0.005<br>29.34 : + | −0.069±0.006<br>−34.33 : − | −0.070±0.006<br>−34.87 : − |
| 12 → 3 | 0.038±0.005<br>21.97 : + | 0.038±0.005<br>21.96 : + | −0.080±0.006<br>−36.76 : − | −0.080±0.006<br>−36.78 : − |
| 3 → 12 | 0.012±0.005<br>6.948 : + | 0.012±0.005<br>6.944 : + | −0.071±0.006<br>−34.15 : − | −0.071±0.006<br>−34.15 : − |
| 1234→4 | 0.022±0.005<br>11.97 : + | 0.021±0.005<br>10.98 : + | −0.067±0.006<br>−33.29 : − | −0.069±0.006<br>−34.36 : − |
| 5 → 4 | −0.009±0.005<br>−4.738 : − | −0.009±0.005<br>−4.923 : − | −0.068±0.006<br>−34.04 : − | −0.068±0.006<br>−34.23 : − |
| 4 → 5 | −0.011±0.005<br>−5.811 : − | −0.011±0.005<br>−5.592 : − | −0.066±0.006<br>−33.46 : − | −0.065±0.006<br>−33.24 : − |

Table S17: Simulation with ancient sample, analyzed with the statistics ${}_S\Delta^*_{1-2-6}$, ${}_S\Delta^*_{2-1-6}$, ${}_S\Delta^*_{3-4-5}$ and ${}_S\Delta^*_{4-3-5}$ — in other words $D_{\mathrm{FOIL}}$-statistics without the singleton patterns, a solution suggested by Pease and Hahn for situations where (2) doesn't hold. For each statistic under each scenario we report the value, the 99% Wald confidence interval with radius $2.576 \times \sqrt{4n(L)n(R)/(n(L)+n(R))^3}$, the $Z$-score and the classification. Misclassifications are highlighted in gray. Compared to Table S10, twenty scenarios were correctly identified. Clearly discarding the singleton patterns was a step to the right direction, as the problems no longer appear to be about the synchronization assumption (2). Because the binomial Δ-statistics are not always consistent under various gene flow events (see Figure 3 in main text), neither are their arbitrary combinations. To attain consistent behaviour, one must also discard some parsimony informative patterns to end up with our ${}_S\Delta^*_{1-6}$, ${}_S\Delta^*_{2-6}$, ${}_S\Delta^*_{3-5}$ and ${}_S\Delta^*_{4-5}$ that are proven to work in Appendix 2.

| | ${}_S\Delta^*_{1-2-6}$ | ${}_S\Delta^*_{2-1-6}$ | ${}_S\Delta^*_{3-4-5}$ | ${}_S\Delta^*_{4-3-5}$ |
|---|---|---|---|---|
| 1 → 3 | 0.314±0.014<br>55.64 : + | 0.098±0.015<br>17.39 : + | 0.150±0.017<br>23.83 : + | −0.118±0.017<br>−18.82 : − |
| 3 → 1 | 0.154±0.016<br>24.57 : + | −0.128±0.017<br>−20.33 : − | 0.319±0.014<br>56.53 : + | 0.091±0.015<br>16.18 : + |
| 1 → 4 | −0.314±0.014<br>−55.59 : − | −0.098±0.015<br>−17.34 : − | −0.113±0.017<br>−17.97 : − | 0.154±0.016<br>24.49 : + |
| 4 → 1 | −0.147±0.016<br>−23.41 : − | 0.121±0.017<br>19.33 : + | 0.095±0.015<br>16.83 : + | 0.311±0.014<br>55.25 : + |
| 2 → 3 | 0.092±0.015<br>16.36 : + | 0.307±0.014<br>54.75 : + | −0.138±0.016<br>−22.00 : − | 0.130±0.016<br>20.78 : + |
| 3 → 2 | −0.122±0.017<br>−19.52 : − | 0.158±0.016<br>25.22 : + | −0.317±0.014<br>−56.24 : − | −0.090±0.015<br>−16.04 : − |
| 2 → 4 | −0.096±0.015<br>−16.99 : − | −0.319±0.014<br>−56.70 : − | 0.123±0.016<br>19.67 : + | −0.153±0.016<br>−24.44 : − |
| 4 → 2 | 0.123±0.017<br>19.57 : + | −0.152±0.016<br>−24.26 : − | −0.091±0.015<br>−16.15 : − | −0.313±0.014<br>−55.54 : − |
| 12345 → 1 | −0.000±0.019<br>−0.028 : 0 | −0.003±0.019<br>−0.396 : 0 | −0.234±0.016<br>−38.63 : − | −0.235±0.016<br>−38.94 : − |
| 34 → 2 | 0.000±0.022<br>0.057 : 0 | −0.003±0.022<br>−0.368 : 0 | −0.198±0.015<br>−35.40 : − | −0.200±0.015<br>−35.70 : − |
| 2 → 34 | 0.016±0.017<br>2.538 : 0 | 0.007±0.017<br>1.117 : 0 | −0.034±0.015<br>−5.873 : − | −0.041±0.015<br>−7.137 : − |
| 1234 → 1 | 0.001±0.019<br>0.168 : 0 | −0.006±0.019<br>−0.883 : 0 | −0.120±0.017<br>−19.04 : − | −0.126±0.017<br>−19.99 : − |
| 5 → 1 | 0.009±0.019<br>1.261 : 0 | 0.010±0.019<br>1.359 : 0 | −0.537±0.011<br>−111.8 : − | −0.536±0.011<br>−111.8 : − |
| 1 → 5 | 0.006±0.019<br>0.823 : 0 | 0.001±0.019<br>0.122 : 0 | −0.135±0.017<br>−21.49 : − | −0.139±0.017<br>−22.10 : − |
| 12345 → 2 | −0.005±0.019<br>−0.686 : 0 | −0.003±0.019<br>−0.431 : 0 | 0.232±0.016<br>38.29 : + | 0.233±0.016<br>38.50 : + |
| 34 → 1 | 0.010±0.022<br>1.260 : 0 | −0.000±0.022<br>−0.016 : 0 | 0.197±0.015<br>34.98 : + | 0.192±0.015<br>34.10 : + |
| 1 → 34 | −0.001±0.017<br>−0.116 : 0 | 0.002±0.017<br>0.360 : 0 | 0.054±0.015<br>9.412 : + | 0.057±0.015<br>9.838 : + |
| 1234 → 2 | −0.002±0.019<br>−0.319 : 0 | −0.000±0.019<br>−0.050 : 0 | 0.110±0.017<br>17.39 : + | 0.112±0.017<br>17.64 : + |
| 5 → 2 | 0.000±0.019<br>0.056 : 0 | −0.004±0.019<br>−0.522 : 0 | 0.543±0.011<br>113.5 : + | 0.541±0.011<br>113.1 : + |
| 2 → 5 | 0.010±0.019<br>1.429 : 0 | −0.003±0.019<br>−0.386 : 0 | 0.135±0.017<br>21.50 : + | 0.125±0.017<br>19.90 : + |
| 12345 → 3 | −0.223±0.016<br>−36.63 : − | −0.218±0.016<br>−35.75 : − | 0.001±0.019<br>0.207 : 0 | 0.009±0.019<br>1.237 : 0 |
| 12 → 4 | −0.203±0.015<br>−36.32 : − | −0.202±0.015<br>−36.05 : − | 0.001±0.022<br>0.107 : 0 | 0.004±0.022<br>0.502 : 0 |
| 4 → 12 | −0.044±0.015<br>−7.692 : − | −0.049±0.015<br>−8.554 : − | 0.004±0.017<br>0.648 : 0 | −0.002±0.017<br>−0.324 : 0 |
| 1234 → 3 | −0.114±0.017<br>−18.02 : − | −0.124±0.017<br>−19.58 : − | 0.009±0.019<br>1.273 : 0 | −0.003±0.019<br>−0.448 : 0 |
| 5 → 3 | −0.542±0.011<br>−112.9 : − | −0.536±0.011<br>−111.6 : − | −0.008±0.019<br>−1.119 : 0 | 0.006±0.019<br>0.795 : 0 |
| 3 → 5 | −0.134±0.016<br>−21.40 : − | −0.130±0.016<br>−20.89 : − | 0.002±0.019<br>0.264 : 0 | 0.006±0.019<br>0.849 : 0 |
| 12345 → 4 | 0.239±0.016<br>39.41 : + | 0.229±0.016<br>37.68 : + | 0.013±0.019<br>1.764 : 0 | −0.002±0.019<br>−0.270 : 0 |
| 12 → 3 | 0.199±0.015<br>35.50 : + | 0.199±0.015<br>35.60 : + | 0.005±0.022<br>0.615 : 0 | 0.006±0.022<br>0.763 : 0 |
| 3 → 12 | 0.047±0.015<br>8.184 : + | 0.046±0.015<br>8.057 : + | −0.004±0.017<br>−0.591 : 0 | −0.005±0.017<br>−0.733 : 0 |
| 1234 → 4 | 0.137±0.017<br>21.82 : + | 0.114±0.017<br>18.08 : + | 0.013±0.019<br>1.810 : 0 | −0.017±0.019<br>−2.373 : 0 |
| 5 → 4 | 0.543±0.011<br>113.2 : + | 0.541±0.011<br>112.9 : + | 0.000±0.019<br>0.014 : 0 | −0.004±0.019<br>−0.508 : 0 |
| 4 → 5 | 0.133±0.017<br>21.23 : + | 0.140±0.016<br>22.27 : + | 0.002±0.019<br>0.229 : 0 | 0.010±0.019<br>1.418 : 0 |

Table S18: Simulation with ancient sample, analyzed with statistics that combine the attempts at Tables S16 and S17, in other words, statistics ${}_S\delta^*_{1-2-6}$, ${}_S\delta^*_{2-1-6}$, ${}_S\delta^*_{3-4-5}$ and ${}_S\delta^*_{4-3-5}$ where the lower case $\delta$ denotes that we have filtered out patterns where 5 carries the derived allele. For each statistic under each scenario we report the value, the 99% Wald confidence interval with radius $2.576 \times \sqrt{4n(L)n(R)/(n(L)+n(R))^3}$, the $Z$-score and the classification. Misclassifications are highlighted in gray. Compared to Table S10, twenty two scenarios were correctly identified. There was no theoretical reason to think this method would work, the example was included only for completeness' sake.

| | ${}_S\delta^*_{1-2-6}$ | ${}_S\delta^*_{2-1-6}$ | ${}_S\delta^*_{3-4-5}$ | ${}_S\delta^*_{4-3-5}$ |
|---|---|---|---|---|
| 1 → 3 | 0.341±0.015<br>56.03 : + | 0.097±0.016<br>15.98 : + | 0.166±0.018<br>24.43 : + | −0.139±0.018<br>−20.45 : − |
| 3 → 1 | 0.171±0.018<br>25.15 : + | −0.144±0.018<br>−21.15 : − | 0.339±0.015<br>55.76 : + | 0.087±0.016<br>14.36 : + |
| 1 → 4 | −0.335±0.015<br>−55.17 : − | −0.093±0.016<br>−15.23 : − | −0.125±0.018<br>−18.34 : − | 0.180±0.018<br>26.45 : + |
| 4 → 1 | −0.162±0.018<br>−23.84 : − | 0.140±0.018<br>20.67 : + | 0.092±0.016<br>15.11 : + | 0.333±0.015<br>54.88 : + |
| 2 → 3 | 0.090±0.016<br>14.88 : + | 0.331±0.015<br>54.77 : + | −0.154±0.018<br>−22.62 : − | 0.151±0.018<br>22.18 : + |
| 3 → 2 | −0.142±0.018<br>−20.89 : − | 0.177±0.018<br>26.03 : + | −0.338±0.015<br>−55.70 : − | −0.084±0.016<br>−13.92 : − |
| 2 → 4 | −0.093±0.016<br>−15.42 : − | −0.342±0.015<br>−56.51 : − | 0.143±0.018<br>21.11 : + | −0.168±0.018<br>−24.85 : − |
| 4 → 2 | 0.141±0.018<br>20.74 : + | −0.170±0.018<br>−25.07 : − | −0.088±0.016<br>−14.51 : − | −0.336±0.015<br>−55.39 : − |
| 12345→1 | 0.003±0.021<br>0.406 : 0 | −0.000±0.021<br>−0.031 : 0 | −0.263±0.017<br>−39.10 : − | −0.265±0.017<br>−39.47 : − |
| 34 → 2 | 0.008±0.024<br>0.823 : 0 | −0.003±0.024<br>−0.329 : 0 | −0.218±0.016<br>−35.04 : − | −0.223±0.016<br>−35.83 : − |
| 2 → 34 | 0.016±0.019<br>2.235 : 0 | 0.008±0.019<br>1.118 : 0 | −0.039±0.017<br>−6.100 : − | −0.045±0.017<br>−7.109 : − |
| 1234→1 | −0.004±0.020<br>−0.543 : 0 | −0.006±0.020<br>−0.760 : 0 | −0.128±0.018<br>−18.16 : − | −0.129±0.018<br>−18.36 : − |
| 5 → 1 | 0.013±0.021<br>1.704 : 0 | 0.011±0.021<br>1.455 : 0 | −0.297±0.017<br>−45.31 : − | −0.299±0.017<br>−45.52 : − |
| 1 → 5 | 0.004±0.021<br>0.479 : 0 | 0.001±0.021<br>0.128 : 0 | −0.015±0.021<br>−1.848 : 0 | −0.017±0.021<br>−2.195 : 0 |
| 12345→2 | −0.011±0.021<br>−1.337 : 0 | −0.005±0.021<br>−0.613 : 0 | 0.255±0.017<br>37.98 : + | 0.259±0.017<br>38.59 : + |
| 34 → 1 | 0.009±0.024<br>1.022 : 0 | 0.001±0.024<br>0.110 : 0 | 0.216±0.016<br>34.53 : + | 0.212±0.016<br>33.91 : + |
| 1 → 34 | −0.001±0.019<br>−0.155 : 0 | 0.004±0.019<br>0.605 : 0 | 0.058±0.017<br>9.155 : + | 0.063±0.017<br>9.844 : + |
| 1234→2 | −0.001±0.021<br>−0.157 : 0 | −0.000±0.021<br>−0.000 : 0 | 0.112±0.019<br>15.93 : + | 0.113±0.019<br>16.07 : + |
| 5 → 2 | 0.002±0.021<br>0.235 : 0 | −0.001±0.021<br>−0.156 : 0 | 0.300±0.017<br>46.04 : + | 0.298±0.017<br>45.72 : + |
| 2 → 5 | 0.010±0.021<br>1.228 : 0 | −0.004±0.021<br>−0.494 : 0 | 0.021±0.021<br>2.688 : + | 0.008±0.021<br>0.995 : 0 |
| 12345→3 | −0.256±0.017<br>−37.83 : − | −0.247±0.017<br>−36.52 : − | −0.002±0.021<br>−0.262 : 0 | 0.010±0.021<br>1.280 : 0 |
| 12 → 4 | −0.225±0.016<br>−36.21 : − | −0.221±0.016<br>−35.47 : − | −0.003±0.024<br>−0.331 : 0 | 0.007±0.024<br>0.771 : 0 |
| 4 → 12 | −0.042±0.017<br>−6.641 : − | −0.051±0.017<br>−7.981 : − | 0.007±0.019<br>0.993 : 0 | −0.004±0.019<br>−0.497 : 0 |
| 1234→3 | −0.118±0.018<br>−16.78 : − | −0.125±0.018<br>−17.83 : − | 0.002±0.021<br>0.225 : 0 | −0.007±0.021<br>−0.940 : 0 |
| 5 → 3 | −0.303±0.017<br>−46.19 : − | −0.296±0.017<br>−45.12 : − | −0.011±0.021<br>−1.376 : 0 | −0.001±0.021<br>−0.101 : 0 |
| 3 → 5 | −0.010±0.020<br>−1.242 : 0 | −0.008±0.020<br>−0.978 : 0 | −0.001±0.021<br>−0.127 : 0 | 0.001±0.021<br>0.143 : 0 |
| 12345→4 | 0.264±0.017<br>39.33 : + | 0.252±0.017<br>37.51 : + | 0.017±0.021<br>2.129 : 0 | −0.000±0.021<br>−0.016 : 0 |
| 12 → 3 | 0.218±0.016<br>35.05 : + | 0.218±0.016<br>35.01 : + | 0.005±0.024<br>0.502 : 0 | 0.004±0.024<br>0.447 : 0 |
| 3 → 12 | 0.049±0.017<br>7.684 : + | 0.049±0.017<br>7.672 : + | −0.002±0.019<br>−0.303 : 0 | −0.002±0.019<br>−0.317 : 0 |
| 1234→4 | 0.140±0.018<br>20.08 : + | 0.114±0.018<br>16.36 : + | 0.013±0.021<br>1.614 : 0 | −0.020±0.021<br>−2.535 : 0 |
| 5 → 4 | 0.305±0.017<br>46.49 : + | 0.300±0.017<br>45.84 : + | 0.002±0.021<br>0.265 : 0 | −0.004±0.021<br>−0.500 : 0 |
| 4 → 5 | 0.020±0.021<br>2.521 : 0 | 0.027±0.021<br>3.411 : + | 0.002±0.021<br>0.239 : 0 | 0.009±0.021<br>1.148 : 0 |

### Appendix 6: Brown bear – polar bear gene flow, an empirical example

To reevaluate previous findings on brown bear – polar bear gene flow direction, we used the same Variant Call Format (VCF) file as in our 2022 work (Lan et al., 2022), adding to it the data (a newly constructed genotype VCF) for the Bruno sample (obtaining its genome data from the Short Read Archive of the National Center for Biotechnology Information Sequence Reads Archive SRS8777210. Data processing, variant calling, and VCF construction and filtering (using the same parameters as for Dataset02) followed the methodology outlined previously (Lan et al., 2022). The final VCF was converted into a variant-major additive component file (–recode A-transpose) using PLINK 2.0 (Chang et al., 2015).

First, we wanted to ensure the consistency of phylogenetic signals between the newly generated dataset and the original dataset. We therefore investigated the relationship between the two ancient polar bear samples, APB and Bruno, and their relationships to modern polar and brown bears. We removed any missing data with vcftools version 0.1.16 (`--max-missing 1.0`), and individuals with depth of coverage less than 8x, as described in Lan et al., 2022. We performed Principal Component Analysis with smartPCA from the eigensoft package version 7.2 (Patterson, Price, & Reich, 2006; Price et al., 2006) and generated a drift maximum likelihood tree using TreeMix version 1.13 (Pickrell & Pritchard, 2012).

The PCA recovered all the brown and polar bear populations reported by Lan et al., 2022. The ancient polar bear samples, Bruno and APB, were placed very close to each other and both samples fell outside the tight cluster of modern polar bear diversity, demonstrating slightly closer relationship to brown bears than modern polar bears did (Figures S1 and S2). To explore the relationships of the ancient polar bears with modern polar bears in greater detail, a new data set was created removing all brown bears (Figure S3). Both PC1 and PC2 separated Bruno and APB from one another and from modern polar bears, the latter still comprising a tight cluster. Along PC1, Bruno was placed closer to modern polar bears than was APB, while along PC2, APB placed closer to modern polar bears than Bruno did.

The initial TreeMix tree, without migration edges, successfully represented all the groupings reported in Lan et al., 2022. In this tree, Bruno was positioned as sister to modern polar bears, with APB as the sister taxon to the clade encompassing Bruno and modern polar bears (Figure S4). The tree with a migration edge set to 0 captures 99.83% of the variance, with only 0.17% unaccounted for. Subsequently, we incrementally added migration edges, up to a total of 5. The first migration event observed was from ABC bears to the ancestor of polar bears, a signal also reflected in the $\Delta$-statistics. Additional migration edges suggested gene flow between various brown bear populations and polar bears, as well as from the ancestral polar bear branch to mainland North American (YB) brown bears.

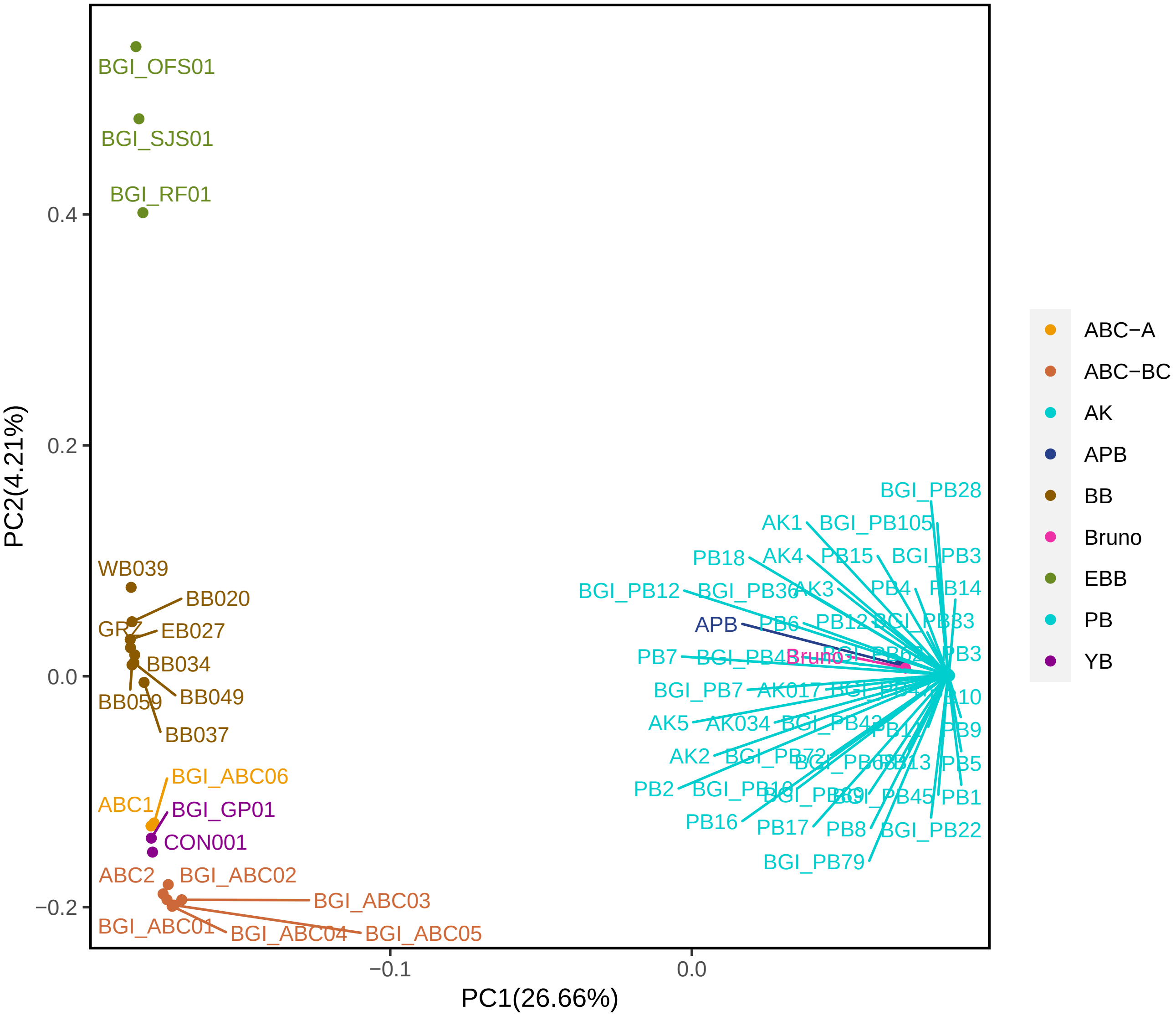


Figure S1: Principal component analysis with sample labels.

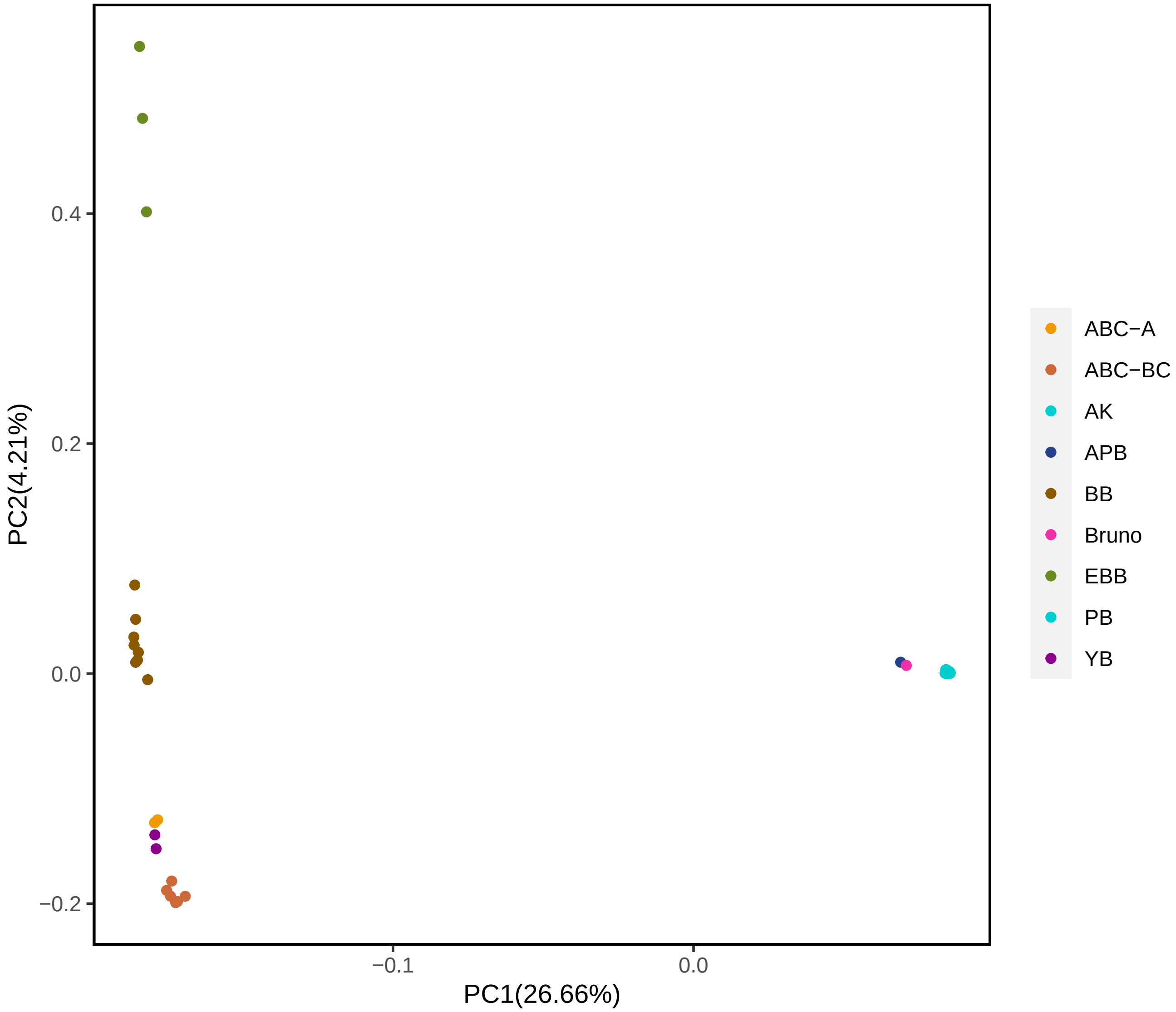


Figure S2: Principal component analysis without sample labels.

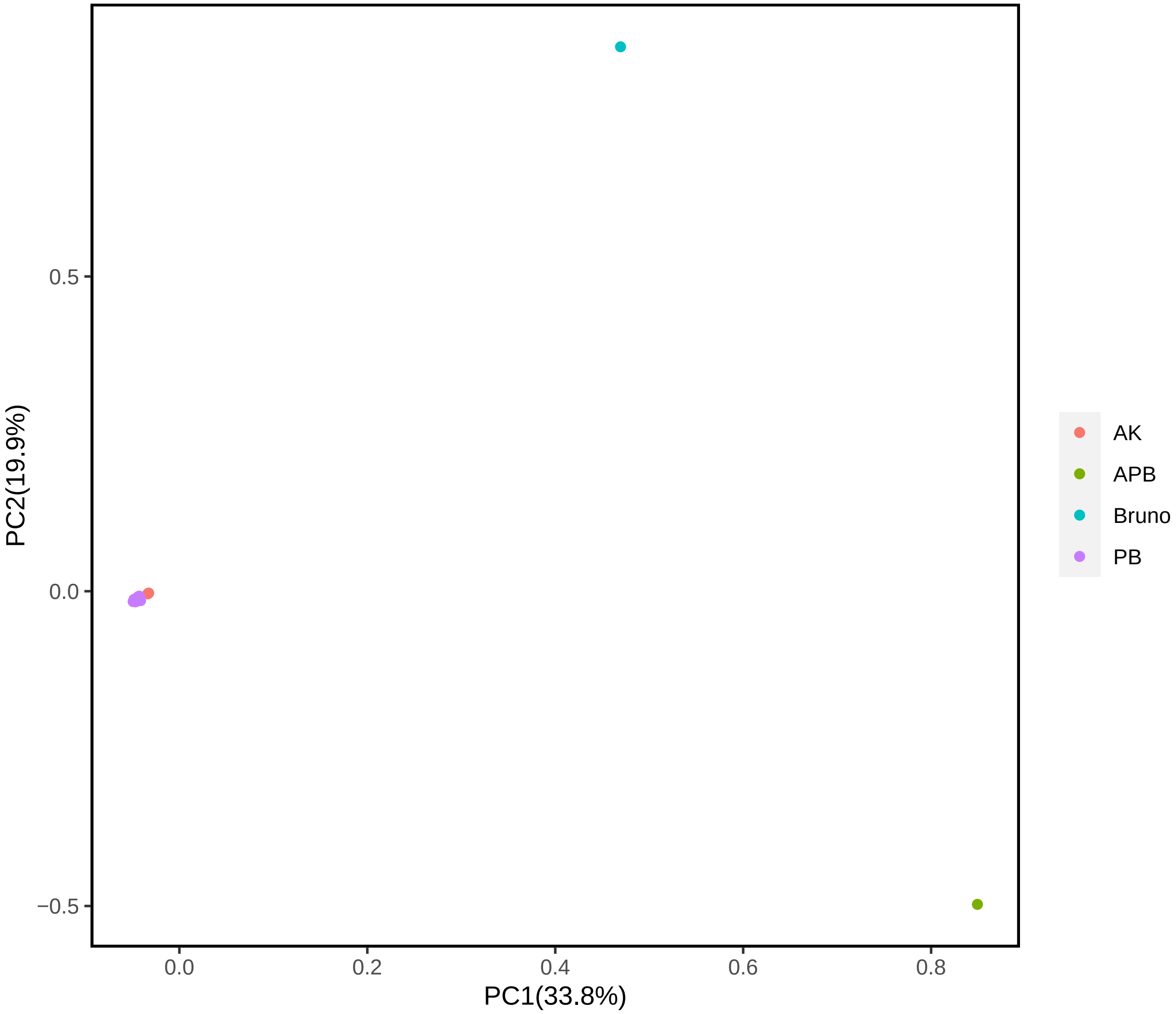


Figure S3: Principal component analysis of polar bears without sample labels.

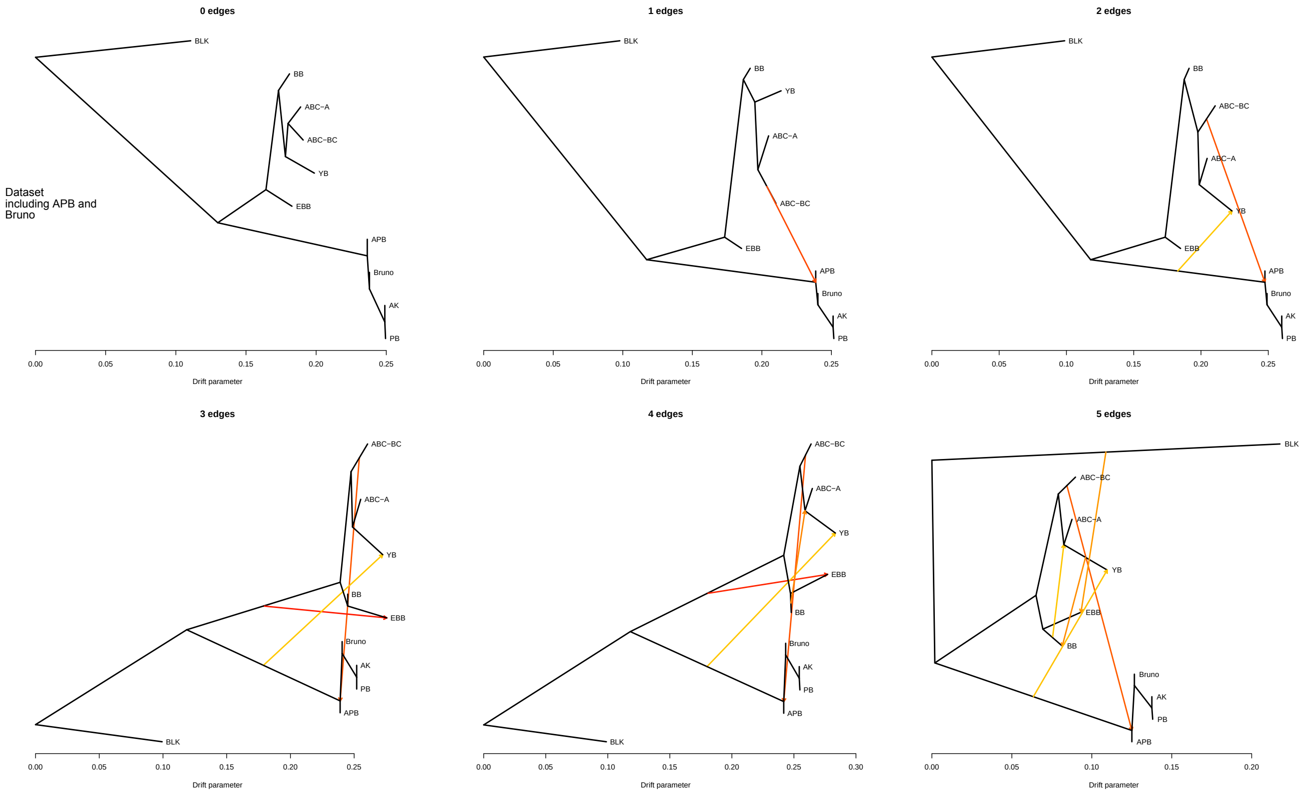


Figure S4: TreeMix analysis.

#### Δ-statistic analyses based on other population sets

To compare our Δ-statistics results that use the ancient Svalbard polar bear sample, APB (main text Figure 4), with combinations using the ancient Alaskan polar bear sample, "Bruno", we ran all symmetric, asymmetric, and quasisymmetric combinations replacing APB alternately with Bruno (Figure S5). The similarity of number of windows in each scenario regardless of the ancient sample used in the combination is noteworthy. Hence, both Bruno and APB provide evidence of ancient bidirectional gene flow between brown and polar bears.

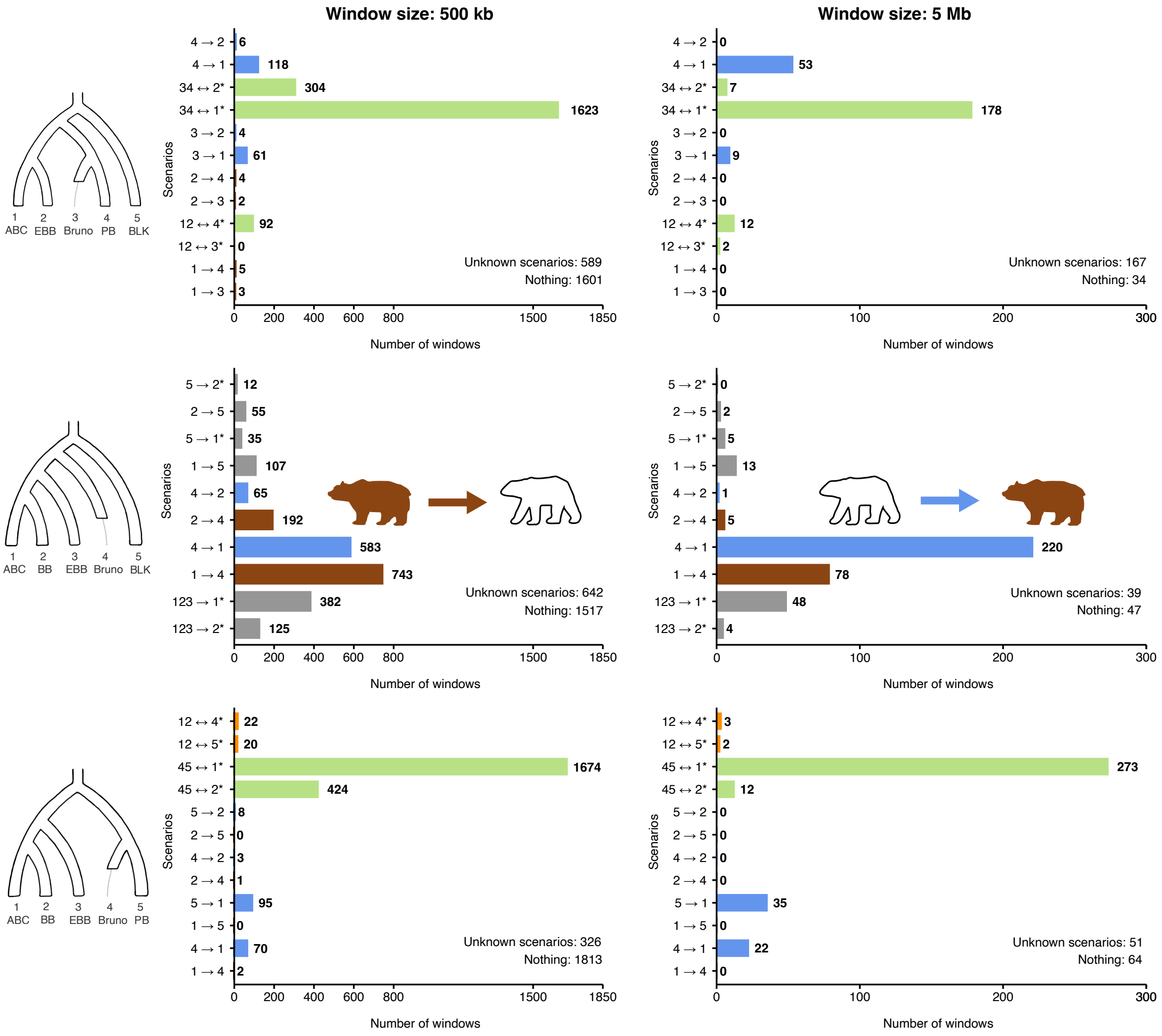


Figure S5: Bar plots of various Δ-statistic analyses of brown bear – polar bear gene flow incorporating a modern and the Alaskan ancient polar bear (Bruno) in place of the Svalbard sample APB (main text Figure 4). Because Bruno is an ancient sample, we cannot make the synchronization assumption (2), so we therefore exclude singletons and use the collapsed Tables S1–S3 in Appendix 3. The asterisk means there are additional gene flow events consistent with the signature to the representative written, for example 34 ↔ 2* in tree S stands for either 12345 → 1, 34 ↔ 2, 1234 → 1 or 1 ↔ 5, according to Table S1 in Appendix 3. The colors of the bars signify the following: Brown — the signature supports only gene flow from brown bears into polar bears; Blue — the signature supports only gene flow from polar bears into brown bears; Orange — the signature supports gene flow in either direction between brown bears and polar bears; Gray — the signature supports some gene flow event that has nothing to do with the brown bear – polar bear question; Green — the signature supports gene flow of either or neither direction between brown bears and polar bears. The windows with zero signature are counted under "Nothing", and windows with non-zero signature are discordant with any row in Tables S1–S3 in Appendix 3 under "Unknown scenarios". See the main text for description of the populations and interpretation.

In Figures S6 and S7 we tried out the full $\Delta$-statistic analysis making the synchronization assumption (2) despite using the ancient samples APB and Bruno, respectively. In tree $S$ where sample 3 is ancient, the binomial statistic ${}_S\Delta_8$ is biased downwards. This bias deteriorates signatures like $(0\,0+\,+\,|\,-\,0\,0\,0)$ (corresponding to $34 \leftrightarrow 1^*$) and $(-\,0+\,+\,|\,-\,+\,0\,0)$ (corresponding to $r \rightarrow 1$) towards $(0\,0+\,+\,|\,-\,-\,0\,-)$ and $(-\,0+\,+\,|\,-\,-\,0\,-)$ (corresponding to nothing). This is clearly visible in the "Unknown scenarios" bin ultimately swallowing up everything. A similar effect happens in tree $Q$ where sample 4 is ancient, as the binomial statistic ${}_Q\Delta_8$ is also biased downwards. In turn, tree $A$ where sample 4 is ancient is not affected by the violation of the synchronization assumption (2). This is because only samples 1 and 2 contribute singleton patterns to the $\Delta$-statistics for $A$, not 4. This is not to say that tree $A$ is inherently more robust than $S$ or $Q$; an ancient sample in populations 1 or 2 would cause an issue.

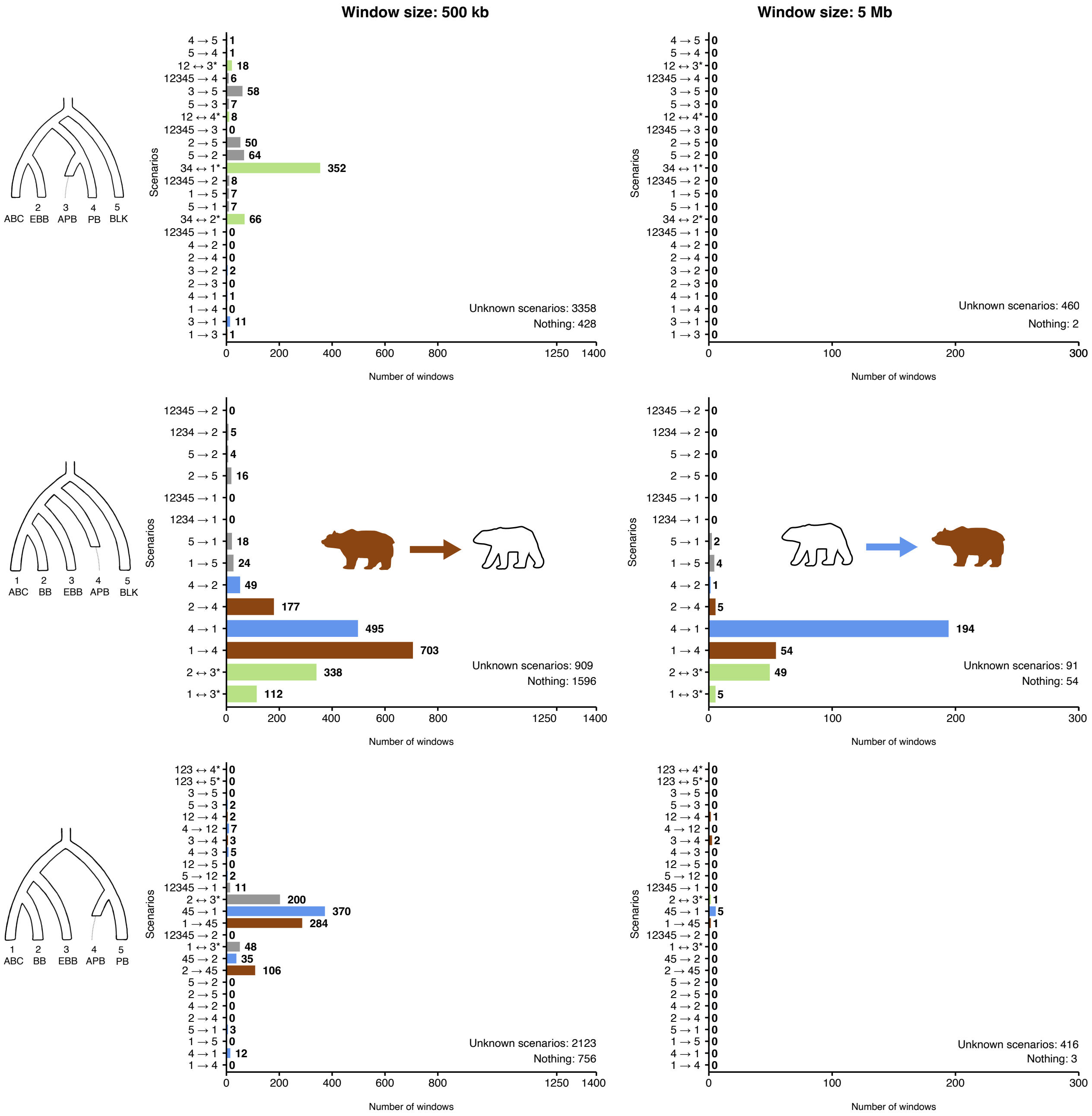


Figure S6: Bar plots of various Δ-statistics analyses concerning brown bear – polar bear gene flow. We commit to the synchronization assumption (2) despite using an ancient sample (APB) and therefore we use the complete Tables 2—4 in the main text; the results are biased. The asterisk means there are additional gene flow events consistent with the signature to the representative written, for example $34 \leftrightarrow 2^*$ in tree $S$ stands for $1234 \to 1$ or $34 \leftrightarrow 2$, according to Table 2 in the main text. The colours of the bars are as in Figure S5. The windows with zero signature are counted under "Nothing", and windows with non-zero signature are discordant with any row in Tables 2–4 in the main text under "Unknown scenarios".

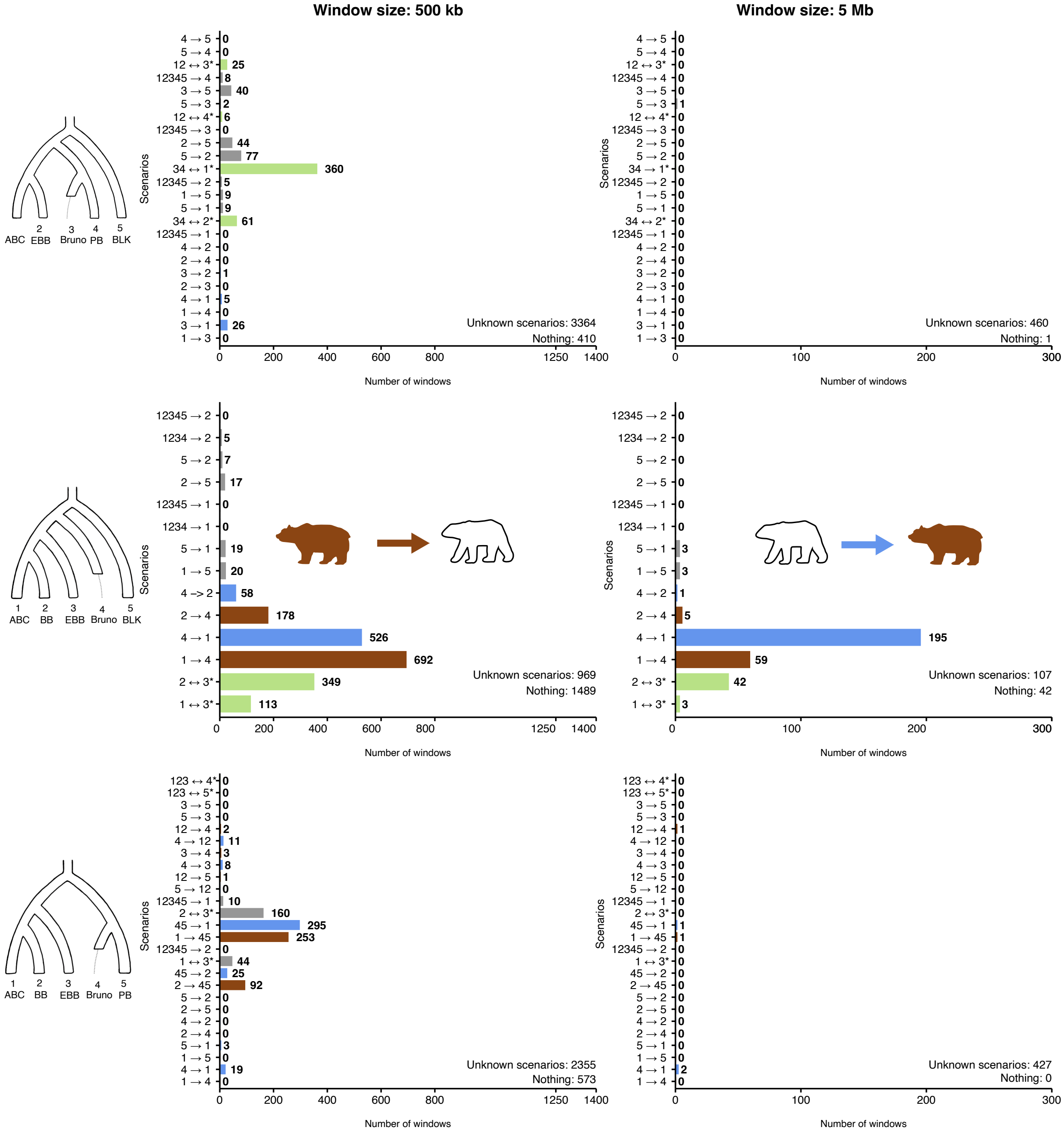


Figure S7: Bar plots of various Δ-statistics analyses concerning brown bear – polar bear gene flow. We commit to the synchronization assumption (2) despite using an ancient sample (Bruno) and therefore we use the complete Tables 2—4 in the main text; the results are biased. The asterisk means there are additional gene flow events consistent with the signature to the representative written, for example $34 \leftrightarrow 2^*$ in tree $S$ stands for $1234 \rightarrow 1$ or $34 \leftrightarrow 2$, according to Table 2 in the main text. The colours of the bars are as in Figure S5. The windows with zero signature are counted under "Nothing", and windows with non-zero signature are discordant with any row in Tables 2–4 in the main text under "Unknown scenarios".

#### Comparisons of empirical results from different methods

A recent article by Wang et al., 2022 used $D_{\text{FOIL}}$ (Pease & Hahn, 2015) on tree $S$ to study brown bear – polar bear gene flow, with results interpreted to principally support unidirectional gene flow from polar bears into brown bears. Here we provide a comparison of the Wang et al. results to our own based on $D_{\text{FOIL}}$- and $\Delta$-statistics.

The results of Wang et al. are visualized in their Figure 3, comparable to the upper panels (tree $S$) of our Figures 4 and 5 in the main text; a bar plot counting the number of 200 kb windows favouring different admixture scenarios according to $D_{\text{FOIL}}$ repeated using different individuals to represent the populations. The populations used (for the gray box plots) are North American brown bears (NA), European brown bears (E), the 100,000 year old ancient polar bear specimen Bruno from Alaska, modern polar bears (MP) and black bears (B). For the sake of comparison, we assume these are roughly equivalent to our samples ABC, EBB, APB, PB and BLK, respectively (our Figure 4 repeated using Bruno in place of APB can be found as Figure S5 in Appendix 6). Note that Figure 3 of Wang et al. is split into two alternative scenarios (panels $a$ and $b$ versus panels $c$ and $d$) according to whether the North American and European brown bears split before or after Bruno and modern polar bears did. Such differentiation is caused by an unnecessary artefact in $D_{\text{FOIL}}$. The original $D_{\text{FOIL}}$ article (Pease & Hahn, 2015) assumes that populations 1 and 2 split closer to the present time than 3 and 4 did, and that consequently gene flow events $12 \leftrightarrow 3$ and $12 \leftrightarrow 4$ are considered a possibility while $34 \leftrightarrow 1$ and $34 \leftrightarrow 2$ are not. While it is true that in this setting, the events $1 \rightarrow 34$ and $2 \rightarrow 34$ are impossible, events $34 \rightarrow 1$ and $34 \rightarrow 2$ can still be mediated by a ghost population. Furthermore, as we show in Appendix 5, gene flow events $12345 \rightarrow 2$, $1234 \rightarrow 2$ and $5 \rightarrow 2$ produce the same $D_{\text{FOIL}}$ signature $(0\,0 + +)$ as $34 \leftrightarrow 1$, and $12345 \rightarrow 1$, $1234 \rightarrow 1$, and $5 \rightarrow 1$ the same $D_{\text{FOIL}}$ signature $(0\,0 - -)$ as $34 \leftrightarrow 2$. We therefore hold the view that it is not appropriate to consider any of the signatures $(0\,0 - -)$, $(0\,0 + +)$, $(- - 0\,0)$ or $(+ + 0\,0)$ as impossible *a priori*, especially when they are ecountered in practice. Indeed, the difference between panels $b$ and $d$ in Figure 3 of Wang et al. is simply $b$ omitting the signatures consistent with gene flow between ancient polar bears and either of the two brown bear samples, and $d$ omitting the signatures consistent with gene flow between ancient brown bears and either of the two polar bear samples — the number of windows consistent with the null hypothesis or inconsistent with all the scenarios shown is not reported. Hence, for the remainder of this comparison we view panels $b$ and $d$ as two incomplete subsets of the complete result, and all statements about Figure 3 will be translated onto the tree $S = (((1 = \text{NA}, 2 = \text{E}), (3 = \text{Bruno/MP}, 4 = \text{MP})), 5 = \text{B})$.

One striking feature of the Figure 3 of Wang et al. is the abundance (note the logarithmic scale) of windows supporting the bidirectional gene flow events $12 \leftrightarrow 3$, $12 \leftrightarrow 4$, $34 \leftrightarrow 1$ and $34 \leftrightarrow 2$. We believe that the singleton patterns used by $D_{\text{FOIL}}$ introduce extensive stochasticity on the short windows of 200 kb in length. Sometimes population NA may happen to have more private mutations than population E, resulting in $D_{\text{FOIL}}$ signature $(0\,0 - -)$, which looks like the event $34 \leftrightarrow 2$, yet equally often is the other way around with $(0\,0 + +)$ , which looks like event $34 \leftrightarrow 1$. Therefore all four signatures are overrepresented. We can also observe the effect of switching Bruno with an extant polar bear sample (the ochre box plots in their Figure 3). With Bruno, many more windows were compatible with $12 \leftrightarrow 3$ than with $12 \leftrightarrow 4$, and with two modern samples the signals become virtually indistinguishable. The same can be observed in our $D_{\text{FOIL}}$ runs in Figures S8 and S9: $12 \leftrightarrow 4$ has very little support and $12 \leftrightarrow 3$ has plenty. Using $\Delta$-statistics without singletons we found no support at all for $12 \leftrightarrow 3$ and only some for $12 \leftrightarrow 4$ (Figure 4 in the main text), and no support for either when using two modern polar bear samples (the second polar bear being AK34, Figure 5 in the main text). This difference in results (whether the ancient — $D_{\text{FOIL}}$ — or the modern — $\Delta$ without the use of singletons — polar bear shows more signs of admixture with all the brown bears) demonstrates how the synchronization assumption (2) can affect the results when singletons are included in the analyses. The $D_{\text{FOIL}}$ method is unsuitable for ancient data; the private mutations in MP compared to Bruno were enough to reverse the result with excess of false signature $(+ + 0\,0)$. On the other hand, stochasticity or the presence of the ancient sample cannot explain why Wang et al. observe more windows compatible with $34 \leftrightarrow 1$ than with $34 \leftrightarrow 2$ (American brown bear shows more signs of admixture with all the polar bears than European brown bears do), because the difference persists when using two modern polar bear samples, the ochre box plots. This pattern is also observed with our Figures 4 and 5, S19 and S20. This is the strongest signal in the data, ambiguous to the direction of gene flow regarding brown bear – polar bear admixture, and even compatible with other explanations ignored by $D_{\text{FOIL}}$ such as cave bear or black bear admixture into European brown bears; see expanded $D_{\text{FOIL}}$ predictions in Table S4 in Appendix 5.

The other gene flow events with noteworthy support in Wang et al.'s results and in our Figures S8 and S9 were $3 \rightarrow 1$ and $4 \rightarrow 1$, present both with Bruno and with two modern polar bear samples. These signals can reasonably be interpreted as modern gene flow from polar bears into North American brown bears. In the top panels of Figure 4 in the main text, we observe clear evidence of $4 \rightarrow 1$ and some evidence of $3 \rightarrow 1$. In the top panels of Figure 5 , contrasting Wang et al., these events have practically no support, likely because the threshold for "modernity" is pulled towards the present day when APB is replaced by AK. Stochasticity in small windows may again play a role, potentially breaking the strong older signature $(0\,0++)$ into equal amounts of $(+\,0++)$ (that looks like $3 \to 1$), $(-\,0++)$ (that looks like $4 \to 1$), $(0+++)$ and $(0-++)$ (which are discarded).

Evidence for recent polar bear $\to$ brown bear gene flow should be accounted for, but not surmount the strongest signal (the signature $(0\,0++)$ consistent with $34 \leftrightarrow 1$) that has several plausible explanations. Research using mitochondrial DNA (Lindqvist et al., 2010) or $f$-statistic fitting (Lan et al., 2022) conflict with an interpretation of predominantly unidirectional polar bear $\to$ brown bear gene flow, and admixture between European brown bears and both cave bears (Barlow et al., 2018) and black bears (Kumar et al., 2017) is also recognized in the literature — as well as supported by our earlier work using $f$-statistic fitting (Lan et al., 2022). The problem is that this particular use of $D_{\mathrm{FOIL}}$ with tree $S$ is theoretically incapable of giving unambiguous support in favour of ancient gene flow from brown bears into polar bears, so long as the admixture is old enough to affect both polar bear samples. The difficulty of finding an unadmixed sister sample for polar bears also challenges earlier attempts ((Cahill et al., 2015); see SI Appendix section S18 in (Lan et al., 2022)) to study the brown bear – polar bear gene flow using the $\hat{f}$-statistic (Green et al., 2010). Ancient samples could be a solution, but it turns out that Bruno and the Svalbard specimen APB are also admixed. This is why switching to the asymmetric tree $A$ is the key, where an unadmixed sister sample to polar bears is no longer required.

In summary, while the $D_{\mathrm{FOIL}}$ results of Wang et al. are largely consistent with ours, the interpretation of mostly unidirectional gene flow from polar bear into brown bear is no longer consistent with $\Delta$-statics on trees $A$ and $Q$. This is partly due to methodological limitations with $D_{\mathrm{FOIL}}$ that we intended to overcome in the present work. There are some signs (events whose prevalence comes in pairs of equal support) that the window size 200 kb introduces stochastical noise, in particular because the $D_{\mathrm{FOIL}}$ statistics employ singleton patterns. The singleton patterns also render ancient data unusable, visibly in the pattern where the proposed polar bear $\to$ brown bear gene flow appears stronger for Bruno than for a modern polar bear sample, even though that sample has been dead for 100,000 years. This order is reversed with $\Delta$-statistics that are applicable for ancient data: support is strong for $4 \to 1$ but weak for $3 \to 1$ in Figure 4 top panels.

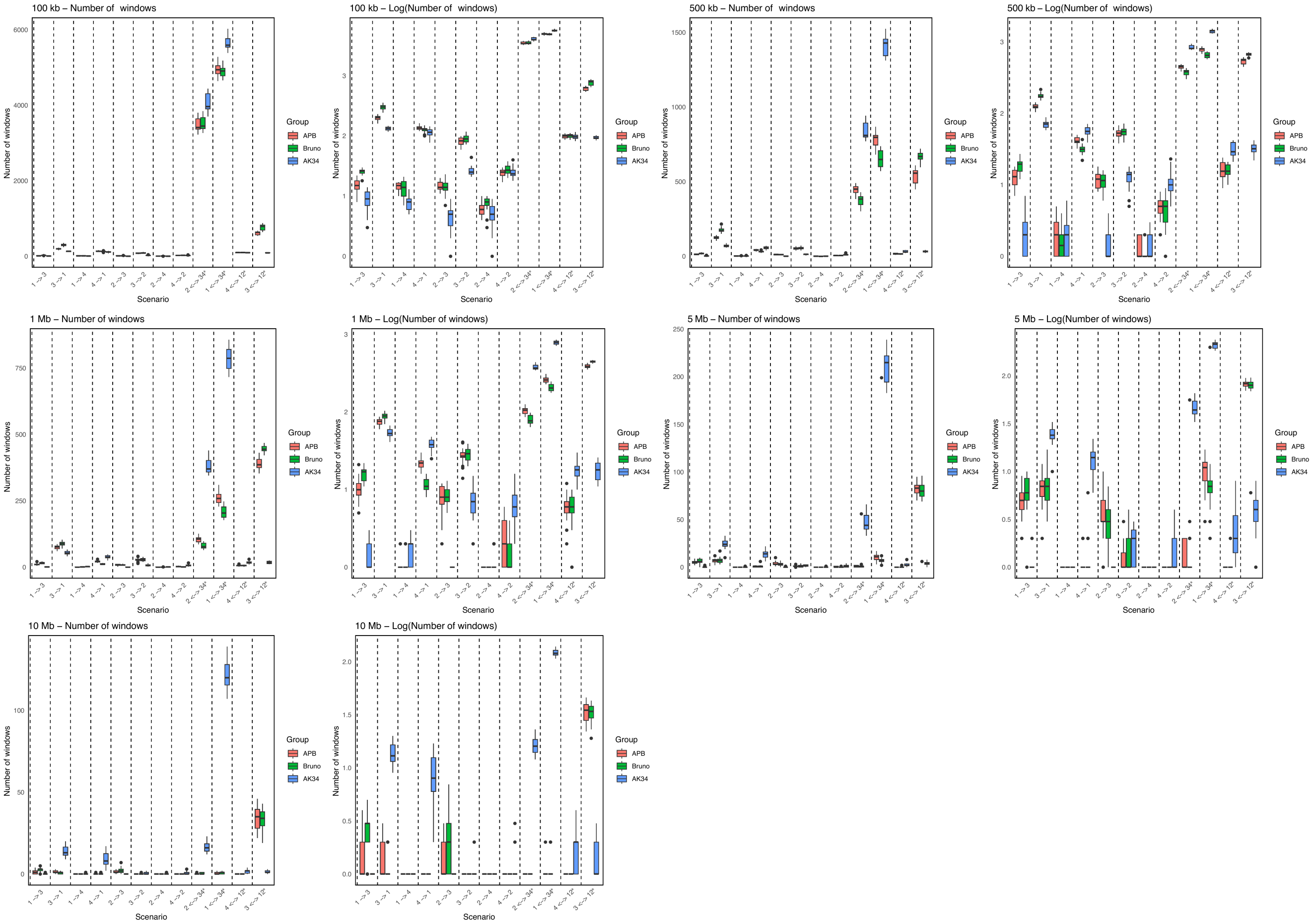


Figure S8: Expanded $D_{\mathrm{FOIL}}$ results from the combination $(((1 = \mathrm{ABC}, 2 = \mathrm{EBB}), (3 = \mathrm{X}, 4 = \mathrm{PB})), 5 = \mathrm{BLK})$, where X represents either APB (coral), Bruno (green), or a modern polar bear AK34 (blue), and ABC, EBB, and PB are represented by three different individuals in independent runs of $D_{\mathrm{FOIL}}$. Analyses were performed using different windows sizes: a) 100 kb, b) 500 kb, c) 1 Mb, d) 5 Mb, and e) 10 Mb. The x-axis represents the various scenarios based on Table S4 in Appendix 5 and the y-axis represents either (left panel) the number of windows or (right panel) $log_{10}$(number of windows).

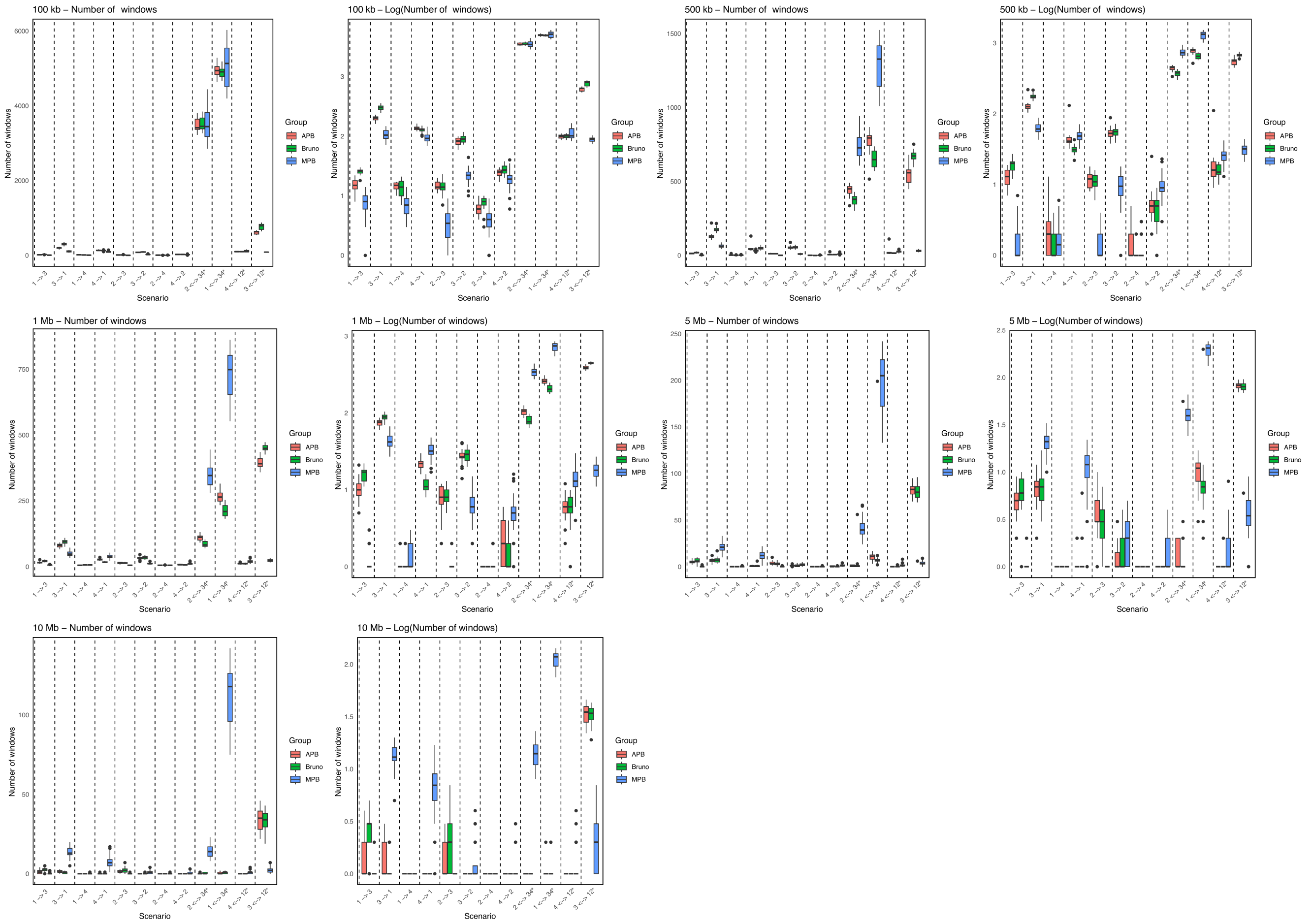


Figure S9: Expanded $D_{\mathrm{FOIL}}$ results from the combination $(((1 = \mathrm{ABC}, 2 = \mathrm{EBB}), (3 = \mathrm{X}, 4 = \mathrm{PB})), 5 = \mathrm{BLK})$, where X represents either APB (coral), Bruno (green), or three different polar bears from the AK individuals (MPB;blue), and ABC, EBB, and PB are represented by three different individuals in independent runs of $D_{\mathrm{FOIL}}$. Analyses were performed using different windows sizes: a) 100 kb, b) 500 kb, c) 1 Mb, d) 5 Mb, and e) 10 Mb. The x-axis represents the various scenarios based on Table S4 in Appendix 3 and the y-axis represents either (left panel) the number of windows or (right panel) $\log_{10}$(number of windows).
